# Comprehensive dissection of GPCR signaling using a NanoBiT-based platform

**DOI:** 10.64898/2026.07.09.737439

**Authors:** Ayaki Saito, So Yamaguchi, Riko Suzuki, Masataka Yanagawa, Ryoji Kise, Asuka Inoue

**Affiliations:** Graduate School of Pharmaceutical Sciences, Kyoto University, Kyoto, 606-8501, Japan; Graduate School of Pharmaceutical Sciences, Tohoku University, Sendai, 980-8578, Japan; Graduate School of Science, Kyoto University, Kyoto, 606-8501, Japan; International Institute for Integrative Sleep Medicine (WPI-IIIS), University of Tsukuba, Tsukuba, 303-0005, Japan

## Abstract

G-protein-coupled receptors (GPCRs) signal through multiple heterotrimeric G proteins, β-arrestins, GPCR kinases (GRKs), and downstream effectors, whose combinatorial interactions shape cellular responses. These events are typically measured with separate assay formats that each capture only part of the network, making comparison across signaling layers difficult. Here, we consolidate a broad set of previously reported GPCR signaling interactions into a single NanoBiT split-luciferase framework, allowing multiple layers of signal transduction to be examined side by side in living cells. We show that rational sensor engineering, in particular the positioning of NanoBiT fragments and targeted modification of the tagged proteins, is essential for detecting transient protein–protein interactions. The framework implements such strategies to measure G-protein dissociation across diverse Gβ–Gγ subtype combinations, β-arrestin recruitment, conformational activation and trafficking, and GRK recruitment. Additionally, we developed new sensors to monitor interactions between transducers and their effector, as well as inter-effector interactions across diverse Gα, β-arrestin, adenylyl cyclase, PLCβ, Rho, and RhoGEF subtypes. It also enables real-time monitoring of the difficult-to-access Gα_12/13_–RhoGEF–RhoA pathway. Together, these assays provide a unified NanoBiT readout for systematic, side-by-side dissection of GPCR signaling.

## Introduction

G-protein–coupled receptors (GPCRs) constitute the largest family of transmembrane receptors and are targeted by approximately one-third of FDA-approved drugs. Upon binding extracellular ligands, GPCRs transduce signals into the intracellular environment through multiple layers of signaling components, including GPCR transducers (G-proteins, β-arrestins, and GRKs) and downstream effectors.

The diverse biological outcomes elicited by GPCRs are primarily mediated by a variety of signal transducers. Heterotrimeric G proteins comprise 16 distinct Gα subtypes, which are categorized into four subfamilies: G_s/olf_, G_i/o_, G_q/11_, and G_12/13_. Upon activation, these G-proteins engage distinct downstream effectors, leading to diverse biological outcomes. Specifically, G_s/olf_ proteins activate adenylyl cyclase, whereas G_i/o_ proteins inhibit it; G_q/11_ proteins activate PLCβ to induce Ca^2+^ mobilization and G_12/13_ proteins induce the RhoA–ROCK pathway via activation of RhoGEF [1]. β- arrestin1 and β-arrestin2 represent another class of GPCR signal transducers and serve as key regulators of receptor trafficking as well as scaffolds for MAPK signaling [2].

Although GPCR signaling has traditionally been depicted as a set of discrete, parallel pathways, recent studies have revealed extensive crosstalk and combinatorial engagement among receptors and multiple transducer classes, complicating the interpretation of downstream signaling assays. One common example is the ability of relatively promiscuous GPCRs to couple to multiple Gα-protein subfamilies [3]. In such cases, the specific Gα-protein coupling profile can be difficult to infer from second-messenger assays alone. Another example is the µ-opioid receptor (MOR), whose recruitment of β-arrestins has been linked to adverse effects of opioid drugs. Although MOR primarily couples to Gα_i/o_ proteins, its β-arrestin recruitment can depend on a more specific signaling pathway involving Gβ_5_ and GRK3 interactions [4]. These examples show that downstream responses do not always reflect which transducers or signaling molecules are involved. Therefore, assays that directly measure proximal GPCR signaling events, such as GPCR–transducer interactions and transducer rearrangements, are needed to define GPCR signaling pathways more clearly.

The need to directly measure proximal GPCR signaling has driven extensive efforts to develop assays that selectively and quantitatively monitor early signaling events downstream of receptor activation. For example, TRUPATH established an open-source BRET-based G-protein biosensor platform by optimizing Gαβγ sensors for 14 Gα subtypes, enabling systematic interrogation of GPCR coupling to heterotrimeric G proteins with single-pathway resolution [5]. In this platform, receptor-catalyzed G-protein activation is detected through changes in BRET associated with heterotrimer rearrangement or dissociation. Effector membrane translocation assay, EMTA, provides another powerful strategy for monitoring proximal GPCR signaling by detecting the recruitment or redistribution of effector domains following activation of specific Gα subfamilies [6]. In addition, G-protein activation can be assessed by detecting the release of free Gβγ dimers through their interaction with GRK-derived BRET biosensor components, thereby providing an indirect readout of heterotrimer dissociation [7]. In addition, bystander BRET configurations have enabled G-protein activation to be monitored without directly modifying the Gα subunit, allowing the use of native Gα proteins [8].

Several assays have also been developed to monitor β-arrestin recruitment to activated GPCRs. One widely used example is the DiscoverX PathHunter assay, which detects GPCR–β-arrestin interactions through β-galactosidase enzyme-fragment complementation [9]. The effector membrane translocation assay (EMTA), described above, has also been adapted to monitor β-arrestin recruitment, although its application to β-arrestin profiling has remained relatively limited. Other platforms, including PRESTO-Tango and Tango-Trio, have enabled systematic profiling of GPCR–β-arrestin interactions across large receptor panels [10,11]. However, these Tango-based platforms rely on endpoint transcriptional readouts and therefore provide limited information about the kinetics and reversibility of β-arrestin recruitment.

Beyond direct interactions with GPCRs, β-arrestin trafficking and conformational changes have also been monitored using resonance energy transfer (RET)-based approaches. β-Arrestin translocation to early endosomes has been assessed using bystander BRET assays that detect its proximity to the FYVE domain of Endofin [12,13]. In parallel, intramolecular FRET- and BRET-based sensors have been developed to monitor receptor-induced conformational rearrangements within β- arrestins without relying on conformation-selective intrabodies, thereby enabling a more direct analysis of β-arrestin structural dynamics [14,15].

Compared with other major GPCR transducers, approaches for monitoring GRK recruitment to GPCRs remain particularly limited. Traditionally, GRK engagement has been inferred from receptor phosphorylation using in vitro kinase assays or phosphosite-specific antibodies [16,17]. In the context of PPI assays, β-arrestin recruitment has been used as an indicator for GRK functions [18]. However, these methods do not reflect GRK direct association with GPCRs.

Recent studies have developed RET-based approaches to monitor interactions between transducers and downstream effectors, as well as interactions among downstream signaling components. For example, FRET- and BRET-based assays have been used to evaluate the association of Gα_q_ with PLCβ [19,20]. BRET has also been applied to detect interactions between Gγ_2_ and adenylyl cyclase [21]. In addition, NanoBRET has been used to monitor the interaction between RAF and MEK1, two downstream components of the MAPK cascade that is frequently engaged by GPCR signaling [22].

Although the above-mentioned approaches have provided valuable insights into individual steps of GPCR signal transduction, interactions across the broader GPCR signaling network have not yet been systematically examined within a single experimental framework. In particular, a unified platform capable of evaluating GPCR interactions with multiple transducer classes—including G proteins, β-arrestins, and GRKs—together with transducer–effector and downstream signaling-component interactions, remains lacking.

Here, we systematically evaluated GPCR signal transduction across multiple signaling layers using a NanoBiT-based assay platform. This platform includes assays for G-protein dissociation, β-arrestin recruitment and subcellular trafficking, GRK recruitment, transducer–effector interactions, and interactions among downstream effector components.

## Results

### Concise guide to design NanoBiT constructs

As with many biosensor platforms, the development of NanoBiT-based assays requires careful probe design to maximize stimulus-dependent changes in signal. Unlike resonance energy transfer-based assays, NanoBiT signals are not constrained by the relative dipole orientation of the sensor components. Instead, signal generation depends on the proximity and productive complementation of two luciferase fragments with low intrinsic affinity for one another. Accordingly, the performance of a NanoBiT assay is determined largely by the positions at which the fragments are fused or inserted into the proteins of interest. Below, we outline practical design principles for optimizing NanoBiT-based assays.

### Maximizing Assay Sensitivity

Preserving the native function of the proteins of interest is important for avoiding artificial signaling outcomes and minimizing the need for additional validation. The N- and C-termini generally provide suitable sites for luciferase-fragment fusion, whereas internal loops may be considered when both termini are functionally constrained. These configurations should first be evaluated to identify a design that preserves protein expression, localization, and signaling activity.

In some cases, signal sensitivity can be further enhanced by introducing point mutations that intentionally modulate protein behavior while preserving core functionality. For instance, mutation of Gγ within its lipid modification site enhances the detectability of G-protein dissociation signals by facilitating its dissociation from the plasma membrane while largely retaining its plasma membrane localization under basal conditions via the heterotrimeric complex. Another example is the use of kinase-deficient GRK mutants [18], which can prolong GRK association with activated GPCRs by preventing receptor phosphorylation and the subsequent progression of the signaling cycle. Similarly, mini-G proteins are modified to stabilize an active, receptor-bound conformation, thereby increasing the lifetime of GPCR–G-protein complexes (Nehmé et al, 2017). Together, these strategies illustrate how targeted modification of signaling proteins can improve detection of transient interactions without eliminating their core biological functionality.

Protein truncations also serve as an alternative method for improving assay sensitivity. A representative example is β-arrestin, truncation of the autoinhibitory C-terminal tail stabilizes a pre-activated state (Kim et al, 2013), substantially enhancing recruitment signals.

To maximize stimulus-dependent signal changes, luciferase fragments should be positioned at sites that are brought into close proximity upon protein–protein interaction. Structural information can therefore provide valuable guidance for identifying suitable fusion or insertion sites. In heterotrimeric G-protein complexes, for example, the N-terminus of the Gγ subunit is located near the α-helical domain of the Gα subunit [25], making these regions suitable for NanoBiT fragment placement. The optimal insertion site may also vary depending on the interaction being monitored.

For Gα_q_ protein, the NanoBiT insertion site used to detect heterotrimer dissociation differs slightly from that used to monitor its interaction with phospholipase Cβ (PLCβ). In the PLCβ interaction assay, the fragment is inserted at residue 123 within the αB–αC loop, because this region lies closer to the Gα_q_–PLCβ interaction interface [26]. The design and application of these constructs are discussed in greater detail in later sections.

NanoBiT assays are well suited for detecting ligand-induced changes in luminescence relative to the prestimulation baseline. In non-transfected cells, basal luminescence is typically maintained at approximately 10^⁴^ relative luminescence units. Although absolute signal intensity varies with assay configuration and experimental conditions, coexpression of the two NanoBiT-tagged proteins generally produces a readily detectable basal signal, often more than tenfold higher than that observed in non-transfected cells. Against this baseline, changes in luminescence as small as approximately 2% can be reproducibly detected. NanoBiT assays are less well suited, however, for quantitatively comparing basal protein–protein interactions in the absence of ligand stimulation. This limitation arises because the luminescence signal is influenced not only by the extent of interaction but also by the expression levels, localization, and complementation efficiency of the two tagged proteins. Because the abundance of each NanoBiT-tagged component cannot be independently determined from a single luminescence signal, differences in basal luminescence among constructs or experimental conditions cannot necessarily be attributed to differences in basal interaction. In contrast, BRET-based assays provide separate donor and acceptor signals, allowing the acceptor-to-donor expression ratio to be estimated and taken into account when interpreting basal interaction measurements.

Optimizing NanoBiT sensor expression levels is critical for achieving high robustness and reproducibility. As in other protein–protein interaction-based assays, lower expression of luciferase fragment sensors can sometimes increase the apparent fold response by reducing basal complementation or background luminescence. However, decreased expression also lowers absolute luminescence intensity, thereby increasing susceptibility to stochastic noise and technical variability. Therefore, fold change alone is insufficient to define optimal assay conditions. When assay responses are unstable or highly variable, we recommend titrating the plasmid amounts and evaluating assay performance using the Z’ factor. The Z′ factor incorporates both the separation between positive and negative control signals and the variability within each condition, thereby providing a quantitative measure of assay robustness. Our direct β-arrestin recruitment assay was validated using Z′ factor analysis. We applied this approach to optimize the direct β-arrestin recruitment assay by testing one-fifth, the original, and fivefold the original plasmid amounts. As expected, basal luminescence increased with increasing plasmid amount (Fig. 1A). The β-arrestin recruitment response was significantly greater at the original and fivefold plasmid amounts than at the one-fifth condition; however, this increase was accompanied by greater variability, as reflected by an increased coefficient of variation (CV) (Fig. 1B–D). Overall, the highest Z′ factor was obtained with the original plasmid amount, indicating that this condition provided the optimal balance between response and experimental variability (Fig. 1E). Thus, plasmid titration combined with Z′-factor analysis enables identification of assay conditions that are not only responsive but also robust and reproducible.

**Figure 1.**
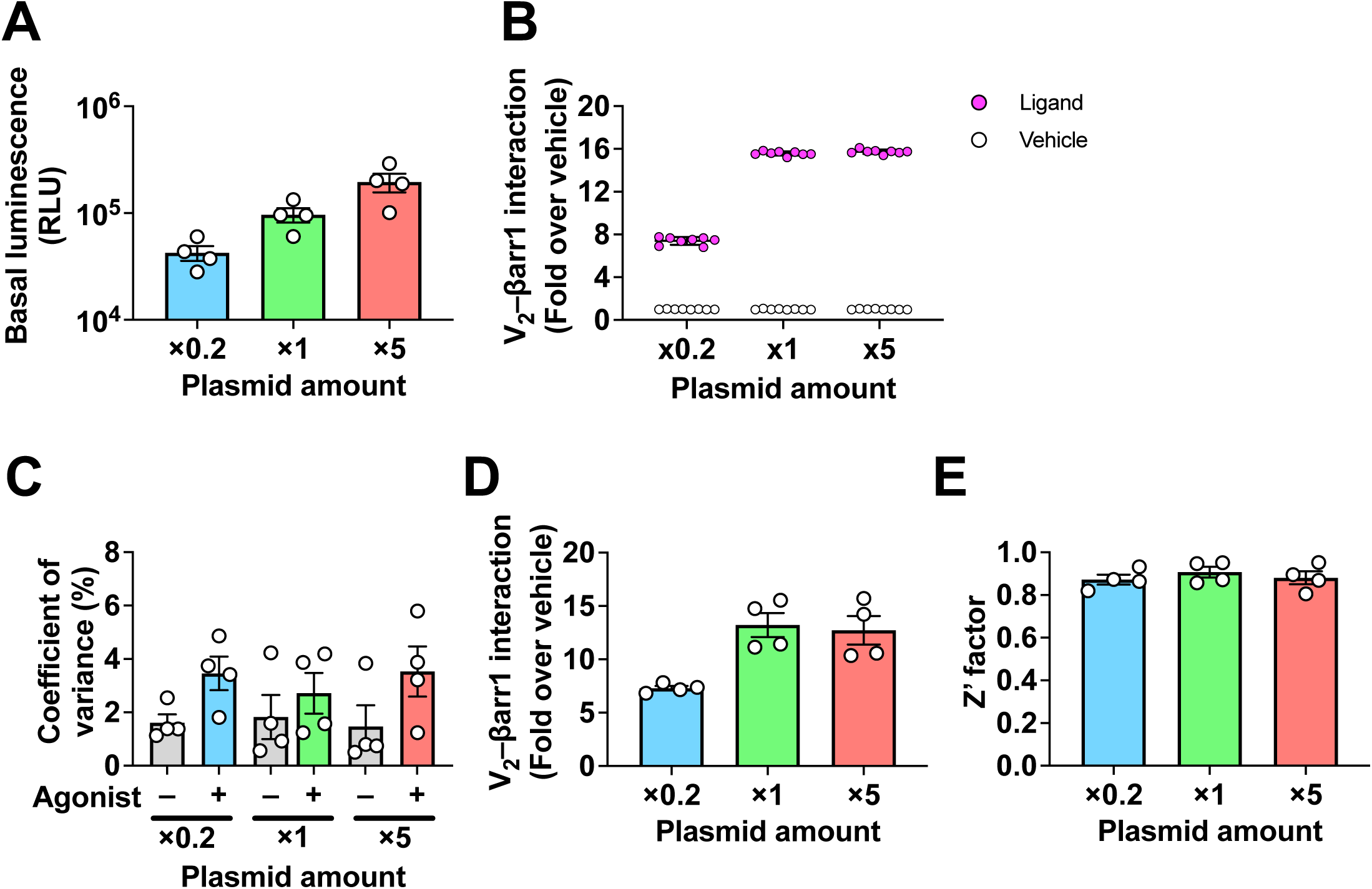
Assay-quality metrics and sensor-expression optimization for the NanoBiT assay. (A–D) Optimization of transfected plasmid amounts using the V_2_-SmBiT and LgBiT-β-arrestin1 constructs in the direct β-arrestin recruitment NanoBiT assay. Assay robustness and related parameters were evaluated using the standard plasmid amount, one-fifth of the standard amount, or fivefold the standard amount. Each condition was assessed in 16 wells (eight vehicle-treated and eight ligand-treated). Symbols and error bars represent the mean ± SEM of four independent experiments, each performed in duplicate. (A) Basal luminescence under each transfection condition. (B) Fold change from a representative independent experiment under each transfection condition. Symbols and error bars represent the mean ± SD of a single independent experiment. (C) Coefficient of variation (%) under each condition, with and without agonist. (D) Ligand-induced response (fold over vehicle) under each condition. (E) Z’ factor calculated from the assay data.

### Monitoring G-protein dissociation across Gα, Gβ, and Gγ subtypes

The G-protein dissociation assay monitors GPCR-mediated G-protein activation using LgBiT fused to Gα and SmBiT fused to Gγ of the Gβγ complex, as previously described [27] (Fig. 2A). Upon receptor activation, Gα-Gβγ dissociation reduces complementation between LgBiT and SmBiT, thereby indicating G-protein activation as a decrease of luminescence. Because this assay depends on productive complementation between engineered Gα and Gβγ in the inactive heterotrimer, the insertion site of LgBiT within Gα must preserve receptor coupling, nucleotide regulation, Gβγ association, and membrane localization.

**Figure 2.**
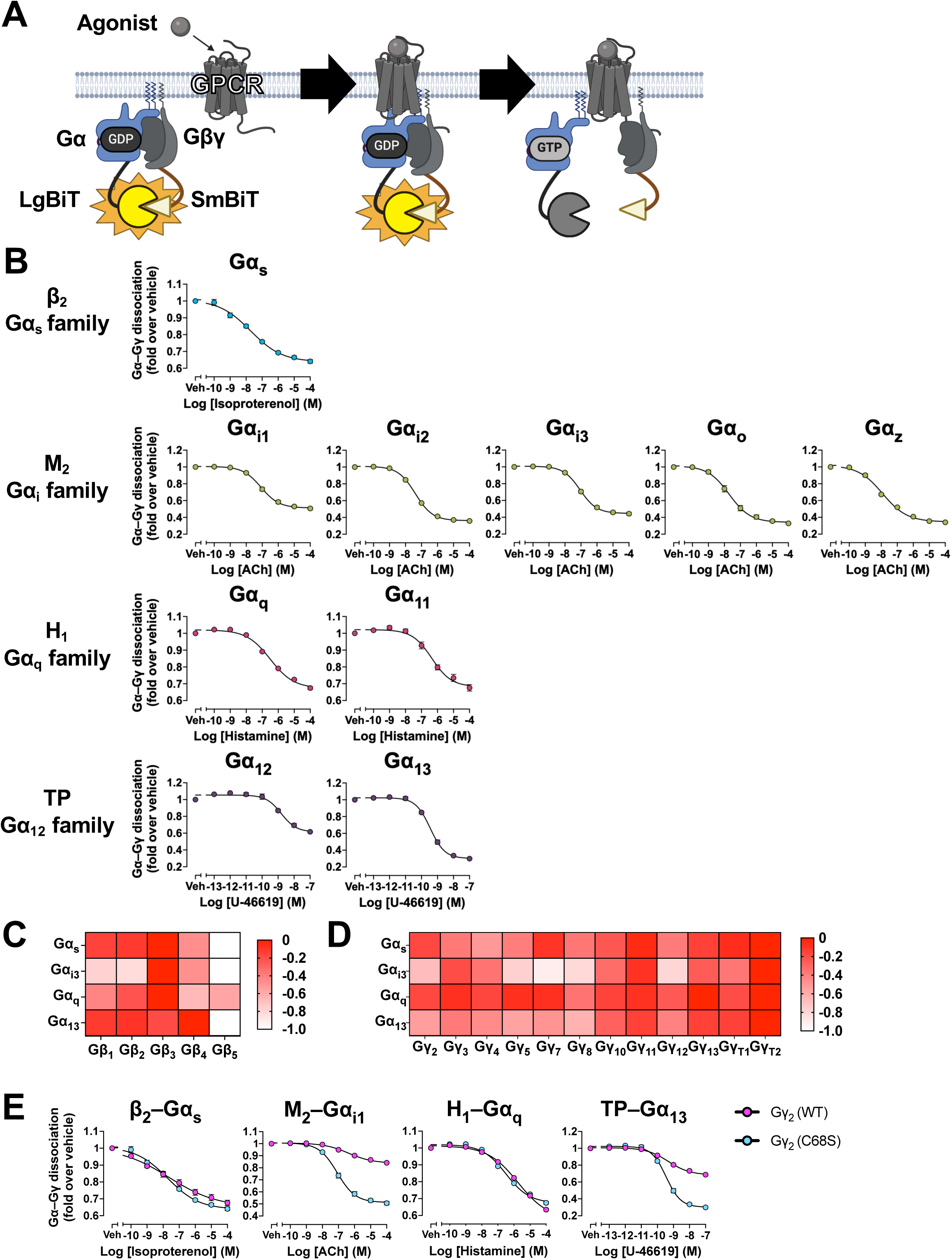
The Gα–Gβγ dissociation assay for profiling Gα, Gβ, and Gγ subtype selectivity. (A) Schematic of the Gα–Gβγ dissociation NanoBiT assay. Gα-LgBiT and SmBiT-Gβ or SmBiT-Gγ reconstitute NanoLuc in the resting heterotrimer; receptor activation dissociates the heterotrimer into Gα and Gβγ, decreasing luminescence. (B) Concentration–response curves of the Gα–Gβγ dissociation assay for representative GPCRs that primarily couple to each of the four Gα subfamilies (β_2_/Gα_s/olf_, M_2_/Gα_i/o_, H_1_/Gα_q/11_, TP/Gα_12/13_). Symbols and error bars represent the mean ± SEM of three independent experiments, each performed in duplicate. (C) Heatmap of Gβ selectivity across five Gβ subunits, measured with Gα-LgBiT and SmBiT-Gβ together with Gγ_2_. Values are ΔLogRAi calculated from *E*_max_ and *EC*_50_; for each Gα dataset, ΔLogRAi was obtained by subtracting the highest LogRAi value across all Gβ conditions. Data represent the mean of three independent experiments, each performed in duplicate. (D) Heatmap of Gγ selectivity across 12 Gγ subunits, measured with Gα-LgBiT and SmBiT-Gγ together with Gβ_1_. Values are ΔLogRAi calculated from *E*_max_ and *EC*_50_, defined as in (C). Data represent the mean of 3–4 independent experiments, each performed in duplicate. (E) Comparison of assay performance using wild-type or C68S Gγ_2_, shown for representative Gα subunits from the four subfamilies (Gα_s_, Gα_i1_, Gα_q_, and Gα_13_). Symbols and error bars represent the mean ± SEM of three independent experiments, each performed in duplicate.

In our Gα constructs, the LgBiT fragment was inserted between the αA and αB helices of Gα. This site was selected because it lies within the α-helical domain, away from major functional elements of the Ras-like domain that mediate GPCR engagement, nucleotide binding, and GTP hydrolysis [28]. In addition, the αA–αB region is positioned in spatial proximity to the N-terminus of Gγ, where SmBiT is fused, thereby favoring efficient NanoBiT complementation in the inactive heterotrimeric state. By contrast, insertion into regions such as the C-terminal α5 helix or the N-terminal αN was avoided because these regions are directly involved in receptor-catalyzed G-protein activation and membrane localization, respectively. Thus, the αA–αB insertion site was chosen to preserve essential Gα functions while allowing dissociation-dependent changes in NanoBiT complementation to be detected.

To validate this design, we generated a panel of Gα-LgBiT sensors covering nearly all Gα subfamilies and examined whether GPCR stimulation induced ligand-dependent decreases in NanoBiT luminescence. Across the tested Gα subtypes, receptor activation produced clear dissociation responses, indicating that insertion of LgBiT into the αA–αB region preserves the ability of Gα to form inactive heterotrimers with Gβγ and to undergo activation-dependent dissociation (Fig. 2B, S1A). These results support the αA–αB insertion site as a generally applicable position for monitoring GPCR-induced G-protein activation across multiple Gα subtypes.

In addition to validating the Gα insertion site, we evaluated whether the NanoBiT G-protein dissociation assay could be extended to different Gβ and Gγ subtypes. Mammalian cells express multiple Gβ and Gγ subtypes with distinct biochemical properties and tissue-expression profiles. Therefore, we asked whether the NanoBiT-based dissociation assay could be applied to other Gβ and Gγ subtypes. To assess Gβ-subtype compatibility, five N-terminal SmBiT-fused Gβ subtypes were tested in combination with Gγ_2_ (Fig. 2C). To assess Gγ-subtype compatibility, 12 N-terminal SmBiT-fused Gγ subtypes were tested in combination with Gβ_1_ (Fig. 2D, S2B, S3B). For both evaluations, we used representative Gα–LgBiT constructs. Among the Gβ subtypes tested, SmBiT-fused Gβ_5_ produced both the lowest basal luminescence and the weakest ligand-induced responses across the receptors examined (Fig. S3A). Gβ_5_ is predominantly expressed in the nervous system and preferentially forms complexes with members of the R7 family of RGS proteins and R7BP rather than with conventional Gγ subunits, which tethers it to the membrane [29]. Its limited ability to form functional complexes with the Gγ subunits used in our assay may therefore have impaired plasma membrane localization and reduced the observed responses. Gβ_3_ produced the largest responses among the Gβ subtypes tested, consistent with a previous study that optimized BRET-based G-protein dissociation assays [5] (Fig. S2A, S3A). Among the Gγ subtypes, Gγ_13_-, GγT_2_-, and in some cases Gγ_11_-SmBiT supported stronger ligand-dependent responses than other Gγ subtypes. Notably, both GγT_2_ and Gγ_11_ are farnesylated with a C15 isoprenoid, whereas most other Gγ subtypes are modified with a C20 geranylgeranyl group [30]. Because farnesylation generally provides weaker membrane anchoring than geranylgeranylation [31], the relatively large ligand-dependent responses obtained with these subtypes may reflect more efficient separation of Gα from the Gβγ complex following receptor activation. These results indicate that the NanoBiT G-protein dissociation assay can be adapted to multiple Gβγ subtype combinations.

For general profiling of receptor-induced G-protein dissociation, we used Gβ_1_/Gγ_2_ as the standard Gβγ pair. This pair provides a practical reference condition for comparing responses across different Gα-LgBiT sensors owing to their broad expression and suitability as representative Gβγ components. Other Gβγ combinations can be used when the aim is to examine subtype-specific Gβγ contributions or model a defined cellular context. Thus, Gβ_1_/Gγ_2_ was adopted as the default configuration for general GPCR–G-protein dissociation measurements, whereas the broader Gβγ subtype panel provides flexibility for subtype-focused applications.

The signal intensity of the G-protein dissociation assay could be further enhanced through targeted modification of G-protein subunits. The C-terminal CAAX motif of Gγ is required for prenylation and membrane anchoring [32]. We therefore introduced a C68S mutation into Gγ_2_ to disrupt this prenylation site and examined whether reduced membrane anchoring would facilitate dissociation from Gα. Compared with wild-type Gγ_2_, the Gγ_2_ C68S mutant significantly increased the magnitude of the ligand-dependent decrease in NanoBiT luminescence for Gα_i1_ and Gα_13_, whereas only a modest enhancement was observed for Gα_s_ (Fig. 2E, S1B). Thus, the effect of impaired Gγ_2_ prenylation on assay dynamic range varied among the Gα subtypes tested. These findings indicate that modulation of Gγ membrane anchoring can enhance the sensitivity of the dissociation assay, although the extent of improvement depends on the Gα-containing heterotrimer.

### Development of NanoBiT sensors for monitoring β-arrestin recruitment, compartmental trafficking and activation

β-arrestin recruitment to the plasma membrane can be monitored using two complementary evaluation modes: direct and bystander (Fig. 3A). In the direct mode, NanoBiT complementation is measured between the GPCR and β-arrestin, with the GPCR tagged with the smaller SmBiT fragment to minimize steric interference. This configuration enables a direct readout of GPCR–β-arrestin interactions and typically yields stronger recruitment signal change than the bystander approach. In contrast, the bystander mode assesses β-arrestin recruitment to the plasma membrane by measuring complementation between SmBiT-tagged β-arrestin and an LgBiT-fused to the C-terminus of membrane-anchored CAAX motif, a configuration designed to maintain stable expression of the CAAX sensor. Although this approach does not directly report GPCR–β-arrestin binding, it offers several advantages. First, the bystander mode allows the use of intact, unmodified GPCRs, thereby preserving the native receptor architecture and enabling assessment of the intrinsic ability of receptors to recruit and activate β-arrestins. Second, it can capture β-arrestin recruitment events that may arise from catalytic activation of β-arrestin by GPCRs, in which β-arrestin transiently interacts with an activated receptor and subsequently translocates to pre-endocytic membrane compartments [33].

**Figure 3.**
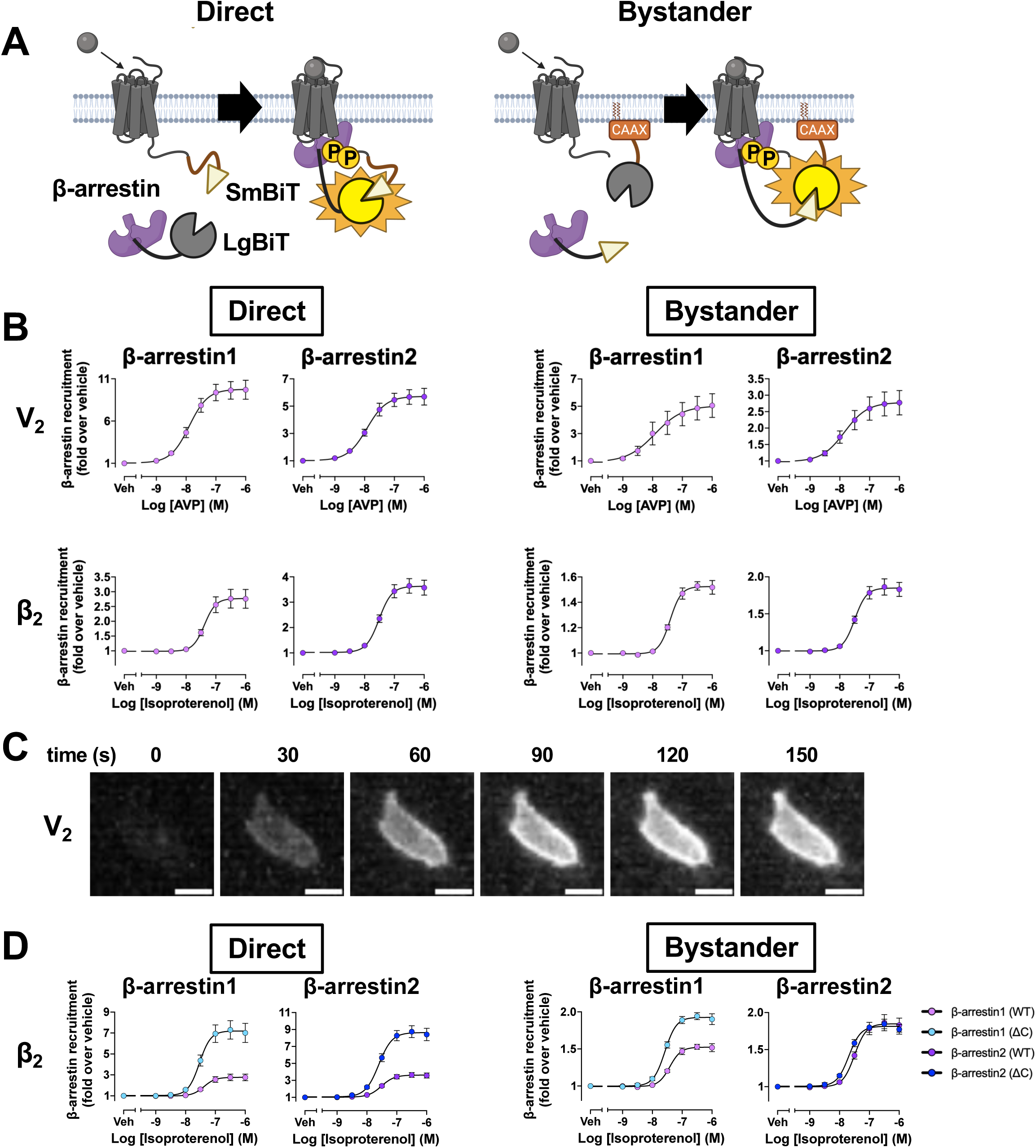
The direct and bystander methods for β-arrestin recruitment to GPCRs and the plasma membrane. (A) Schematic of the two configurations. Left (direct): GPCR-SmBiT and LgBiT-β-arrestin reconstitute NanoLuc upon agonist-induced β-arrestin recruitment to the activated receptor. Right (bystander): membrane-anchored LgBiT-CAAX and SmBiT-β-arrestin detect β-arrestin recruitment to the plasma membrane, increasing luminescence. (B) Concentration–response curves for β-arrestin1 and β-arrestin2 recruitment to V_2_ and β_2_ receptors, measured by the direct (left) and bystander (right) assays. Symbols and error bars represent the mean ± SEM of 3–5 independent experiments, each performed in duplicate. (C) Representative bioluminescence microscopy images of direct β-arrestin recruitment to V2R after stimulation with 1 μM AVP. Scale bar, 10 μm. (D) Effect of C-terminal truncation, comparing full-length β-arrestin with the β-arrestin(ΔC) mutant lacking residues after position 382, for the β_2_ receptor in the direct (left) and bystander (right) assays. The β-arrestin1 (WT) and β-arrestin2 (WT) data from (B) are replotted together with β-arrestin (ΔC) data acquired in the same experimental sets. Symbols and error bars represent the mean ± SEM of 3– 5 independent experiments, each performed in duplicate.

For both the direct and bystander β-arrestin assays, we fused the LgBiT fragment to the N-terminus of β-arrestin to preserve native signaling behavior while enabling robust and stable NanoBiT complementation. Compared with the C-terminal region, which undergoes dynamic displacement upon interaction with the phosphorylated receptor C-tail, the N-terminal region of β-arrestin is expected to be less affected by the conformational rearrangements associated with receptor engagement [34,35]. We therefore reasoned that the N-terminal region would provide a favorable site for LgBiT fusion.

Using LgBiT-fused β-arrestin1/2, we evaluated β-arrestin recruitment to the receptor and the plasma membrane in direct and bystander assay configurations, respectively. For this evaluation, we employed V_2_R as a model receptor because it is known to robustly recruit β-arrestins [36,37]. In both direct and bystander configurations, we observed a concentration-dependent increase in luminescence (Fig. 3B, S4A). By contrast, β_2_AR, which recruits β-arrestins less efficiently than V_2_R, produced only a modest increase in luminescence. We next examined whether β-arrestin recruitment could also be visualized by luminescence microscopy (Fig. 3C). Following V_2_R stimulation, an increase in β-arrestin localization at the plasma membrane became apparent within approximately 60 seconds. Together, these results indicate that the LgBiT-fused β-arrestin1/2 sensors retain functional responsiveness and enable detection of receptor-dependent β-arrestin recruitment in both direct and bystander assay formats, as well as visualization of recruitment dynamics by microscopy.

When evaluating GPCRs that induce only weak β-arrestin recruitment, the limited response of β-arrestin recruitment assay can be overcome by truncation of β-arrestin C-terminal tail (ΔC). Under basal conditions, the C-tail of β-arrestin engages the positively charged groove within the N-domain, functioning as an autoinhibitory element and truncation of this C-tail promotes a pre-activated conformational state that increases its propensity to interact with GPCRs [38,39]. We applied this concept to our NanoBiT system and engineered LgBiT-fused β-arrestin constructs lacking the C-tail to enhance the sensitivity of the receptor recruitment assay. Upon β_2_AR ligand stimulation, a substantial increase in responses was detected for the direct β-arrestin receptor recruitment assay compared to that of wild-type β-arrestin in both subtypes (Fig. 3D, S4C). In contrast, only a modest increase in signal was observed in the bystander configuration. We also performed similar evaluations downstream of V_2_R. Interestingly, β-arrestin (ΔC) resulted in an apparent decrease in fold change over basal compared with wild-type β-arrestin in both the direct and bystander configurations (Fig. S4B, S4D). This apparent decrease reflects increased basal luminescence for receptors with intrinsically strong β-arrestin recruitment, which reduces the dynamic range of ligand-induced responses. Together, these results indicate that β-arrestin (ΔC) constructs are useful for sensitively detecting β-arrestin recruitment downstream of receptors with relatively weak β-arrestin recruitment efficacy, whereas their utility may depend on the receptor and assay configuration.

Beyond plasma membrane localization, β-arrestins translocate to endosomes, a process that is particularly prominent for trafficking class B GPCRs [40]. Using V_2_R as a representative trafficking class B receptor, we evaluated β-arrestin endosomal recruitment by measuring NanoBiT complementation between SmBiT–β-arrestin1 and Endofin, an early endosomal marker [41], fused to LgBiT (Endo–LgBiT) (Fig. 4A). We observed an increase in luminescence upon ligand stimulation (Fig. 4B, S5A). Because the NanoBiT assay does not provide spatial information, we verified that the observed response reflected endosomal translocation rather than complementation with a fraction of Endo–LgBiT potentially mislocalized to the plasma membrane. To this end, we examined the effect of Dyngo-4a, an inhibitor of dynamin-dependent endocytosis (McCluskey et al, 2013). We confirmed that these ligand-dependent complementation signals were strongly attenuated by treatment with Dyngo-4a, indicating that the observed responses reflect β-arrestin early endosomal translocation. Similar results were observed with Rab5, another early endosomal marker [43], downstream of both V_2_R and AT_1_R (Fig. 4C, S5B). We next evaluated other endosomal markers including Rab4, Rab7, and Rab11, which label early/recycling, late, and recycling endosomal compartments, respectively [44,45]. Rab4- and Rab11-based LgBiT sensors produced robust ligand-dependent responses, and these responses were attenuated by Dyngo-4a treatment, consistent with their dependence on endocytic trafficking. In contrast, the Rab7-based sensor did not yield a detectable response, potentially due to its relatively low expression suggested by its basal luminescence. Collectively, these findings demonstrate that β-arrestin translocation to endosomal compartments can be monitored using Endofin- or Rab-based NanoBiT sensors, although Rab7 was not suitable under the present experimental conditions.

**Figure 4.**
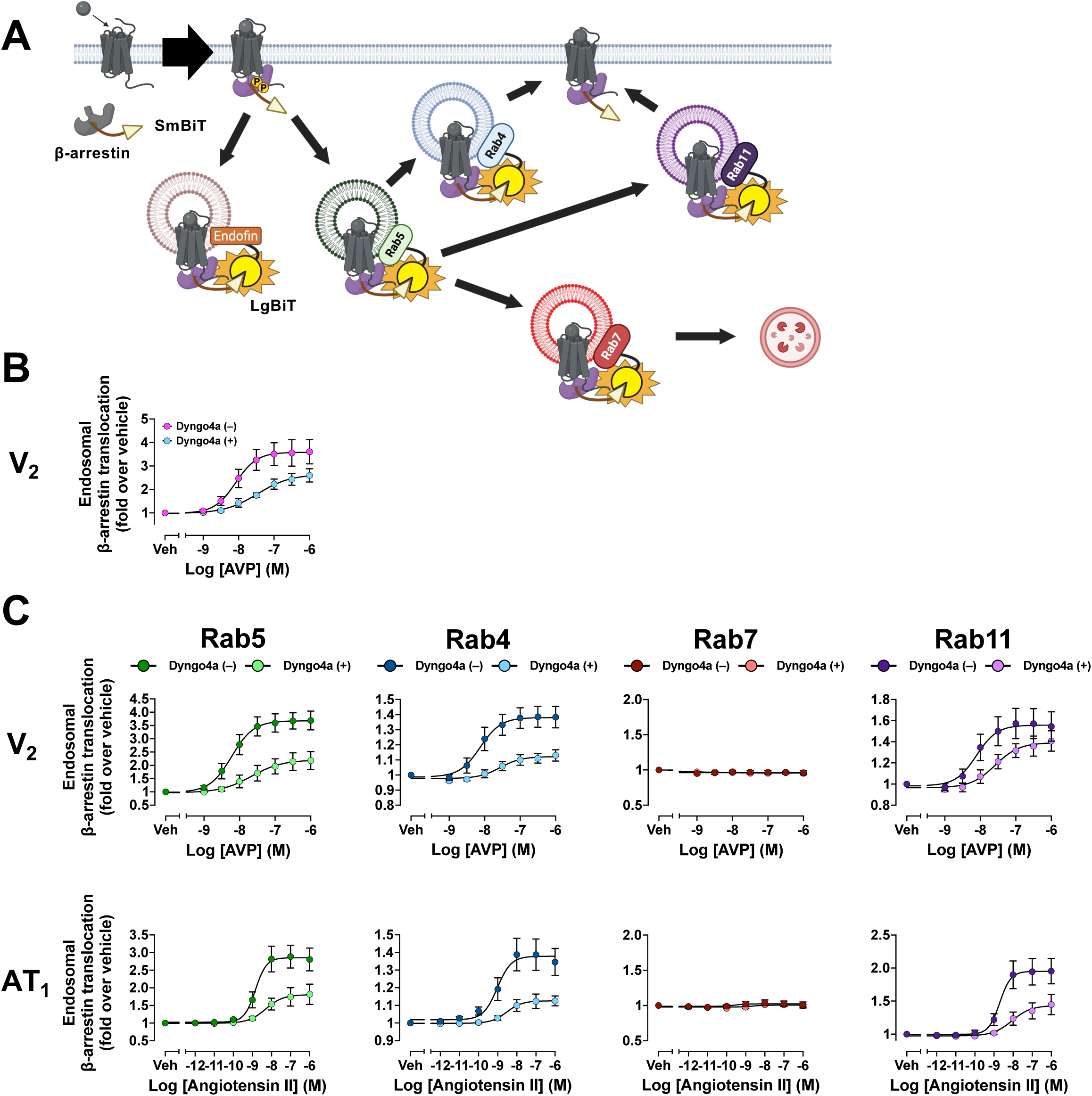
The endosomal β-arrestin translocation assay. (A) Schematic of the endosomal translocation assay. LgBiT-fused endosomal markers (Endofin or Rab proteins) and SmBiT-β-arrestin1 are co-expressed; increased luminescence reflects proximity of β-arrestin1 to endosomal compartments following agonist-induced receptor internalization. (B) â-arrestin1 translocation to Endofin-positive compartments at the V_2_ receptor, with or without the dynamin inhibitor Dyngo-4a. Symbols and error bars represent the mean ± SEM of three independent experiments, each performed in duplicate. (C) â-arrestin1 translocation to Rab4-, Rab5-, Rab7-, and Rab11-positive compartments at the V_2_ and AT_1_ receptors, with or without Dyngo-4a. Symbols and error bars represent the mean ± SEM of three independent experiments, each performed in duplicate.

In addition to intracellular localization, β-arrestins have recently been reported to directly interact with distinct membrane compositions, thereby contributing to their localization within specific membrane domains [46]. Based on this finding, we sought to expand our NanoBiT sensor platform to monitor β-arrestin localization within defined plasma membrane microdomains using membrane-domain markers (Fig. 5A). To this end, we employed Lyn and GAP43 as lipid raft-associated markers and CD86 as a non-lipid raft marker [47,48]. Using LgBiT-fused constructs for each membrane marker, we observed ligand-dependent increases in luminescence to varying degrees, with CD86-based sensors producing the largest response for β-arrestin1 (Fig. 5B, S6A). The ligand-dependent increase observed with the GAP43–LgBiT and SmBiT–β-arrestin1 pair was consistent with a previous study in which a GAP43-based split NanoLuc sensor was used to monitor β-arrestin recruitment to the plasma membrane [49]. These results indicate that, following GPCR activation, β-arrestins can access membrane environments associated with both raft and non-raft markers.

**Figure 5.**
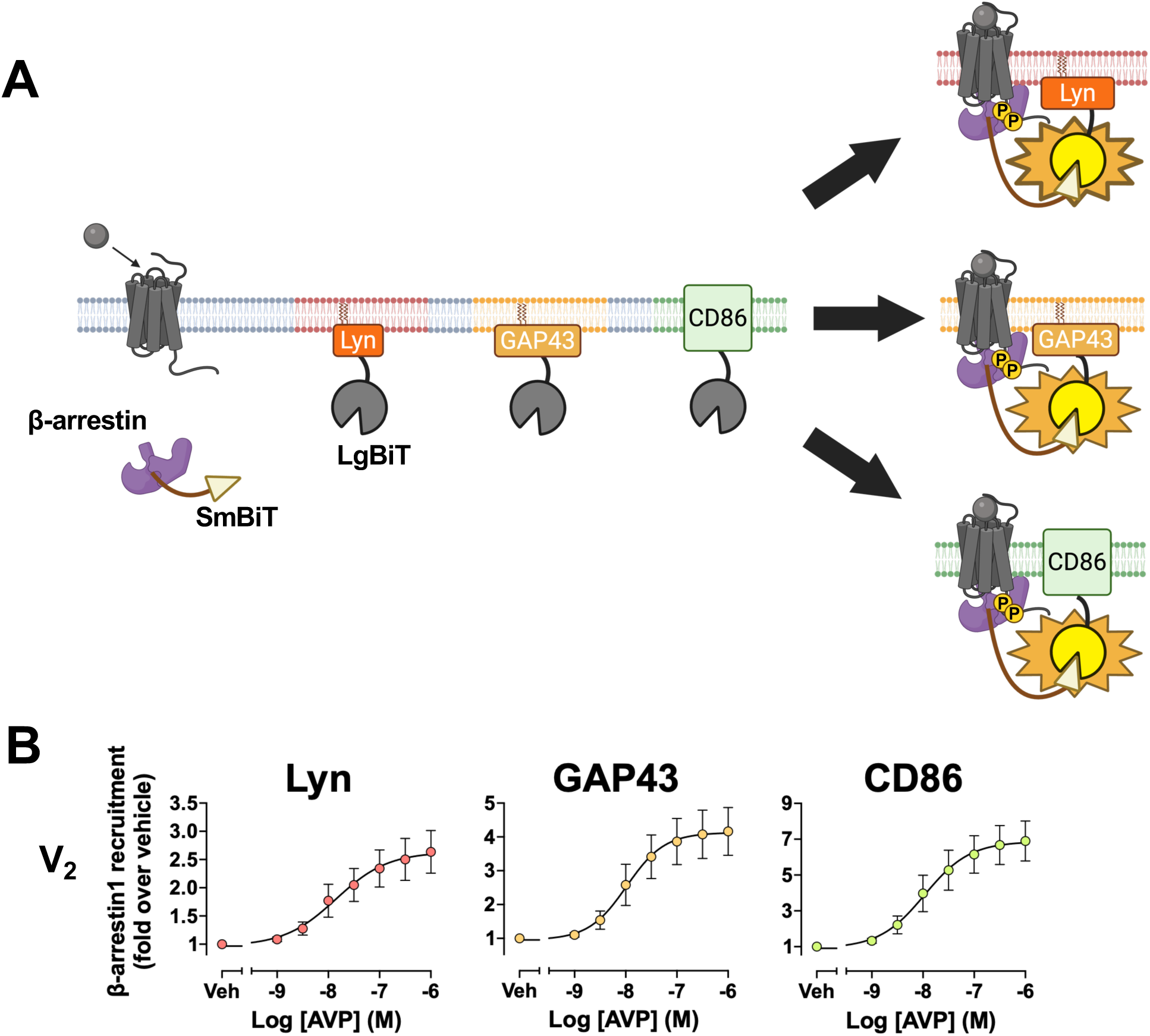
The bystander methods to monitor β-arrestin accumulations in the membrane domains. (A) Schematic of the bystander assay using LgBiT-fused plasma-membrane markers with distinct microdomain localization: the lipid-raft markers Lyn and GAP43 and the non-raft marker CD86. β-arrestin1 recruitment to each microdomain is detected as increased luminescence. (B) Concentration–response curves for β-arrestin1 recruitment to Lyn-, GAP43-, and CD86-defined microdomains at the V_2_ receptor. Symbols and error bars represent the mean ± SEM of 3–4 independent experiments, each performed in duplicate.

β-arrestin NanoBiT sensors can also be extended to monitor additional β-arrestin-specific mechanisms, including conformational activation and its function as a scaffold for MAPK signaling [50]. We first established intrabody-based NanoBiT sensors using Ib4 and Ib30, which have been developed to recognize distinct active conformational states of β-arrestin [34,51]. To accommodate the molecular size of these intrabodies and support stable expression, the LgBiT fragment was fused to the N-terminus of each intrabody, while the complementary SmBiT fragment was fused to the N-terminus of β-arrestin1. Using these sensor pairs, we observed robust agonist concentration-dependent increases in luminescence with both intrabodies, indicating successful detection of β-arrestin conformational activation (Fig. S7A, 7B, 8A). We next examined whether the scaffold function of β- arrestins could be detected using this NanoBiT-based approach. For this purpose, LgBiT was fused to the extreme N-terminus of Src, and we evaluated its interaction with SmBiT–β-arrestin1/2 downstream of V_2_R (Fig. S7C). Ligand stimulation induced a concentration-dependent increase in β- arrestin1/2 association with Src (Fig. S7D, 8B). Together, these results demonstrate that the NanoBiT platform can be used not only to monitor β-arrestin recruitment and subcellular localization, but also to detect conformational activation and interactions associated with its scaffolding functions.

### Monitoring GRK recruitment to GPCRs

The GRK recruitment assay is based on complementation between SmBiT-tagged GPCRs and LgBiT-tagged GRKs. Although GRK subfamilies differ in their basal subcellular localization— GRK2/3 are predominantly cytosolic, whereas GRK5/6 are constitutively associated with the plasma membrane—receptor activation results in increased NanoBiT complementation for both groups. Accordingly, an increase in luminescence serves as a robust readout of GPCR–GRK interaction (Fig. 6A).

**Figure 6.**
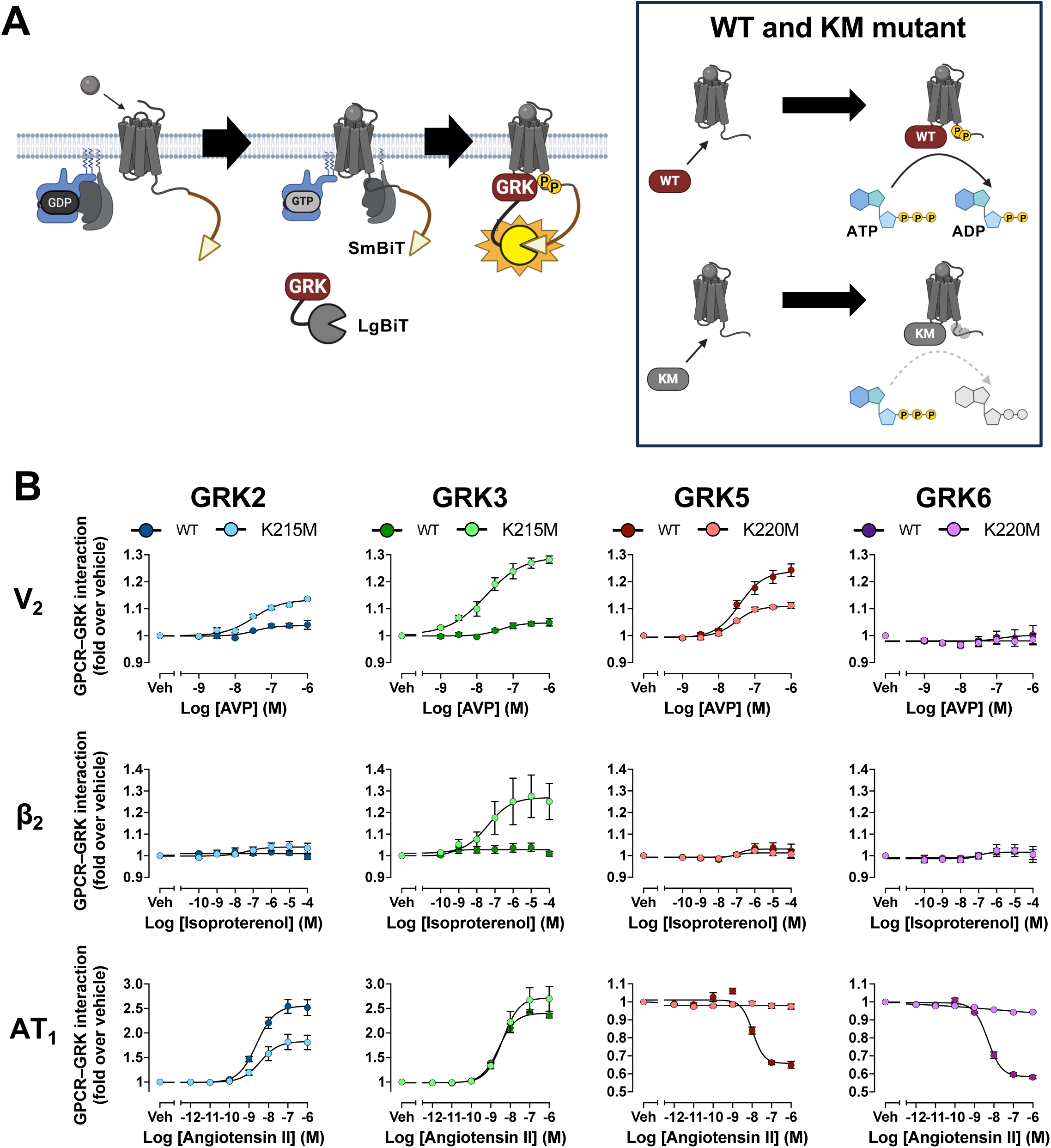
The GRK recruitment assays. (A) Schematic of the GPCR–GRK recruitment assay for the cytosolic GRK2/3 and the membrane-associated GRK5/6 subfamilies. GPCR-SmBiT and GRK-LgBiT detect agonist-induced recruitment of GRKs to the activated receptor. (B) Concentration–response curves for recruitment of GRK2, GRK3, GRK5, and GRK6 to the V_2_, β_2_, and AT_1_ receptors, derived from kinetic data averaged between 1 and 5 min after ligand addition. Kinase-inactivating KM mutants (K220M for GRK2/3, K215M for GRK5/6), which slow dissociation from the receptor, were tested in parallel. Symbols and error bars represent the mean ± SEM of three independent experiments, each performed in duplicate.

To minimize disruption of functionally important regions of GRKs, we fused the LgBiT fragment to the C-terminal region. Previous BRET sensors have successfully fused NanoLuc distal to the C-terminal Gβγ-binding region of GRK3 while retaining its ability to interact with activated Gβγ [7], supporting the feasibility of C-terminal fusion. In contrast, several reports have demonstrated that the N-terminal region of GRKs is critical for their kinase activity [52–54]. Based on these findings, we reasoned that C-terminal fusion of LgBiT would be less likely to interfere with GRK activation while still permitting GRK recruitment to GPCRs.

We evaluated GPCR–GRK interactions using GRK–LgBiT constructs together with V_2_R, β_2_AR, and AT_1_R, each fused to SmBiT at its C-terminus (Fig. 6B, S9A). V_2_R showed clear ligand-dependent recruitment of GRK2, GRK3, and GRK5, whereas no detectable response was observed with GRK6. For the responsive GRKs, the NanoBiT signal declined continuously after reaching its peak, possibly reflecting GRK dissociation following receptor phosphorylation. Downstream of β_2_AR, ligand-dependent recruitment was detected for all GRKs tested, with GRK3 producing the largest response. The response profiles differed downstream of AT1R: GRK2 and GRK3 exhibited robust recruitment, whereas the GRK5 response was comparatively modest, and no detectable response was observed with GRK6. The preferential responses of GRK2 and GRK3 over GRK5 and GRK6 are in line with a previous report showing that Gα_q_ activation downstream of AT_1_R preferentially promotes receptor engagement by GRK2/3 [55]. Collectively, these results demonstrate that GPCR–GRK recruitment can be monitored using the GRK–LgBiT constructs, although assay performance varies depending on the receptor–GRK combination.

The sensitivity of the GRK recruitment assay may be enhanced for certain receptors through targeted modification of the GRK kinase domain. Protein kinases, including GRKs, contain a highly conserved catalytic domain in which the invariant lysine within the VAIK motif of the N-lobe is essential for phosphotransfer activity [56,57]. Accordingly, mutation of the corresponding lysine in several GRK subtypes has been shown to abolish kinase activity toward GPCRs [58–60]. We therefore generated kinase-deficient GRK mutants by substituting this conserved lysine with methionine (hereafter referred to as KM): K220M in GRK2 and GRK3, and K215M in GRK5 and GRK6. We hypothesized that preventing receptor phosphorylation might prolong the GPCR–GRK interaction while preserving the ability of GRKs to recognize activated receptors, thereby increasing the assay window. Using LgBiT-tagged GRK KM mutants, we re-evaluated GRK responses downstream of V_2_R, β_2_AR, and AT_1_R (Fig. 6B, S9A). Compared with wild-type GRK5, the kinase-dead GRK5 (KM) mutant produced a larger ligand-dependent response with V_2_R. The kinetic profile of GRK5 (KM) also showed a more gradual decline than that of wild-type GRK5, suggesting that the KM mutation prolonged the receptor-associated interaction. Downstream of β_2_AR, none of the GRK KMs produced a detectable ligand-dependent recruitment response. In contrast, the KMs produced distinct response profiles downstream of AT_1_R. GRK2(KM) showed the largest improvement in response magnitude relative to its wild-type counterpart, whereas GRK3 (KM) showed no significant change. Unexpectedly, GRK5 (KM) and GRK6 (KM) exhibited ligand-dependent decreases in luminescence, suggesting increased separation of these GRKs from AT_1_R following receptor activation. Previous studies have reported that GRK5 can form preassembled clusters with AT_1_R under basal conditions and that its interaction with phosphatidylinositol 4,5-bisphosphate (PIP_2_) contributes to its plasma membrane localization [55,61]. One possible explanation is therefore that GRK5 and GRK6 are associated with AT_1_R-containing membrane domains before stimulation and subsequently redistribute following AT_1_R-mediated activation of the Gα_q_–PLCβ pathway and PIP_2_ hydrolysis. Overall, these results demonstrate that kinase-inactivating mutations can enhance the sensitivity of the GRK recruitment assay in a receptor- and GRK subtype-dependent manner.

### Evaluating Gα_s/olf_- and Gα_i/o_-protein activity via adenylyl cyclase association assay

As an additional readout of Gα_s/olf_- and Gα_i/o_-protein activity, we extended our NanoBiT platform to monitor interactions between Gα proteins and each of the nine transmembrane adenylyl cyclase (AC) subtypes [62]. Unlike conventional cAMP-based assays, in which Gα_i/o_ activity is typically assessed as inhibition of forskolin-stimulated cAMP production [63], this assay directly detects Gα–AC interactions without requiring forskolin pre-stimulation. Luminescence is generated by complementation between LgBiT-fused Gα proteins and SmBiT-fused AC constructs, thereby enabling quantitative measurement of ligand-dependent Gα–AC interactions (Fig. 7A).

**Figure 7.**
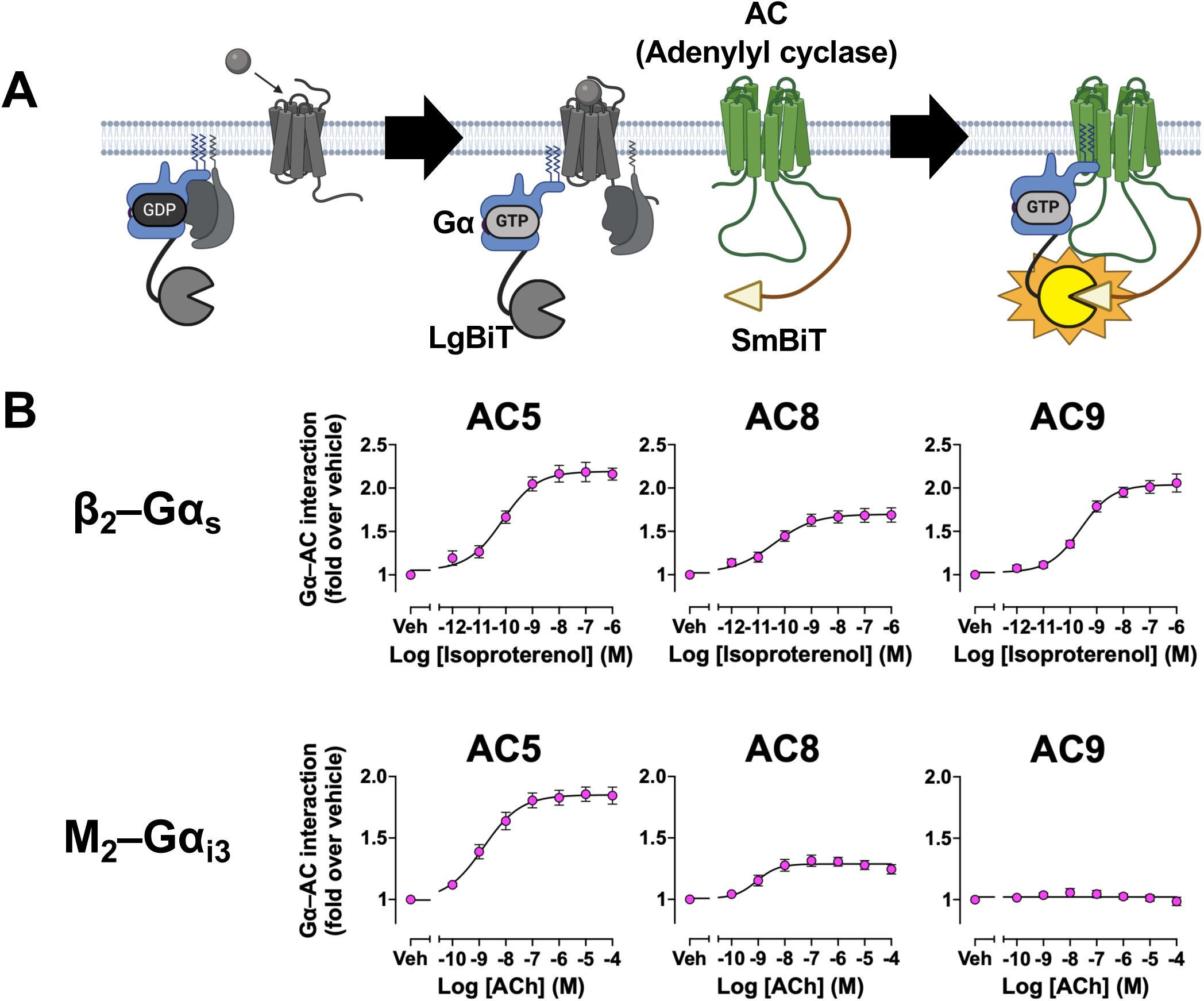
The Gα–adenylyl cyclase association assay. (A) Schematic of the Gα–adenylyl cyclase (AC) interaction assay. Gα-LgBiT and SmBiT-AC detect agonist-induced proximity between activated Gα_s_ or Gα_i_ and AC. (B) Concentration–response curves for the Gα_s_–AC interaction at the β_2_ receptor and the Gα_i3_–AC interaction at the M_2_ receptor, shown for the responsive isoforms AC5, AC8, and AC9. Symbols and error bars represent the mean ± SEM of three independent experiments, each performed in duplicate.

The two catalytic domains of AC, C1a and C2a, are located in the intracellular regions between and downstream of the two transmembrane clusters, respectively. Therefore, we fused SmBiT to the N terminus of each AC isoform to minimize potential interference with catalytic-domain function. A previous study has also shown that AC8 retains functional expression when fused to eGFP at its N terminus [64]. On this basis, we generated a series of SmBiT–AC1–9 constructs. Using representative SmBiT–AC sensors together with LgBiT-fused Gα_s_ or Gα_i_ proteins, we readily detected ligand-dependent interactions between Gα proteins and specific AC subtypes (Fig. 7B, S10A). Robust responses were observed between Gα_s_ and AC5, AC8, and AC9. Gα_i3_ also produced clear responses with AC5 and AC8, whereas no detectable interaction was observed with AC9. By contrast, no ligand-dependent interaction was detected between Gα_i3_ and AC9. This finding is consistent with the previous report that AC9 is not directly inhibited by Gα_i/o_ proteins [65]. The absence of a detectable Gα_i3_–AC9 interaction may therefore reflect subtype-specific differences in the regulation of AC9, although the underlying structural determinants remain unclear. We subsequently examined interactions between the tested Gα proteins and the remaining AC subtypes (Fig. S10B, 10B). Under the assay conditions used, no clear ligand-dependent responses were detected for these additional combinations. This may reflect differences among AC subtypes in their subcellular localization, expression level, accessibility to activated Gα proteins, or intrinsic G-protein selectivity [66]. In addition, the relative positioning of the fused NanoBiT fragments may not have been favorable for productive complementation in some sensor combinations. Thus, failure to detect a response does not necessarily indicate the absence of a functional interaction. Collectively, these results demonstrate that the NanoBiT platform can directly detect selected Gα–AC interactions in living cells.

### Evaluating Gα_q/11_-protein signaling activity via phospholipase Cβ (PLCβ) association assay

As a method to evaluate Gα_q_ protein downstream activity, we applied our NanoBiT system to detect Gα_q_–PLCβ interactions. The PLCβ association assay enables this evaluation by detecting increased luminescence resulting from complementation between LgBiT-fused Gα_q_ proteins and SmBiT-fused PLCβ [67] (Fig. 8A).

**Figure 8.**
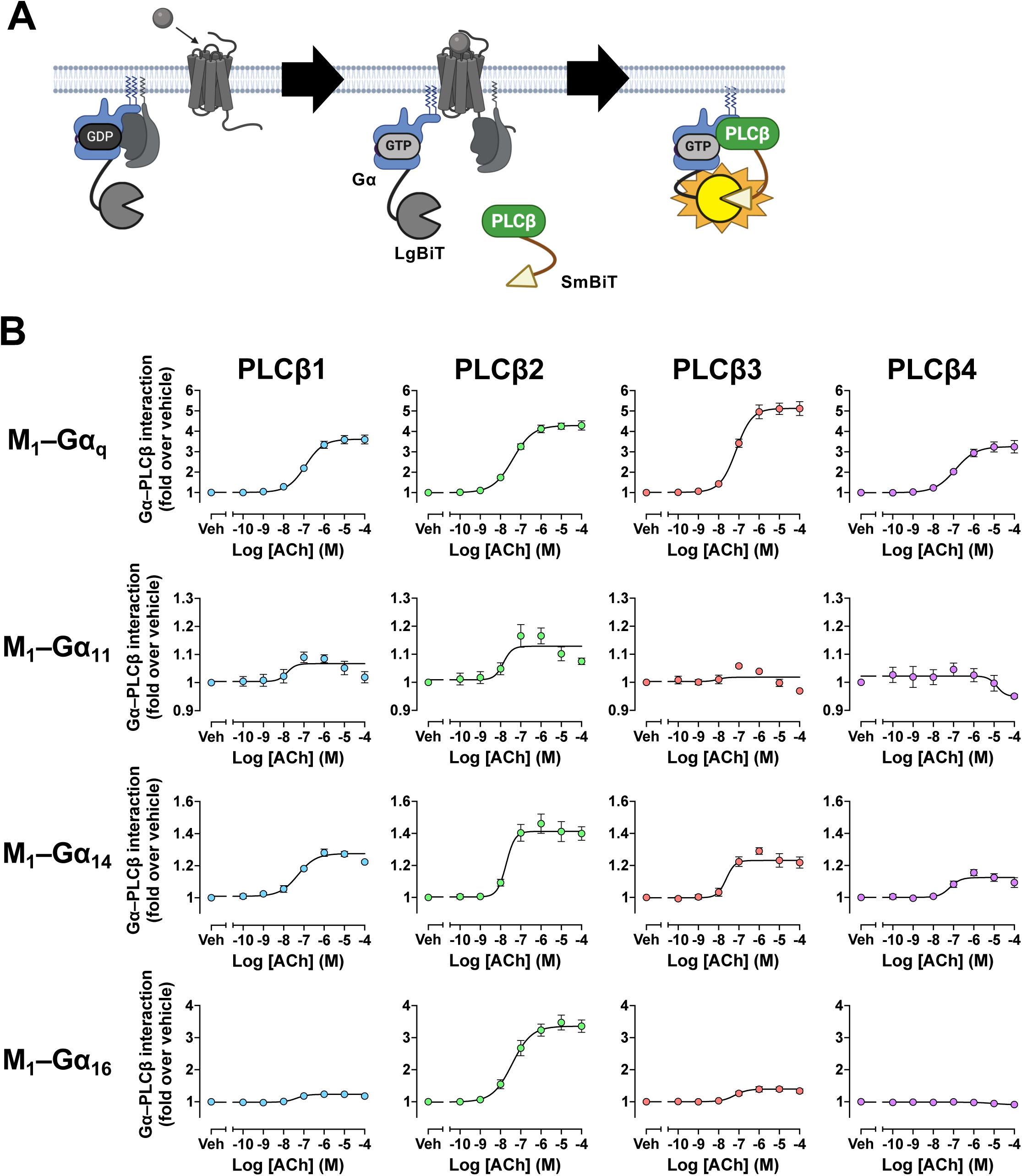
The Gα–PLCβ association assay. (A) Schematic of the Gα_q_–PLCβ interaction assay. Gα_q_-LgBiT (fragment inserted at position 123) and SmBiT-PLCβ detect agonist-induced proximity between activated Gα_q_ and PLCβ. (B) Concentration–response curves for interactions of Gα_q_, Gα_11_, Gα_14_, and Gα_16_ with PLCβ1, PLCβ2, PLCβ3, and PLCβ4, measured at the M_1_ receptor (ssHF-M_1_) and derived from kinetic data averaged between 3 and 5 min after ligand addition. Symbols and error bars represent the mean ± SEM of three independent experiments, each performed in duplicate.

In the PLCβ association assay, we employed LgBiT-fused Gα_q_ proteins together with SmBiT fused to the N-terminus of PLCβ. For Gα_q_, the NanoBiT insertion strategy differed from that used in our G-protein dissociation assay, in which the tag was inserted within the αA–αB loop. Instead, LgBiT was fused at position 123 within the αB–αC helices of Gα_q_, based on a previous split-luciferase complementation assay for Gα_q_–PLCβ3 using click beetle luciferase (Pyrophorus plagiophthalamus) [68]. Comparison of constructs with insertions at position 97 within the αA–αB loop and position 123 within the αB–αC loop revealed that Gα_q_ (αB–αC) exhibited a higher signal-to-background (S/B) ratio, which is thought to be due to the closer proximity of both tags based on the Gα_q_-PLCβ interaction site [69,70] thereby improving the signal intensity. The choice of the N-terminal insertion site for PLCβ was selected owing to the critical regulatory functions of its C-terminal region, which undergoes substantial conformational rearrangements upon activation and is therefore not suitable for tag insertion [71].

We evaluated the functionality of the engineered sensors by pairing SmBiT-tagged PLCβ1– 4 with each LgBiT-fused member of the Gα_q/11_ subfamily and measuring their interactions downstream of the M_1_ muscarinic acetylcholine (M_1_) receptor activation (Fig. 8B, S11). Unexpectedly, the four Gα proteins displayed distinct PLCβ interaction profiles. Gα_q_ and Gα_14_ exhibited relatively high promiscuity against all PLCβ subtypes, whereas Gα_16_ showed a more restricted preference for PLCβ2 and PLCβ3. By contrast, Gα_11_ produced little or no detectable response across the PLCβ subtypes tested. The comparatively narrow interaction profile of Gα_16_ may partly reflect its greater sequence divergence from the other Gα_q/11_-family members, although the structural basis of this selectivity remains to be determined. Collectively, these results demonstrate that the PLCβ association assay can be applied across multiple PLCβ subtypes and enables comparative analysis of subtype-specific interaction preferences within the Gα_q/11_ family.

### Additional Gα_q_ protein activity via Gα_q_–p63-RhoGEF association assay

In addition to PLCβ, Gα_q_ activity can be monitored through its interaction with p63-RhoGEF [72]. Similar to other RhoGEFs, activation of p63-RhoGEF by Gα_q_ promotes downstream RhoA– ROCK signaling. Unlike the RGS-containing RhoGEFs regulated by Gα_12/13_, p63-RhoGEF lacks an RGS domain and interacts with activated Gα_q_ through its DH/PH module [73]. We therefore fused SmBiT to the extreme N-terminus of p63-RhoGEF to minimize potential interference with the Gα_q_- binding interface within the DH/PH region.

To evaluate the functionality of this sensor, we measured ligand-dependent complementation between Gα_q_–LgBiT and SmBiT–p63-RhoGEF following activation of the histamine H_1_ receptor (H_1_) and the M_1_ receptor (Fig. 9A, 9B, S12A). Stimulation of either receptor produced a clear increase in NanoBiT luminescence, indicating recruitment of p63-RhoGEF to activated Gα_q_. These results are consistent with previous reports of receptor-dependent Gα_q_–p63- RhoGEF engagement [74] and provide proof of concept for the functionality of the engineered sensor pair.

**Figure 9.**
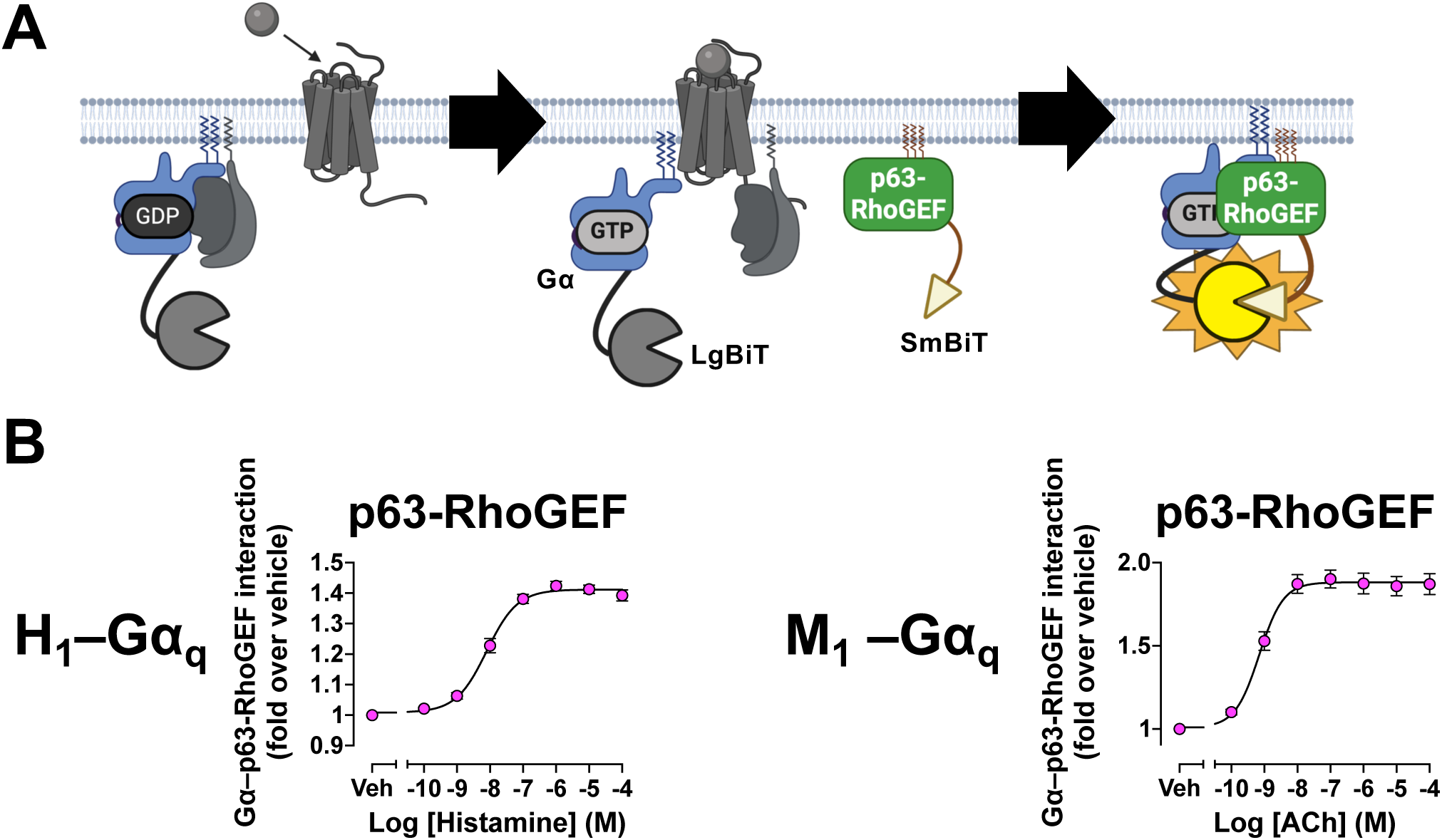
The Gα_q_–p63-RhoGEF association assay. (A) Schematic of the Gα_q_–p63-RhoGEF interaction assay. Gα_q_-LgBiT and SmBiT-p63-RhoGEF detect agonist-induced proximity between activated Gα_q_ and p63-RhoGEF. (B) Concentration–response curves for the Gα_q_–p63-RhoGEF interaction at the H_1_ and M_1_ receptors. Symbols and error bars represent the mean ± SEM of three independent experiments, each performed in duplicate.

### Evaluation of Gα_12/13_ protein activity via Gα_12/13_–RhoGEF association assay

Upon activation, Gα_12/13_ proteins interact with Rho guanine nucleotide exchange factors (RhoGEFs), promoting the exchange of GDP for GTP on Rho GTPases and thereby triggering downstream responses such as cytoskeletal remodeling [75]. Gα_12/13_ proteins are known to engage three major RGS-containing RhoGEFs—p115-RhoGEF, PDZ-RhoGEF, and LARG [76]. Selective engagement of Gα_12/13_ proteins with individual RhoGEF subtypes is an important determinant of downstream signaling.

To develop NanoBiT sensors for these interactions, SmBiT was fused to the extreme N-terminus of each RhoGEF. Both p115-RhoGEF and PDZ-RhoGEF interact with Gα_12/13_ primarily through their RGS domains located in the N-terminal region [77]. We therefore reasoned that positioning SmBiT at the N-terminus would place the tag in close proximity to the Gα_12/13_-bound tag while minimizing possible steric hindrance. LARG also interacts with Gα_13_ through its N-terminal RGS domain, but additional interaction interfaces have been reported within the DH/PH module and C-terminal region [78]. We therefore placed SmBiT at the extreme N-terminus of LARG to minimize potential interference with these additional binding regions while maintaining a sensor configuration consistent with those used for p115-RhoGEF and PDZ-RhoGEF.

To evaluate interactions between Gα_12/13_ proteins and the engineered RhoGEF sensors, we paired SmBiT-tagged ARHGEF1, ARHGEF11, and ARHGEF12, corresponding to p115-RhoGEF, PDZ-RhoGEF, and LARG, respectively, with LgBiT-fused Gα_12/13_ constructs originally developed for the G-protein dissociation assay (Fig. 10A). Following stimulation of the thromboxane A_2_ receptor (TP), a receptor that efficiently activates Gα_12/13_ signaling, we detected ligand-dependent interactions between Gα_13_ and all three RhoGEFs (Fig. 10B, S13A). Responses were modest for PDZ-RhoGEF and LARG, whereas the interaction with p115-RhoGEF was substantially more robust. By contrast, Gα_12_ displayed a more restricted interaction profile, with only a modest response detected for p115- RhoGEF. The distinct kinetics observed for the interactions of Gα_13_ with p115-RhoGEF and LARG were consistent with a previous FRET-based study showing that Gα_13_ dissociates more slowly from LARG than from p115-RhoGEF [19]. The limited responses observed with Gα_12_ may reflect a greater dependence on cellular context and additional regulatory mechanisms. For example, efficient activation of LARG by Gα_12_ has been reported to require tyrosine phosphorylation of LARG [79], whereas the interaction of Gα_12_ with p115-RhoGEF has previously been described as relatively weak compared with that of Gα_13_ [80]. Thus, the restricted interaction profile observed for Gα_12_ is broadly consistent with previous studies. Overall, these results support the functionality of the NanoBiT-based RhoGEF sensors and reveal distinct interaction profiles within the Gα_12/13_ family. Under the conditions tested, Gα_13_ engaged multiple RhoGEF subtypes, whereas detectable Gα_12_ coupling was largely restricted to p115-RhoGEF.

**Figure 10.**
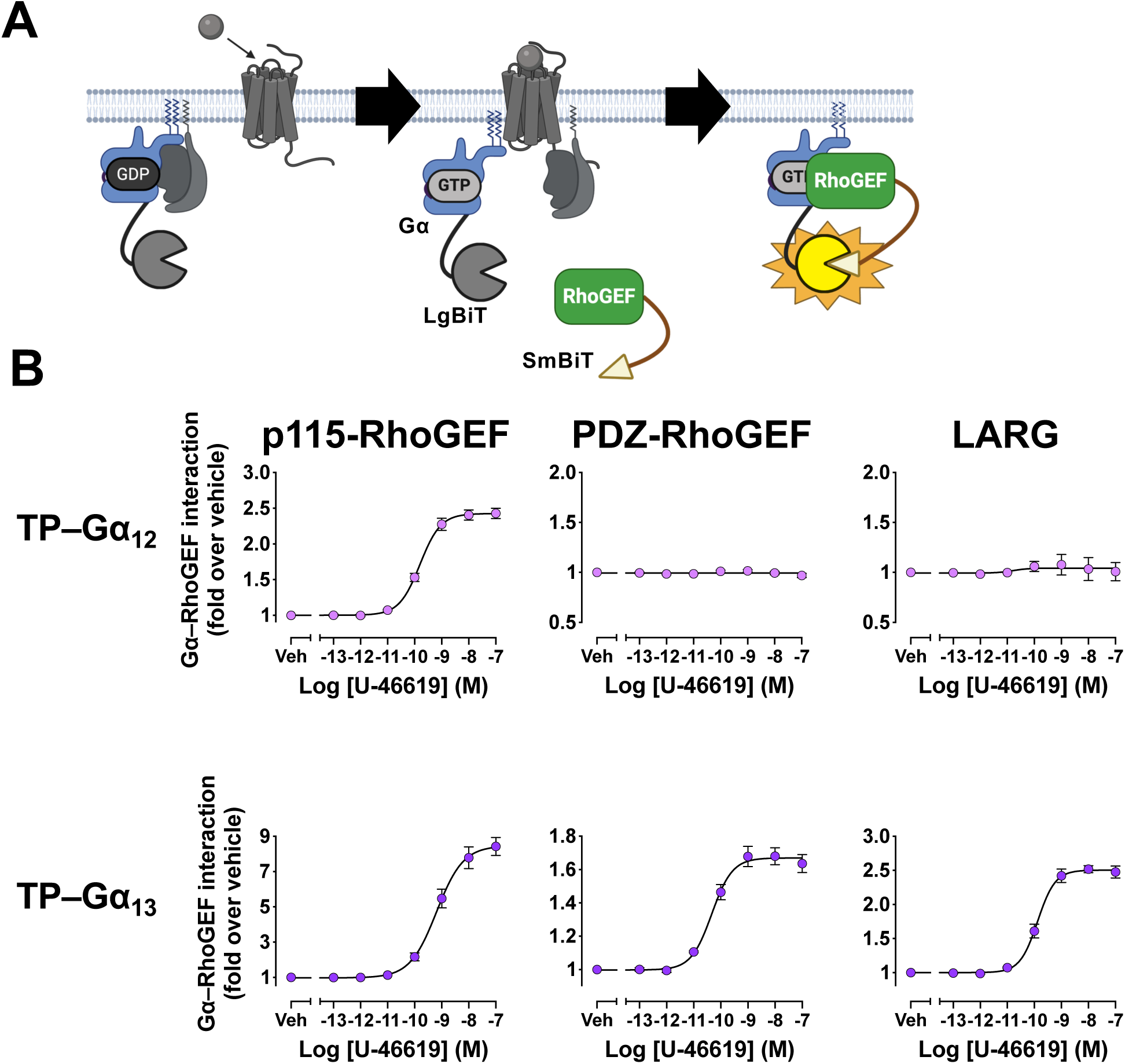
The Gα_12_–RhoGEF assay. (A) Schematic of the Gα_12/13_–RhoGEF interaction assay. Gα_12/13_-LgBiT and SmBiT-RhoGEF detect agonist-induced recruitment of the RGS-containing RhoGEFs p115-RhoGEF, PDZ-RhoGEF, and LARG to activated Gα_12/13_. (B) Concentration–response curves for interactions of Gα_12_ and Gα_13_ with p115-RhoGEF, PDZ-RhoGEF, and LARG at the TP receptor. Symbols and error bars represent the mean ± SEM of three independent experiments, each performed in duplicate.

### Detecting inter-effector associations

We next investigated whether the NanoBiT platform could be extended to detect interactions among downstream signaling components. As an initial model, we examined the association of RhoA with RhoGEFs. Such interactions have traditionally been assessed using biochemical approaches such as pull-down assays [81], prompting us to develop a higher-throughput live-cell assay based on NanoBiT complementation.

To generate a RhoA sensor, LgBiT was fused to the extreme N-terminus of RhoA. RhoGEFs engage the Switch I and Switch II regions of RhoA [82,83], and the sensor design was adapted from previously reported split-firefly-luciferase RhoA constructs by replacing the firefly luciferase fragments with NanoBiT fragments [84]. Using the SmBiT-tagged RhoGEF constructs developed for the Gα_12/13_–RhoGEF interaction assay, we then evaluated ligand-dependent interactions of RhoA with p115-RhoGEF, PDZ-RhoGEF, and LARG (Fig. 11A). No detectable responses were observed with PDZ-RhoGEF or LARG, whereas p115-RhoGEF produced a modest ligand-dependent response (Fig. 11B; S14A).

**Figure 11.**
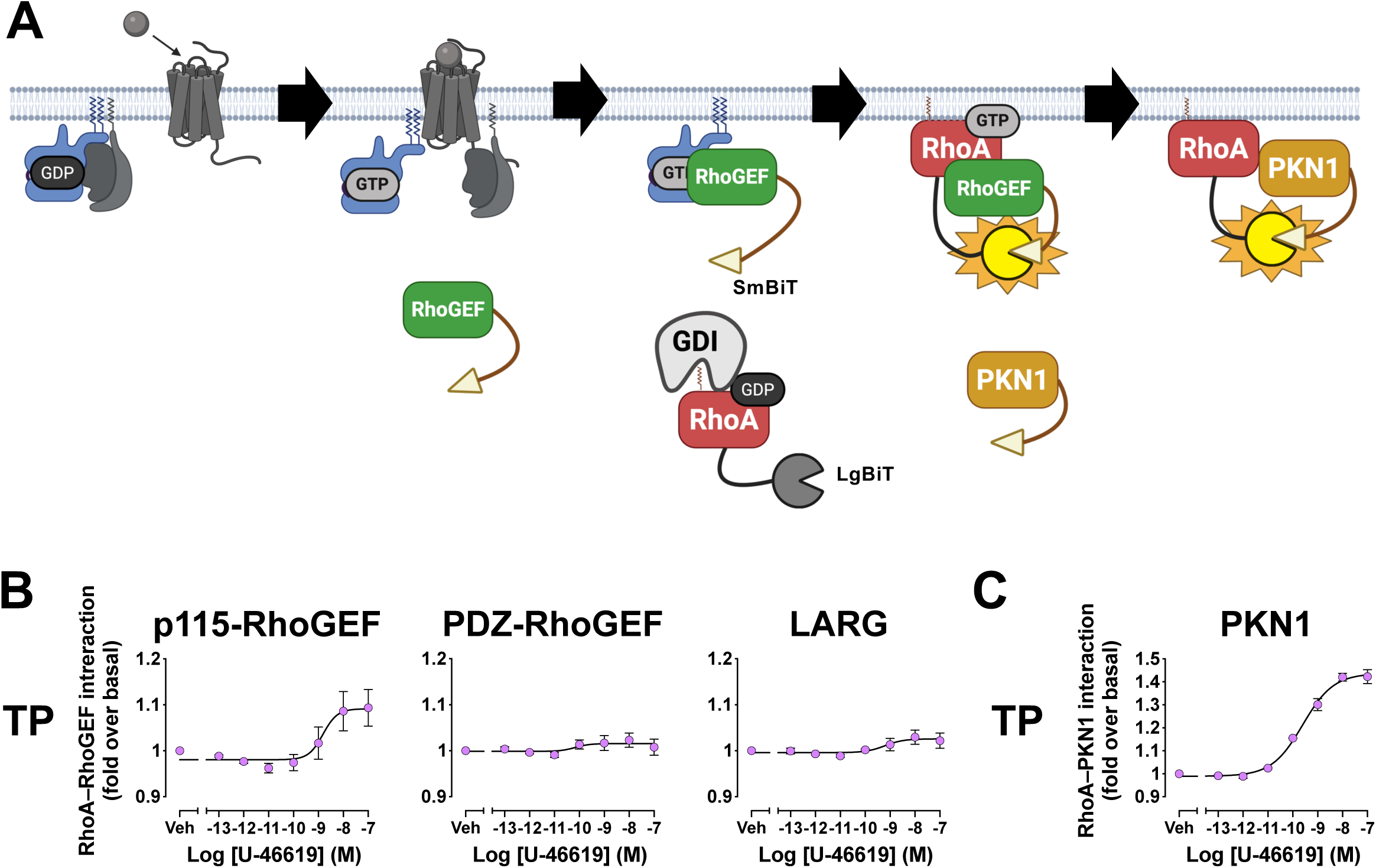
Inter-effector NanoBiT assays for the Gα_12/13_–RhoA pathway. (A) Schematic of the assays monitoring interactions downstream of Gα_12/13_. The RhoA–RhoGEF assay reports recruitment of RhoA to activated RhoGEFs, and the RhoA–PKN1 assay reports engagement of active RhoA by the RhoA-binding domain of the effector PKN1 (PKN1-GBD). (B) Concentration–response curves for RhoA interactions with p115-RhoGEF, PDZ-RhoGEF, and LARG at the TP receptor. Symbols and error bars represent the mean ± SEM of three independent experiments, each performed in duplicate. (C) Concentration–response curve for the RhoA–PKN1-GBD interaction at the TP receptor. Symbols and error bars represent the mean ± SEM of three independent experiments, each performed in duplicate.

We further examined RhoA activity by monitoring the interaction of activated RhoA with its downstream effector protein kinase N1 (PKN1). For this purpose, we used the GTPase-binding domain of PKN1, hereafter referred to as PKN1-GBD, which selectively recognizes the active, GTP-bound form of RhoA. As previously described [27], SmBiT was fused to the extreme N-terminus of PKN1-GBD, and its interaction with LgBiT–RhoA was measured following activation of the TP. Ligand stimulation produced a robust increase in NanoBiT luminescence, demonstrating that the sensor can detect receptor-induced RhoA activation through its association with PKN1-GBD (Fig. 11C, S14B). These results validate the functionality of the NanoBiT-based RhoA activity assay.

## Discussion

Here, we describe the construction and optimization of NanoBiT-based assays for monitoring signaling events downstream of a broad range of GPCRs. Although several protein–protein interaction assays have previously been developed to examine individual components of GPCR signaling, systematic evaluation across multiple receptors, transducers, and downstream signaling proteins has remained limited. Our assay platform enables the monitoring of G-protein dissociation, β-arrestin recruitment and trafficking, and GRK recruitment. We further extended the platform to detect interactions between G proteins and their downstream effectors, as well as interactions among signaling proteins downstream of G-protein activation. In particular, the successful monitoring of interactions within the Gα_12/13_ signaling pathway highlights the expanded coverage and utility of this assay platform.

Engineering signaling proteins to stabilize otherwise transient molecular interactions can substantially improve their detection using NanoBiT assays. GPCR signaling involves numerous short-lived interactions, including the direct engagement of activated receptors with G proteins and GRKs. Mini-G proteins are widely used to assess GPCR–G-protein coupling because they are engineered to form stable complexes with activated receptors (Nehmé et al, 2017). A complete panel of Venus-tagged mini-G proteins has been applied in BRET assays using Rluc8-tagged GPCRs, enabling direct detection of receptor coupling while largely preserving G-protein-subfamily selectivity. This strategy has subsequently been adapted to NanoBiT-based assays, further demonstrating its utility for monitoring direct GPCR–G-protein coupling (Wan *et al*, 2018; Mönnich *et al*, 2024). A conceptually similar strategy can be applied to GRKs. GRK-subtype selectivity has often been inferred indirectly by measuring β-arrestin recruitment in cells lacking individual GRKs, based on the requirement for GRK-mediated receptor phosphorylation in β-arrestin engagement [18,85]. Although such approaches have provided important insights into GRK-dependent receptor regulation, they do not directly measure interactions between a receptor and a specific GRK subtype. In our assay, though it depends on the condition, kinase-dead GRK mutants produced more sustained receptor-associated NanoBiT responses than their wild-type counterparts. This effect may reflect stabilization of the receptor–GRK complex by preventing receptor phosphorylation and the subsequent progression of the signaling complex toward GRK disengagement and β-arrestin recruitment. Thus, stabilizing mutations are a practical strategy that can prolong transient signaling interactions and enable their robust detection using NanoBiT assays.

Membrane-targeted markers can be used to assess the ability of GPCR transducers to translocate to, or interact within, membrane microdomains. Such membrane environments are increasingly recognized as important regulators of GPCR–transducer function and their lateral diffusion [86–88]. In the present study, we demonstrate that our assay platform can detect β-arrestin recruitment to membrane markers associated with both lipid-raft and non-raft regions. However, the current assay configuration is limited to determining whether β-arrestin can access these broad membrane domains and does not identify the specific lipid compositions that are preferentially associated with β-arrestin recruitment or required for its function. Expanding the platform to include markers that selectively recognize defined lipid membrane environments, together with conformation-selective nanobodies for activated transducers, may enable a more detailed analysis of how membrane lipid composition regulates transducer localization and function.

Another important advantage of our approach is its ability to resolve the time course of protein–protein interactions. Kinetic measurements can provide information that is not accessible from endpoint assays, including differences in the onset, duration, and termination of signaling. Such temporal distinctions have been reported for G-protein activation downstream of class A and class B GPCRs [89]. By contrast, kinetic analysis of Gα_12/13_ signaling has remained comparatively limited, largely because few tools are available for monitoring this pathway in real time. Gα_12/13_ activity has traditionally been evaluated using endpoint measurements of RhoA activation, including Rhotekin-based pull-down assays and G-LISA [90–92]. Although these approaches provide robust measurements of pathway activation at selected time points, they do not directly capture the continuous kinetics of signaling and therefore provide limited information about the duration, reversibility, and temporal contribution of individual molecular interactions. More recently, live-cell imaging approaches have begun to address this limitation. For example, a FRET-based strategy was developed to visualize conformational changes associated with RhoA activation in real time [93]. Our NanoBiT platform provides a complementary approach by enabling continuous monitoring of receptor-dependent interactions involving Gα_12/13_, RhoGEFs, and downstream RhoA signaling components. Together, these emerging methods should facilitate a more detailed understanding of the temporal organization of GPCR signaling.

The distinct activities observed among different heterotrimeric G-protein combinations may reflect their physiological significance. For example, knockdown of Gβ_1_ in zebrafish embryos was reported to markedly impair cardiac contractility [94]. This phenotype was accompanied by reduced Gα_s_ protein expression and a consequent decrease in cAMP production, suggesting that Gβ_1_ contributes to the stability or function of Gα_s_-containing heterotrimers in vivo. Consistent with this observation, our G-protein dissociation assay showed that Gβ_1_ exhibited high specificity with Gα_s_ proteins relative to other Gα protein subfamilies. This result raises the possibility that Gβ_1_ has functional importance in Gα_s_-mediated signaling. More generally, the subtype-dependent responses observed for different Gβ and Gγ combinations may reflect physiologically relevant differences in heterotrimer assembly, stability, localization, or signaling efficiency.

Expression profiles of GPCR signaling components may provide a basis for reconstructing pathway usage from omics data. In principle, ligand-induced transcriptomic or proteomic changes could be integrated with information on receptor expression, Gα-, Gβ-, Gγ-, and β-arrestin-subtype abundance, and their known functional compatibilities to infer which transducer pathways are preferentially engaged in a given cellular context. A conceptually related strategy has already been used to infer pathway activity from transcriptional signatures. For example, stimulation of P2Y2 and P2X4 receptors in mouse mesenchymal stromal cells induced expression of the immediate-early transcription factor Fos together with activation of ERK/MAPK signaling [95]. Similarly, gene ontology analysis of Fos-positive dentate granule cells from mice revealed enrichment of genes associated with MAPK signaling and neuronal activation [96]. These studies illustrate how pathway-associated transcriptional changes can be used to connect receptor stimulation with downstream signaling and cellular responses. By extending this concept to GPCR signaling components, RNA-sequencing or other omics-based analyses may help predict which Gα- or β-arrestin-dependent pathways are activated by a given ligand–receptor pair and how those pathways contribute to the resulting cellular phenotype.

NanoBiT differs from other assay formats in several important respects. NanoBiT is not a ratiometric assay, and its absolute luminescence is therefore strongly influenced by sensor abundance, fragment complementation efficiency, and substrate availability [97]. This makes direct comparison of basal luminescence across different sensor pairs or expression conditions difficult. By contrast, BRET measurements are expressed as an acceptor-to-donor emission ratio, which partially normalizes for differences in total donor signal and probe abundance and can therefore provide a more robust basis for comparing basal states across conditions [98]. Further efforts to improve control over sensor stoichiometry have led to the development of platforms such as G-CASE and ONE-GO, in which multiple BRET sensor components are encoded on a single vector plasmid to promote more consistent relative expression [99,100]. Moreover, NanoBiT assays do not completely reflect the physiological conditions. NanoBiT sensors are usually overexpressed to maximize the detection potential of intermolecular interactions. Therefore, whether a certain signaling protein serves an important physiological role must be examined through in vivo investigations. A further limitation is that many NanoBiT configurations require direct fusion of LgBiT or SmBiT to the proteins of interest. Validation using orthogonal assays is therefore important for interpreting NanoBiT-based measurements.

In conclusion, the NanoBiT platform provides a robust and highly versatile framework for monitoring multiple layers of GPCR signaling. We demonstrate representative sensor combinations and assay conditions for each signaling event examined. By applying a common detection principle across diverse signaling processes, this platform enables parallel evaluation of multiple GPCR signaling layers without requiring separate assay formats for each event. We further demonstrated that the platform can be extended beyond plate-based measurements to bioluminescence microscopy, enabling spatial visualization of GPCR signaling processes in living cells. Microscopic analysis can therefore serve as a valuable approach for validating changes in subcellular localization and confirming the spatial basis of luminescence responses. Overall, applying this NanoBiT-based platform across diverse GPCRs and cellular contexts should facilitate systematic mapping of the GPCR signalome and improve our understanding of how receptor-, transducer-, and effector-level interactions collectively shape cellular responses.

### Cloning

All plasmids used in this study were generated in the pCAGGS expression vector, unless otherwise stated. GPCR expression plasmids were reported previously (Inoue *et al*, Cell 2019), except for following constructs. M_1_ was N-terminally fused with a hemagglutinin-derived signal sequence, a FLAG epitope tag, and a 15-amino-acid flexible linker (ssHA-FLAG; MKTIIALSYIFCLVFA<u>DYKDDDDK</u>GGSGGGGSGGSSSGGG; FLAG epitope tag underlined). The same N-terminal ssHA-FLAG tag was introduced into the V_2_ construct carrying a C-terminal SmBiT fusion (V_2_-SmBiT), in which V2 and SmBiT was the 15-amino-acid flexible linker described above. For the Gα-LgBiT constructs, ORFs encoding Gα proteins with LgBiT inserted at the positions indicated in Table 1 were cloned into pcDNA3.1. The LgBiT sequence was flanked on both sides by the 15-amino-acid flexible linker. For the other LgBiT- or SmBiT-fused constructs, the NanoBiT fragments were fused to the N or C terminus via the 15-amino-acid flexible linker. Defined regions or partial sequences rather than full-length ORFs were used for some constructs; the full nucleotide sequences of all constructs are provided in Supplementary Information.

### Cell culture

HEK293A cells (Thermo Fisher Scientific) were maintained in Dulbecco’s Modified Eagle Medium (DMEM; Shimadzu Diagnostics) supplemented with 5% fetal bovine serum (Sigma-Aldrich), 100 U/mL penicillin (Sigma-Aldrich), 100 µg/mL streptomycin (Gibco) and 2 mM L-glutamine (Gibco) at 37°C in a humidified incubator with 5% CO_2_.

### Transfection

Transfection was performed using a cationic polyethyleneimine solution (PEI Max, Polysciences). Typically, HEK293A cells were seeded in 6-well plates at a cell density of 2.0 × 10^5^ cells/mL in 2 mL of DMEM supplemented with 5% fetal bovine serum (Sigma-Aldrich), 100 U/mL penicillin (Sigma-Aldrich), 100 µg/mL streptomycin (Gibco) and 2 mM L-glutamine (Gibco). A transfection solution was mixed by combining plasmid solution diluted in 100 μL of Opti-MEM and 100 μL of Opti-MEM solution containing 5 μg of PEI. The transfected cells were further incubated for one day before being subjected to the assays as described below.

### NanoBiT assay

For the NanoBiT assay, plasmid transfection was performed in 6-well plates using the plasmid mixtures listed in Supplementary File XX. When the total amount of DNA was lower than 1 µg per well, the empty pCAGGS vector was added upto 1 µg. Twenty-four hours after transfection, the cells were harvested with 1 mL of Dulbecco’s phosphate-buffered saline (D-PBS) containing 0.53 mM EDTA, followed by the addition of 1 mL of Hank’s balanced salt solution (HBSS) containing 5 mM HEPES at pH7.4 (FUJIFILM Wako Pure Chemical). The cells were pelleted by centrifugation at 190 × g for 5 min and resuspended in 2 mL of assay medium consisting of HBSS supplemented with 5 mM HEPES (FUJIFILM Wako Pure Chemical) and 0.01% bovine serum albumin (BSA; SERVA). The cell suspension was dispensed into each well of a white 96-well plate (80 μL per well), followed by the addition of 20 μL of 50 µM coelenterazine (Angene) solution, and incubated at room temperature. After 2 h of incubation, 20 μL of 6× ligand solution diluted in assay medium was added to each well. Immediately thereafter, luminescence was measured using a SpectraMax L microplate reader (Molecular Devices) operated with SoftMax Pro software. Measurements were taken at 35-sec intervals for 15 or 30 min, or at 20-sec intervals for 10 min. Unless otherwise stated, dose–response curves were generated using luminescence values measured between 11 and 15 min after ligand addition.

### NanoBiT imaging

V_2_-SmBiT and LgBiT-βarr1 plasmids were transfected into parental cells seeded on coverslips using Lipofectamine 3000 one day before imaging. The amount of plasmid DNA was the same as that used for the NanoBiT assay. The coverslip (Matsunami, 25 mm round, No. 1) was mounted in an Attofluor cell chamber (Thermo Fisher Scientific), and the extracellular solution was replaced with 300 µL of HBSS. After buffer exchange, 100 µL of 10 µM furimazine solution was added before imaging. Immediately before image acquisition, 100 µL of ligand solution was added to the chamber on the microscope to stimulate the cells with AVP at a final concentration of 1 µM.

NanoBiT time-lapse images were taken by a homemade microscope system with the following configurations (microscope: LV200, Olympus, objective: UPLXAPO40XO 40×, NA 1.40, Olympus, imaging lens: 36 mm with 0.2× magnification, camera: MTR3CMOS09000KPA, ToupTek). Microscopy acquisition was controlled using ToupView software (ToupTek Photonics). Images were acquired as color images, and the blue channel was used for subsequent analysis.

Time-lapse images were acquired as 1496 ×1500-pixel images with a pixel size of 0.47 µm/pixel under the following settings: exposure time, 10 s; Gain, 10,000; time interval, 10 s; total acquisition time, 2.5 min. Image processing, including channel splitting and cropping of regions of interest, was performed using ImageJ.

---

**Figure S1.**
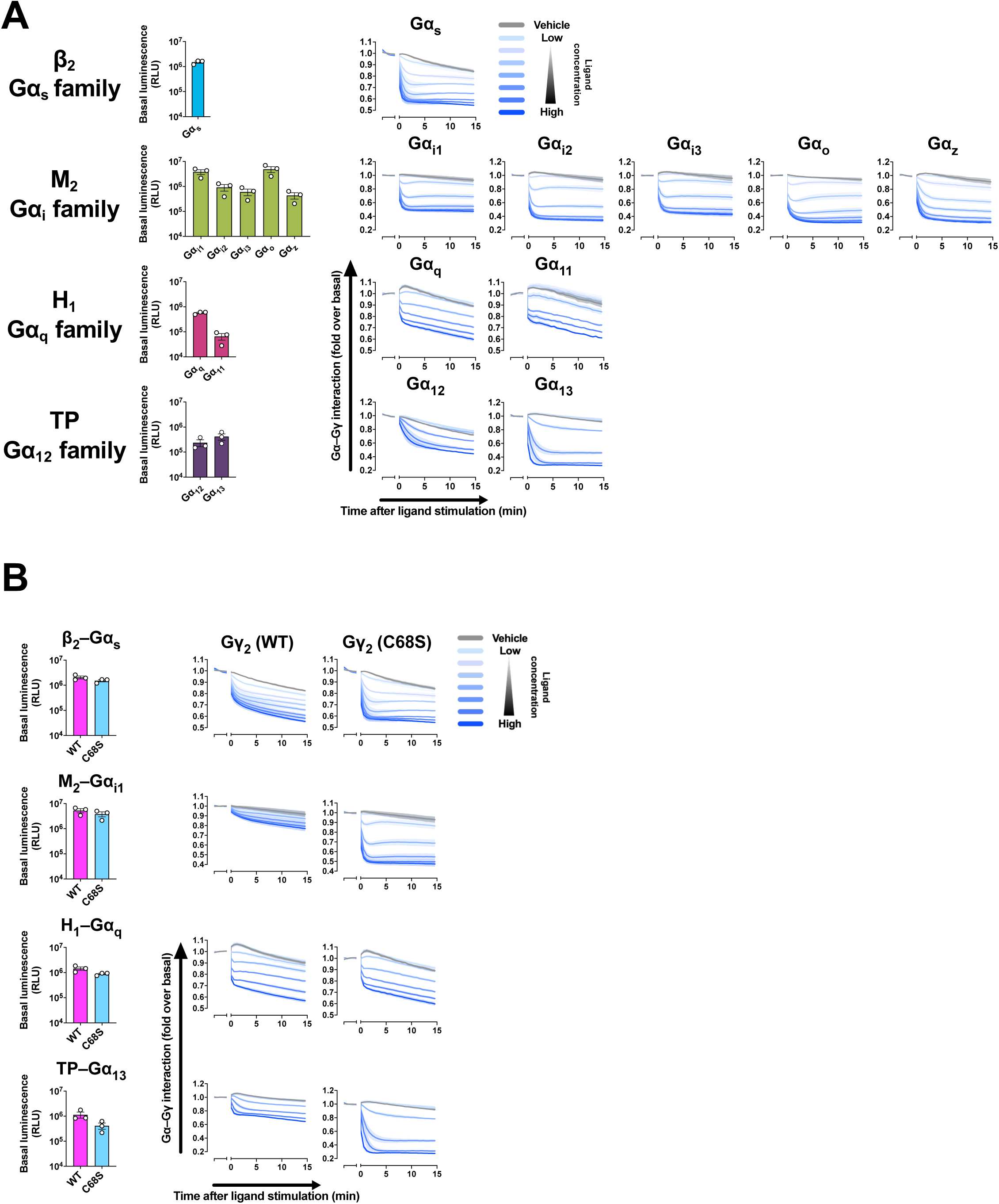
Basal luminescence and real-time kinetics of the Gα–Gβγ dissociation assay. (A) Basal luminescence and dissociation kinetics for representative GPCR–Gα coupling conditions spanning the four Gα subfamilies. Symbols and error bars represent the mean ± SEM of three independent experiments, each performed in duplicate. (B) Basal luminescence and dissociation kinetics using wild-type or C68S Gγ_2_. Symbols and error bars represent the mean ± SEM of three independent experiments, each performed in duplicate.

**Figure S2.**
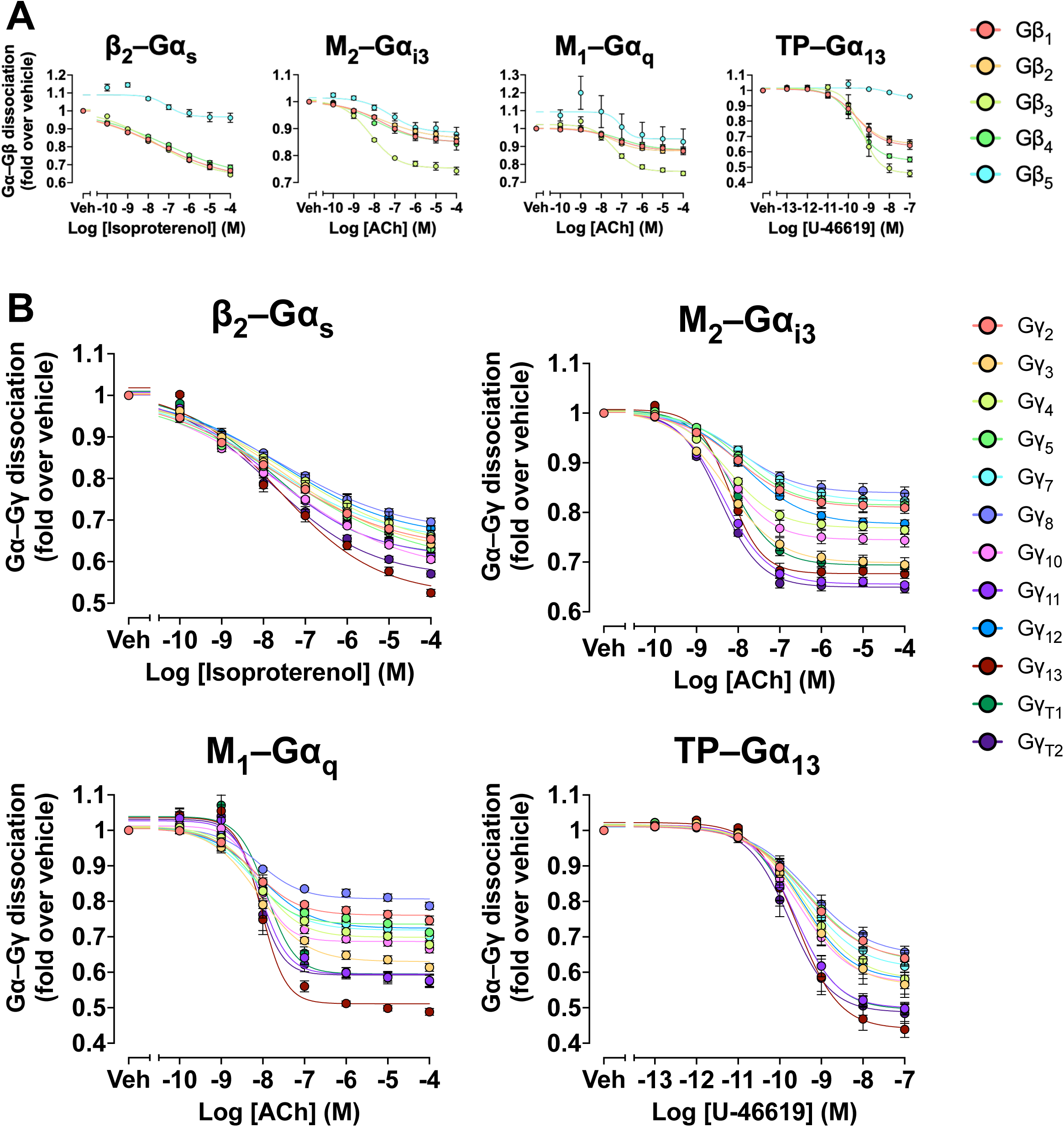
Concentration–response curves of the the Gβ- and Gγ-subtype profiling experiments. (A) Concentration–response curves for Gβ selectivity across five Gβ subunits. Symbols and error bars represent the mean ± SEM of three independent experiments, each performed in duplicate. (B) Concentration–response curves for Gγ selectivity across 12 Gγ subunits. Symbols and error bars represent the mean ± SEM of 3–4 independent experiments, each performed in duplicate.

**Figure S3.**
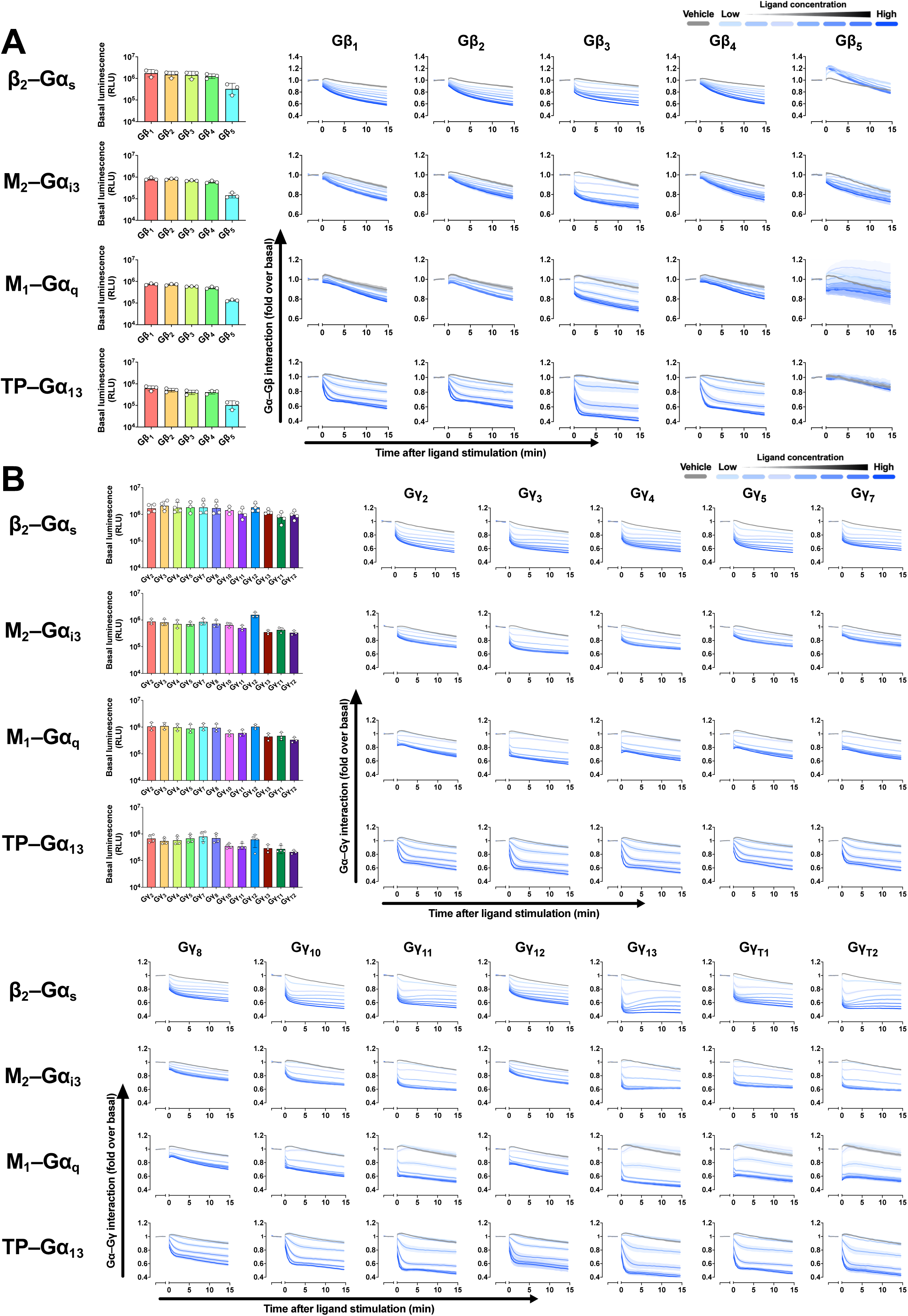
Basal luminescence and real-time kinetics of the Gβ- and Gγ-subtype profiling experiments. (A, B) Basal luminescence and dissociation kinetics for Gβ (A) and Gγ (B) subtype profiling by the Gα–Gβγ dissociation assay. Symbols and error bars represent the mean ± SEM of 3–4 independent experiments, each performed in duplicate.

**Figure S4.**
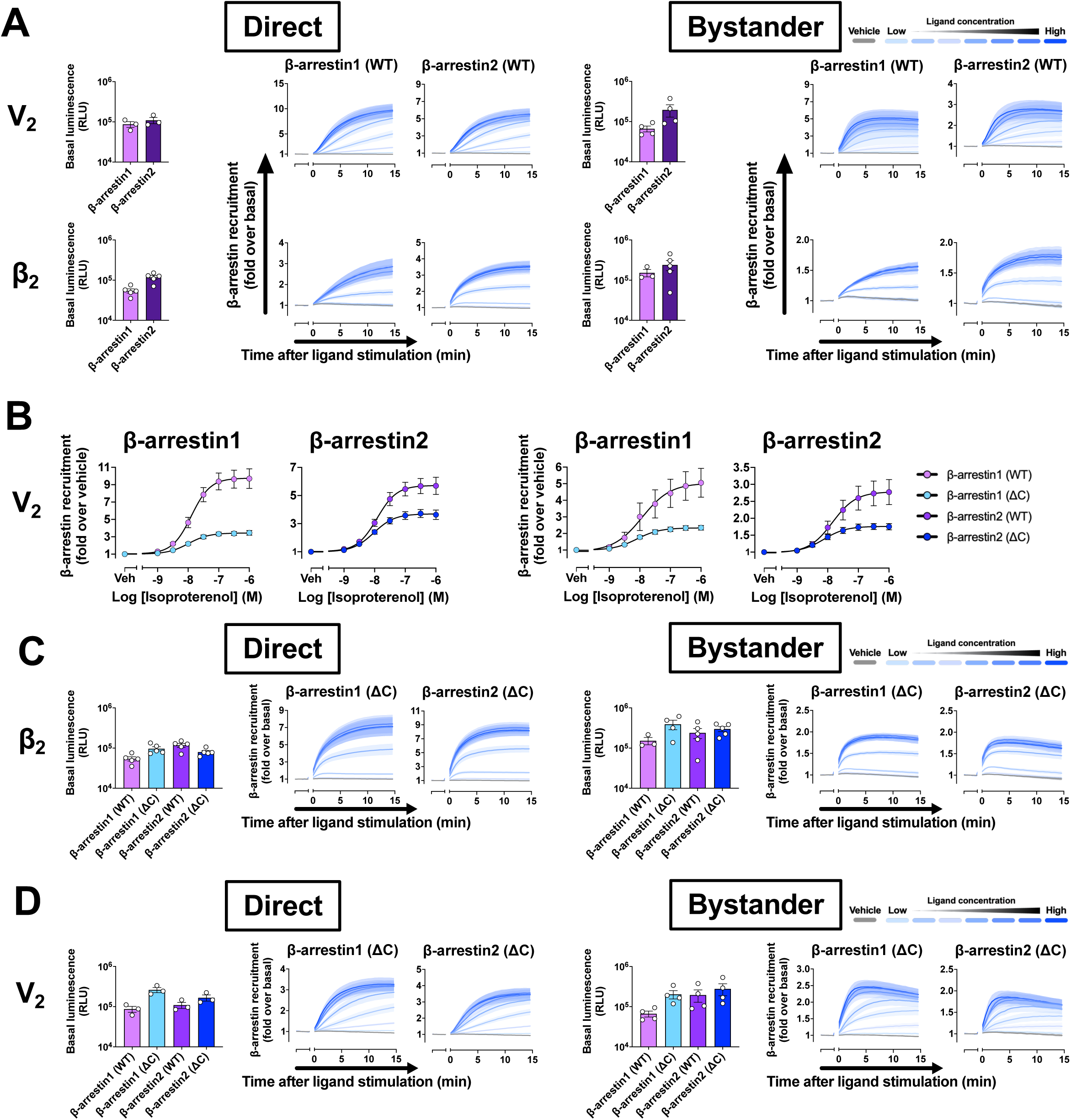
Basal luminescence and real-time kinetics of the direct and bystander β-arrestin recruitment assays. (A) Basal luminescence and recruitment kinetics for β-arrestin1 and β-arrestin2 in the direct (left) and bystander (right) assays. Symbols and error bars represent the mean ± SEM of 3–5 independent experiments, each performed in duplicate. (B) Concentration–response comparison of full-length β-arrestin and the β-arrestin(ΔC) mutant in the direct (left) and bystander (right) assays. The β-arrestin1 (WT) and β-arrestin2 (WT) data from Fig. 3B are replotted together with β-arrestin(ΔC) data acquired in the same experimental sets. Symbols and error bars represent the mean ± SEM of 3–5 independent experiments, each performed in duplicate. (C, D) Basal luminescence and recruitment kinetics for full-length β-arrestin and β-arrestin(ΔC) in the direct (left) and bystander (right) assays. Symbols and error bars represent the mean ± SEM of 3–5 independent experiments, each performed in duplicate.

**Figure S5.**
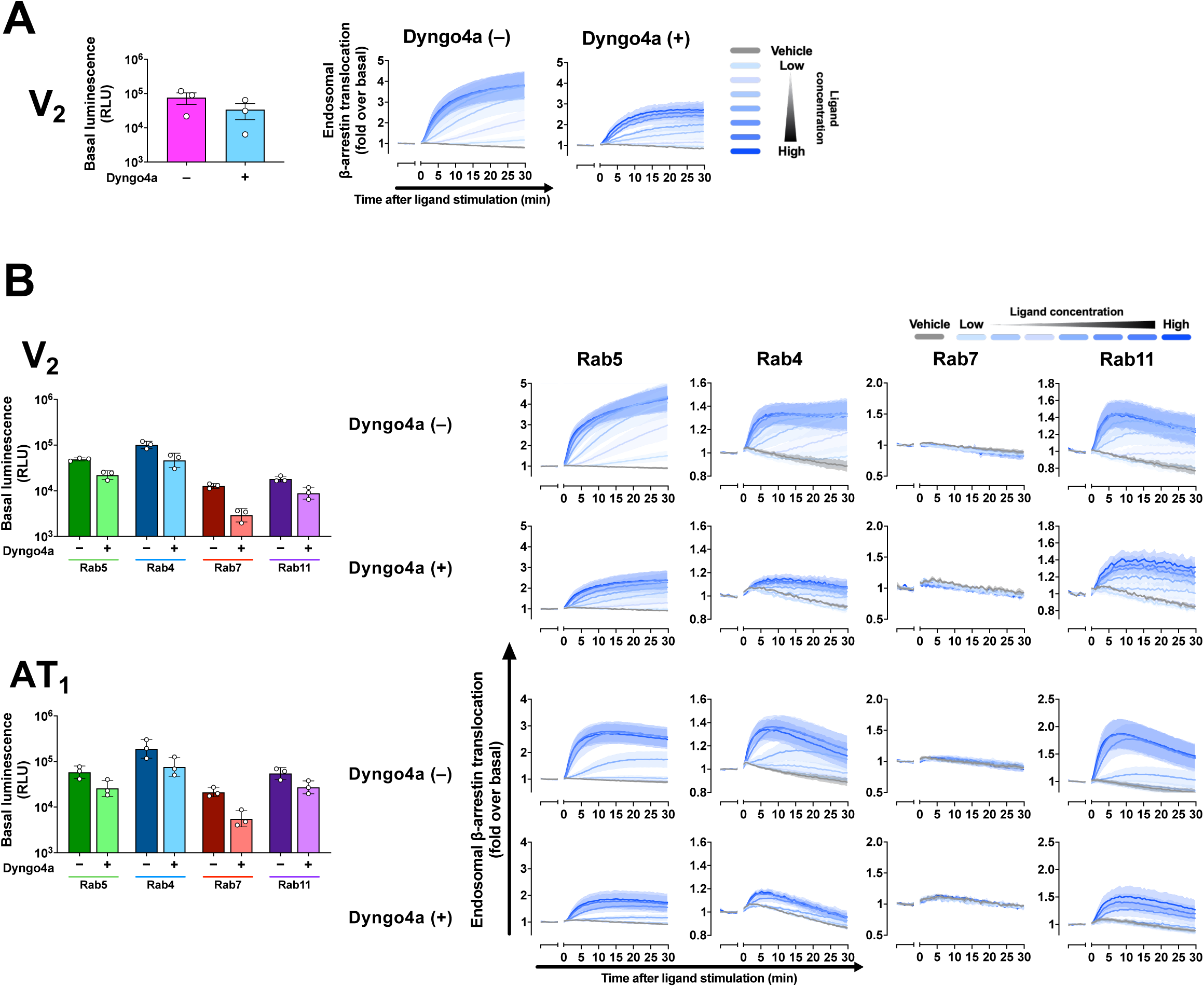
Basal luminescence and real-time kinetics of the β-arrestin endosomal translocation assay. (A–C) Basal luminescence and translocation kinetics of β-arrestin1 to Endofin-positive (A) and Rab-positive (B, C) endosomal compartments at the V_2_ and AT_1_ receptors, with or without Dyngo-4a. Symbols and error bars represent the mean ± SEM of three independent experiments, each performed in duplicate.

**Figure S6.**
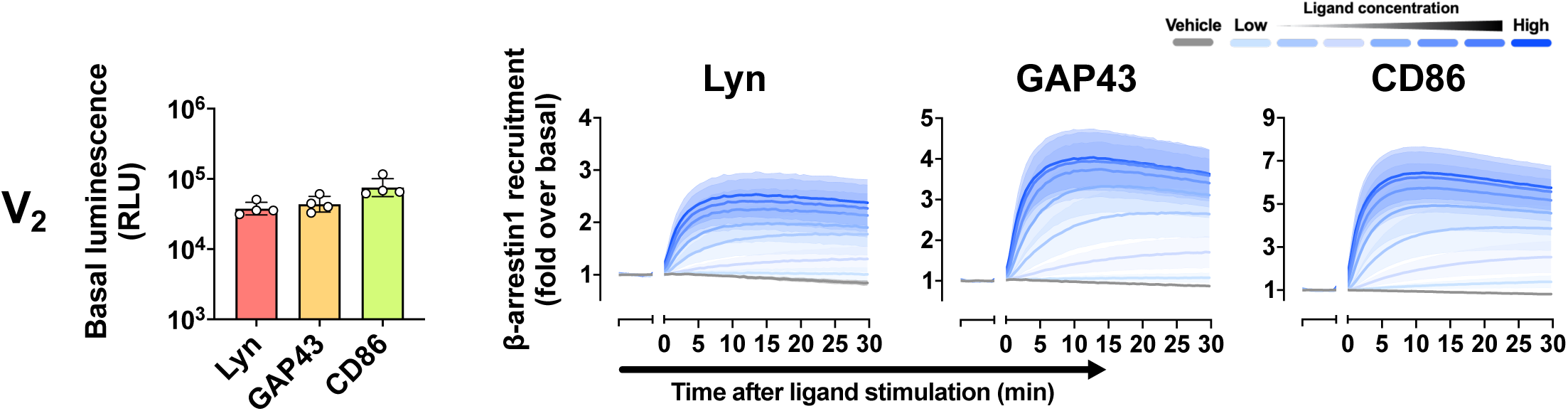
Basal luminescence and real-time kinetics of the β-arrestin membrane domain accumulation assay. Basal luminescence and recruitment kinetics of β-arrestin1 to Lyn-, GAP43-, and CD86-defined microdomains at the V_2_ receptor. Symbols and error bars represent the mean ± SEM of 3–4 independent experiments, each performed in duplicate.

**Figure S7.**
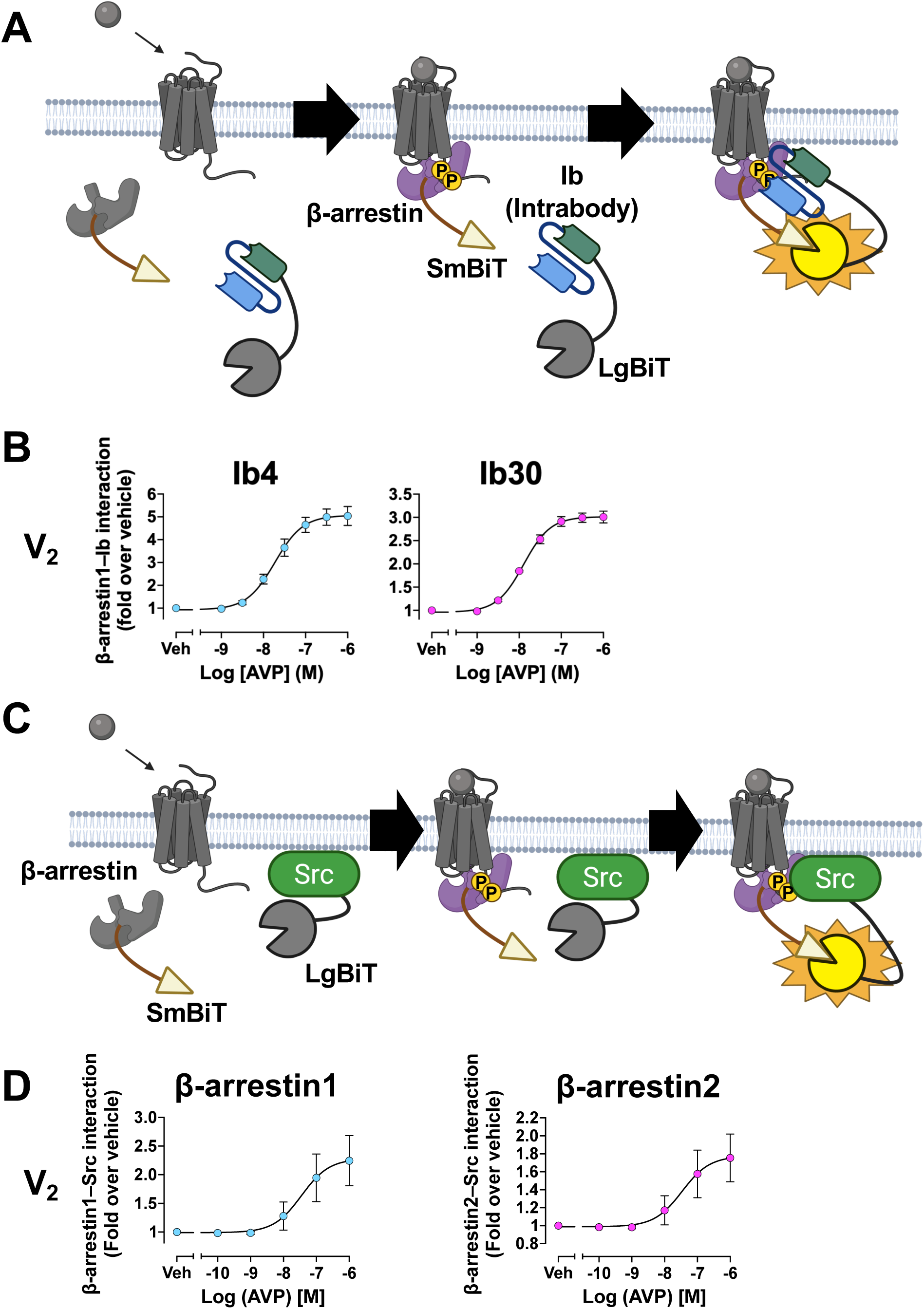
The β-arrestin conformational activation assay and β-arrestin–Src scaffolding assay. (A) Schematic of the intrabody (Ib)-based assay for detecting the active conformation of β-arrestin. LgBiT-fused intrabodies that recognize activated β-arrestin are used with SmBiT-β-arrestin; receptor activation drives intrabody engagement and increases luminescence. (B) Concentration–response curves for the intrabodies Ib4 and Ib30 with β-arrestin1 at the V_2_ receptor. Symbols and error bars represent the mean ± SEM of three independent experiments, each performed in duplicate. (C) Schematic of the β-arrestin–Src interaction assay, which detects agonist-induced proximity between β-arrestin and the scaffolded effector Src. (D) Concentration–response curves for β-arrestin1–Src and β-arrestin2–Src interactions at the V_2_ receptor. Symbols and error bars represent the mean ± SEM of three independent experiments, each performed in duplicate.

**Figure S8.**
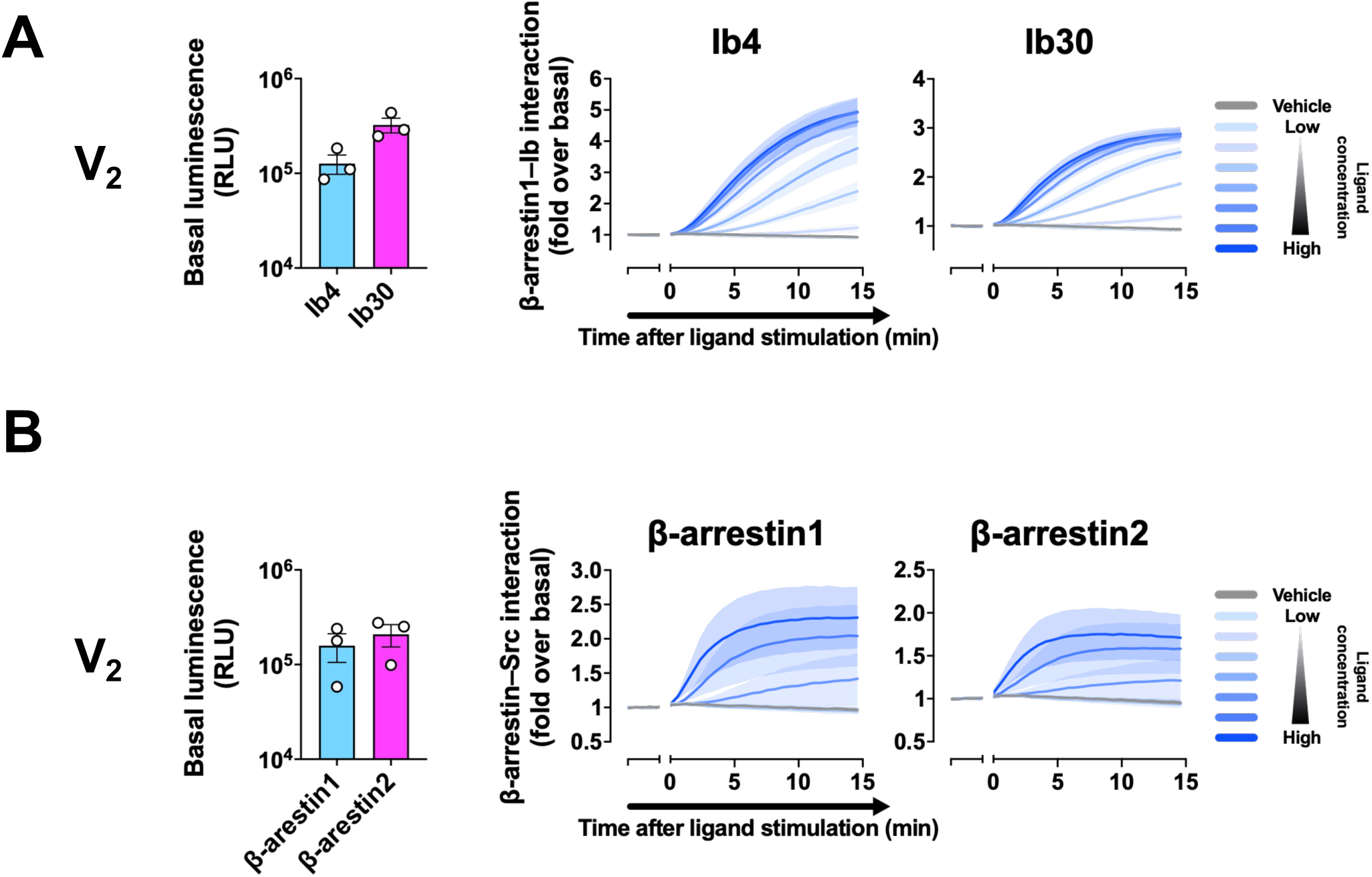
Basal luminescence and real-time kinetics of the β-arrestin conformational-activation and β-arrestin–Src assays. (A) Basal luminescence and kinetics of the intrabody-based conformational-activation assay using Ib4 and Ib30. Symbols and error bars represent the mean ± SEM of three independent experiments, each performed in duplicate. (B) Basal luminescence and kinetics of the β-arrestin1–Src and β-arrestin2–Src interaction assays. Symbols and error bars represent the mean ± SEM of three independent experiments, each performed in duplicate.

**Figure S9.**
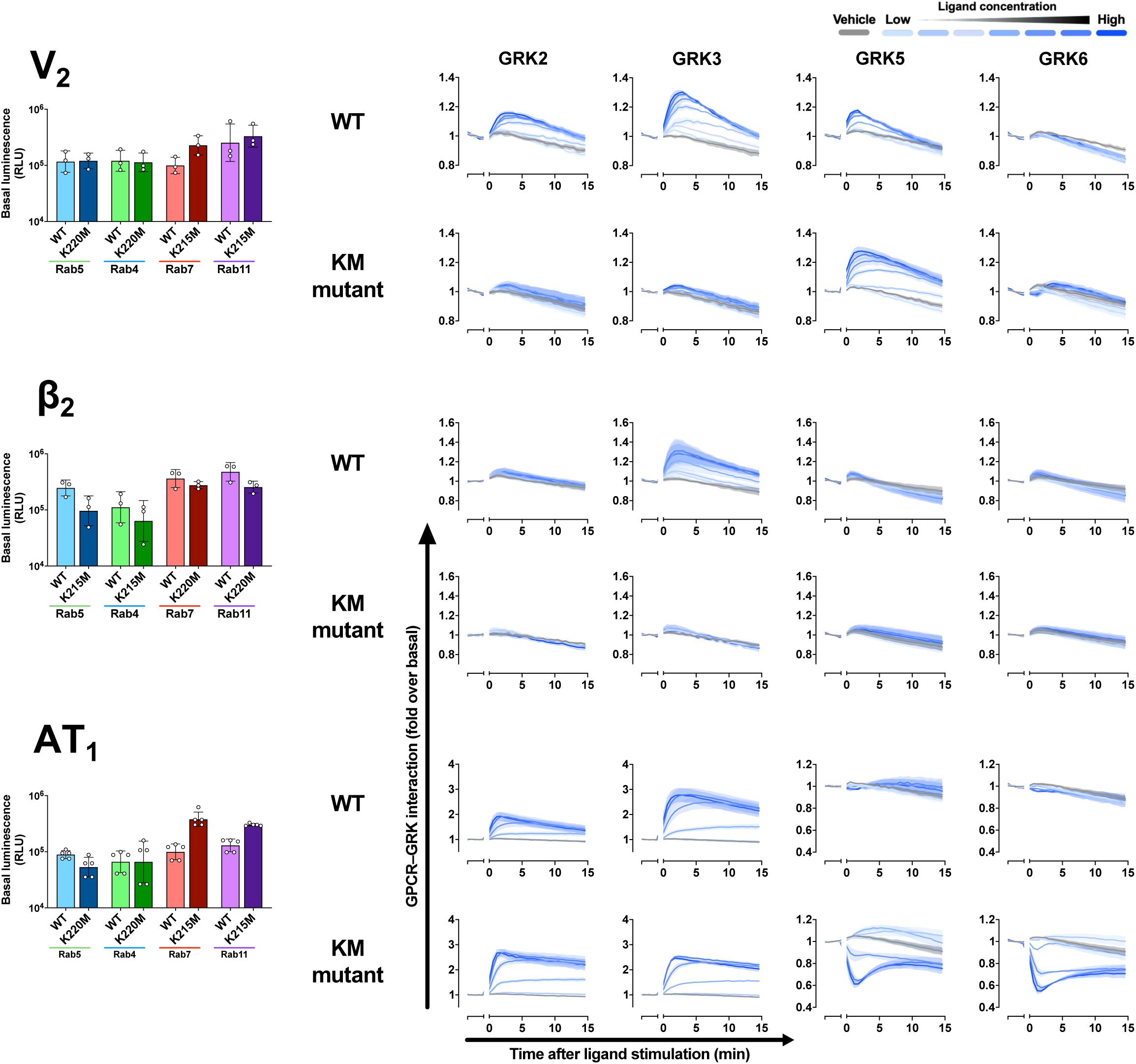
Basal luminescence and real-time kinetics of the GRK recruitment assay. Basal luminescence and recruitment kinetics for GRK2, GRK3, GRK5, GRK6, and their KM mutants at the V_2_, β_2_, and AT_1_ receptors. Symbols and error bars represent the mean ± SEM of 3–5 independent experiments, each performed in duplicate.

**Figure S10.**
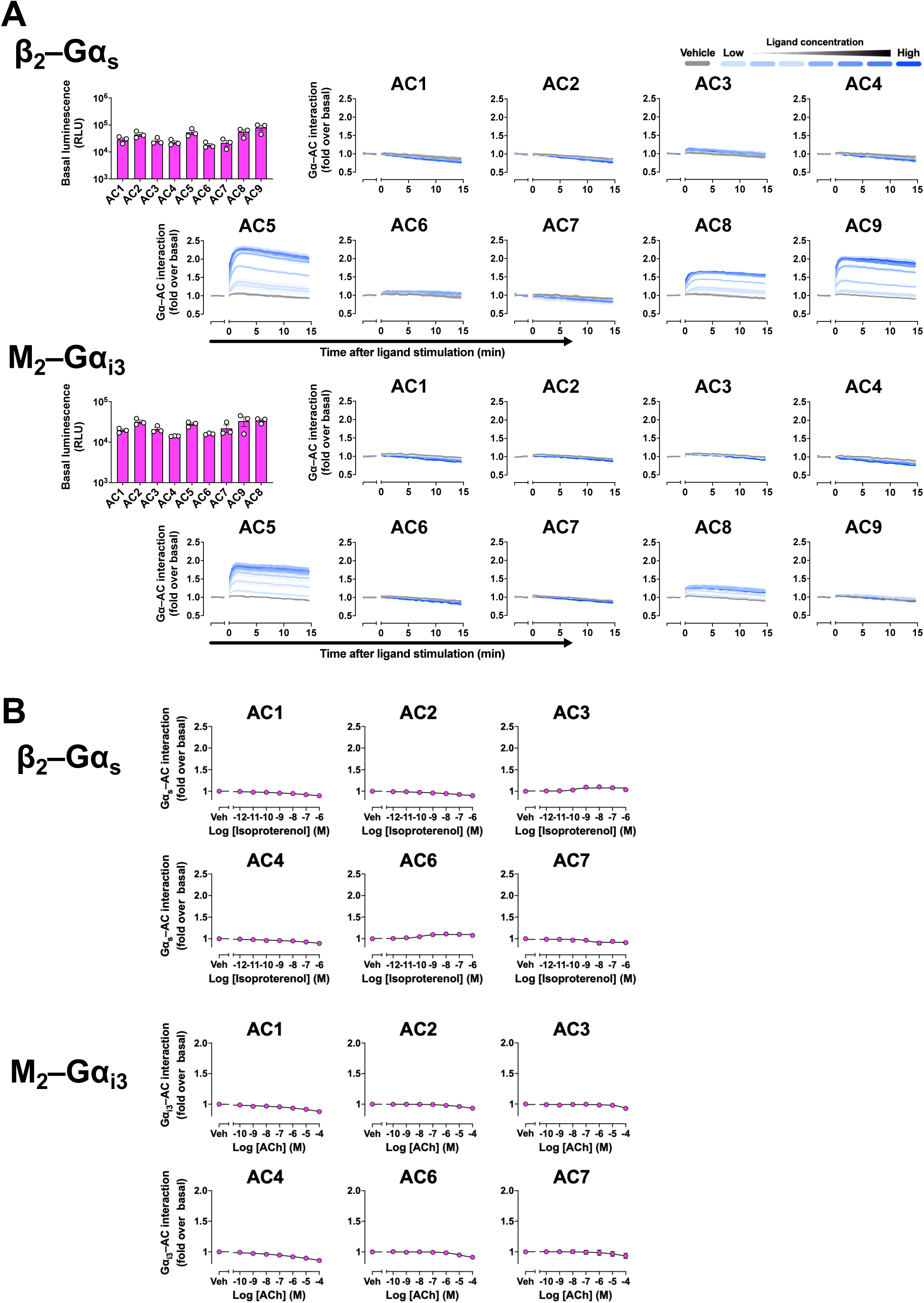
Extended profiling and real-time kinetics of the Gα–adenylyl cyclase association assay. (A) Concentration–response curves and basal luminescence for Gα_s_–AC and Gα_i3_–AC interactions across the AC isoforms other than the responsive AC5, AC8, and AC9 shown in Fig. 7B. Symbols and error bars represent the mean ± SEM of three independent experiments, each performed in duplicate. (B) Interaction kinetics of the Gα–AC assay across AC1–AC9. Symbols and error bars represent the mean ± SEM of three independent experiments, each performed in duplicate.

**Figure S11.**
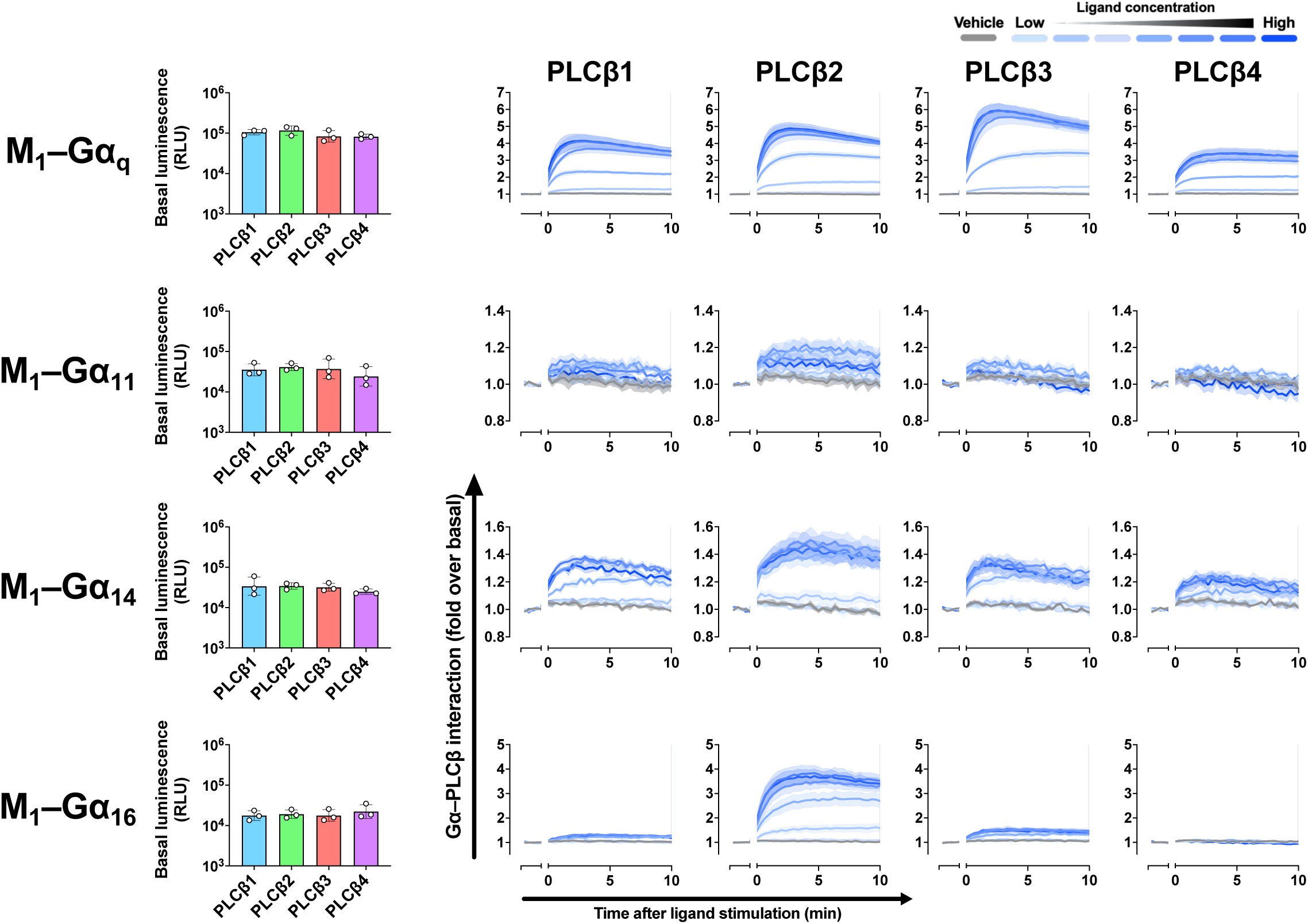
Basal luminescence and real-time kinetics of the Gα_q/_–PLCβ association assay. Basal luminescence and interaction kinetics for Gα_q_, Gα_11_, Gα_14_, and Gα_16_ with PLCβ1–PLCβ4 at the M_1_ receptor. Symbols and error bars represent the mean ± SEM of three independent experiments, each performed in duplicate.

**Figure S12.**
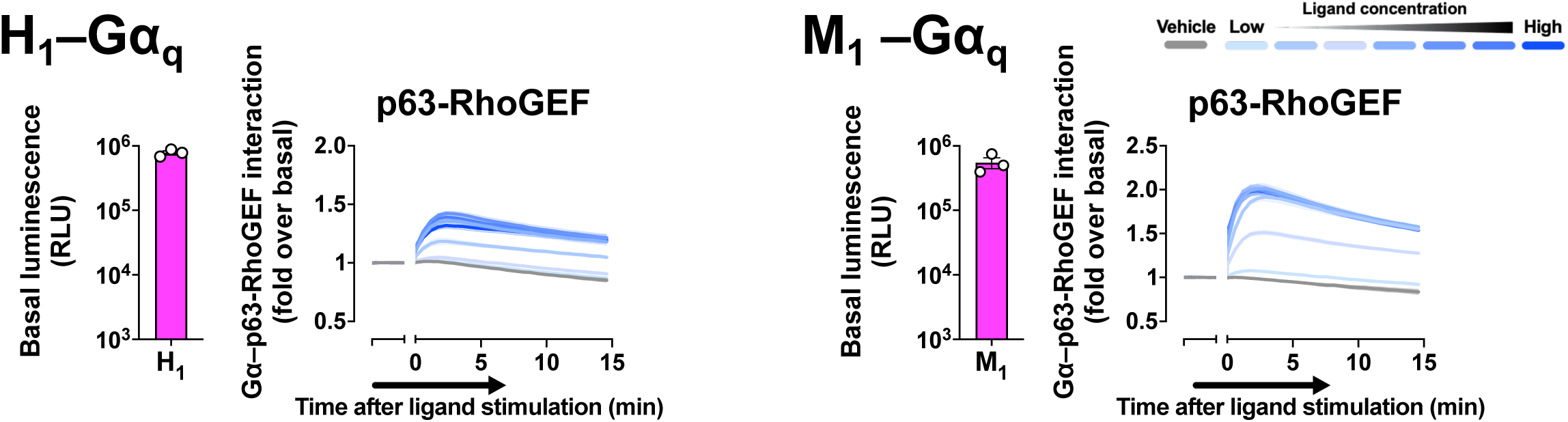
Basal luminescence and real-time kinetics of the Gα_q_–p63-RhoGEF association assay. Basal luminescence and interaction kinetics of the Gα_q_–p63-RhoGEF assay at the H_1_ and M_1_ receptors. Symbols and error bars represent the mean ± SEM of three independent experiments, each performed in duplicate.

**Figure S13.**
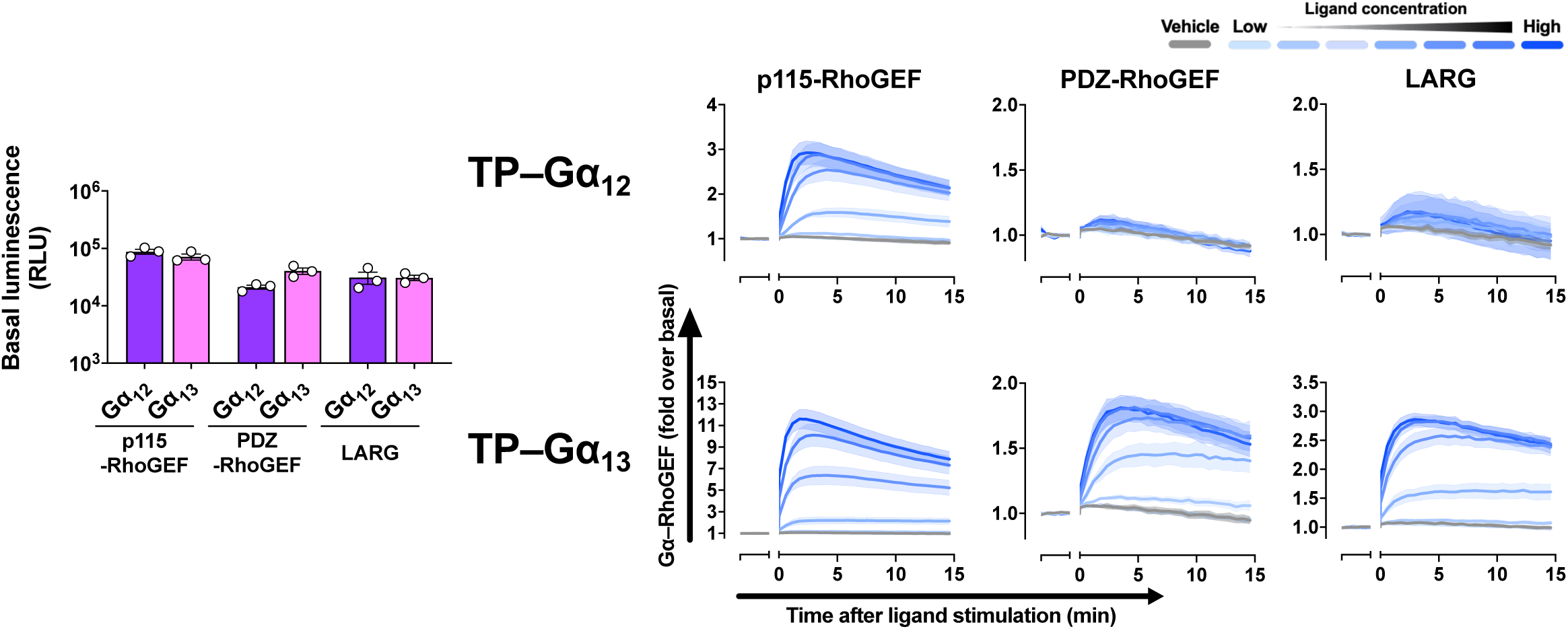
Basal luminescence and real-time kinetics of the Gα_12/13_–RhoGEF association assay. Basal luminescence and interaction kinetics for Gα_12_ and Gα_13_ with p115-RhoGEF, PDZ-RhoGEF, and LARG at the TP receptor. Symbols and error bars represent the mean ± SEM of three independent experiments, each performed in duplicate.

**Figure S14.**
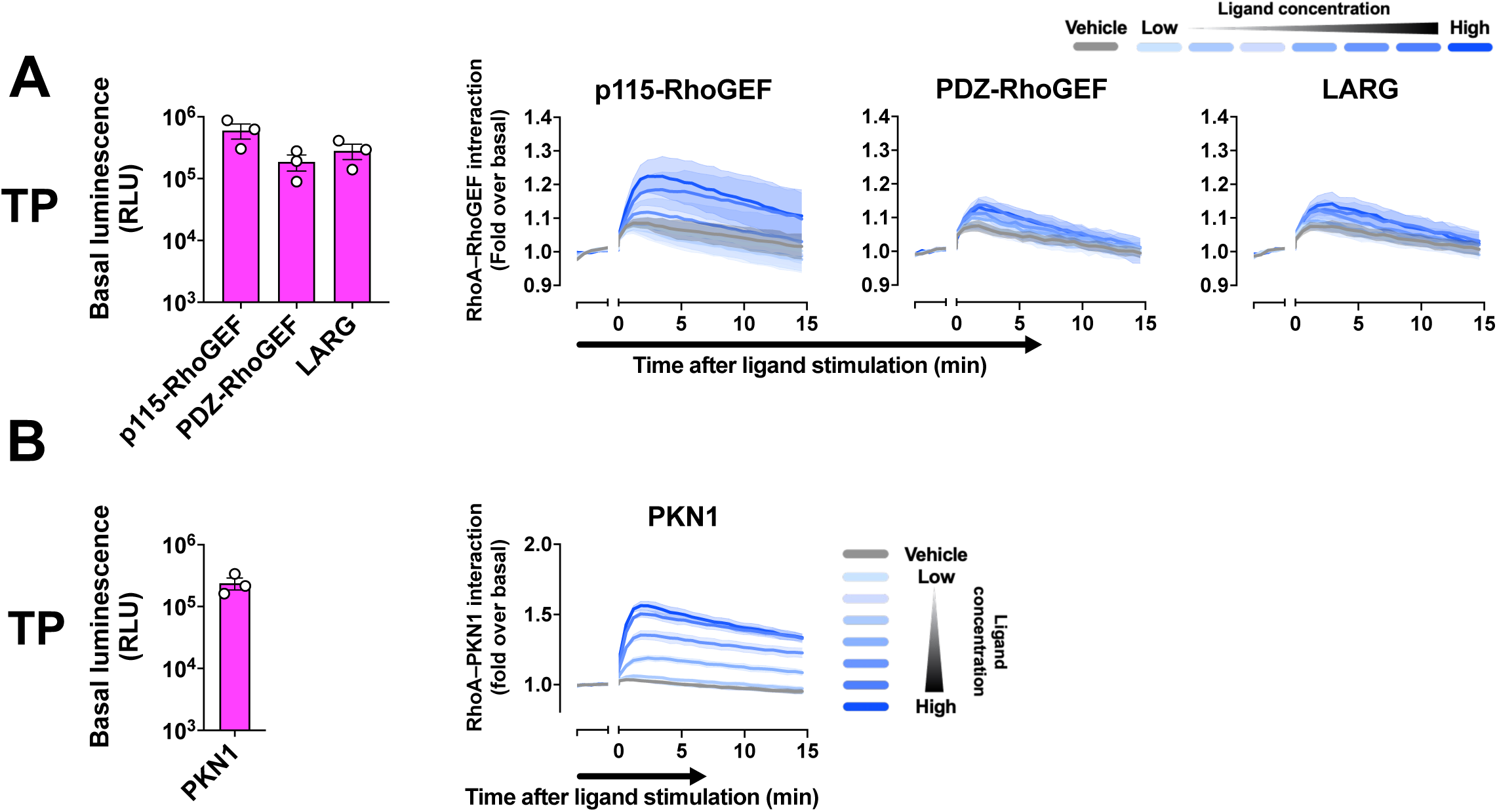
Basal luminescence and real-time kinetics of the RhoA–RhoGEF and the RhoA– PKN1 association assays. Basal luminescence and interaction kinetics of the RhoA–RhoGEF assay (p115-RhoGEF, PDZ-RhoGEF, and LARG) and the RhoA–PKN1-GBD assay at the TP receptor. Symbols and error bars represent the mean ± SEM of three independent experiments, each performed in duplicate.

## Acknowledgements

We thank Kouki Kawakami for helpful discussion of the initial project and members of the A.I. laboratory for helpful discussion and manuscript editing. This work was supported by KAKENHI JP24K21281 (A.I.), JP25H01016 (A.I.), JP24K01982 (M.Y.), JP24H01266 (M.Y.) and JP25H01328 (M.Y.) from the Japan Society for the Promotion of Science; JPMJFR215T (A.I.), JPMJMS2023 (A.I.), JPMJPR20EF (M.Y.) and JPMJMI22H5 (A.I.) from the Japan Science and Technology Agency; JP22ama121038 (A.I.) and JP22zf0127007 (A.I.) from the Japan Agency for Medical Research and Development; the Takeda Science Foundation (A.I.); the Uehara Memorial Foundation (A.I.); the Lotte Foundation (M.Y.); the Kobayashi Foundation (M.Y.).

## Author contributions

Conceptualization, AI; Methodology, AS, RK, AI; Investigation, SY, AS, MY, RS; Formal analysis, SY, AS, RK; Validation, SY, AS, RK; Visualization, SY, AS, RK; Writing - original draft, AS, RK, AI; Writing - review & editing, AS, RK, AI; Funding acquisition, AI; Supervision, RK, AI.

