## Supplementary Information Figures and Tables for "Comprehensive dissection of GPCR signaling using a NanoBiT-based platform"

### S1 Figure

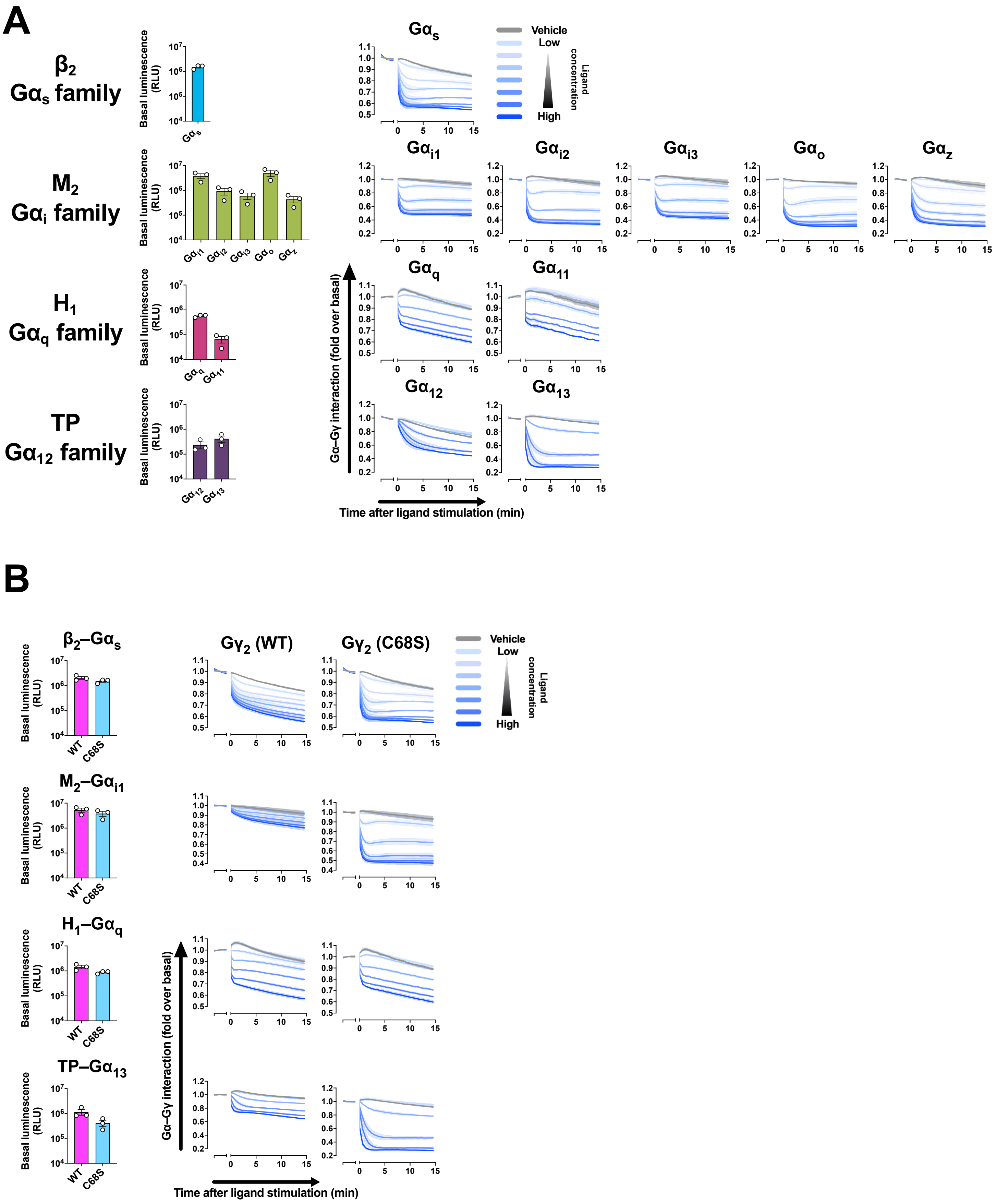
A
β2
Gαs family
M2
Gαi family
H1
Gαq family
TP
Gα12 family
Basal luminescence (RLU)
Gαs
Gαi1
Gαi2
Gαi3
Gαo
Gαz
Gαq
Gα11
Gα12
Gα13
Vehicle
Low
High
Ligand concentration
Gα–Gγ interaction (fold over basal)
Time after ligand stimulation (min)
B
β2–Gαs
M2–Gαi1
H1–Gαq
TP–Gα13
WT
C68S
Gγ2 (WT)
Gγ2 (C68S)

### S2 Figure

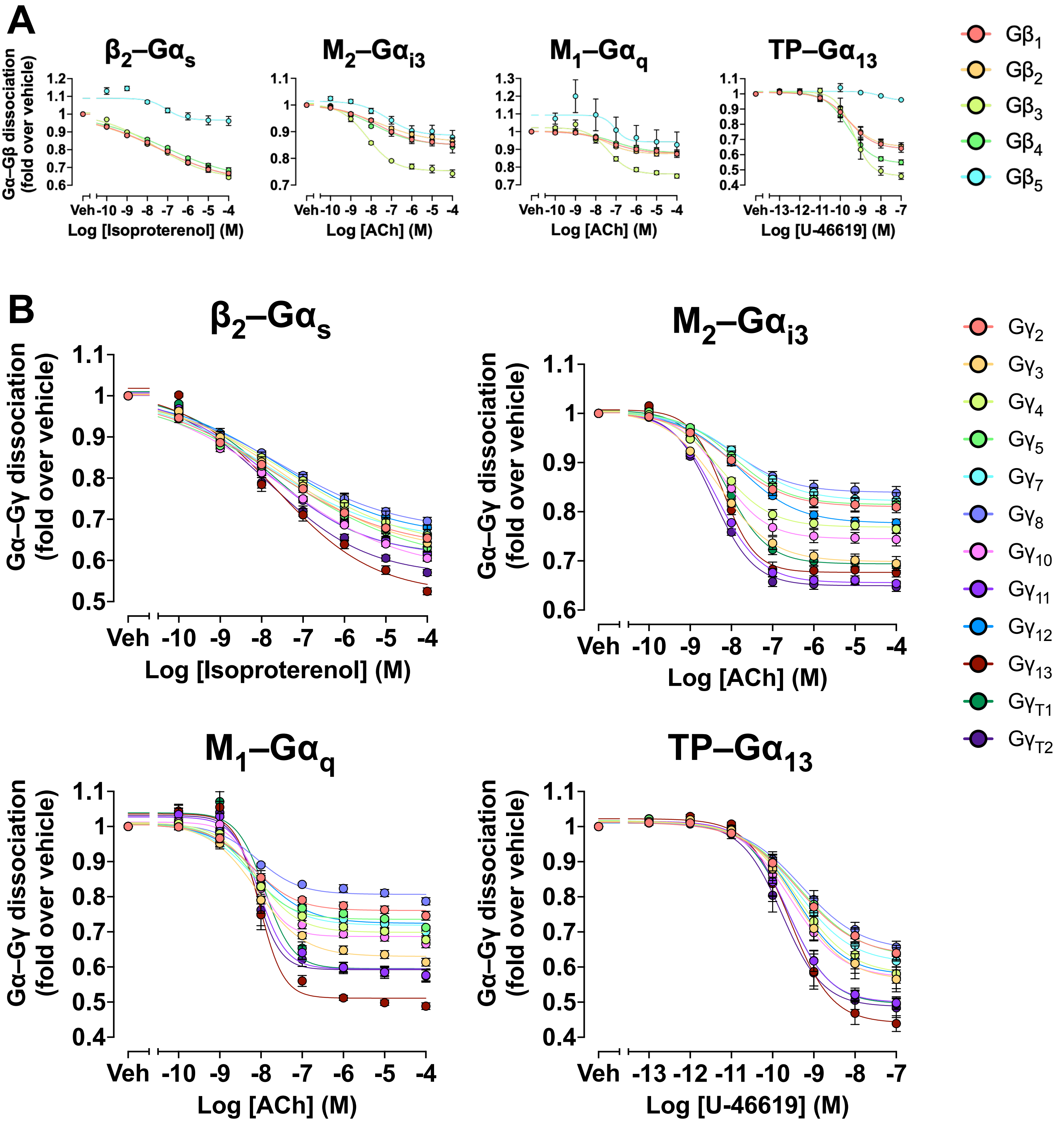

A
β2–Gαs
M2–Gαi3
M1–Gαq
TP–Gα13
Gα–Gβ dissociation (fold over vehicle)
Log [Isoproterenol] (M)
Log [ACh] (M)
Log [ACh] (M)
Log [U-46619] (M)
Gβ1
Gβ2
Gβ3
Gβ4
Gβ5
B
β2–Gαs
M2–Gαi3
M1–Gαq
TP–Gα13
Gα–Gγ dissociation (fold over vehicle)
Log [Isoproterenol] (M)
Log [ACh] (M)
Log [ACh] (M)
Log [U-46619] (M)
Gγ2
Gγ3
Gγ4
Gγ5
Gγ7
Gγ8
Gγ10
Gγ11
Gγ12
Gγ13
GγT1
GγT2

### S3 Figure

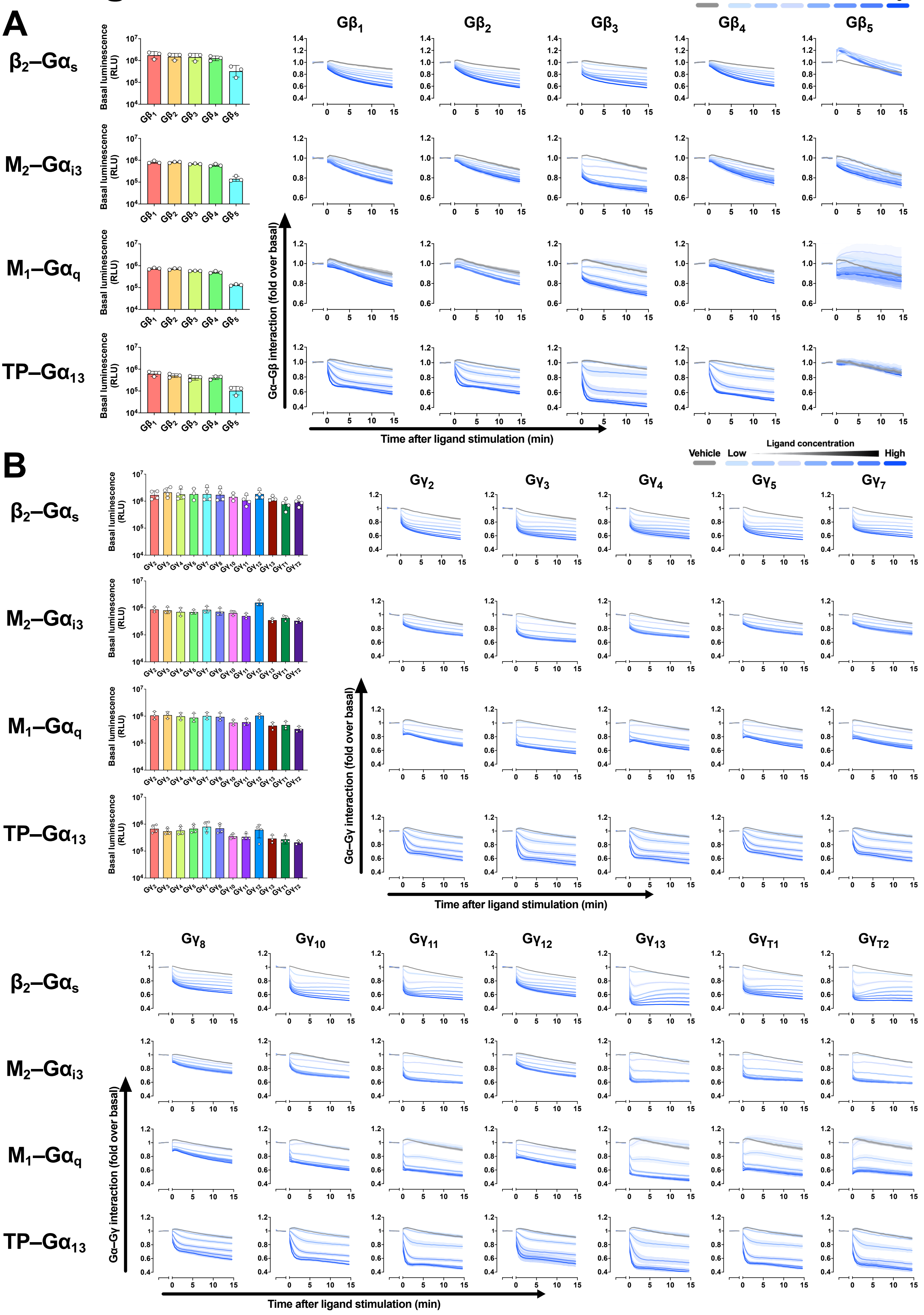
A
B
Vehicle
Low
Ligand concentration
High
β2–Gαs
M2–Gαi3
M1–Gαq
TP–Gα13
Basal luminescence (RLU)
Gβ1
Gβ2
Gβ3
Gβ4
Gβ5
Gα–Gβ interaction (fold over basal)
Time after ligand stimulation (min)
Gγ2
Gγ3
Gγ4
Gγ5
Gγ7
Gγ8
Gγ10
Gγ11
Gγ12
Gγ13
GγT1
GγT2
Gα–Gγ interaction (fold over basal)

### S4 Figure

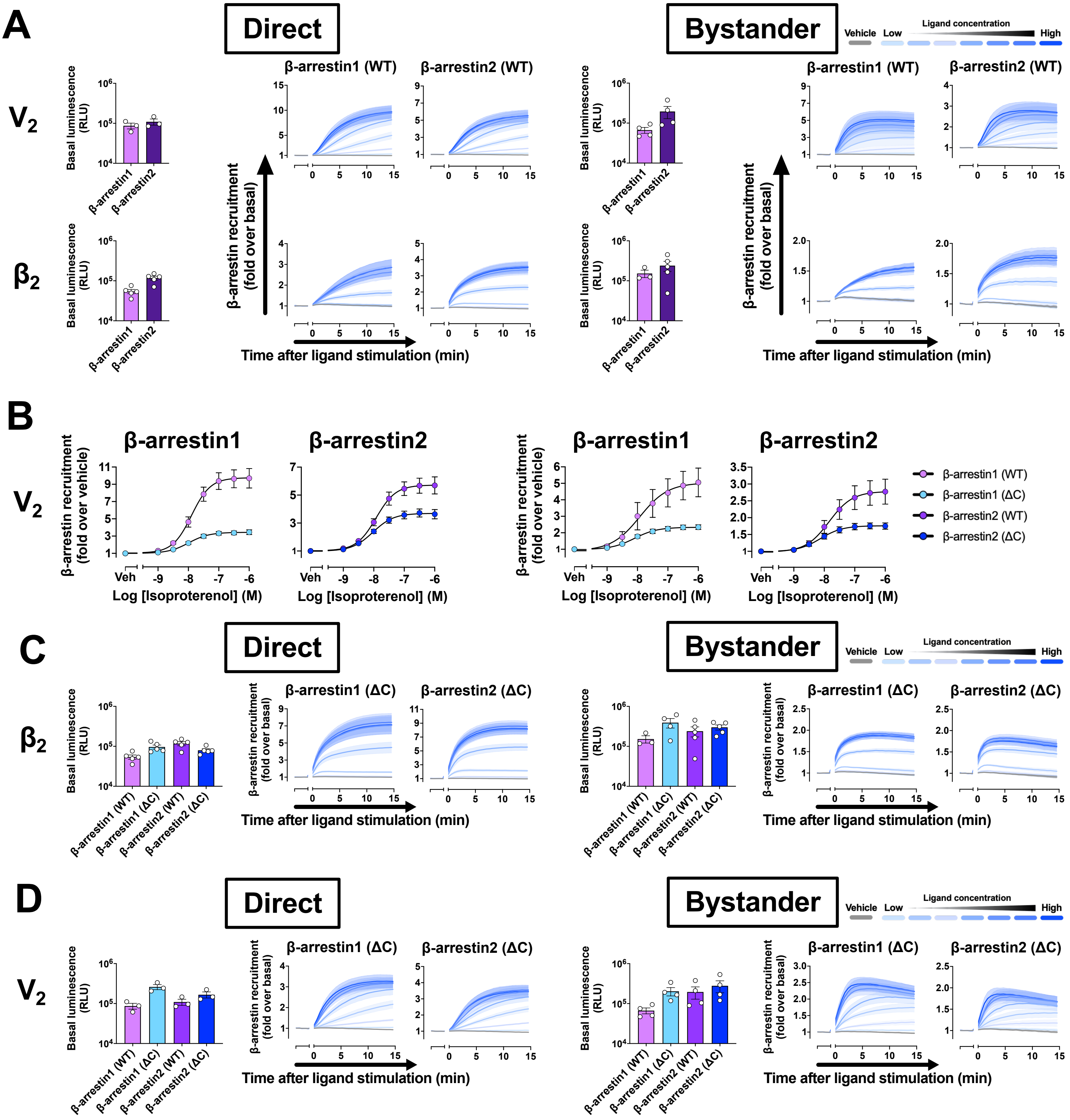

A
Direct
Bystander
Vehicle
Low
Ligand concentration
High
V2
β2
Basal luminescence (RLU)
β-arrestin1
β-arrestin2
β-arrestin1 (WT)
β-arrestin2 (WT)
β-arrestin recruitment (fold over basal)
Time after ligand stimulation (min)
B
β-arrestin1
β-arrestin2
β-arrestin recruitment (fold over vehicle)
Veh
Log [Isoproterenol] (M)
β-arrestin1 (WT)
β-arrestin1 (ΔC)
β-arrestin2 (WT)
β-arrestin2 (ΔC)
C
D

### S5 Figure

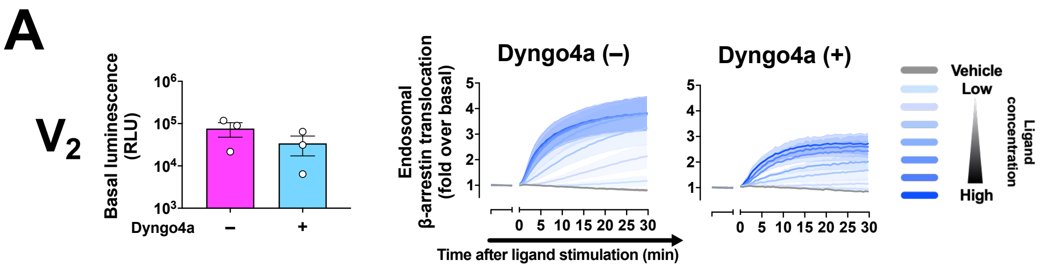

A
V2
Basal luminescence (RLU)
Dyngo4a
Dyngo4a (–)
Dyngo4a (+)
Endosomal β-arrestin translocation (fold over basal)
Time after ligand stimulation (min)
Vehicle
Low
High
Ligand concentration


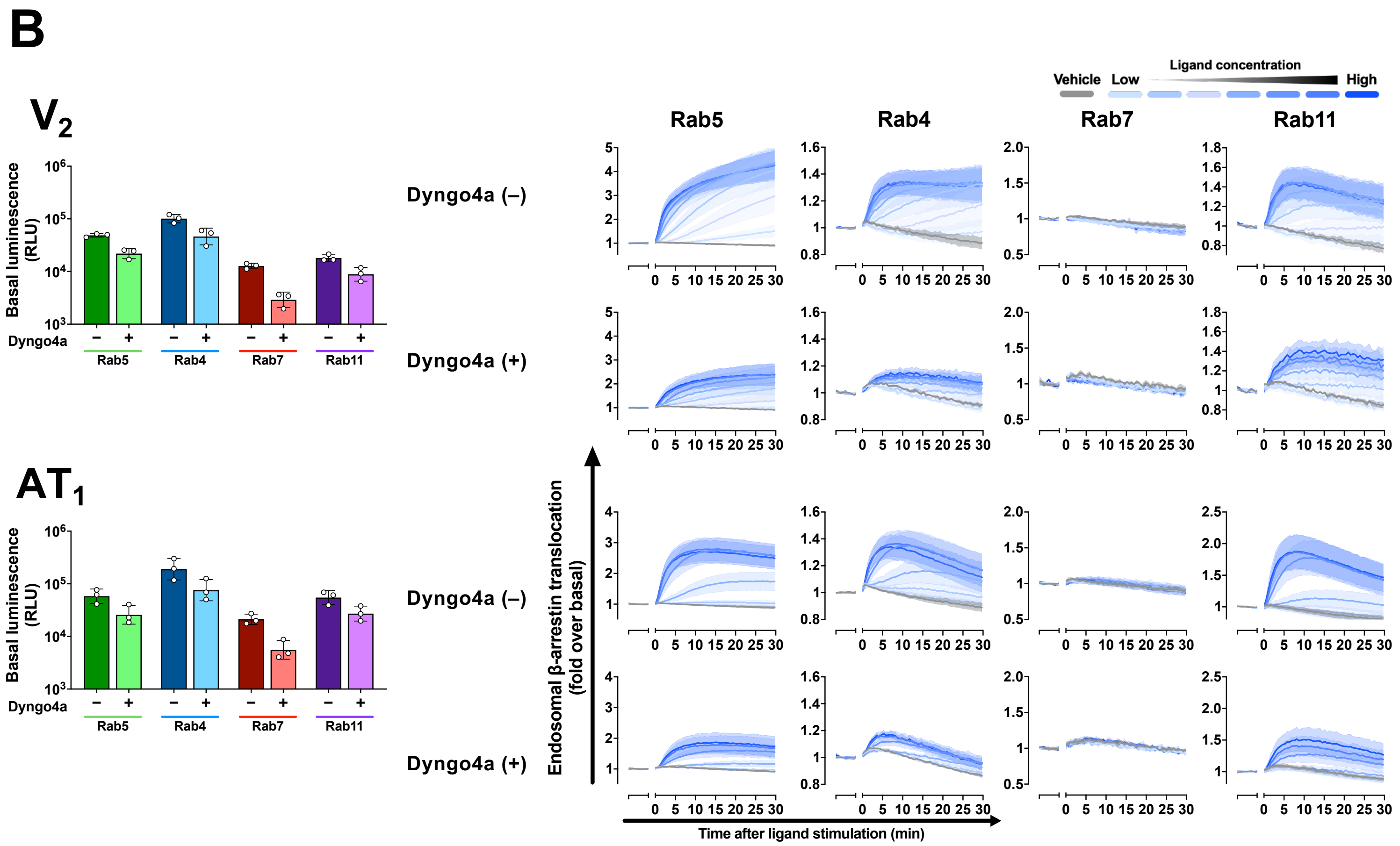

B
V2
AT1
Basal luminescence (RLU)
Dyngo4a
Rab5
Rab4
Rab7
Rab11
Dyngo4a (–)
Dyngo4a (+)
Vehicle
Low
Ligand concentration
High
Endosomal β-arrestin translocation (fold over basal)
Time after ligand stimulation (min)

### S6 Figure

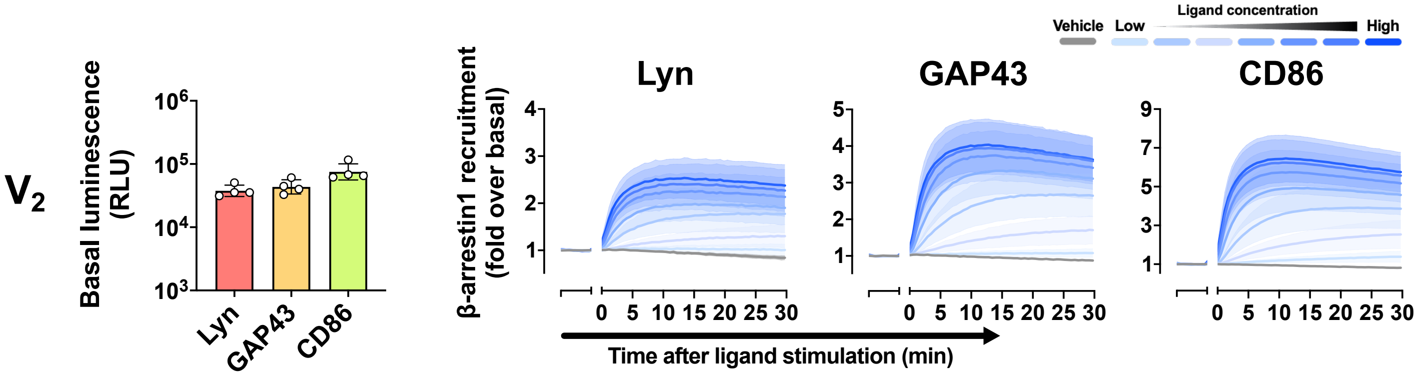

Vehicle
Low
Ligand concentration
High
V2
Basal luminescence (RLU)
10^6
10^5
10^4
10^3
Lyn
GAP43
CD86
β-arrestin1 recruitment (fold over basal)
Lyn
GAP43
CD86
0 5 10 15 20 25 30
Time after ligand stimulation (min)

### S7 Figure

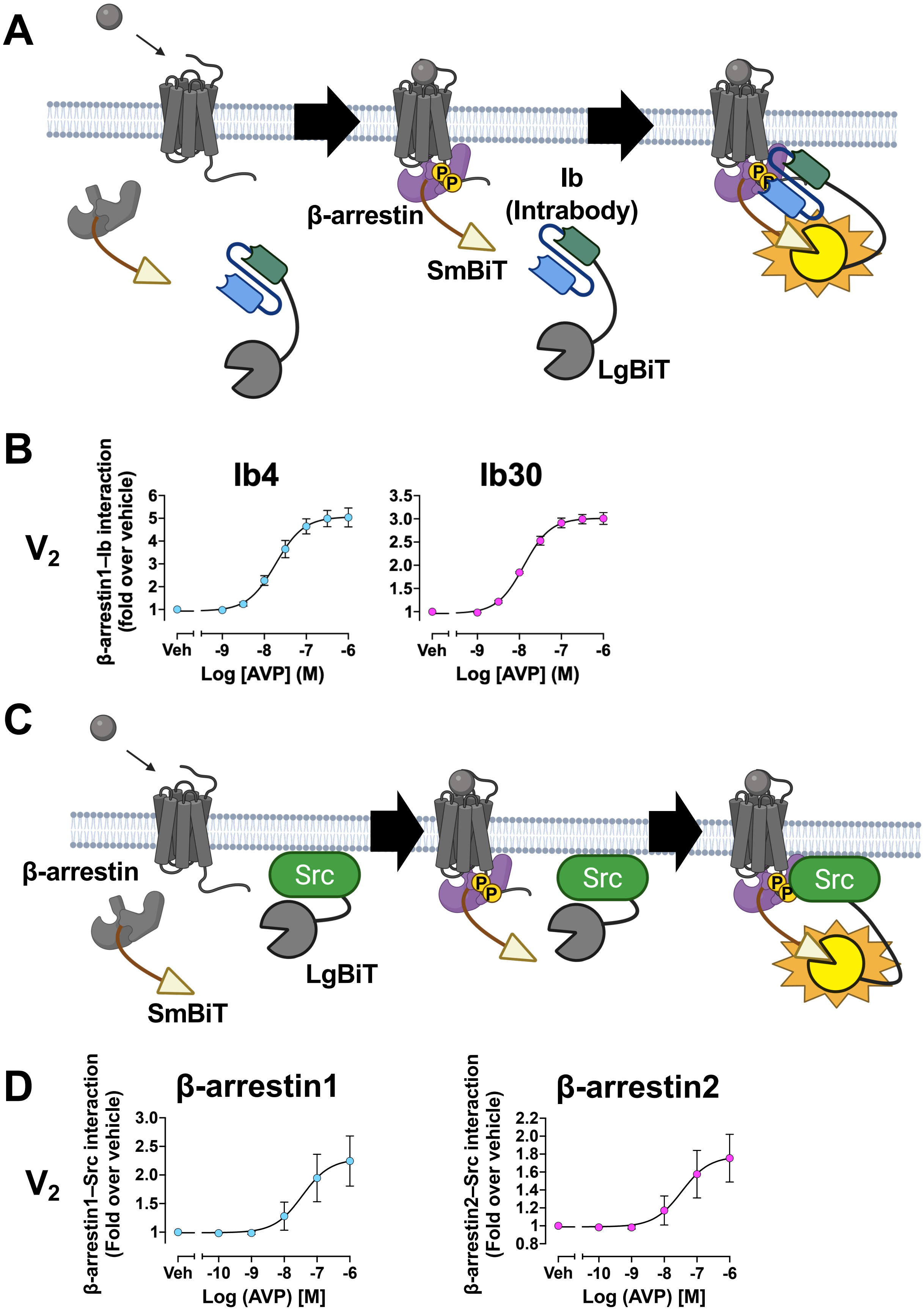
A
β-arrestin
SmBiT
Ib
(Intrabody)
LgBiT
P
B
V2
Ib4
Ib30
β-arrestin1–Ib interaction
(fold over vehicle)
Log [AVP] (M)
Veh
-9
-8
-7
-6
C
β-arrestin
SmBiT
Src
LgBiT
D
V2
β-arrestin1
β-arrestin2
β-arrestin1–Src interaction
(Fold over vehicle)
β-arrestin2–Src interaction
(Fold over vehicle)
Log (AVP) [M]
Veh
-10
-9
-8
-7
-6

### S8 Figure

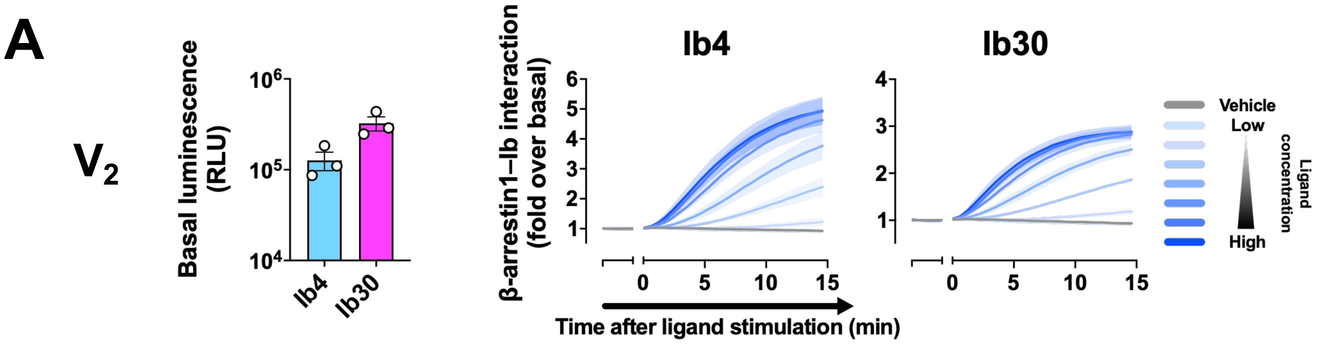
A
V2
Basal luminescence (RLU)
10^6
10^5
10^4
Ib4
Ib30
β-arrestin1–Ib interaction (fold over basal)
Ib4
Ib30
6
5
4
3
2
1
0
5
10
15
Time after ligand stimulation (min)
Vehicle
Low
High
Ligand concentration

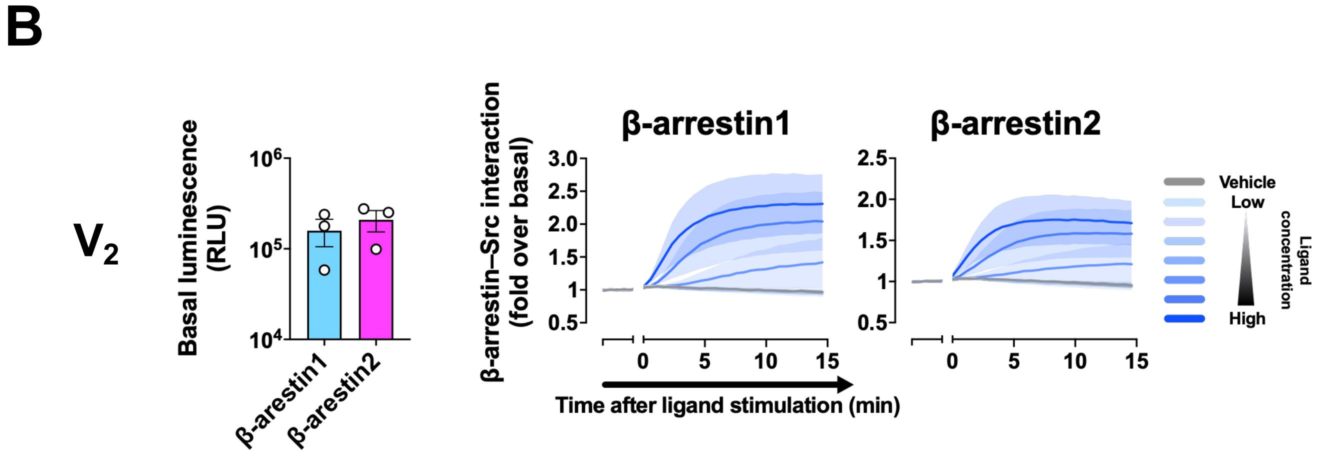
B
V2
Basal luminescence (RLU)
10^6
10^5
10^4
β-arestin1
β-arestin2
β-arrestin–Src interaction (fold over basal)
β-arrestin1
β-arrestin2
3.0
2.5
2.0
1.5
1
0.5
0
5
10
15
Time after ligand stimulation (min)
Vehicle
Low
High
Ligand concentration

### S9 Figure

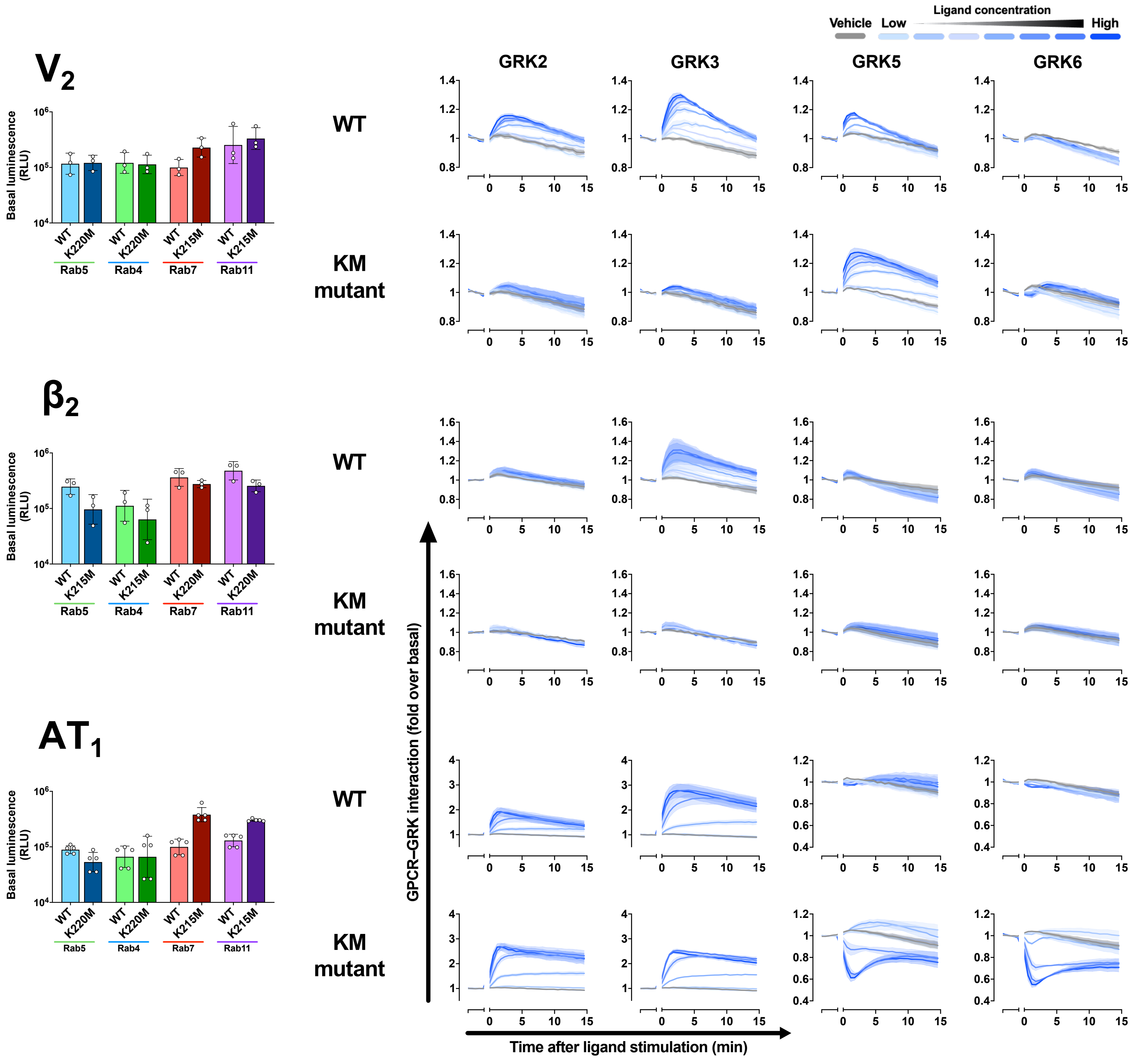

Vehicle
Low
Ligand concentration
High
V2
β2
AT1
GRK2
GRK3
GRK5
GRK6
WT
KM mutant
Basal luminescence (RLU)
WT
K220M
K215M
Rab5
Rab4
Rab7
Rab11
GPCR–GRK interaction (fold over basal)
Time after ligand stimulation (min)

### S10 Figure

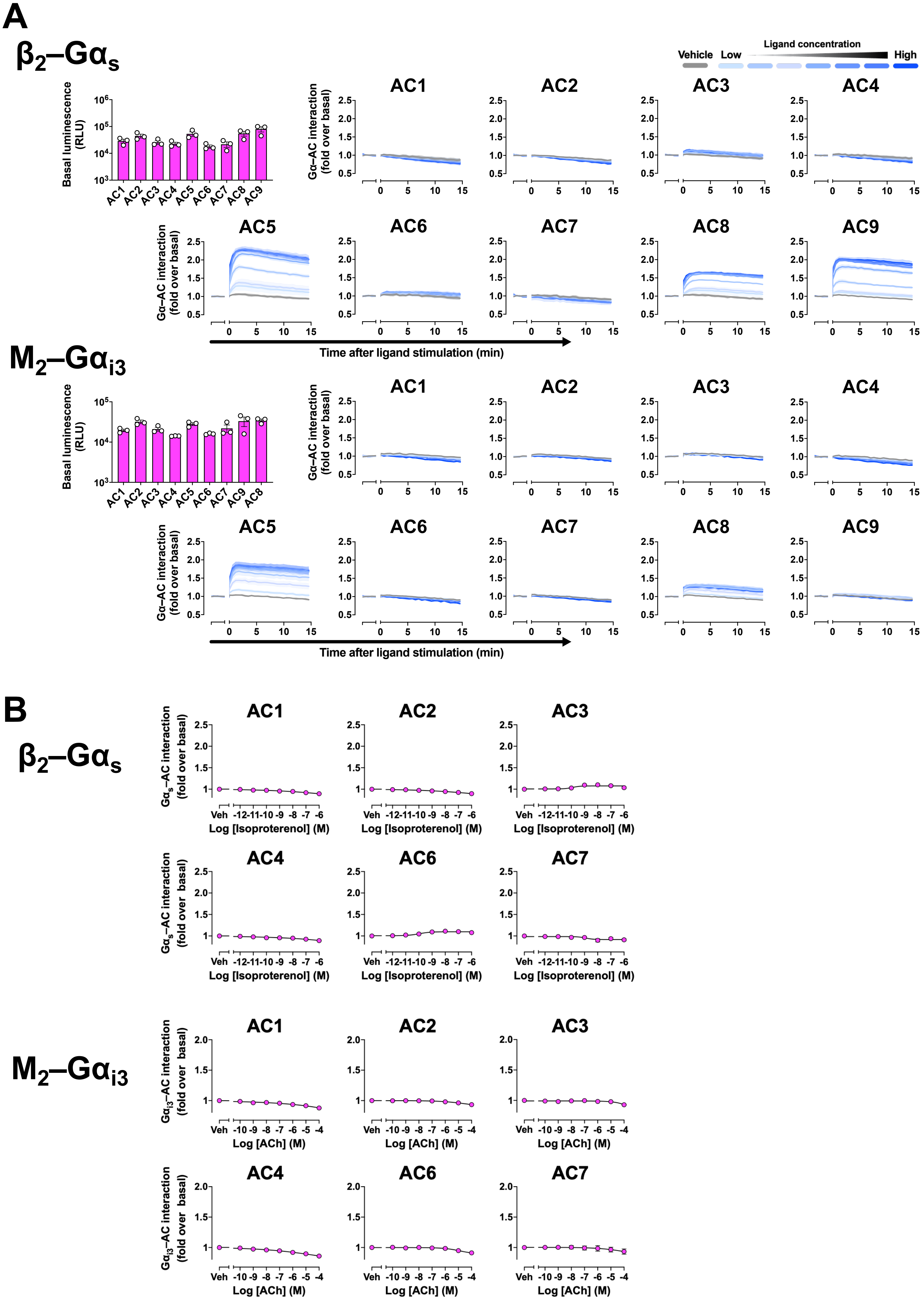
A
β2–Gαs
Vehicle
Low
Ligand concentration
High
Basal luminescence (RLU)
AC1 AC2 AC3 AC4 AC5 AC6 AC7 AC8 AC9
Gα–AC interaction (fold over basal)
Time after ligand stimulation (min)
M2–Gαi3
AC1 AC2 AC3 AC4 AC5 AC6 AC7 AC9 AC8
B
β2–Gαs
Gαs–AC interaction (fold over basal)
Log [Isoproterenol] (M)
Veh -12 -11 -10 -9 -8 -7 -6
M2–Gαi3
Gαi3–AC interaction (fold over basal)
Log [ACh] (M)
Veh -10 -9 -8 -7 -6 -5 -4

### S11 Figure

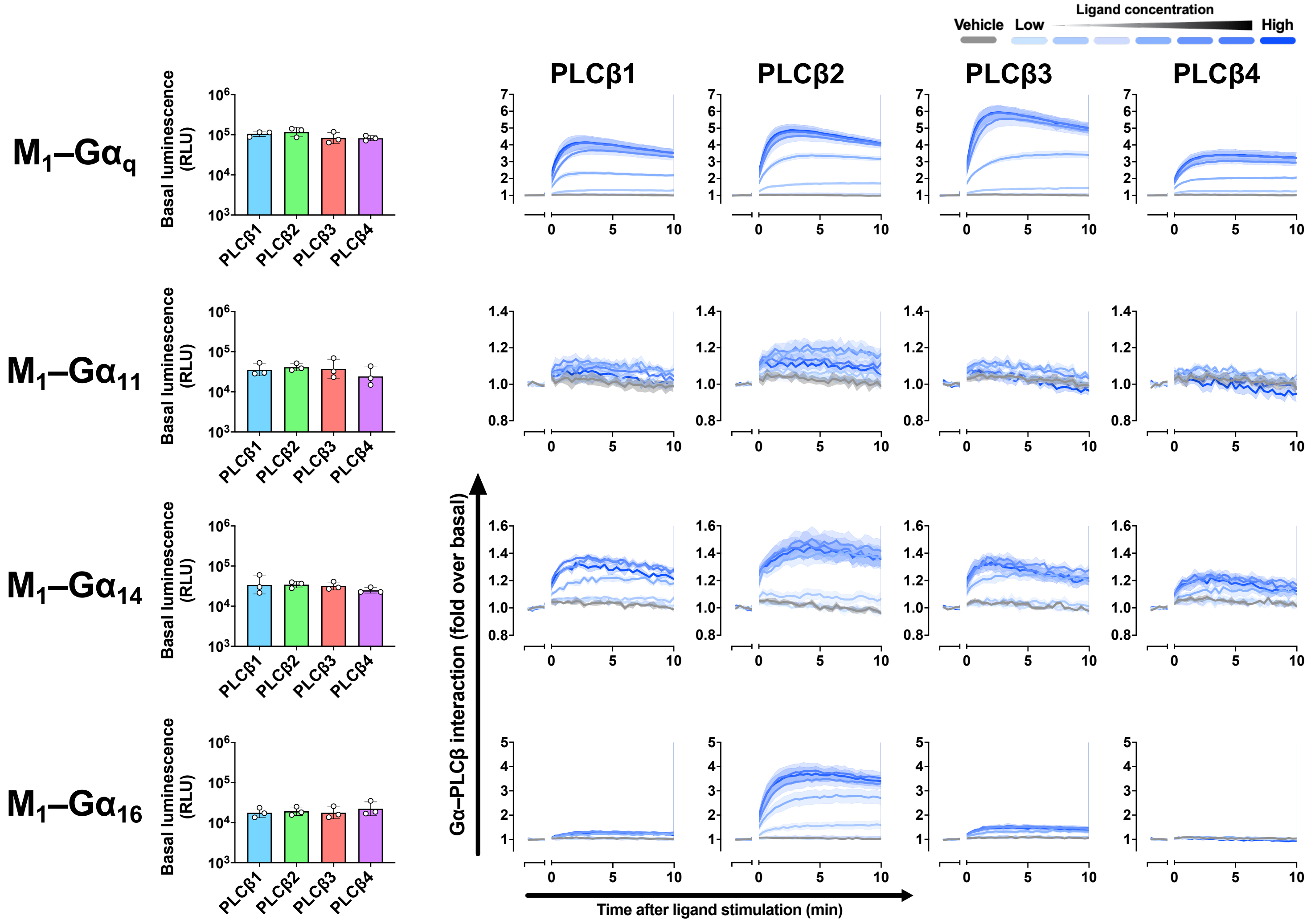

Vehicle
Low
Ligand concentration
High
PLCβ1
PLCβ2
PLCβ3
PLCβ4
M1–Gαq
M1–Gα11
M1–Gα14
M1–Gα16
Basal luminescence (RLU)
Gα–PLCβ interaction (fold over basal)
Time after ligand stimulation (min)

### S12 Figure

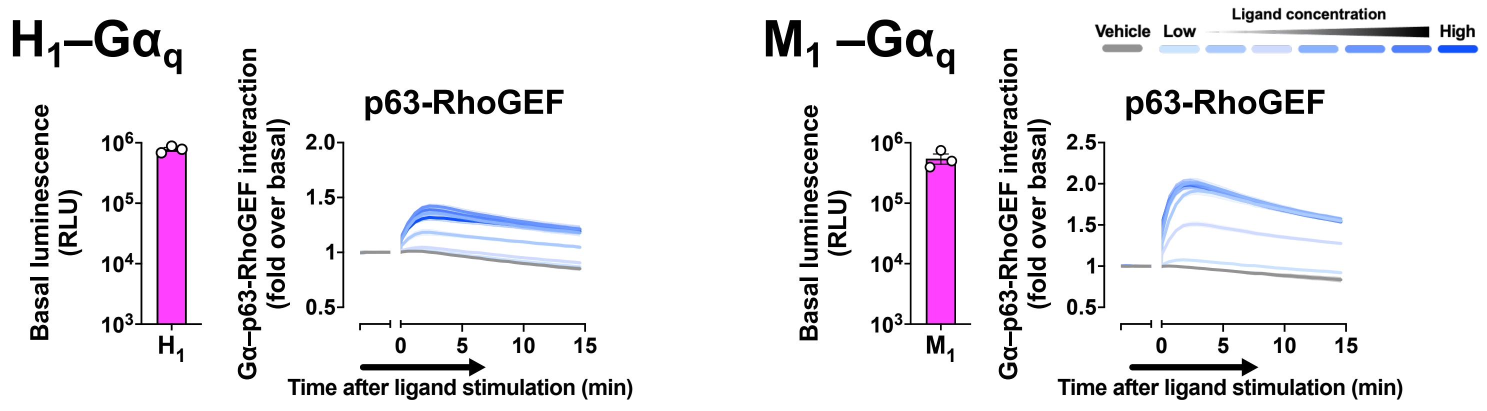
H1–Gαq
Basal luminescence (RLU)
10^6
10^5
10^4
10^3
H1
Gα–p63-RhoGEF interaction (fold over basal)
p63-RhoGEF
2.0
1.5
1
0.5
0
5
10
15
Time after ligand stimulation (min)
M1 –Gαq
Basal luminescence (RLU)
M1
Gα–p63-RhoGEF interaction (fold over basal)
p63-RhoGEF
2.5
2.0
1.5
1
0.5
Time after ligand stimulation (min)
Vehicle
Low
Ligand concentration
High

### S13 Figure

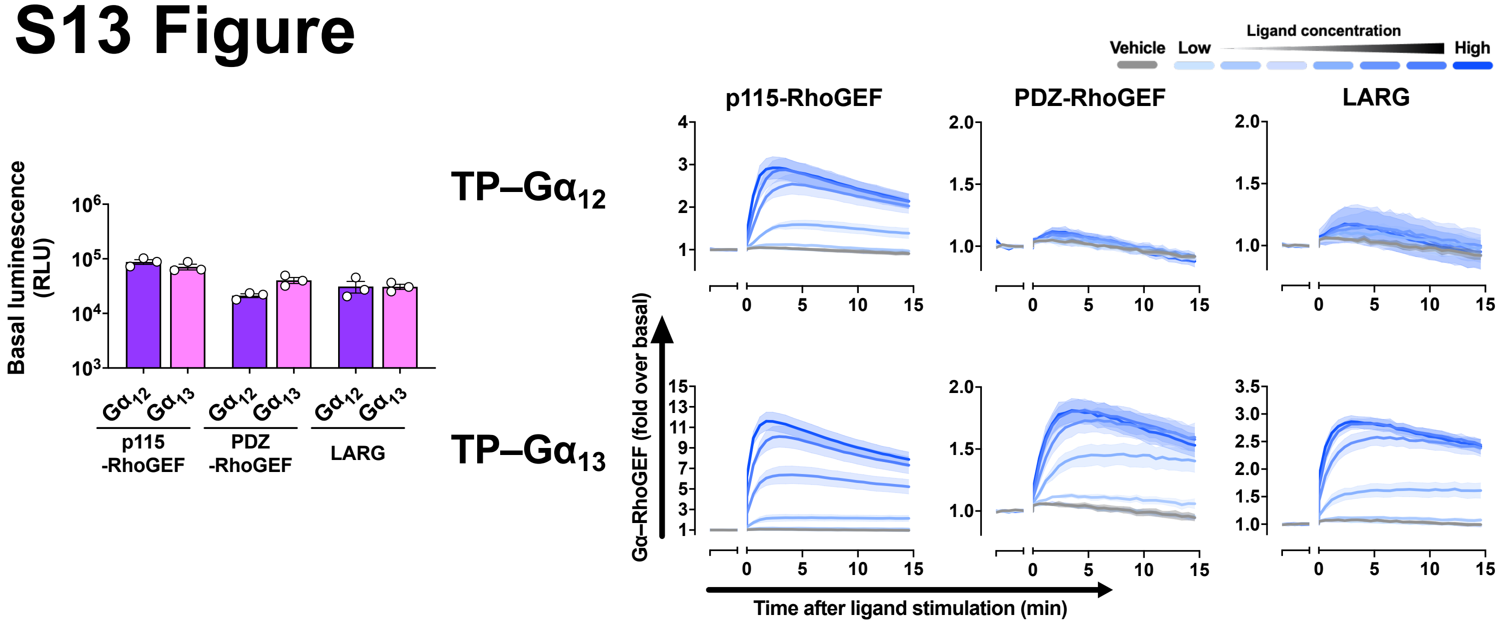

Basal luminescence (RLU)
p115 -RhoGEF
PDZ -RhoGEF
LARG
TP–Gα12
TP–Gα13
Vehicle
Low
Ligand concentration
High
p115-RhoGEF
PDZ-RhoGEF
LARG
Gα–RhoGEF (fold over basal)
Time after ligand stimulation (min)

### S14 Figure

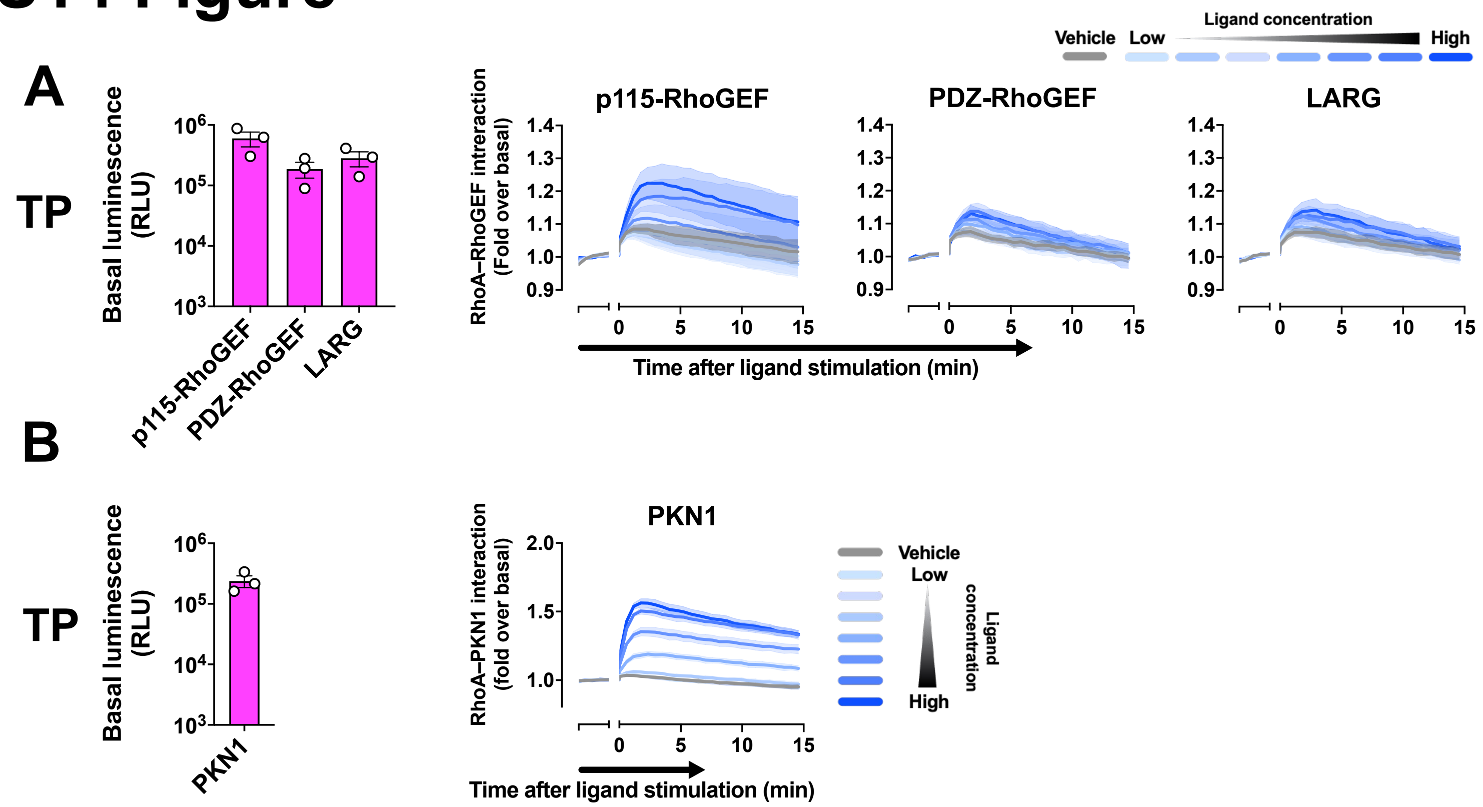

A
TP
Basal luminescence (RLU)
p115-RhoGEF
PDZ-RhoGEF
LARG
Vehicle
Low
Ligand concentration
High
RhoA–RhoGEF interaction (Fold over basal)
Time after ligand stimulation (min)
B
PKN1
RhoA–PKN1 interaction (fold over basal)
Ligand concentration
