## Supplementary material for "Comprehensive dissection of GPCR signaling using a NanoBiT-based platform": Table

**Table 1 LgBiT-fused Gα plasmids used in this study and their LgBiT insertion sites**

| Protein class | Plasmid Name | LgBiT insertion site |
| --- | --- | --- |
| Gα_s/olf_ family | Gα_s_-LgBiT | between positions 99 and 100 |
|  | Gα_olf_-LgBiT | between positions 100 and 101 |
| Gα_i/o_ family | Gα_i1_-LgBiT | between positions 91 and 92 |
|  | Gα_i2_-LgBiT | between positions 91 and 92 |
|  | Gα_i3_-LgBiT | between positions 91 and 92 |
|  | Gα_o_-LgBiT | between positions 91 and 92 |
|  | Gα_z_-LgBiT | between positions 91 and 92 |
| Gα_q/11_ family | Gα_q_-LgBiT | between positions 97 and 98 |
|  | Gα_q_-LgBiT (αB–αC) | between positions 123 and 124 |
|  | Gα_11_-LgBiT | between positions 97 and 98 |
|  | Gα_14_-LgBiT (αB–αC) | between positions 119 and 120 |
|  | Gα_16_-LgBiT (αB–αC) | between positions 126 and 127 |
| Gα_12/13_ family | Gα_12_-LgBiT | between positions 115 and 116 |
|  | Gα_13_-LgBiT | between positions 106 and 107 |

**Table 2 Reagent and tool table**

| **Reagent/resource** | **Reference or source** | **Identifier or catalog number** |
| --- | --- | --- |
| **Experimental models** | | |
| HEK293A cells | Thermo Fisher Scientific | R70507 |
| **Recombinant DNA** | | |
| Gα_s_-LgBiT | Inoue *et al.*, *Cell*, 2019 | - |
| Gα_olf_-LgBiT | This study | - |
| Gα_i1_-LgBiT | Inoue *et al.*, *Cell*, 2019 | - |
| Gα_i2_-LgBiT | Inoue *et al.*, *Cell*, 2019 | - |
| Gα_i3_-LgBiT | Inoue *et al.*, *Cell*, 2019 | - |
| Gα_o_-LgBiT | Inoue *et al.*, *Cell*, 2019 | - |
| Gα_z_-LgBiT | This study | - |
| Gα_q_-LgBiT | Inoue *et al.*, *Cell*, 2019 | - |
| Gα_q_-LgBiT (αB–αC) | Heo *et al.*, *PLos Biol.*, 2022 | - |
| Gα_11_-LgBiT | This study | - |
| Gα_14_-LgBiT (αB–αC) | This study | - |
| Gα_16_-LgBiT (αB–αC) | This study | - |
| Gα_12_-LgBiT | Inoue *et al.*, *Cell*, 2019 | - |
| Gα_13_-LgBiT | Inoue *et al.*, *Cell*, 2019 | - |
| SmBiT-Gβ_1_ | Inoue *et al.*, *Cell*, 2019 | - |
| SmBiT-Gβ_2_ | C.M.C. Carino *et al.*, *Eur J Pharmacol*., 2025 | - |
| SmBiT-Gβ_3_ | Inoue *et al.*, *Cell*, 2019 | - |
| SmBiT-Gβ_4_ | C.M.C. Carino *et al.*, *Eur J Pharmacol*., 2025 | - |
| SmBiT-Gβ_5_ | Inoue *et al.*, *Cell*, 2019 | - |
| SmBiT-Gγ_2_ | Inoue *et al.*, *Cell*, 2019 | - |
| SmBiT-Gγ_2_ (C68S) | Kato *et al.*, *Nature.*, 2019 | - |
| SmBiT-Gγ_3_ | This study | - |
| SmBiT-Gγ_4_ | This study | - |
| SmBiT-Gγ_5_ | This study | - |
| SmBiT-Gγ_7_ | This study | - |
| SmBiT-Gγ_8_ | This study | - |
| SmBiT-Gγ_10_ | This study | - |
| SmBiT-Gγ_11_ | This study | - |
| SmBiT-Gγ_12_ | This study | - |
| SmBiT-Gγ_13_ | This study | - |
| SmBiT-Gγ_T1_ | Inoue *et al.*, *Cell*, 2019 | - |
| SmBiT-Gγ_T2_ | This study | - |
| SmBiT-AC1 | This study | - |
| SmBiT-AC2 | This study | - |
| SmBiT-AC3 | This study | - |
| SmBiT-AC4 | This study | - |
| SmBiT-AC5 | This study | - |
| SmBiT-AC6 | This study | - |
| SmBiT-AC7 | This study | - |
| SmBiT-AC8 | This study | - |
| SmBiT-AC9 | This study | - |
| SmBiT-PLCβ1 | This study |  |
| SmBiT-PLCβ2 | Heo *et al.*, *PLos Biol.*, 2022 |  |
| SmBiT-PLCβ3 | This study |  |
| SmBiT-PLCβ4 | This study |  |
| LgBiT-RhoA | Inoue *et al.*, *Cell*, 2019 | - |
| SmBiT-PKN1–GBD | Inoue *et al.*, *Cell*, 2019 | - |
| SmBiT-p115-RhoGEF | This study | - |
| SmBiT-PDZ-RhoGEF | This study | - |
| SmBiT-LARG | This study | - |
| SmBiT-p63-RhoGEF | This study | - |
| LgBiT-Ib4 | This study | - |
| LgBiT-Ib30 | Baidya *et al.*, *EMBO Rep.*, 2020 | - |
| LgBiT-β-arrestin1 | Shihoya *et al.*, *Nat Commun.*, 2018  Dixon *et al.*, *ACS Chem Biol.*, 2016 | - |
| LgBiT-β-arrestin1 (ΔC382) | This study | - |
| LgBiT-β-arrestin2 | Shihoya *et al.*, *Nat Commun.*, 2018  Dixon *et al.*, *ACS Chem Biol.*, 2016 | - |
| LgBiT-β-arrestin2 (ΔC382) | This study | - |
| SmBiT-β-arrestin1 | Shihoya *et al.*, *Nat Commun.*, 2018  Dixon *et al.*, *ACS Chem Biol.*, 2016 | - |
| SmBiT-β-arrestin1 (ΔC382) | This study | - |
| SmBiT-β-arrestin2 | Shihoya *et al.*, *Nat Commun.*, 2018  Dixon *et al.*, *ACS Chem Biol.*, 2016 | - |
| SmBiT-β-arrestin2 (ΔC382) | This study | - |
| LgBiT-CAAX | Xu *et al.*, *Nat Cehm Biol.*, 2022 | - |
| Lyn-LgBiT | This study | - |
| GAP43-LgBiT | This study | - |
| CD86-LgBiT | This study | - |
| Endofin-LgBiT | Kawakami *et al., Nat Commun.*, 2022 | - |
| LgBiT-Rab4 | This study | - |
| LgBiT-Rab5 | This study | - |
| LgBiT-Rab7 | This study | - |
| LgBiT-Rab11 | This study | - |
| GRK2-LgBiT | Kawakami *et al., Nat Commun.*, 2022 | - |
| GRK2-LgBiT (K220M) | Tiwari *et al.*, *Mol Cell.*, 2026 | - |
| GRK3-LgBiT | Kawakami *et al., Nat Commun.*, 2022 | - |
| GRK3-LgBiT (K220M) | Tiwari *et al.*, *Mol Cell.*, 2026 | - |
| GRK5-LgBiT | Kawakami *et al., Nat Commun.*, 2022 | - |
| GRK5-LgBiT (K215M) | Tiwari *et al.*, *Mol Cell.*, 2026 | - |
| GRK6-LgBiT | Kawakami *et al., Nat Commun.*, 2022 | - |
| GRK6-LgBiT (K215M) | Tiwari *et al.*, *Mol Cell.*, 2026 | - |
| SRC-LgBiT | This study | - |
| **Antibodies** | - | - |
| **Oligonucleotides and other sequence-based reagents** | | |
| **Chemicals, enzymes and other reagents** | | |
| Angiotensin II (human) | Peptide Institute | 4001-v |
| [Arg8]-vasopressin | Peptide Institute | 4085-v |
| Acetylcholine chloride | Peptide Institute | A6625 |
| (–)-isoproterenol hydrochloride | Cayman Chemicals | I6504 |
| U-46619 | Cayman Chemicals | 16450 |
| Histamine dihydrochloride | FUJIFILM Wako Pure Chemical | 087-03553 |
| **Software** |  |  |
| GraphPad Prism 10 | GraphPad Software | - |
| ImageJ | National Institutes of Health | - |
