## Appendix for "Comprehensive dissection of GPCR signaling using a NanoBiT-based platform"

Gα_s_-linker-LgBiT

ATGGGATGCCTGGGAAACAGTAAAACCGAAGATCAGAGGAACGAAGAAAAAGCACAGAGGGAAGCCAACAAGAAGATTGAGAAACAGCTCCAGAAGGATAAGCAGGTGTACCGGGCCACACACAGACTGCTGCTGCTGGGAGCAGGAGAGTCCGGCAAGTCTACCATCGTGAAGCAGATGAGAATCCTGCACGTGAACGGCTTCAATGGCGACAGCGAGAAGGCCACAAAGGTGCAGGACATCAAGAACAATCTGAAGGAGGCCATCGAGACAATCGTGGCCGCCATGTCCAATCTGGGAGGCTCTGGAGGAGGAGGCAGCGGAGGCAGCTCCTCTGGCGGCGTGTTCACACTGGAGGACTTTGTGGGCGATTGGGAGCAGACCGCCGCCTACAACCTGGACCAGGTGCTGGAGCAGGGAGGCGTGAGCAGCCTGCTCCAGAATCTGGCCGTGTCCGTGACCCCAATCCAGAGGATCGTGCGCTCTGGCGAGAACGCCCTGAAGATCGATATCCACGTGATCATCCCCTATGAGGGCCTGAGCGCCGACCAGATGGCACAGATCGAGGAGGTGTTCAAGGTGGTGTACCCTGTGGACGATCACCACTTCAAAGTGATCCTGCCATATGGCACCCTGGTCATCGATGGCGTGACCCCAAACATGCTGAATTACTTCGGCAGGCCCTATGAGGGCATCGCCGTGTTTGACGGCAAGAAGATCACCGTGACAGGCACCCTGTGGAACGGCAATAAGATCATCGACGAGCGGCTGATCACACCCGATGGCTCCATGCTGTTCAGAGTGACCATCAATAGCGGAGGCTCCGGCGGAGGAGGCTCTGGAGGCTCTAGCTCCGGCGGCGTGCCCCCTGTGGAGCTGGCCAACCCCGAGAATCAGTTCCGCGTGGATTACATCCTGAGCGTGATGAACGTGCCTGACTTCGATTTTCCACCCGAGTTTTATGAGCACGCAAAGGCCCTGTGGGAGGACGAGGGCGTGAGGGCATGCTACGAGCGGAGCAATGAGTATCAGCTCATCGATTGTGCCCAGTACTTTCTGGACAAGATCGATGTGATCAAGCAGGCCGACTATGTGCCTTCTGACCAGGATCTGCTGCGGTGCAGAGTGCTGACAAGCGGCATCTTCGAGACAAAGTTTCAGGTGGATAAGGTGAACTTCCACATGTTTGACGTGGGCGGCCAGAGAGATGAGCGGAGAAAGTGGATTCAGTGTTTCAACGATGTGACAGCCATCATCTTTGTGGTGGCCTCTAGCTCCTACAATATGGTCATCAGGGAGGACAACCAGACCAATCGCCTCCAGGAGGCCCTGAACCTGTTCAAGAGCATCTGGAACAATCGGTGGCTGAGAACAATCTCCGTGATCCTGTTTCTGAATAAGCAGGATCTGCTGGCCGAGAAGGTGCTGGCCGGCAAGTCCAAGATCGAGGACTACTTCCCCGAGTTTGCCCGGTATACCACACCTGAGGACGCCACACCTGAGCCAGGAGAGGACCCCCGGGTGACCCGCGCCAAGTATTTCATCCGGGATGAGTTTCTGAGAATCTCTACCGCCAGCGGCGACGGCAGACACTACTGCTATCCACACTTCACATGTGCCGTGGACACCGAGAACATCAGGCGCGTGTTCAACGATTGTAGAGACATCATCCAGAGAATGCACCTGAGACAGTATGAGCTGCTGTGA

Gα_i1_-linker-LgBiT

ATGGGATGCACCCTGTCCGCCGAGGATAAAGCCGCAGTCGAGAGGAGCAAAATGATTGATAGGAACCTGAGAGAGGATGGAGAGAAAGCCGCCAGGGAGGTGAAGCTGCTGCTGCTGGGAGCAGGAGAGTCTGGCAAGAGCACAATCGTGAAGCAGATGAAGATCATCCACGAGGCCGGCTACAGCGAGGAGGAGTGCAAGCAGTACAAGGCCGTGGTGTATAGCAATACCATCCAGTCCATCATCGCCATCATCAGGGCAATGGGCAGACTGGGAGGCTCTGGAGGAGGAGGCAGCGGAGGCAGCTCCTCTGGCGGCGTGTTCACACTGGAGGACTTTGTGGGCGATTGGGAGCAGACCGCCGCCTACAATCTGGACCAGGTGCTGGAGCAGGGAGGCGTGAGCAGCCTGCTCCAGAACCTGGCCGTGTCTGTGACCCCAATCCAGCGGATCGTGAGAAGCGGCGAGAACGCCCTGAAGATCGACATCCACGTGATCATCCCCTATGAGGGCCTGTCCGCCGATCAGATGGCCCAGATCGAGGAGGTGTTCAAGGTGGTGTACCCCGTGGACGATCACCACTTCAAAGTGATCCTGCCTTATGGCACCCTGGTCATCGACGGCGTGACCCCTAACATGCTGAATTACTTCGGCAGACCATATGAGGGCATCGCCGTGTTTGATGGCAAGAAGATCACCGTGACAGGCACCCTGTGGAACGGCAATAAGATCATCGACGAGCGGCTGATCACACCTGATGGCTCCATGCTGTTCAGAGTGACCATCAATTCCGGCGGCTCTGGCGGCGGCGGCAGCGGAGGCTCTAGCTCCGGAGGCAAGATCGACTTTGGCGATTCCGCCAGGGCCGACGATGCAAGGCAGCTCTTCGTGCTGGCAGGAGCCGCCGAGGAGGGCTTTATGACCGCCGAGCTGGCAGGCGTGATCAAGAGGCTGTGGAAGGACTCTGGCGTGCAGGCCTGTTTCAACCGGAGCAGAGAGTACCAGCTCAACGATTCCGCCGCCTACTATCTGAACGACCTGGATCGCATCGCCCAGCCCAACTATATCCCTACACAGCAGGACGTGCTGAGGACCCGCGTGAAGACCACAGGCATCGTGGAGACACACTTCACCTTTAAGGACCTGCACTTCAAGATGTTTGATGTGGGCGGCCAGAGGTCTGAGCGCAAGAAGTGGATTCACTGCTTCGAGGGCGTGACAGCCATCATCTTCTGCGTGGCCCTGAGCGACTACGATCTGGTGCTGGCCGAGGACGAGGAGATGAATCGGATGCACGAGTCCATGAAGCTGTTCGACTCTATCTGCAACAATAAGTGGTTTACAGATACCAGCATCATCCTGTTCCTGAACAAGAAGGATCTGTTTGAGGAGAAGATCAAGAAGTCCCCACTGACAATCTGCTACCCCGAGTATGCCGGCTCTAATACCTACGAGGAGGCTGCCGCCTATATCCAGTGTCAGTTCGAGGACCTGAACAAGAGAAAGGATACCAAGGAAATCTACACACACTTCACCTGTGCCACAGATACCAAGAATGTGCAGTTTGTGTTCGATGCTGTGACCGATGTGATTATCAAGAATAACCTGAAAGACTGCGGGCTGTTTTGA

Gα_i2_-linker-LgBiT

ATGGGATGCACAGTGAGCGCAGAGGACAAAGCCGCCGCCGAAAGGAGCAAAATGATTGACAAGAACCTGAGAGAGGACGGGGAAAAAGCAGCCCGGGAGGTGAAGCTGCTGCTGCTGGGAGCAGGAGAGTCTGGCAAGAGCACAATCGTGAAGCAGATGAAGATCATCCACGAGGACGGCTACAGCGAGGAGGAGTGCAGGCAGTACAGGGCAGTGGTGTATTCTAACACCATCCAGAGCATCATGGCCATCGTGAAGGCCATGGGCAATCTGGGAGGCTCCGGCGGAGGAGGCTCTGGAGGCAGCTCCTCTGGCGGCGTGTTCACACTGGAGGACTTTGTGGGCGATTGGGAGCAGACCGCCGCCTATAACCTGGATCAGGTGCTGGAGCAGGGAGGCGTGAGCAGCCTGCTCCAGAATCTGGCCGTGTCCGTGACCCCAATCCAGAGGATCGTGCGGAGCGGAGAGAACGCCCTGAAGATCGACATCCACGTGATCATCCCTTACGAGGGCCTGTCCGCCGATCAGATGGCCCAGATCGAGGAGGTGTTCAAGGTGGTGTATCCTGTGGACGATCACCACTTCAAAGTGATCCTGCCATACGGCACCCTGGTCATCGACGGAGTGACCCCAAACATGCTGAATTACTTCGGCAGGCCCTATGAGGGCATCGCCGTGTTTGATGGCAAGAAGATCACCGTGACAGGCACCCTGTGGAACGGCAATAAGATCATCGACGAGAGGCTGATCACACCTGATGGCTCTATGCTGTTCCGCGTGACCATCAATTCCGGAGGCTCTGGAGGAGGAGGCAGCGGAGGCTCTAGCTCCGGCGGCCAGATCGACTTCGCCGATCCAAGCCGGGCCGACGATGCAAGACAGCTCTTTGCACTGTCCTGCACAGCCGAGGAGCAGGGCGTGCTGCCAGACGATCTGTCTGGCGTGATCCGGAGACTGTGGGCAGACCACGGAGTGCAGGCATGTTTCGGCCGGAGCCGGGAGTATCAGCTCAACGACAGCGCCGCCTACTATCTGAATGATCTGGAGCGCATCGCCCAGTCCGACTACATCCCCACACAGCAGGATGTGCTGCGGACCAGAGTGAAGACCACAGGCATCGTGGAGACACACTTCACCTTTAAGGACCTGCACTTCAAGATGTTTGATGTGGGCGGCCAGAGGAGCGAGCGCAAGAAGTGGATTCACTGCTTCGAGGGCGTGACCGCCATCATCTTCTGCGTGGCCCTGTCCGCCTATGACCTGGTGCTGGCCGAGGATGAGGAGATGAACCGGATGCACGAGAGCATGAAGCTGTTCGACTCCATCTGCAACAATAAGTGGTTTACAGATACCTCCATCATCCTGTTCCTGAATAAGAAGGACCTGTTTGAGGAGAAGATCACACACTCTCCCCTGACCATCTGTTTCCCTGAGTACACAGGCGCCAACAAGTATGATGAGGCCGCCAGCTACATCCAGTCCAAGTTTGAGGACCTGAATAAGAGAAAGGATACCAAGGAAATCTACACACACTTCACCTGTGCCACAGACACCAAGAACGTGCAGTTTGTGTTCGACGCTGTGACCGATGTGATTATCAAAAATAACCTGAAAGATTGTGGGCTGTTCTGA

Gα_i3_-linker-LgBiT

ATGGGATGCACCCTGTCCGCCGAGGATAAAGCCGCAGTCGAGAGGAGCAAAATGATTGATAGGAACCTGAGAGAGGATGGAGAGAAAGCCGCCAAGGAGGTGAAGCTGCTGCTGCTGGGAGCAGGAGAGTCTGGCAAGAGCACAATCGTGAAGCAGATGAAGATCATCCACGAGGACGGCTACAGCGAGGATGAGTGCAAGCAGTACAAGGTGGTGGTGTATAGCAATACCATCCAGTCCATCATCGCCATCATCAGGGCAATGGGCAGACTGGGAGGCTCTGGAGGAGGAGGCAGCGGAGGCAGCTCCTCTGGCGGCGTGTTCACACTGGAGGACTTTGTGGGCGATTGGGAGCAGACCGCCGCCTACAATCTGGACCAGGTGCTGGAGCAGGGAGGCGTGAGCAGCCTGCTCCAGAACCTGGCCGTGTCTGTGACCCCAATCCAGCGGATCGTGAGAAGCGGCGAGAACGCCCTGAAGATCGACATCCACGTGATCATCCCCTATGAGGGCCTGTCCGCCGATCAGATGGCCCAGATCGAGGAGGTGTTCAAGGTGGTGTACCCCGTGGACGATCACCACTTCAAAGTGATCCTGCCTTATGGCACCCTGGTCATCGACGGCGTGACCCCTAACATGCTGAATTACTTCGGCAGACCATATGAGGGCATCGCCGTGTTTGATGGCAAGAAGATCACCGTGACAGGCACCCTGTGGAACGGCAATAAGATCATCGACGAGCGGCTGATCACACCTGATGGCTCCATGCTGTTCAGAGTGACCATCAATTCCGGCGGCTCTGGCGGCGGCGGCAGCGGAGGCTCTAGCTCCGGAGGCAAGATCGACTTTGGCGAGGCAGCCAGGGCCGACGATGCAAGGCAGCTCTTCGTGCTGGCAGGAAGCGCCGAGGAGGGCGTGATGACCCCAGAGCTGGCAGGCGTGATCAAGAGGCTGTGGAGAGACGGAGGCGTGCAGGCCTGTTTCTCTCGGAGCAGAGAGTACCAGCTCAACGATTCCGCCTCTTACTATCTGAACGACCTGGATCGCATCAGCCAGTCCAACTATATCCCTACACAGCAGGACGTGCTGAGGACCCGCGTGAAGACCACAGGCATCGTGGAGACACACTTCACCTTTAAGGACCTGTACTTCAAGATGTTTGATGTGGGCGGCCAGAGGTCTGAGCGCAAGAAGTGGATTCACTGCTTCGAGGGCGTGACAGCCATCATCTTCTGCGTGGCCCTGAGCGACTACGATCTGGTGCTGGCCGAGGACGAGGAGATGAATCGGATGCACGAGTCCATGAAGCTGTTCGACTCTATCTGCAACAATAAGTGGTTTACAGAGACAAGCATCATCCTGTTCCTGAACAAGAAGGATCTGTTTGAGGAGAAGATCAAGCGGTCCCCACTGACAATCTGCTACCCCGAGTATACAGGCTCTAATACCTACGAGGAGGCTGCCGCCTATATCCAGTGTCAGTTCGAGGACCTGAACCGGAGAAAGGATACCAAGGAAATCTACACACACTTCACCTGTGCCACAGATACCAAGAATGTGCAGTTTGTGTTTGATGCTGTGACCGATGTGATTATCAAGAATAACCTGAAAGAATGCGGGCTGTATTGA

Gα_o_-linker-LgBiT

ATGGGATGCACCCTGTCCGCCGAGGAAAGAGCAGCACTGGAAAGAAGCAAGGCTATTGAGAAGAACCTGAAAGAAGATGGGATTAGTGCCGCCAAGGACGTGAAGCTGCTGCTGCTGGGAGCAGGAGAGTCTGGCAAGAGCACAATCGTGAAGCAGATGAAGATCATCCACGAGGACGGCTTCAGCGGCGAGGATGTGAAGCAGTACAAGCCCGTGGTGTATTCTAACACAATCCAGAGCCTGGCAGCAATCGTGCGGGCAATGGATACCCTGGGAGGCTCCGGAGGAGGAGGCTCTGGAGGCAGCTCCTCTGGCGGCGTGTTCACACTGGAGGACTTTGTGGGCGATTGGGAGCAGACCGCCGCCTATAACCTGGACCAGGTGCTGGAGCAGGGAGGCGTGAGCAGCCTGCTCCAGAATCTGGCCGTGTCTGTGACCCCCATCCAGCGGATCGTGAGAAGCGGCGAGAATGCCCTGAAGATCGATATCCACGTGATCATCCCTTACGAGGGCCTGTCCGCCGACCAGATGGCACAGATCGAGGAGGTGTTCAAGGTGGTGTATCCTGTGGACGATCACCACTTCAAAGTGATCCTGCCATACGGCACCCTGGTCATCGATGGCGTGACCCCAAACATGCTGAATTACTTCGGCAGGCCCTATGAGGGCATCGCCGTGTTTGACGGCAAGAAGATCACCGTGACAGGCACCCTGTGGAACGGCAATAAGATCATCGATGAGCGGCTGATCACACCTGACGGCTCCATGCTGTTCAGAGTGACCATCAACTCCGGAGGCTCTGGCGGCGGAGGCTCTGGAGGCTCTAGCTCCGGAGGAGGAATCGAGTACGGCGACAAGGAGAGGAAGGCCGATGCCAAGATGGTGTGCGACGTGGTGAGCCGCATGGAGGATACAGAGCCTTTCAGCGCCGAGCTGCTGTCCGCCATGATGAGGCTGTGGGGCGATAGCGGCATCCAGGAGTGTTTTAACCGGTCCAGAGAGTATCAGCTCAACGACTCCGCCAAGTACTATCTGGACTCTCTGGATAGAATCGGCGCCGCCGATTACCAGCCTACAGAGCAGGACATCCTGAGGACCCGCGTGAAGACCACAGGCATCGTGGAGACACACTTCACCTTTAAGAACCTGCACTTCAGGCTGTTTGATGTGGGCGGCCAGAGGTCTGAGCGCAAGAAGTGGATTCACTGCTTCGAGGACGTGACCGCCATCATCTTCTGCGTGGCCCTGTCCGGCTATGATCAGGTGCTGCACGAGGACGAGACAACAAATCGCATGCACGAGTCTCTGATGCTGTTTGATAGCATCTGCAACAATAAGTTCTTTATCGACACAAGCATCATCCTGTTCCTGAACAAGAAGGATCTGTTTGGCGAGAAGATCAAGAAGTCCCCACTGACCATCTGTTTCCCCGAGTACACAGGCCCTAATACCTATGAGGACGCTGCCGCCTACATCCAGGCCCAGTTTGAGAGCAAGAACCGGTCCCCCAATAAGGAAATCTACTGCCACATGACCTGTGCCACAGACACCAACAATATCCAGGTGGTCTTTGACGCTGTGACCGACATCATCATCGCCAATAACCTGAGAGGCTGCGGGCTGTATTGA

Gα_z_-linker-LgBiT

ATGGGATGTAGACAGTCCAGTGAAGAGAAAGAAGCAGCAAGAAGGAGCAGACGGATTGATAGACACCTGAGAAGTGAAAGCCAGAGGCAGCGGAGAGAGATCAAGCTGCTGCTGCTGGGCACATCCAATTCTGGCAAGTCCACCATCGTGAAGCAGATGAAGATCATCCACTCTGGCGGCTTCAATCTGGAGGCCTGCAAGGAGTACAAGCCCCTGATCATCTATAACGCCATCGATAGCCTGACACGGATCATCAGAGCCCTGGCCGCCCTGGGAGGATCTGGCGGAGGAGGCAGCGGCGGCAGCTCCTCTGGCGGCGTGTTCACACTGGAGGACTTTGTGGGCGATTGGGAGCAGACCGCCGCCTATAATCTGGACCAGGTGCTGGAGCAGGGAGGCGTGAGCTCCCTGCTGCAGAACCTGGCCGTGTCTGTGACCCCAATCCAGAGGATCGTGCGCAGCGGCGAGAACGCCCTGAAGATCGATATCCACGTGATCATCCCCTACGAGGGCCTGTCCGCCGACCAGATGGCACAGATCGAGGAGGTGTTCAAGGTGGTGTATCCCGTGGACGATCACCACTTCAAAGTGATCCTGCCTTACGGCACCCTGGTCATCGATGGCGTGACCCCAAATATGCTGAACTACTTCGGCAGACCCTATGAGGGCATCGCCGTGTTTGACGGCAAGAAGATCACCGTGACAGGCACCCTGTGGAATGGCAACAAGATCATCGACGAGCGGCTGATCACACCTGATGGCTCTATGCTGTTCAGAGTGACCATCAATAGCGGAGGATCCGGAGGAGGAGGAAGCGGAGGCTCTAGCTCCGGCGGCAGGATCGATTTCCACAACCCAGACCGCGCCTATGATGCCGTGCAGCTGTTTGCACTGACAGGACCTGCAGAGAGCAAGGGAGAGATCACCCCAGAGCTGCTGGGCGTGATGAGGCGCCTGTGGGCAGACCCAGGAGCACAGGCATGCTTTTCCAGGTCTAGCGAGTATCACCTGGAGGACAATGCCGCCTACTATCTGAACGATCTGGAGAGGATCGCAGCAGCAGACTACATCCCTACCGTGGAGGACATCCTGAGGAGCCGCGATATGACCACAGGCATCGTGGAGAACAAGTTCACCTTCAAGGAGCTGACCTTCAAGATGGTGGACGTGGGAGGACAGCGGTCCGAGAGAAAGAAGTGGATCCACTGCTTCGAGGGCGTGACAGCCATCATCTTCTGCGTGGAGCTGTCTGGCTACGACCTGAAGCTGTATGAGGATAACCAGACCTCCAGGATGGCCGAGTCTCTGCGCCTGTTTGATAGCATCTGCAACAATAACTGGTTCATCAATACCTCCCTGATCCTGTTTCTGAACAAGAAGGACCTGCTGGCCGAGAAGATCCGGAGAATCCCCCTGACAATCTGTTTCCCTGAGTACAAGGGCCAGAATACCTATGAGGAGGCCGCCGTGTACATCCAGCGGCAGTTTGAGGATCTGAATAGAAACAAGGAGACAAAGGAGATCTACAGCCACTTCACCTGTGCCACAGACACCTCCAACATCCAGTTTGTCTTTGATGCTGTGACCGACGTGATTATCCAGAACAACCTGAAATACATTGGGCTGTGCTGA

Gα_q_-linker-LgBiT

ATGACCCTGGAATCAATTATGGCATGTTGTCTGAGCGAGGAAGCAAAGGAGGCAAGGAGAATCAATGACGAAATCGAGAGGCAACTGCGGAGAGACAAGAGAGATGCCAGGCGCGAGCTGAAGCTGCTGCTGCTGGGCACAGGAGAGAGCGGCAAGTCCACCTTCATCAAGCAGATGAGAATCATCCACGGCTCTGGCTACAGCGACGAGGATAAGAGGGGCTTCACAAAGCTGGTGTATCAGAACATCTTTACCGCCATGCAGGCCATGATCAGGGCCATGGATACACTGGGAGGCAGCGGAGGAGGAGGCTCCGGAGGCAGCTCCTCTGGCGGCGTGTTCACACTGGAGGACTTTGTGGGCGATTGGGAGCAGACCGCCGCCTACAACCTGGACCAGGTGCTGGAGCAGGGAGGCGTGAGCAGCCTGCTCCAGAATCTGGCCGTGTCTGTGACCCCAATCCAGAGGATCGTGCGGAGCGGAGAGAATGCCCTGAAGATCGATATCCACGTGATCATCCCTTATGAGGGCCTGAGCGCCGACCAGATGGCACAGATCGAGGAGGTGTTCAAGGTGGTGTACCCAGTGGACGATCACCACTTCAAAGTGATCCTGCCCTATGGCACCCTGGTCATCGATGGCGTGACCCCCAACATGCTGAATTACTTCGGCCGCCCTTATGAGGGCATCGCCGTGTTTGACGGCAAGAAGATCACCGTGACAGGCACCCTGTGGAACGGCAATAAGATCATCGACGAGCGCCTGATCACACCAGATGGCTCTATGCTGTTCCGGGTGACCATCAACTCCGGAGGCTCTGGCGGCGGAGGCTCCGGAGGCTCTAGCTCCGGAGGCAAGATCCCCTACAAGTATGAGCACAATAAGGCACACGCACAGCTCGTGCGGGAGGTGGATGTGGAGAAGGTGTCCGCCTTTGAGAACCCTTACGTGGACGCCATCAAGTCTCTGTGGAATGATCCAGGCATCCAGGAGTGCTACGACCGGAGAAGGGAGTATCAGCTCTCCGACTCTACAAAGTACTATCTGAACGACCTGGATAGGGTGGCCGATCCCGCATACCTGCCTACCCAGCAGGACGTGCTGAGAGTGAGGGTGCCTACCACAGGCATCATCGAGTATCCCTTCGACCTCCAGAGCGTGATCTTTCGCATGGTGGACGTGGGAGGACAGCGGTCCGAGAGGCGGAAGTGGATTCACTGTTTCGAGAATGTGACCTCCATCATGTTTCTGGTGGCCCTGTCTGAGTACGACCAGGTGCTGGTGGAGAGCGATAACGAGAATAGAATGGAGGAGTCCAAGGCCCTGTTCAGGACAATCATCACCTACCCTTGGTTCCAGAACTCTAGCGTGATCCTGTTTCTGAATAAGAAGGATCTGCTGGAGGAGAAGATCATGTATTCCCACCTGGTGGACTACTTTCCAGAGTATGACGGACCACAGAGAGATGCACAGGCAGCCAGGGAGTTCATCCTGAAGATGTTTGTGGATCTGAACCCCGACTCTGATAAGATCATCTACAGCCACTTCACCTGCGCCACAGACACCGAGAATATCAGATTTGTGTTTGCCGCCGTGAAAGATACAATCCTGCAACTGAACCTGAAAGAATACAACCTGGTGTGA

Gα_q_-linker-LgBiT (αB-αC)

ATGACTCTGGAAAGCATTATGGCATGTTGTCTGTCCGAGGAGGCAAAGGAGGCAAGGAGAATTAACGACGAGATTGAACGGCAACTGCGGAGAGACAAGAGAGATGCCAGGCGCGAGCTGAAGCTGCTGCTGCTGGGCACCGGAGAGTCCGGCAAGTCTACATTCATCAAGCAGATGAGAATCATCCACGGCAGCGGCTACTCCGACGAGGATAAGAGGGGCTTCACCAAGCTGGTGTATCAGAACATCTTTACAGCCATGCAGGCCATGATCCGCGCCATGGATACCCTGAAGATCCCTTACAAGTATGAGCACAATAAGGCACACGCACAGCTCGTGCGGGAGGTGGACGTGGAGAAGGTGTCTGCCGGAGGCAGCGGAGGAGGAGGCTCCGGAGGCAGCTCCTCTGGCGGCGTGTTTACTCTGGAGGACTTCGTCGGAGACTGGGAACAGACTGCTGCTTACAATCTGGATCAGGTGCTGGAACAGGGGGGGGTCAGCTCCCTGCTCCAGAACCTGGCCGTGTCTGTGACACCTATCCAGCGGATCGTGAGAAGCGGCGAGAATGCCCTGAAGATCGACATCCACGTGATCATCCCATACGAGGGCCTGTCCGCCGATCAGATGGCCCAGATCGAGGAGGTGTTCAAGGTGGTGTACCCAGTGGACGATCACCACTTCAAAGTGATCCTGCCCTATGGCACCCTGGTCATCGACGGAGTGACCCCAAACATGCTGAATTACTTCGGCAGGCCTTATGAGGGCATCGCCGTGTTTGATGGCAAGAAGATCACCGTGACAGGCACCCTGTGGAACGGCAATAAGATCATCGACGAGCGGCTGATCACCCCCGATGGCTCTATGCTGTTCAGAGTGACCATCAATAGCGGAGGCTCTGGCGGCGGAGGCTCCGGAGGCTCTAGCTCCGGAGGCTTTGAGAACCCCTACGTGGACGCCATCAAGAGCCTGTGGAATGATCCTGGCATCCAGGAGTGCTACGACCGGAGAAGGGAGTATCAGCTCTCTGATAGCACCAAGTACTATCTGAACGACCTGGATAGGGTGGCCGATCCCGCCTACCTGCCTACACAGCAGGACGTGCTGAGAGTGAGGGTGCCTACCACAGGCATCATCGAGTATCCCTTCGACCTCCAGTCCGTGATCTTTAGAATGGTGGACGTGGGAGGACAGAGGTCTGAGAGGAGGAAGTGGATTCACTGTTTCGAGAATGTGACCTCCATCATGTTTCTGGTGGCCCTGTCTGAGTACGACCAGGTGCTGGTGGAGAGCGATAACGAGAATCGCATGGAGGAGTCCAAGGCCCTGTTCCGGACCATCATCACATACCCATGGTTCCAGAACAGCTCCGTGATCCTGTTTCTGAATAAGAAGGATCTGCTGGAGGAGAAGATCATGTATAGCCACCTGGTGGACTACTTTCCAGAGTATGACGGACCACAGAGGGATGCACAGGCAGCCCGGGAGTTCATCCTGAAGATGTTTGTGGATCTGAACCCCGACTCTGATAAGATCATCTATAGCCACTTCACATGCGCCACCGACACAGAGAATATCCGGTTTGTGTTTGCCGCCGTGAAAGATACAATCCTTCAGCTTAACCTGAAAGAATACAACCTGGTGTGA

Gα_11_-linker-LgBiT

ATGACCCTGGAGAGTATGATGGCATGTTGTCTGAGCGACGAAGTGAAGGAGAGCAAGAGGATTAACGCCGAGATTGAGAAGCAGCTGCGGAGAGACAAGAGAGATGCCAGGCGCGAGCTGAAGCTGCTGCTGCTGGGCACAGGAGAGAGCGGCAAGTCCACCTTCATCAAGCAGATGAGAATCATCCACGGCGCCGGCTACAGCGAGGAGGACAAGAGGGGCTTCACAAAGCTGGTGTATCAGAACATCTTTACCGCCATGCAGGCCATGATCAGGGCCATGGAGACACTGGGAGGCTCCGGAGGAGGAGGCTCTGGAGGCAGCTCCTCTGGCGGCGTGTTCACACTGGAGGACTTTGTGGGCGATTGGGAGCAGACCGCCGCCTACAACCTGGATCAGGTGCTGGAGCAGGGAGGCGTGAGCAGCCTGCTCCAGAATCTGGCCGTGAGCGTGACCCCAATCCAGAGGATCGTGCGGTCCGGAGAGAATGCCCTGAAGATCGACATCCACGTGATCATCCCCTATGAGGGCCTGTCCGCCGATCAGATGGCCCAGATCGAGGAGGTGTTCAAGGTGGTGTACCCCGTGGACGATCACCACTTCAAAGTGATCCTGCCTTATGGCACCCTGGTCATCGACGGCGTGACCCCTAACATGCTGAATTACTTCGGCCGCCCATATGAGGGCATCGCCGTGTTTGATGGCAAGAAGATCACCGTGACAGGCACCCTGTGGAACGGCAATAAGATCATCGACGAGCGCCTGATCACACCCGATGGCTCTATGCTGTTCCGGGTGACCATCAACTCTGGAGGCAGCGGAGGAGGAGGCAGCGGAGGCTCTAGCTCCGGAGGCAAGATCCTGTACAAGTATGAGCAGAACAAGGCCAATGCCCTGCTGATCCGCGAGGTGGACGTGGAGAAGGTGACCACATTTGAGCACCAGTACGTGAGCGCCATCAAGACACTGTGGGAGGACCCTGGCATCCAGGAGTGCTACGATCGGAGAAGGGAGTATCAGCTCTCCGACTCTGCCAAGTACTATCTGACCGACGTGGATCGGATCGCCACACTGGGCTACCTGCCAACCCAGCAGGATGTGCTGAGAGTGAGGGTGCCAACCACAGGCATCATCGAGTATCCCTTCGACCTGGAGAACATCATCTTTCGCATGGTGGATGTGGGAGGACAGCGGTCCGAGAGGCGGAAGTGGATTCACTGTTTCGAGAATGTGACCTCTATCATGTTTCTGGTGGCCCTGAGCGAGTACGACCAGGTGCTGGTGGAGTCCGATAACGAGAATAGAATGGAGGAGTCTAAGGCCCTGTTCAGGACAATCATCACCTATCCCTGGTTCCAGAACTCTAGCGTGATCCTGTTTCTGAATAAGAAGGACCTGCTGGAGGATAAGATCCTGTACTCTCACCTGGTGGACTATTTCCCCGAGTTTGACGGACCTCAGAGAGATGCACAGGCAGCCAGGGAGTTCATCCTGAAGATGTTTGTGGACCTGAACCCTGACAGCGATAAGATCATCTACTCCCACTTCACCTGCGCCACAGATACCGAGAATATCAGATTTGTGTTCGCCGCCGTGAAAGACACTATCCTGCAGCTGAATCTGAAAGAATACAACCTGGTCTGA

Gα_14_-linker-LgBiT (αB-αC)

ATGGCCGGCTGCTGCTGCCTGTCCGCGGAGGAGAAGGAGTCGCAGCGCATCAGCGCGGAGATCGAGCGACAGCTTCGTCGGGACAAGAAGGACGCGCGCCGTGAGCTTAAGCTGCTGCTGCTGGGAACTGGTGAAAGTGGGAAAAGCACCTTTATCAAGCAGATGAGAATTATCCATGGGTCTGGTTACAGCGACGAAGACAGAAAGGGGTTCACGAAGCTGGTTTACCAAAACATATTCACCGCCATGCAAGCCATGATCAGAGCGATGGACACGCTAAGGATACAGTATGTGTGTGAACAGAATAAGGAAAATGCCCAGATAATCAGAGAAGTGGAAGTGGACAAGGTCTCCATGGGAGGCAGCGGAGGAGGAGGCTCCGGAGGCAGCTCCTCTGGCGGCGTGTTTACTCTGGAGGACTTCGTCGGAGACTGGGAACAGACTGCTGCTTACAATCTGGATCAGGTGCTGGAACAGGGGGGGGTCAGCTCCCTGCTCCAGAACCTGGCCGTGTCTGTGACACCTATCCAGCGGATCGTGAGAAGCGGCGAGAATGCCCTGAAGATCGACATCCACGTGATCATCCCATACGAGGGCCTGTCCGCCGATCAGATGGCCCAGATCGAGGAGGTGTTCAAGGTGGTGTACCCAGTGGACGATCACCACTTCAAAGTGATCCTGCCCTATGGCACCCTGGTCATCGACGGAGTGACCCCAAACATGCTGAATTACTTCGGCAGGCCTTATGAGGGCATCGCCGTGTTTGATGGCAAGAAGATCACCGTGACAGGCACCCTGTGGAACGGCAATAAGATCATCGACGAGCGGCTGATCACCCCCGATGGCTCTATGCTGTTCAGAGTGACCATCAATAGCGGAGGCTCTGGCGGCGGAGGCTCCGGAGGCTCTAGCTCCGGAGGCCTCTCCAGGGAGCAGGTGGAGGCCATCAAGCAGCTCTGGCAAGATCCAGGCATCCAGGAGTGTTACGACAGGAGGAGGGAGTACCAGCTGTCGGACTCTGCCAAATATTACCTGACTGACATTGACCGCATCGCCACACCATCATTCGTGCCTACCCAACAAGATGTGCTTCGCGTCCGAGTGCCCACCACCGGCATCATTGAGTATCCATTTGACTTGGAAAACATCATCTTTCGGATGGTGGATGTTGGTGGCCAACGATCGGAAAGACGGAAGTGGATTCACTGCTTTGAGAGTGTCACCTCCATTATTTTCTTGGTTGCTCTGAGTGAATATGACCAGGTCCTGGCTGAGTGTGACAACGAGAATCGCATGGAAGAGAGCAAAGCCTTATTTAAAACCATCATCACCTACCCCTGGTTTCTGAATTCGTCTGTGATTTTATTCTTGAACAAGAAGGATCTTTTGGAAGAGAAAATCATGTACTCTCATCTAATTAGCTATTTCCCAGAATACACAGGACCGAAACAGGATGTCAGAGCTGCCAGAGACTTTATCCTGAAGCTTTACCAAGATCAGAATCCTGACAAAGAGAAAGTCATCTACTCTCACTTCACATGTGCTACAGATACAGACAATATTCGCTTTGTGTTTGCTGCTGTCAAAGACACAATTCTACAGCTAAACCTAAGGGAATTCAACCTTGTCTGA

Gα_16_-linker-LgBiT (αB-αC)

ATGGCCCGCTCGCTGACCTGGCGCTGCTGCCCCTGGTGCCTGACGGAGGATGAGAAGGCCGCCGCCCGGGTGGACCAGGAGATCAACAGGATCCTCTTGGAGCAGAAGAAGCAGGACCGCGGGGAGCTGAAGCTGCTGCTTTTGGGCCCAGGCGAGAGCGGGAAGAGCACCTTCATCAAGCAGATGCGGATCATCCACGGCGCCGGCTACTCGGAGGAGGAGCGCAAGGGCTTCCGGCCCCTGGTCTACCAGAACATCTTCGTGTCCATGCGGGCCATGATCGAGGCCATGGAGCGGCTGCAGATTCCATTCAGCAGGCCCGAGAGCAAGCACCACGCTAGCCTGGTCATGAGCCAGGACCCCTATAAAGTGACCACGGGAGGCAGCGGAGGAGGAGGCTCCGGAGGCAGCTCCTCTGGCGGCGTGTTTACTCTGGAGGACTTCGTCGGAGACTGGGAACAGACTGCTGCTTACAATCTGGATCAGGTGCTGGAACAGGGGGGGGTCAGCTCCCTGCTCCAGAACCTGGCCGTGTCTGTGACACCTATCCAGCGGATCGTGAGAAGCGGCGAGAATGCCCTGAAGATCGACATCCACGTGATCATCCCATACGAGGGCCTGTCCGCCGATCAGATGGCCCAGATCGAGGAGGTGTTCAAGGTGGTGTACCCAGTGGACGATCACCACTTCAAAGTGATCCTGCCCTATGGCACCCTGGTCATCGACGGAGTGACCCCAAACATGCTGAATTACTTCGGCAGGCCTTATGAGGGCATCGCCGTGTTTGATGGCAAGAAGATCACCGTGACAGGCACCCTGTGGAACGGCAATAAGATCATCGACGAGCGGCTGATCACCCCCGATGGCTCTATGCTGTTCAGAGTGACCATCAATAGCGGAGGCTCTGGCGGCGGAGGCTCCGGAGGCTCTAGCTCCGGAGGCTTTGAGAAGCGCTACGCTGCGGCCATGCAGTGGCTGTGGAGGGATGCCGGCATCCGGGCCTACTATGAGCGTCGGCGGGAATTCCACCTGCTCGATTCAGCCGTGTACTACCTGTCCCACCTGGAGCGCATCACCGAGGAGGGCTACGTCCCCACAGCTCAGGACGTGCTCCGCAGCCGCATGCCCACCACTGGCATCAACGAGTACTGCTTCTCCGTGCAGAAAACCAACCTGCGGATCGTGGACGTCGGGGGCCAGAAGTCAGAGCGTAAGAAATGGATCCATTGTTTCGAGAACGTGATCGCCCTCATCTACCTGGCCTCACTGAGTGAATACGACCAGTGCCTGGAGGAGAACAACCAGGAGAACCGCATGAAGGAGAGCCTCGCATTGTTTGGGACTATCCTGGAACTACCCTGGTTCAAAAGCACATCCGTCATCCTCTTTCTCAACAAAACCGACATCCTGGAGGAGAAAATCCCCACCTCCCACCTGGCTACCTATTTCCCCAGTTTCCAGGGCCCTAAGCAGGATGCTGAGGCAGCCAAGAGGTTCATCCTGGACATGTACACGAGGATGTACACCGGGTGCGTGGACGGCCCCGAGGGCAGCAAGAAGGGCGCACGATCCCGACGCCTCTTCAGCCACTACACATGTGCCACAGACACACAGAACATCCGCAAGGTCTTCAAGGACGTGCGGGACTCGGTGCTCGCCCGCTACCTGGACGAGATCAACCTGCTGTGA

Gα_12_-linker-LgBiT

ATGTCAGGAGTCGTGCGAACACTGTCAAGATGTCTGCTGCCCGCTGAAGCTGGAGGAGCCCGAGAGAGGCGAGCCGGATCAGGAGCCAGAGACGCAGAGAGGGAGGCACGGAGAAGGTCCAGAGACATCGATGCACTGCTGGCCAGGGAGAGGAGGGCCGTGAGAAGGCTGGTGAAGATCCTGCTGCTGGGAGCAGGAGAGTCTGGCAAGAGCACATTCCTGAAGCAGATGCGCATCATCCACGGCCGGGAGTTTGATCAGAAGGCCCTGCTGGAGTTCAGAGACACCATCTTTGATAACATCCTGAAGGGCAGCCGCGTGCTGGTGGACGCCCGGGATAAGCTGGGAGGCTCTGGAGGAGGAGGCAGCGGAGGCAGCTCCTCTGGCGGCGTGTTCACACTGGAGGACTTTGTGGGCGATTGGGAGCAGACCGCCGCCTACAACCTGGACCAGGTGCTGGAGCAGGGAGGCGTGAGCAGCCTGCTCCAGAATCTGGCCGTGAGCGTGACACCAATCCAGAGAATCGTGAGGTCCGGCGAGAATGCCCTGAAGATCGACATCCACGTGATCATCCCCTATGAGGGCCTGTCCGCCGATCAGATGGCCCAGATCGAGGAGGTGTTCAAGGTGGTGTACCCCGTGGACGATCACCACTTCAAAGTGATCCTGCCTTATGGCACCCTGGTCATCGACGGCGTGACCCCTAACATGCTGAATTACTTCGGCAGGCCATATGAGGGCATCGCCGTGTTTGATGGCAAGAAGATCACCGTGACAGGCACCCTGTGGAACGGCAATAAGATCATCGACGAGCGCCTGATCACACCCGATGGCAGCATGCTGTTCCGGGTGACCATCAACTCCGGCGGCTCTGGCGGAGGAGGCTCTGGAGGCTCTAGCTCCGGAGGAGGCATCCCTTGGCAGTACAGCGAGAACGAGAAGCACGGCATGTTCCTGATGGCCTTTGAGAATAAGGCAGGCCTGCCAGTGGAGCCTGCCACCTTCCAGCTCTATGTGCCAGCCCTGTCCGCCCTGTGGAGAGACTCTGGCATCAGGGAGGCCTTCAGCCGCCGGTCCGAGTTTCAGCTCGGCGAGTCTGTGAAGTACTTCCTGGACAACCTGGATAGAATCGGCCAGCTCAACTATTTTCCTAGCAAGCAGGATATCCTGCTGGCCAGGAAGGCCACAAAGGGCATCGTGGAGCACGACTTCGTGATCAAGAAGATCCCCTTCAAGATGGTGGATGTGGGAGGACAGCGGAGCCAGCGGCAGAAGTGGTTCCAGTGCTTTGACGGCATCACCTCTATCCTGTTCATGGTGTCTAGCTCCGAGTACGACCAGGTGCTGATGGAGGATAGAAGGACAAACCGGCTGGTGGAGAGCATGAATATCTTCGAGACAATCGTGAACAATAAGCTGTTCTTTAACGTGTCCATCATCCTGTTTCTGAATAAGATGGATCTGCTGGTGGAGAAGGTGAAGACAGTGTCCATCAAGAAGCACTTCCCAGACTTTCGCGGCGATCCCCACCGGCTGGAGGACGTGCAGAGATACCTGGTGCAGTGTTTCGATAGAAAGCGCCGGAACAGGTCTAAGCCTCTGTTCCACCACTTTACCACAGCCATCGACACCGAGAATGTGCGGTTTGTGTTCCACGCTGTGAAGGATACTATTCTGCAAGAAAATCTGAAAGACATTATGCTCCAGTGA

Gα_13_-linker-LgBiT

ATGGCCGATTTCCTGCCCTCAAGAAGCGTCCTGAGCGTGTGCTTTCCTGGATGTCTGCTGACTAGCGGAGAAGCCGAACAGCAGAGAAAGAGCAAGGAGATCGACAAGTGCCTGTCCAGAGAGAAGACATACGTGAAGAGGCTGGTGAAGATCCTGCTGCTGGGAGCAGGAGAGTCTGGCAAGAGCACCTTCCTGAAGCAGATGAGAATCATCCACGGACAGGACTTCGATCAGCGCGCCCGGGAGGAGTTTAGGCCTACAATCTATTCCAACGTGATCAAGGGAATGCGCGTGCTGGTGGACGCCCGGGAGAAGCTGGGAGGCTCCGGCGGAGGAGGCTCTGGAGGCAGCTCCTCTGGCGGCGTGTTCACACTGGAGGACTTTGTGGGCGATTGGGAGCAGACCGCCGCCTACAACCTGGATCAGGTGCTGGAGCAGGGAGGCGTGAGCAGCCTGCTCCAGAATCTGGCCGTGTCTGTGACCCCTATCCAGAGGATCGTGCGGAGCGGAGAGAATGCCCTGAAGATCGACATCCACGTGATCATCCCATATGAGGGCCTGAGCGCCGATCAGATGGCCCAGATCGAGGAGGTGTTCAAGGTGGTGTACCCTGTGGACGATCACCACTTCAAAGTGATCCTGCCATATGGCACCCTGGTCATCGACGGCGTGACCCCAAACATGCTGAATTACTTCGGCAGACCCTATGAGGGCATCGCCGTGTTTGATGGCAAGAAGATCACCGTGACAGGCACCCTGTGGAACGGCAATAAGATCATCGACGAGAGACTGATCACACCAGATGGCAGCATGCTGTTCAGGGTGACCATCAACTCCGGAGGCTCTGGAGGAGGAGGCTCCGGAGGCTCTAGCTCCGGCGGCCACATCCCCTGGGGCGACAACTCCAATCAGCAGCACGGCGACAAGATGATGTCTTTCGATACAAGGGCCCCAATGGCAGCACAGGGAATGGTGGAGACACGGGTGTTCCTCCAGTACCTGCCAGCAATCCGGGCCCTGTGGGCAGACTCTGGCATCCAGAACGCCTATGATCGGAGAAGGGAGTTCCAGCTCGGCGAGAGCGTGAAGTACTTTCTGGACAATCTGGATAAGCTGGGCGAGCCCGACTATATCCCTTCCCAGCAGGACATCCTGCTGGCTAGACGGCCCACAAAGGGCATCCACGAGTACGACTTCGAGATCAAGAACGTGCCTTTTAAGATGGTGGATGTGGGAGGACAGCGGAGCGAGAGAAAGAGGTGGTTCGAGTGTTTTGACAGCGTGACCTCCATCCTGTTCCTGGTGTCTAGCTCCGAGTTTGACCAGGTGCTGATGGAGGATAGACTGACAAACAGGCTGACCGAGTCCCTGAATATCTTCGAGACAATCGTGAACAATCGGGTGTTCTCTAACGTGAGCATCATCCTGTTTCTGAATAAGACCGACCTGCTGGAGGAGAAGGTGCAGATCGTGAGCATCAAGGATTACTTCCTGGAGTTTGAGGGCGACCCCCACTGCCTGAGGGATGTGCAGAAGTTCCTGGTGGAGTGTTTTCGGAATAAGAGAAGGGATCAGCAGCAGAAGCCTCTGTATCACCACTTTACCACAGCCATCAACACCGAGAATATCAGACTGGTCTTTAGGGATGTGAAGGACACCATTCTGCACGACAATCTGAAACAGCTCATGCTTCAGTGA

SmBiT-linker-Gβ_1_

ATGGTCACCGGCTATCGGCTGTTTGAAGAGATTCTGGGGGGGTCAGGAGGAGGAGGCTCAGGGGGCTCATCATCAGGAGGAAGTGAGCTTGACCAGTTACGGCAGGAGGCCGAGCAACTTAAGAACCAGATTCGAGACGCCAGGAAAGCATGTGCAGATGCAACTCTCTCTCAGATCACAAACAACATCGACCCAGTGGGAAGAATCCAAATGCGCACGAGGAGGACACTGCGGGGGCACCTGGCCAAGATCTACGCCATGCACTGGGGCACAGACTCCAGGCTTCTCGTCAGTGCCTCGCAGGATGGTAAACTTATCATCTGGGACAGCTACACCACCAACAAGGTCCACGCCATCCCTCTGCGCTCCTCCTGGGTCATGACCTGTGCATATGCCCCTTCTGGGAACTATGTGGCCTGCGGTGGCCTGGATAACATTTGCTCCATTTACAATCTGAAAACTCGTGAGGGGAACGTGCGCGTGAGTCGTGAGCTGGCAGGACACACAGGTTACCTGTCCTGCTGCCGATTCCTGGATGACAATCAGATCGTCACCAGCTCTGGAGACACCACGTGTGCCCTGTGGGACATCGAGACCGGCCAGCAGACGACCACGTTTACCGGACACACTGGAGATGTCATGAGCCTTTCTCTTGCTCCTGACACCAGACTGTTCGTCTCTGGTGCTTGTGATGCTTCAGCCAAACTCTGGGATGTGCGAGAAGGCATGTGCCGGCAGACCTTCACTGGCCACGAGTCTGACATCAATGCCATTTGCTTCTTTCCAAATGGCAATGCATTTGCCACTGGCTCAGACGACGCCACCTGCAGGCTGTTTGACCTTCGTGCTGACCAGGAGCTCATGACTTACTCCCATGACAACATCATCTGCGGGATCACCTCTGTCTCCTTCTCCAAGAGCGGGCGCCTCCTCCTTGCTGGGTACGACGACTTCAACTGCAACGTCTGGGATGCACTCAAAGCCGACCGGGCAGGTGTCTTGGCTGGGCATGACAACCGCGTCAGCTGCCTGGGCGTGACTGACGATGGCATGGCTGTGGCGACAGGGTCCTGGGATAGCTTCCTCAAGATCTGGAACTGA

SmBiT-linker-Gβ_2_

ATGGTCACCGGCTATCGGCTGTTTGAAGAGATTCTGGGGGGGTCAGGAGGAGGAGGCTCAGGGGGCTCATCATCAGGAGGAAGTGAGCTGGAGCAACTGAGACAGGAGGCCGAGCAGCTCCGGAACCAGATCCGGGATGCCCGAAAAGCATGTGGGGACTCAACACTGACCCAGATCACAGCTGGGCTGGACCCAGTGGGGAGAATCCAGATGAGGACCCGGAGGACCCTCCGTGGGCACCTGGCAAAGATCTATGCCATGCACTGGGGGACCGACTCAAGGCTGCTGGTCAGCGCCTCCCAGGATGGGAAGCTCATCATCTGGGACAGCTACACCACCAACAAGGTCCACGCCATCCCGCTGCGCTCCTCCTGGGTAATGACCTGTGCCTACGCGCCCTCAGGGAACTTTGTGGCCTGTGGGGGGTTGGACAACATCTGCTCCATCTACAGCCTCAAGACCCGCGAGGGCAACGTCAGGGTCAGCCGGGAGCTGCCTGGCCACACTGGGTACCTGTCGTGTTGCCGCTTCCTGGATGACAACCAAATCATCACCAGCTCTGGGGATACCACCTGTGCCCTGTGGGACATTGAGACAGGCCAGCAGACAGTGGGTTTTGCTGGACACAGTGGGGATGTGATGTCCCTGTCCCTGGCCCCCGATGGCCGCACGTTTGTGTCAGGCGCCTGTGATGCCTCTATCAAGCTGTGGGACGTGCGGGATTCCATGTGCCGACAGACCTTCATCGGCCATGAATCCGACATCAATGCAGTGGCTTTCTTCCCCAACGGCTACGCCTTCACCACCGGCTCTGACGACGCCACGTGCCGCCTCTTCGACCTGCGGGCCGATCAGGAGCTCCTCATGTACTCCCATGACAACATCATCTGTGGCATCACCTCTGTTGCCTTCTCGCGCAGCGGACGGCTGCTGCTCGCTGGCTACGACGACTTCAACTGCAACATCTGGGATGCCATGAAGGGCGACCGTGCAGGAGTCCTCGCTGGCCACGACAACCGCGTGAGCTGCCTCGGGGTCACCGACGATGGCATGGCTGTGGCCACGGGCTCCTGGGACTCCTTCCTCAAGATCTGGAACTGA

SmBiT-linker-Gβ_3_

ATGGTCACCGGCTATCGGCTGTTTGAAGAGATTCTGGGGGGGTCAGGAGGAGGAGGCTCAGGGGGCTCATCATCAGGAGGAGGGGAGATGGAGCAACTGCGTCAGGAAGCGGAGCAGCTCAAGAAGCAGATTGCAGATGCCAGGAAAGCCTGTGCTGACGTTACTCTGGCAGAGCTGGTGTCTGGCCTAGAGGTGGTGGGACGAGTCCAGATGCGGACGCGGCGGACGTTAAGGGGACACCTGGCCAAGATTTACGCCATGCACTGGGCCACTGATTCTAAGCTGCTGGTAAGTGCCTCGCAAGATGGGAAGCTGATCGTGTGGGACAGCTACACCACCAACAAGGTGCACGCCATCCCACTGCGCTCCTCCTGGGTCATGACCTGTGCCTATGCCCCATCAGGGAACTTTGTGGCATGTGGGGGGCTGGACAACATGTGTTCCATCTACAACCTCAAATCCCGTGAGGGCAATGTCAAGGTCAGCCGGGAGCTTTCTGCTCACACAGGTTATCTCTCCTGCTGCCGCTTCCTGGATGACAACAATATTGTGACCAGCTCGGGGGACACCACGTGTGCCTTGTGGGACATTGAGACTGGGCAGCAGAAGACTGTATTTGTGGGACACACGGGTGACTGCATGAGCCTGGCTGTGTCTCCTGACTTCAATCTCTTCATTTCGGGGGCCTGTGATGCCAGTGCCAAGCTCTGGGATGTGCGAGAGGGGACCTGCCGTCAGACTTTCACTGGCCACGAGTCGGACATCAACGCCATCTGTTTCTTCCCCAATGGAGAGGCCATCTGCACGGGCTCGGATGACGCTTCCTGCCGCTTGTTTGACCTGCGGGCAGACCAGGAGCTGATCTGCTTCTCCCACGAGAGCATCATCTGCGGCATCACGTCCGTGGCCTTCTCCCTCAGTGGCCGCCTACTATTCGCTGGCTACGACGACTTCAACTGCAATGTCTGGGACTCCATGAAGTCTGAGCGTGTGGGCATCCTCTCTGGCCACGATAACAGGGTGAGCTGCCTGGGAGTCACAGCTGACGGGATGGCTGTGGCCACAGGTTCCTGGGACAGCTTCCTCAAAATCTGGAACTGA

SmBiT-linker-Gβ_4_

ATGGTCACCGGCTATCGGCTGTTTGAAGAGATTCTGGGGGGGTCAGGAGGAGGAGGCTCAGGGGGCTCATCATCAGGAGGAAGCGAACTGGAACAGTTGAGGCAAGAAGCAGAACAACTGCGGAATCAGATTCAGGATGCTCGGAAAGCATGTAATGATGCAACGCTTGTTCAGATTACATCAAATATGGACTCTGTGGGTCGAATACAAATGCGAACAAGACGTACACTGAGGGGCCACCTAGCTAAAATCTATGCTATGCATTGGGGATACGATTCCAGGCTGCTAGTCAGTGCTTCTCAAGATGGAAAATTAATTATTTGGGATAGCTATACAACAAATAAGATGCATGCTATTCCTTTGAGGTCCTCCTGGGTGATGACCTGTGCTTATGCTCCCTCTGGTAATTATGTTGCCTGTGGAGGCTTGGACAACATCTGCTCTATATATAACTTAAAGACCAGAGAGGGAAATGTGAGAGTAAGCCGAGAGTTGCCAGGTCACACAGGGTACTTGTCCTGCTGTCGTTTTTTAGATGACAGCCAAATTGTTACAAGTTCAGGAGATACAACTTGTGCTTTATGGGACATCGAAACTGCCCAGCAGACCACCACATTCACTGGGCATTCTGGAGATGTGATGAGTCTTTCTTTGAGTCCTGACATGAGGACTTTTGTTTCTGGTGCTTGTGATGCCTCTTCCAAATTATGGGATATTCGAGATGGAATGTGTAGACAGTCTTTCACGGGACATGTCTCAGATATCAATGCTGTCAGTTTTTTCCCAAATGGATATGCCTTCGCCACTGGCTCTGATGATGCCACTTGCCGGCTCTTTGACCTTCGTGCAGATCAAGAGTTATTATTGTATTCTCATGACAATATCATCTGTGGAATCACTTCTGTAGCCTTCTCAAAAAGTGGGCGTCTCTTGTTGGCTGGTTACGATGACTTTAATTGTAATGTATGGGACACGCTAAAAGGAGATCGTGCAGGTGTCCTTGCTGGTCATGACAACCGTGTGAGCTGCTTAGGTGTAACTGATGATGGCATGGCTGTGGCAACAGGCTCTTGGGACAGTTTTCTTAGAATCTGGAATTGA

SmBiT-linker-Gβ_5_

ATGGTCACCGGCTATCGGCTGTTTGAAGAGATTCTGGGGGGGTCAGGAGGAGGAGGCTCAGGGGGCTCATCATCAGGAGGATGTGATCAGACCTTTCTCGTTAATGTATTTGGCTCATGTGACAAATGTTTCAAACAACGAGCTCTGAGACCAGTTTTCAAGAAGTCTCAACAACTCAGCTACTGTTCAACATGTGCAGAAATTATGGCAACCGAGGGGCTGCACGAGAACGAGACGCTGGCGTCGCTGAAGAGCGAGGCCGAGAGCCTCAAGGGCAAGCTGGAGGAGGAGCGAGCCAAGCTGCACGATGTGGAGCTGCACCAGGTGGCGGAGCGGGTGGAGGCCCTGGGGCAGTTTGTCATGAAGACCAGAAGGACCCTCAAAGGCCACGGGAACAAAGTCCTGTGCATGGACTGGTGCAAAGATAAGAGGAGGATCGTGAGCTCGTCACAGGATGGGAAGGTGATCGTGTGGGATTCCTTCACCACAAACAAGGAGCACGCGGTCACCATGCCCTGCACGTGGGTGATGGCATGTGCTTATGCCCCATCGGGATGTGCCATTGCTTGTGGTGGTTTGGATAATAAGTGTTCTGTGTACCCCTTGACGTTTGACAAAAATGAAAACATGGCTGCCAAAAAGAAGTCTGTTGCTATGCACACCAACTACCTGTCGGCCTGCAGCTTCACCAACTCTGACATGCAGATCCTGACAGCGAGCGGCGATGGCACATGTGCCCTGTGGGACGTGGAGAGCGGGCAGCTGCTGCAGAGCTTCCACGGACATGGGGCTGACGTCCTCTGCTTGGACCTGGCCCCCTCAGAAACTGGAAACACCTTCGTGTCTGGGGGATGTGACAAGAAAGCCATGGTGTGGGACATGCGCTCCGGCCAGTGCGTGCAGGCCTTTGAAACACATGAATCTGACATCAACAGTGTCCGGTACTACCCCAGTGGAGATGCCTTTGCTTCAGGGTCAGATGACGCTACGTGTCGCCTCTATGACCTGCGGGCAGATAGGGAGGTTGCCATCTATTCCAAAGAAAGCATCATATTTGGAGCATCCAGCGTGGACTTCTCCCTCAGTGGTCGCCTGCTGTTTGCTGGATACAATGATTACACTATCAACGTCTGGGATGTTCTCAAAGGGTCCCGGGTCTCCATCCTGTTTGGACATGAAAACCGCGTTAGCACTCTACGAGTTTCCCCCGATGGGACTGCTTTCTGCTCTGGATCATGGGATCATACCCTCAGAGTCTGGGCCTGA

SmBiT-linker-Gγ_2_

ATGGTCACCGGCTATCGGCTGTTTGAAGAGATTCTGGGGGGGTCAGGAGGAGGAGGCTCAGGGGGCTCATCATCAGGAGGAGCCAGCAACAACACCGCCAGCATAGCACAAGCCAGGAAGCTGGTAGAGCAGCTTAAGATGGAAGCCAATATCGACAGGATAAAGGTGTCCAAGGCAGCTGCAGATTTGATGGCCTACTGTGAAGCACATGCCAAGGAAGACCCCCTCCTGACCCCTGTTCCGGCTTCAGAAAACCCGTTTAGGGAGAAGAAGTTTTTCTGTGCCATCCTTTGA

SmBiT-linker-Gγ_2_ (C68S)

ATGGTCACCGGCTATCGGCTGTTTGAAGAGATTCTGGGGGGGTCAGGAGGAGGAGGCTCAGGGGGCTCATCATCAGGAGGAGCCAGCAACAACACCGCCAGCATAGCACAAGCCAGGAAGCTGGTAGAGCAGCTTAAGATGGAAGCCAATATCGACAGGATAAAGGTGTCCAAGGCAGCTGCAGATTTGATGGCCTACTGTGAAGCACATGCCAAGGAAGACCCCCTCCTGACCCCTGTTCCGGCTTCAGAAAACCCGTTTAGGGAGAAGAAGTTTTTCAGCGCCATCCTTTGA

SmBiT-linker-Gγ_3_

ATGGTCACCGGCTATCGGCTGTTTGAAGAGATTCTGGGGGGGTCAGGAGGAGGAGGCTCAGGGGGCTCATCATCAGGAGGAAAAGGTGAGACCCCGGTGAACAGCACTATGAGTATTGGGCAAGCACGCAAGATGGTGGAACAGCTTAAGATTGAAGCCAGCTTGTGTCGGATAAAGGTGTCCAAGGCAGCAGCAGACCTGATGACTTACTGTGATGCCCACGCCTGTGAGGATCCCCTCATCACCCCTGTGCCCACTTCGGAGAACCCCTTCCGGGAGAAGAAGTTCTTCTGTGCTCTCCTCTGA

SmBiT-linker-Gγ_4_

ATGGTCACCGGCTATCGGCTGTTTGAAGAGATTCTGGGGGGGTCAGGAGGAGGAGGCTCAGGGGGCTCATCATCAGGAGGAAAAGAGGGCATGTCTAATAACAGCACCACTAGCATCTCCCAAGCCAGGAAAGCTGTGGAGCAGCTAAAGATGGAAGCCTGTATGGACAGGGTCAAGGTCTCCCAGGCAGCTGCGGACCTCCTGGCCTACTGTGAAGCTCACGTGCGGGAAGATCCTCTCATCATTCCAGTGCCTGCATCAGAAAACCCCTTTCGCGAGAAGAAGTTCTTTTGTACCATTCTCTGA

SmBiT-linker-Gγ_5_

ATGGTCACCGGCTATCGGCTGTTTGAAGAGATTCTGGGGGGGTCAGGAGGAGGAGGCTCAGGGGGCTCATCATCAGGAGGATCTGGCTCCTCCAGCGTCGCCGCTATGAAGAAAGTGGTTCAACAGCTCCGGCTGGAGGCCGGACTCAACCGCGTAAAAGTTTCCCAGGCAGCTGCAGACTTGAAACAGTTCTGTCTGCAGAATGCTCAACATGACCCTCTGCTGACTGGAGTATCTTCAAGTACAAATCCCTTCAGACCCCAGAAAGTCTGTTCCTTTTTGTGA

SmBiT-linker-Gγ_7_

ATGGTCACCGGCTATCGGCTGTTTGAAGAGATTCTGGGGGGGTCAGGAGGAGGAGGCTCAGGGGGCTCATCATCAGGAGGATCAGCCACTAACAACATAGCCCAGGCCCGGAAGCTGGTGGAACAGCTACGCATAGAAGCCGGGATTGAGCGCATCAAGGTCTCCAAAGCGGCGTCTGACCTCATGAGCTACTGTGAGCAACATGCCCGGAACGACCCCCTGCTGGTCGGAGTCCCTGCCTCGGAGAACCCCTTTAAGGACAAGAAACCTTGTATTATTTTATGA

SmBiT-linker-Gγ_8_

ATGGTCACCGGCTATCGGCTGTTTGAAGAGATTCTGGGGGGGTCAGGAGGAGGAGGCTCAGGGGGCTCATCATCAGGAGGATCCAACAACATGGCCAAGATTGCCGAGGCCCGCAAGACGGTGGAACAGCTGAAGCTGGAGGTGAACATCGACCGCATGAAGGTGTCGCAGGCAGCAGCGGAACTCCTGGCTTTCTGCGAGACGCATGCCAAAGATGACCCGCTGGTGACGCCAGTACCCGCCGCGGAGAACCCCTTCCGCGACAAGCGCCTCTTTTGTGTTCTGCTCTGA

SmBiT-linker-Gγ_10_

ATGGTCACCGGCTATCGGCTGTTTGAAGAGATTCTGGGGGGGTCAGGAGGAGGAGGCTCAGGGGGCTCATCATCAGGAGGATCCTCCGGGGCTAGCGCGAGCGCCCTGCAGCGCTTGGTAGAGCAGCTCAAGTTGGAGGCTGGCGTGGAGAGGATCAAGGTCTCTCAGGCAGCTGCAGAGCTTCAACAGTACTGTATGCAGAATGCCTGCAAGGATGCCCTGCTGGTGGGTGTTCCAGCTGGAAGTAACCCCTTCCGGGAGCCTAGATCCTGTGCTTTACTCTGA

SmBiT-linker-Gγ_11_

ATGGTCACCGGCTATCGGCTGTTTGAAGAGATTCTGGGGGGGTCAGGAGGAGGAGGCTCAGGGGGCTCATCATCAGGAGGACCTGCCCTTCACATCGAAGATTTGCCAGAGAAGGAAAAACTGAAAATGGAAGTTGAGCAGCTTCGCAAAGAAGTGAAGTTGCAGAGACAACAAGTGTCTAAATGTTCTGAAGAAATAAAGAACTATATTGAAGAACGTTCTGGAGAGGATCCTCTAGTAAAGGGAATTCCAGAAGACAAGAACCCCTTTAAAGAAAAAGGCAGCTGTGTTATTTCATGA

SmBiT-linker-Gγ_12_

ATGGTCACCGGCTATCGGCTGTTTGAAGAGATTCTGGGGGGGTCAGGAGGAGGAGGCTCAGGGGGCTCATCATCAGGAGGATCCAGCAAAACAGCAAGCACCAACAATATAGCCCAGGCAAGGAGAACTGTGCAGCAGTTAAGATTAGAAGCCTCCATTGAAAGAATAAAGGTTTCGAAGGCATCAGCGGACCTCATGTCCTACTGTGAGGAACATGCCAGGAGTGACCCTTTGCTGATAGGAATACCAACTTCAGAAAACCCTTTCAAGGATAAAAAAACTTGCATCATCTTATGA

SmBiT-linker-Gγ_13_

ATGGTCACCGGCTATCGGCTGTTTGAAGAGATTCTGGGGGGGTCAGGAGGAGGAGGCTCAGGGGGCTCATCATCAGGAGGAGAGGAGTGGGACGTGCCACAGATGAAGAAAGAGGTGGAGAGCCTCAAGTACCAGCTGGCCTTCCAGCGGGAGATGGCGTCCAAGACCATCCCCGAGCTGCTGAAGTGGATCGAGGACGGGATCCCCAAGGACCCCTTCCTGAACCCCGACCTGATGAAGAACAACCCATGGGTGGAAAAGGGCAAATGCACCATCCTGTGA

SmBiT-linker-Gγ_T1_

ATGGTCACCGGCTATCGGCTGTTTGAAGAGATTCTGGGGGGGTCAGGAGGAGGAGGCTCAGGGGGCTCATCATCAGGAGGACCAGTAATCAATATTGAGGACCTGACAGAAAAGGACAAATTGAAGATGGAAGTTGACCAGCTCAAGAAAGAAGTGACACTGGAAAGAATGCTAGTTTCCAAATGTTGTGAAGAAGTAAGAGATTACGTTGAAGAACGATCTGGCGAGGATCCACTGGTAAAGGGCATCCCAGAGGACAAAAATCCCTTCAAGGAGCTCAAAGGAGGCTGTGTGATTTCATGA

SmBiT-linker-Gγ_T2_

ATGGTCACCGGCTATCGGCTGTTTGAAGAGATTCTGGGGGGGTCAGGAGGAGGAGGCTCAGGGGGCTCATCATCAGGAGGAGCCCAGGATCTCAGCGAGAAGGACCTGTTGAAGATGGAGGTGGAGCAGCTGAAGAAAGAAGTGAAAAACACAAGAATTCCGATTTCCAAAGCGGGAAAGGAAATCAAGGAGTACGTGGAGGCCCAAGCAGGAAACGATCCTTTTCTCAAAGGCATCCCTGAGGACAAGAATCCCTTCAAGGAGAAAGGTGGCTGTCTGATAAGCTGA

SmBiT-linker-AC1

ATGGTCACCGGCTATCGGCTGTTTGAAGAGATTCTGGGGGGGTCAGGAGGAGGAGGCTCAGGGGGCTCATCATCAGGAGGAGCGGGGGCGCCGCGCGGCGGAGGCGGCGGCGGAGGCGGCGCGGGCGAGCCCGGGGGCGCCGAGCGGGCGGCCGGGACAAGCCGCCGGCGCGGGCTCCGGGCGTGCGACGAGGAGTTCGCTTGCCCAGAGCTGGAGGCGCTGTTCCGCGGCTACACGCTGCGGCTGGAGCAGGCGGCCACGCTGAAGGCGCTGGCCGTTCTCAGCCTGCTGGCGGGCGCGCTGGCGCTGGCCGAGCTGCTGGGCGCGCCGGGGCCCGCGCCCGGCCTGGCCAAGGGCTCACACCCGGTGCACTGCGTCCTCTTCCTGGCGCTGCTCGTGGTAACCAACGTCCGGTCCCTGCAGGTGCCCCAGCTGCAGCAGGTCGGCCAGCTGGCGCTGCTCTTCAGCCTCACCTTCGCGCTGCTCTGCTGTCCTTTCGCGCTGGGCGGCCCCGCCCGGGGTTCCGCCGGGGCCGCTGGGGGGCCAGCGACCGCCGAACAAGGGGTTTGGCAGCTCCTTTTGGTCACCTTCGTGTCCTATGCCTTGCTGCCCGTGCGCAGCCTGCTGGCCATAGGCTTTGGGCTCGTGGTGGCTGCGTCGCACTTGCTGGTCACAGCCACCTTGGTCCCCGCCAAGCGCCCACGTCTCTGGAGGACGCTCGGTGCCAATGCCTTGCTCTTCGTCGGTGTGAACATGTATGGGGTCTTTGTGCGGATTCTGACTGAGCGTTCACAGAGGAAGGCGTTCCTGCAGGCCCGGAGCTGCATTGAGGACCGACTGAGGCTGGAGGATGAGAACGAGAAGCAGGAGCGGCTCCTCATGAGCCTCCTGCCCCGGAACGTTGCCATGGAGATGAAGGAGGACTTCCTGAAGCCCCCTGAGAGGATTTTCCACAAGATTTACATCCAGAGGCACGACAATGTGAGCATCCTGTTTGCTGACATCGTGGGTTTCACGGGCTTGGCATCCCAGTGCACAGCCCAGGAGCTGGTGAAACTCCTCAATGAGCTCTTCGGCAAGTTCGATGAATTAGCCACGGAGAACCACTGTCGCCGCATCAAGATTCTCGGGGACTGCTACTACTGCGTGTCGGGCCTCACCCAGCCCAAGACTGACCATGCCCACTGCTGTGTGGAGATGGGACTCGACATGATTGATACCATCACATCTGTGGCTGAAGCCACCGAGGTGGATCTGAACATGCGTGTGGGTCTGCACACGGGCAGGGTCCTCTGTGGTGTCCTGGGCTTGCGCAAGTGGCAGTACGACGTGTGGTCCAATGATGTGACCTTGGCCAATGTCATGGAAGCCGCTGGCCTGCCAGGGAAGGTTCATATCACAAAGACGACCCTAGCGTGCTTGAATGGGGACTACGAGGTAGAACCGGGTTACGGACATGAGAGGAACAGTTTCTTGAAAACTCATAACATCGAAACCTTTTTTATTGTGCCATCCCATCGCCGAAAGATATTTCCAGGCCTGATTCTCTCAGATATAAAACCGGCCAAAAGGATGAAGTTCAAGACTGTCTGCTACCTGCTGGTGCAGCTCATGCACTGCCGGAAAATGTTCAAGGCCGAGATCCCCTTCTCCAATGTCATGACCTGCGAGGACGATGACAAGCGGAGGGCATTAAGAACAGCCTCGGAAAAACTCAGAAACCGCTCATCTTTTTCTACCAACGTTGTCTACACCACCCCGGGCACTCGCGTCAACAGGTACATCAGCCGCCTCTTAGAAGCCCGCCAGACAGAGCTGGAGATGGCAGACCTGAACTTCTTTACCCTGAAGTACAAACATGTCGAACGGGAGCAAAAGTACCACCAGCTTCAGGACGAGTATTTCACCAGCGCCGTTGTCCTCACCCTCATCCTGGCTGCCTTATTTGGCCTTGTCTACCTTCTAATATTCCCACAGAGTGTGGTCGTCCTGCTCCTGCTAGTATTCTGCATCTGCTTCCTGGTGGCCTGTGTCCTGTACCTGCACATCACCCGGGTCCAGTGTTTTCCAGGGTGCCTGACGATTCAGATTCGCACTGTCCTGTGTATTTTCATAGTGGTCTTAATCTACTCAGTAGCCCAAGGTTGTGTGGTGGGCTGCCTGCCTTGGGCCTGGAGCTCCAAGCCCAACAGTTCCCTGGTGGTCCTTTCGTCTGGGGGCCAGCGCACAGCCCTGCCCACCCTGCCCTGCGAGTCTACACACCATGCCCTGCTCTGCTGCCTGGTGGGCACCCTCCCGCTAGCCATATTTTTCCGGGTGTCCTCCTTGCCAAAAATGATCCTGCTCTCCGGGCTCACCACGTCCTACATCCTCGTTCTGGAGCTCAGCGGATACACCAGGACTGGGGGTGGTGCCGTCTCCGGGCGCAGCTACGAGCCGATTGTGGCCATCCTGCTCTTCTCCTGTGCGCTGGCCCTGCATGCCAGGCAGGTGGACATCAGGCTGAGGCTGGACTACCTCTGGGCCGCACAGGCAGAGGAGGAGCGAGAGGACATGGAGAAGGTGAAGCTGGACAACAGGCGCATCCTCTTCAACCTCCTGCCGGCCCACGTCGCCCAGCACTTCCTCATGTCCAACCCTCGGAACATGGACCTCTACTACCAGTCCTACTCCCAGGTGGGCGTCATGTTTGCCTCCATCCCCAACTTCAATGACTTCTACATCGAGCTGGACGGCAACAACATGGGGGTGGAGTGTCTGCGGCTTCTCAACGAGATCATCGCCGACTTTGACGAGCTCATGGAAAAAGACTTTTACAAGGACATAGAGAAGATCAAGACCATCGGGAGCACCTACATGGCCGCTGTGGGGCTAGCGCCCACCTCGGGGACCAAGGCTAAGAAGTCCATCTCCTCCCACCTGAGCACGCTGGCGGACTTTGCCATTGAGATGTTTGACGTTCTGGATGAAATCAACTACCAGTCTTACAACGACTTTGTCCTCCGAGTTGGCATCAATGTTGGCCCTGTGGTGGCTGGAGTGATTGGCGCTCGCAGGCCCCAGTACGACATCTGGGGAAACACAGTCAACGTGGCCAGTCGGATGGATAGCACAGGGGTCCAGGGCAGAATCCAGGTGACTGAGGAAGTCCACCGGCTGCTGAGAAGGTGCCCCTACCACTTTGTGTGCCGAGGCAAAGTCAGTGTCAAGGGCAAAGGCGAGATGTTGACATACTTTCTAGAAGGCAGGACTGATGGAAACGGCTCCCAAATCAGGTCCCTGGGCTTGGATCGGAAAATGTGTCCATTTGGGAGAGCTGGCCTTCAGGGCAGACGTCCCCCCGTGTGCCCCATGCCTGGCGTCTCAGTCAGGGCTGGGCTCCCTCCACACTCCCCAGGCCAGTACCTGCCCTCTGCAGCAGCTGGGAAGGAGGCTTGA

SmBiT-linker-AC2

ATGGTCACCGGCTATCGGCTGTTTGAAGAGATTCTGGGGGGGTCAGGAGGAGGAGGCTCAGGGGGCTCATCATCAGGAGGATGGCAGGAGGCGATGCGGCGCCGCCGCTACCTGCGGGACCGCTCCGAGGAGGCGGCGGGCGGCGGAGACGGGCTGCCGCGGTCCCGGGACTGGCTCTACGAGTCCTACTACTGCATGAGCCAGCAGCACCCGCTCATCGTCTTCCTGCTGCTCATCGTCATGGGCTCCTGCCTCGCCCTGCTCGCCGTCTTCTTCGCGCTCGGGCTGGAAGTTGAAGACCATGTGGCGTTTCTAATAACAGTTCCAACTGCCCTGGCGATTTTCTTTGCGATATTTATCCTGGTCTGCATCGAGTCTGTGTTTAAGAAGCTGCTGCGCCTCTTCTCGTTGGTGATATGGATATGCCTTGTTGCCATGGGATACCTGTTCATGTGTTTTGGAGGCACCGTCTCTCCCTGGGACCAGGTATCGTTCTTCCTCTTCATCATCTTCGTGGTGTACACCATGCTGCCCTTCAACATGCGAGACGCCATCATTGCCAGCGTCCTCACCTCCTCCTCCCACACCATCGTGCTTAGCGTCTGCCTGTCTGCAACACCGGGAGGCAAGGAGCACCTGGTCTGGCAGATCCTGGCCAATGTGATCATTTTCATCTGTGGGAACCTGGCGGGAGCCTACCATAAGCACCTCATGGAACTCGCTCTTCAGCAAACATATCAGGACACCTGTAATTGCATCAAGTCGCGGATCAAGTTGGAATTTGAAAAACGTCAACAGGAGCGGCTTCTGCTCTCCCTGCTGCCGGCCCACATCGCCATGGAGATGAAAGCGGAGATCATCCAGAGGCTGCAGGGCCCCAAGGCGGGCCAGATGGAGAACACAAATAACTTCCACAACCTGTATGTGAAGCGGCATACAAACGTGAGCATCTTATACGCTGACATCGTTGGCTTTACCCGGCTGGCAAGTGACTGCTCCCCGGGAGAACTAGTCCACATGCTGAATGAGCTCTTTGGAAAGTTTGATCAAATTGCAAAGGAGAATGAATGCATGAGAATTAAAATTTTAGGAGACTGCTACTACTGTGTATCTGGACTCCCTATATCTCTCCCTAACCATGCCAAGAACTGTGTGAAAATGGGGCTGGACATGTGTGAAGCCATAAAGAAAGTGAGGGATGCTACTGGAGTTGATATCAACATGCGCGTGGGCGTGCATTCTGGGAATGTCCTGTGTGGCGTGATTGGTCTGCAGAAGTGGCAATATGATGTGTGGTCACATGATGTGACCTTGGCCAACCACATGGAAGCTGGAGGGGTCCCTGGACGTGTTCACATTTCTTCTGTCACCCTGGAGCACTTGAATGGCGCTTATAAAGTGGAGGAGGGAGATGGTGACATTAGGGACCCATATTTAAAACAGCACCTGGTGAAAACCTACTTTGTGATCAACCCCAAGGGAGAACGACGGAGCCCCCAGCATCTCTTCAGACCTCGCCACACCCTTGATGGAGCCAAAATGAGGGCCTCGGTCCGCATGACCCGGTACTTGGAGTCCTGGGGGGCAGCCAAGCCCTTTGCACACCTACATCACAGGGACAGCATGACCACAGAGAACGGCAAGATCAGCACCACGGATGTACCCATGGGTCAGCATAATTTTCAAAATCGCACCTTAAGAACCAAGTCACAAAAGAAGAGATTTGAAGAAGAATTGAATGAAAGGATGATTCAAGCAATTGATGGGATTAATGCACAGAAGCAATGGCTCAAGTCTGAAGACATTCAGAGAATCTCACTGCTTTTCTATAACAAAGTACTAGAAAAAGAGTACCGGGCCACGGCACTGCCAGCGTTCAAGTATTATGTGACTTGTGCCTGTCTCATATTCTTCTGCATCTTCATTGTGCAGATTCTCGTGCTGCCAAAAACGTCCGTCCTGGGCATCTCCTTTGGGGCTGCGTTTCTCTTGCTGGCCTTCATCCTCTTCGTCTGCTTTGCTGGACAGCTTCTGCAATGCAGCAAAAAAGCCTCTCCCCTGCTCATGTGGCTTTTGAAGTCCTCGGGCATCATTGCCAACCGCCCCTGGCCACGGATCTCTCTCACGATCATCACCACAGCCATCATATTAATGATGGCCGTGTTCAACATGTTTTTCCTGAGTGACTCAGAGGAAACAATCCCTCCAACTGCCAACACAACAAACACAAGCTTTTCAGCCTCAAATAATCAGGTGGCGATTCTGCGTGCGCAGAATTTATTTTTCCTCCCGTACTTTATCTACAGCTGCATTCTGGGACTGATATCCTGTTCCGTGTTCCTGCGGGTAAACTATGAGCTGAAGATGTTGATCATGATGGTGGCCTTGGTGGGCTACAACACCATCCTACTCCACACCCACGCCCACGTCCTGGGCGACTACAGCCAGGTCTTATTTGAGAGACCAGGCATTTGGAAAGACCTGAAGACCATGGGCTCTGTGTCTCTCTCTATATTCTTCATCACACTGCTTGTTCTGGGTAGACAGAATGAATATTACTGTAGGTTAGACTTCTTATGGAAGAACAAATTCAAAAAAGAGCGGGAGGAGATAGAGACCATGGAGAACCTGAACCGCGTGCTGCTGGAGAACGTGCTTCCCGCGCACGTGGCTGAGCACTTCCTGGCCAGGAGCCTGAAGAATGAGGAGCTATACCACCAGTCCTATGACTGCGTCTGCGTCATGTTTGCCTCCATTCCGGATTTCAAAGAATTTTATACAGAATCCGACGTGAACAAGGAGGGCTTGGAATGCCTTCGGCTCCTGAACGAGATCATCGCTGACTTTGATGATCTTCTTTCCAAGCCAAAATTCAGTGGAGTTGAAAAGATTAAGACCATTGGCAGCACATACATGGCAGCAACAGGTCTGAGCGCTGTGCCCAGCCAGGAGCACTCCCAGGAGCCCGAGCGGCAGTACATGCACATTGGCACCATGGTGGAGTTTGCTTTTGCCCTGGTAGGGAAGCTGGATGCCATCAACAAGCACTCCTTCAACGACTTCAAATTGCGAGTGGGTATTAACCATGGACCTGTGATAGCTGGTGTGATTGGAGCTCAGAAGCCACAATATGATATCTGGGGCAACACTGTCAATGTGGCCAGTAGGATGGACAGCACCGGAGTCCTGGACAAAATACAGGTTACCGAGGAGACGAGCCTCGTCCTGCAGACCCTCGGATACACGTGCACCTGTCGAGGAATAATCAACGTGAAAGGAAAGGGGGACCTGAAGACGTACTTTGTAAACACAGAAATGTCAAGGTCCCTTTCCCAGAGCAACGTGGCATCCTGA

SmBiT-linker-AC3

ATGGTCACCGGCTATCGGCTGTTTGAAGAGATTCTGGGGGGGTCAGGAGGAGGAGGCTCAGGGGGCTCATCATCAGGAGGACCGAGGAACCAGGGCTTCTCCGAGCCCGAATACTCGGCCGAGTACTCAGCCGAGTACTCCGTCAGCCTGCCCTCCGACCCTGACCGCGGGGTGGGCCGGACCCATGAAATCTCGGTCCGGAACTCGGGCTCCTGCCTGTGCCTGCCTCGCTTCATGCGGCTGACTTTCGTGCCGGAGTCCTTGGAGAACCTCTACCAGACCTACTTCAAAAGGCAGCGCCACGAGACCCTGCTGGTGCTGGTGGTCTTTGCAGCCCTCTTTGACTGCTACGTGGTGGTCATGTGTGCTGTGGTCTTCTCCAGCGACAAGCTGGCTTCCCTCGCCGTGGCTGGAATTGGACTGGTGTTGGACATCATCCTCTTCGTGCTCTGCAAAAAGGGGCTGCTCCCGGACCGGGTCACCCGCAGAGTGCTGCCCTACGTGCTGTGGCTGCTCATAACCGCCCAGATCTTCTCCTACCTGGGCCTGAACTTCGCGCGTGCCCACGCGGCTAGTGACACGGTGGGCTGGCAGGTCTTCTTTGTCTTCTCCTTCTTCATCACGCTGCCCCTCAGCCTCAGCCCCATCGTGATCATCTCCGTGGTCTCCTGTGTGGTGCACACGTTGGTCCTGGGGGTCACCGTGGCCCAGCAGCAGCAGGAGGAGCTCAAGGGGATGCAGCTGCTGCGGGAGATCCTGGCCAACGTCTTCCTCTACCTGTGCGCCATCGCTGTGGGCATCATGTCCTACTACATGGCTGACCGCAAGCACCGCAAGGCCTTCCTGGAGGCCCGCCAGTCGCTGGAGGTGAAGATGAACCTGGAAGAGCAGAGCCAGCAGCAGGAGAACCTCATGCTTTCCATCCTGCCCAAGCACGTGGCTGACGAGATGCTGAAAGACATGAAGAAAGACGAGAGCCAGAAGGACCAGCAGCAGTTCAACACCATGTACATGTACCGTCACGAGAACGTCAGCATCCTCTTTGCCGACATCGTGGGCTTTACCCAGCTGTCTTCTGCCTGCAGTGCCCAGGAGCTTGTGAAGCTGCTCAACGAGCTCTTTGCCCGCTTTGACAAGCTGGCAGCTAAATACCACCAGCTGCGGATTAAGATCCTGGGCGACTGCTACTACTGCATCTGCGGCTTGCCCGACTACCGGGAGGACCACGCCGTCTGCTCCATCCTCATGGGGCTGGCCATGGTGGAGGCCATCTCGTATGTGCGGGAGAAGACCAAGACTGGGGTGGACATGCGTGTGGGGGTGCACACGGGCACCGTGCTGGGGGGCGTCCTGGGCCAGAAGCGCTGGCAGTACGACGTGTGGTCGACTGATGTCACTGTAGCCAACAAGATGGAGGCCGGCGGCATCCCTGGGCGCGTGCACATCTCCCAGAGCACCATGGACTGCCTGAAAGGGGAGTTTGATGTGGAGCCAGGCGATGGGGGCAGCCGCTGTGATTACCTAGAAGAGAAGGGTATTGAAACCTACCTCATCATTGCCTCCAAGCCAGAGGTGAAGAAAACAGCCACCCAGAATGGCCTCAATGGCTCGGCCCTGCCCAATGGAGCACCAGCTTCCTCAAAGTCCAGCTCCCCTGCCCTCATTGAGACCAAGGAGCCCAACGGGAGTGCCCACAGCAGTGGGTCCACGTCGGAGAAGCCCGAGGAGCAGGATGCCCAGGCCGACAACCCCTCATTCCCCAACCCACGCCGGAGGCTGCGCCTGCAGGACCTGGCTGACCGAGTGGTGGATGCCTCTGAAGATGAGCACGAGCTCAACCAGCTGCTCAACGAGGCCCTGCTTGAGCGAGAGTCCGCCCAAGTAGTAAAGAAGAGAAACACCTTCCTCTTGTCCATGCGGTTCATGGACCCCGAGATGGAAACCCGCTACTCGGTGGAGAAGGAGAAGCAGAGTGGGGCTGCCTTCAGCTGCTCCTGCGTCGTCCTGCTCTGCACGGCCCTGGTCGAGATACTCATCGACCCCTGGCTAATGACAAACTATGTGACCTTCATGGTGGGGGAGATTCTGCTCCTCATCCTGACCATCTGCTCCCTGGCTGCCATCTTTCCCCGGGCCTTTCCTAAGAAGCTTGTGGCCTTCTCAACTTGGATTGACCGGACCCGCTGGGCCAGGAACACCTGGGCCATGCTCGCCATCTTCATCCTGGTGATGGCAAATGTCGTGGACATGCTCAGCTGTCTCCAGTACTACACGGGACCCAGCAATGCAACGGCAGGGATGGAAACGGAGGGCAGCTGCCTGGAGAACCCCAAGTATTACAACTATGTGGCCGTGCTGTCCCTCATCGCCACCATCATGCTGGTGCAGGTCAGCCACATGGTGAAGCTCACGCTCATGCTGCTCGTCGCAGGCGCCGTGGCCACCATCAACCTCTATGCCTGGCGTCCCGTCTTTGATGAATACGACCACAAGCGTTTTCGGGAGCACGACTTACCTATGGTGGCCTTAGAGCAGATGCAAGGATTCAACCCTGGGCTCAATGGCACTGACAGGCTGCCCCTGGTGCCTTCCAAGTACTCTATGACGGTGATGGTGTTCCTCATGATGCTCAGCTTCTACTACTTCTCCCGCCACGTAGAAAAACTGGCACGGACACTTTTCTTGTGGAAGATTGAGGTCCACGACCAGAAGGAACGTGTCTATGAGATGCGACGCTGGAACGAGGCCTTGGTCACCAACATGTTGCCTGAGCACGTGGCACGCCATTTCCTGGGGTCCAAGAAGAGAGATGAGGAGCTGTATAGCCAGACGTATGATGAGATTGGAGTCATGTTTGCCTCCCTGCCCAACTTTGCTGACTTCTACACAGAGGAGAGCATCAACAATGGTGGTATTGAGTGTCTGCGTTTCCTCAATGAAATCATCTCAGATTTTGACTCTCTCCTGGACAATCCCAAGTTCCGGGTGATCACCAAGATCAAAACCATTGGCAGCACGTATATGGCGGCTTCAGGAGTCACCCCCGATGTCAACACCAATGGCTTTGCCAGCTCCAACAAGGAAGACAAGTCCGAGAGAGAGCGCTGGCAGCACCTGGCTGACCTGGCCGACTTCGCGCTGGCCATGAAGGATACGCTCACCAACATCAACAACCAGTCCTTCAATAACTTCATGCTGCGCATAGGCATGAACAAAGGCGGGGTTCTGGCTGGGGTCATCGGAGCCCGGAAACCACACTACGACATCTGGGGCAATACAGTCAATGTAGCCAGCAGGATGGAGTCCACGGGGGTCATGGGCAACATTCAGGTGGTAGAAGAAACCCAAGTCATCCTCCGAGAGTACGGCTTCCGCTTTGTGAGGCGAGGCCCCATCTTTGTGAAGGGGAAGGGGGAGCTGCTGACCTTCTTCTTGAAGGGGCGGGATAAGCTAGCCACCTTCCCCAATGGCCCCTCTGTCACACTGCCCCACCAGGTGGTGGACAACTCCTGA

SmBiT-linker-AC4

ATGGTCACCGGCTATCGGCTGTTTGAAGAGATTCTGGGGGGGTCAGGAGGAGGAGGCTCAGGGGGCTCATCATCAGGAGGAGCCCGCCTCTTCAGCCCCCGGCCGCCCCCCAGCGAAGACCTCTTCTACGAGACCTACTACAGCCTGAGCCAGCAGTACCCGCTGCTGCTGCTGCTGCTGGGGATCGTGCTCTGTGCGCTCGCGGCGCTGCTCGCAGTGGCCTGGGCCAGCGGCAGGGAGCTGACCTCAGACCCGAGCTTCCTGACCACTGTGCTGTGCGCGCTGGGCGGCTTCTCGCTGCTGCTGGGCCTCGCTTCCCGGGAGCAGCGACTGCAGCGCTGGACGCGTCCCCTGTCCGGCTTGGTATGGGTCGCGCTGCTAGCGCTAGGCCACGCCTTCCTGTTCACCGGGGGCGTGGTGAGCGCCTGGGACCAGGTGTCCTATTTTCTCTTCGTCATCTTCACGGCGTATGCCATGCTGCCCTTGGGCATGCGGGACGCCGCCGTCGCGGGCCTCGCCTCCTCACTCTCGCATCTGCTGGTCCTCGGGCTGTATCTTGGGCCACAGCCGGACTCACGGCCTGCACTGCTGCCGCAGTTGGCAGCAAACGCAGTGCTGTTCCTGTGCGGGAACGTGGCAGGAGTGTACCACAAGGCGCTGATGGAGCGCGCCCTGCGGGCCACGTTCCGGGAGGCACTCAGCTCCCTGCACTCACGCCGGCGGCTGGACACCGAGAAGAAGCACCAGGAACACCTTCTCTTGTCCATCCTTCCTGCCTACCTGGCCCGAGAGATGAAGGCAGAGATCATGGCACGGCTGCAGGCAGGACAGGGGTCACGGCCAGAGAGCACTAACAATTTCCACAGCCTCTATGTCAAGAGGCACCAGGGAGTCAGCGTGCTGTATGCTGACATCGTGGGCTTCACGCGGCTGGCCAGCGAGTGTTCCCCTAAGGAGCTGGTGCTCATGCTCAATGAGCTCTTTGGCAAGTTCGACCAGATTGCCAAGGAGCATGAATGCATGCGGATCAAGATCCTGGGGGACTGTTACTACTGTGTCTCTGGGCTGCCACTCTCACTGCCAGACCATGCCATCAACTGCGTGCGCATGGGCCTGGACATGTGCCGGGCCATCAGGAAACTGCGGGCAGCCACTGGCGTGGACATCAACATGCGTGTGGGCGTGCACTCAGGCAGCGTACTGTGTGGAGTCATCGGGCTGCAGAAGTGGCAGTACGACGTTTGGTCACATGATGTCACACTGGCTAACCACATGGAGGCAGGCGGTGTACCAGGGCGAGTGCACATCACAGGGGCTACCCTGGCCCTGCTGGCAGGGGCTTATGCTGTGGAGGACGCAGGCATGGAGCATCGGGACCCCTACCTTCGGGAGCTAGGGGAGCCTACCTATCTGGTCATCGATCCACGGGCAGAGGAGGAGGATGAGAAGGGCACTGCAGGAGGCTTGCTGTCCTCGCTTGAGGGCCTCAAGATGCGTCCATCACTGCTGATGACCCGTTACCTGGAGTCCTGGGGCGCAGCCAAGCCTTTTGCCCACCTGAGCCACGGAGACAGCCCTGTGTCCACCTCCACCCCTCTCCCGGAGAAGACCCTGGCTTCCTTCAGCACCCAGTGGAGCCTGGATCGGAGCCGTACCCCCCGGGGACTAGATGATGAACTGGACACCGGGGATGCCAAGTTCTTCCAGGTCATTGAGCAGCTCAACTCGCAGAAACAGTGGAAGCAGTCGAAGGACTTCAACCCACTGACACTGTACTTCAGAGAGAAGGAGATGGAGAAAGAGTACCGACTCTCTGCAATCCCCGCCTTCAAATACTATGAAGCCTGCACCTTCCTGGTTTTTCTCTCCAACTTCATCATCCAGATGCTAGTGACAAACAGGCCCCCAGCTCTGGCCATCACGTATAGCATCACCTTCCTCCTCTTCCTCCTCATCCTTTTTGTCTGCTTCTCAGAGGACCTGATGAGGTGTGTCCTGAAAGGCCCCAAGATGCTGCACTGGCTGCCTGCACTGTCTGGCCTGGTGGCCACACGACCAGGACTGAGAATAGCCTTGGGCACCGCCACCATCCTCCTTGTCTTTGCCATGGCCATTACCAGCCTGTTCTTCTTCCCAACATCATCAGACTGCCCTTTCCAAGCTCCCAATGTGTCCTCCATGATTTCCAACCTCTCCTGGGAGCTCCCTGGGTCTCTGCCTCTCATCAGTGTCCCATACTCCATGCACTGCTGCACGCTGGGCTTCCTCTCCTGCTCCCTCTTTCTGCACATGAGCTTCGAGCTGAAGCTGCTGCTGCTCCTGCTGTGGCTGGCGGCATCCTGCTCCCTCTTCCTGCACTCCCATGCCTGGCTGTCGGAATGCCTCATCGTCCGCCTCTATCTGGGCCCCTTGGACTCCAGGCCCGGAGTGCTGAAGGAGCCCAAACTGATGGGTGCTATCTCCTTCTTCATCTTCTTCTTCACCCTCCTTGTCCTGGCTCGCCAGAATGAGTACTACTGCCGCCTGGACTTCCTGTGGAAGAAGAAGCTGAGGCAGGAGAGGGAGGAGACAGAGACGATGGAGAACCTGACTCGGCTGCTCTTGGAGAACGTGCTCCCTGCACACGTGGCCCCCCAGTTCATTGGCCAGAACCGGCGCAACGAGGATCTCTACCACCAGTCCTATGAATGCGTTTGTGTCCTCTTCGCCTCAGTCCCAGACTTCAAGGAGTTCTACTCTGAATCCAACATCAATCATGAGGGCCTAGAGTGTCTGAGGCTGCTCAATGAGATAATTGCTGATTTTGATGAGCTGCTCTCCAAGCCCAAGTTCAGTGGGGTGGAGAAGATCAAGACCATCGGCAGCACCTACATGGCAGCCACAGGCTTAAATGCCACCTCTGGACAGGATGCACAACAGGATGCTGAACGGAGCTGCAGCCACCTTGGCACTATGGTGGAATTTGCCGTGGCCCTGGGGTCTAAGCTGGACGTCATCAACAAGCATTCATTCAACAACTTCCGCCTGCGAGTGGGGTTGAACCATGGACCCGTAGTAGCTGGAGTTATTGGGGCCCAGAAGCCGCAATATGACATTTGGGGCAACACAGTGAACGTGGCCAGCCGCATGGAGAGTACAGGAGTCCTTGGCAAAATCCAAGTGACTGAGGAGACAGCATGGGCCCTACAGTCCCTGGGCTACACCTGCTACAGCCGGGGTGTCATCAAGGTGAAAGGCAAAGGGCAGCTCTGCACCTACTTCCTGAACACAGACTTGACACGAACTGGACCTCCTTCAGCTACCCTAGGCTGA

SmBiT-linker-AC5

ATGGTCACCGGCTATCGGCTGTTTGAAGAGATTCTGGGGGGGTCAGGAGGAGGAGGCTCAGGGGGCTCATCATCAGGAGGATCCGGCTCCAAAAGCGTGAGCCCCCCGGGCTACGCGGCGCAGAAGACTGCGGCGCCGGCGCCCCGGGGAGGCCCCGAACACCGCTCTGCGTGGGGCGAGGCCGATTCCCGCGCGAATGGCTACCCCCATGCCCCCGGGGGCTCTGCCCGCGGCTCCACCAAGAAACCCGGGGGGGCGGTGACCCCGCAGCAGCAGCAGCGCCTGGCCAGCCGCTGGCGCAGCGACGACGACGACGATCCTCCGCTGAGCGGTGACGACCCCCTGGCCGGGGGCTTCGGCTTCAGCTTCCGCTCCAAGTCCGCCTGGCAGGAGCGCGGCGGCGACGACTGCGGTCGCGGCAGCCGCCGGCAGCGGCGGGGCGCGGCCAGCGGGGGCAGCACCCGGGCGCCCCCTGCGGGCGGCGGCGGCGGCTCGGCGGCGGCGGCTGCCTCGGCGGGCGGGACGGAGGTGCGCCCTCGCTCGGTGGAGGTGGGTCTGGAGGAGCGGCGGGGCAAGGGGCGCGCGGCCGACGAGCTGGAGGCCGGCGCCGTCGAGGGCGGCGAGGGGTCCGGGGATGGCGGCAGCTCGGCGGACTCGGGCTCGGGCGCGGGGCCCGGCGCGGTGCTGTCCCTGGGCGCCTGCTGCCTGGCGTTGCTGCAGATATTCCGCTCCAAGAAGTTCCCGTCGGACAAACTGGAGCGGCTGTACCAGCGCTACTTCTTCCGCCTGAACCAGAGCAGCCTCACCATGCTCATGGCCGTGCTGGTGCTCGTGTGCCTGGTCATGTTGGCCTTCCACGCGGCGCGGCCCCCGCTCCAGCTGCCCTACCTGGCCGTGCTGGCGGCCGCCGTCGGCGTGATCCTCATCATGGCTGTGCTTTGCAACCGCGCCGCCTTCCACCAGGACCACATGGGCCTGGCCTGCTATGCGCTCATCGCCGTGGTGCTGGCCGTCCAGGTGGTGGGCCTGCTGCTGCCGCAGCCACGCAGCGCCTCTGAGGGCATCTGGTGGACCGTGTTCTTCATCTACACCATCTACACGCTGCTGCCCGTGCGCATGCGGGCCGCAGTGCTCAGCGGGGTGCTCCTGTCCGCCCTCCACCTGGCCATCGCCCTGCGCACCAACGCCCAGGACCAGTTCCTGCTGAAGCAGCTTGTCTCCAATGTTCTCATTTTCTCCTGCACCAACATCGTGGGTGTCTGCACCCACTATCCGGCTGAGGTCTCCCAGAGACAGGCTTTCCAGGAGACCCGAGAGTGCATCCAGGCGCGGCTCCACTCGCAGCGGGAGAACCAGCAGCAGGAACGGCTCCTGCTGTCTGTCCTTCCCCGTCATGTTGCCATGGAGATGAAAGCAGACATCAACGCCAAGCAGGAGGATATGATGTTCCATAAGATTTACATCCAGAAACATGACAACGTGAGCATCCTGTTTGCTGACATCGAGGGCTTCACCAGCCTGGCGTCCCAGTGCACTGCACAGGAACTGGTCATGACCCTCAACGAGCTCTTCGCCCGCTTTGACAAGCTGGCCGCAGAGAATCACTGTTTACGTATTAAGATCCTTGGGGATTGTTATTACTGCGTCTCGGGGCTGCCTGAAGCAAGGGCTGACCACGCCCACTGCTGTGTGGAGATGGGCATGGACATGATCGAGGCCATCTCGTTGGTCCGGGAGGTGACAGGGGTGAACGTGAACATGCGTGTGGGAATTCACAGCGGGCGAGTACACTGCGGTGTCCTTGGTCTCAGGAAGTGGCAGTTCGACGTCTGGTCTAACGATGTCACGCTAGCCAACCACATGGAGGCTGGCGGCAAGGCAGGACGCATCCACATCACCAAGGCTACACTCAACTACCTGAATGGGGACTACGAGGTGGAGCCAGGCTGTGGGGGCGAGCGCAACGCCTACCTCAAGGAGCACAGTATCGAGACCTTCCTCATCCTGCGCTGCACCCAGAAGCGGAAAGAAGAGAAGGCCATGATCGCCAAGATGAACCGCCAGAGAACCAACTCCATCGGGCACAACCCACCACACTGGGGGGCTGAGCGCCCCTTCTACAACCACCTGGGTGGCAACCAGGTGTCCAAGGAGATGAAGCGGATGGGCTTTGAAGACCCCAAGGACAAGAACGCCCAGGAGAGTGCGAACCCTGAGGATGAAGTGGATGAGTTTCTGGGCCGTGCCATTGACGCCAGGAGCATTGATAGGCTTCGGTCTGAGCACGTCCGCAAGTTCCTCCTGACCTTCAGGGAGCCTGACTTAGAGAAGAAGTACTCCAAGCAGGTAGACGACCGATTTGGTGCCTATGTGGCGTGTGCCTCGCTCGTCTTCCTCTTCATCTGCTTTGTCCAGATCACCATCGTGCCCCACTCCATATTCATGCTCAGCTTCTACCTGACCTGTTCCCTGCTGCTGACCTTGGTGGTGTTTGTGTCTGTGATCTACTCCTGCGTAAAGCTCTTCCCCTCCCCACTGCAGACCCTCTCCAGGAAGATCGTGCGGTCCAAGATGAACAGCACCCTGGTTGGGGTGTTCACCATCACCCTGGTGTTCCTGGCGGCTTTTGTCAACATGTTCACGTGCAACTCCAGGGACCTGCTGGGCTGCTTGGCACAGGAGCACAACATCAGCGCGAGCCAGGTCAACGCGTGTCACGTGGCGGAGTCGGCCGTCAACTACAGCCTGGGCGATGAGCAGGGCTTCTGTGGCAGCCCCTGGCCCAACTGCAACTTCCCCGAGTACTTCACCTACAGCGTGCTGCTCAGCCTGCTGGCCTGCTCCGTGTTCCTGCAGATCAGCTGCATCGGGAAGCTGGTGCTCATGCTGGCCATCGAGCTCATCTACGTGCTCATCGTGGAGGTGCCAGGTGTCACGCTCTTCGACAACGCCGACCTGCTGGTCACCGCCAACGCCATAGACTTCTTCAACAACGGGACCTCCCAGTGCCCTGAGCATGCAACCAAGGTGGCATTGAAGGTGGTGACGCCCATCATCATCTCAGTCTTTGTGCTGGCCCTGTACCTGCACGCCCAGCAGGTGGAGTCCACTGCCCGCCTCGACTTCCTCTGGAAACTGCAGGCCACAGAGGAGAAAGAGGAGATGGAGGAGCTGCAGGCCTACAACCGGCGGCTGCTGCACAACATCCTGCCCAAGGACGTGGCCGCTCACTTCCTGGCCCGCGAGCGGCGCAATGATGAGCTCTACTATCAGTCCTGTGAGTGTGTGGCGGTCATGTTCGCCTCCATCGCCAACTTCTCCGAGTTCTACGTTGAGCTGGAGGCCAACAACGAGGGTGTCGAGTGCCTGCGGCTACTCAATGAGATCATCGCTGACTTTGATGAGATCATCAGCGAGGATCGGTTCCGGCAGCTGGAGAAGATCAAGACCATCGGCAGCACCTACATGGCTGCCTCCGGCCTCAACGACTCTACCTACGACAAGGTGGGCAAGACCCACATCAAGGCACTGGCCGACTTTGCCATGAAGCTGATGGACCAGATGAAGTACATCAATGAGCACTCCTTCAACAACTTCCAGATGAAGATCGGGCTCAACATCGGCCCCGTGGTGGCCGGGGTGATAGGGGCACGAAAGCCTCAGTACGACATCTGGGGCAATACCGTGAACGTGGCCAGCCGCATGGACAGCACCGGTGTACCCGACCGCATCCAGGTCACCACAGACATGTACCAGGTGCTGGCTGCCAACACGTACCAGCTGGAGTGCCGGGGCGTGGTCAAGGTCAAGGGCAAAGGCGAGATGATGACCTACTTCCTCAATGGAGGGCCCCCGCTCAGTTGA

SmBiT-linker-AC6

ATGGTCACCGGCTATCGGCTGTTTGAAGAGATTCTGGGGGGGTCAGGAGGAGGAGGCTCAGGGGGCTCATCATCAGGAGGATCATGGTTTAGTGGCCTCCTGGTCCCTAAAGTGGATGAACGGAAAACAGCCTGGGGTGAACGCAATGGGCAGAAGCGTTCGCGGCGCCGTGGCACTCGGGCAGGTGGCTTCTGCACGCCCCGCTATATGAGCTGCCTCCGGGATGCAGAGCCACCCAGCCCCACCCCTGCGGGCCCCCCTCGGTGCCCCTGGCAGGATGACGCCTTCATCCGGAGGGGCGGCCCAGGCAAGGGCAAGGAGCTGGGGCTGCGGGCAGTGGCCCTGGGCTTCGAGGATACCGAGGTGACAACGACAGCGGGCGGGACGGCTGAGGTGGCGCCCGACGCGGTGCCCAGGAGTGGGCGATCCTGCTGGCGCCGTCTGGTGCAGGTGTTCCAGTCGAAGCAGTTCCGTTCGGCCAAGCTGGAGCGCCTGTACCAGCGGTACTTCTTCCAGATGAACCAGAGCAGCCTGACGCTGCTGATGGCGGTGCTGGTGCTGCTCACAGCGGTGCTGCTGGCTTTCCACGCCGCACCCGCCCGCCCTCAGCCTGCCTATGTGGCACTGTTGGCCTGTGCCGCCGCCCTGTTCGTGGGGCTCATGGTGGTGTGTAACCGGCATAGCTTCCGCCAGGACTCCATGTGGGTGGTGAGCTACGTGGTGCTGGGCATCCTGGCGGCAGTGCAGGTCGGGGGCGCTCTCGCAGCAGACCCGCGCAGCCCCTCTGCGGGCCTCTGGTGCCCTGTGTTCTTTGTCTACATCGCCTACACGCTCCTCCCCATCCGCATGCGGGCTGCCGTCCTCAGCGGCCTGGGCCTCTCCACCTTGCATTTGATCTTGGCCTGGCAACTTAACCGTGGTGATGCCTTCCTCTGGAAGCAGCTCGGTGCCAATGTGCTGCTGTTCCTCTGCACCAACGTCATTGGCATCTGCACACACTATCCAGCAGAGGTGTCTCAGCGCCAGGCCTTTCAGGAGACCCGCGGTTACATCCAGGCCCGGCTCCACCTGCAGCATGAGAATCGGCAGCAGGAGCGGCTGCTGCTGTCGGTATTGCCCCAGCACGTTGCCATGGAGATGAAAGAAGACATCAACACAAAAAAAGAAGACATGATGTTCCACAAGATCTACATACAGAAGCATGACAATGTCAGCATCCTGTTTGCAGACATTGAGGGCTTCACCAGCCTGGCATCCCAGTGCACTGCGCAGGAGCTGGTCATGACCCTGAATGAGCTCTTTGCCCGGTTTGACAAGCTGGCTGCGGAGAATCACTGCCTGAGGATCAAGATCTTGGGGGACTGTTACTACTGTGTGTCAGGGCTGCCGGAGGCCCGGGCCGACCATGCCCACTGCTGTGTGGAGATGGGGGTAGACATGATTGAGGCCATCTCGCTGGTACGTGAGGTGACAGGTGTGAATGTGAACATGCGCGTGGGCATCCACAGCGGGCGCGTGCACTGCGGCGTCCTTGGCTTGCGGAAATGGCAGTTCGATGTGTGGTCCAATGATGTGACCCTGGCCAACCACATGGAGGCAGGAGGCCGGGCTGGCCGCATCCACATCACTCGGGCAACACTGCAGTACCTGAACGGGGACTACGAGGTGGAGCCAGGCCGTGGTGGCGAGCGCAACGCGTACCTCAAGGAGCAGCACATTGAGACTTTCCTCATCCTGGGCGCCAGCCAGAAACGGAAAGAGGAGAAGGCCATGCTGGCCAAGCTGCAGCGGACTCGGGCCAACTCCATGGAAGGGCTGATGCCGCGCTGGGTTCCTGATCGTGCCTTCTCCCGGACCAAGGACTCCAAGGCCTTCCGCCAGATGGGCATTGATGATTCCAGCAAAGACAACCGGGGCACCCAAGATGCCCTGAACCCTGAGGATGAGGTGGATGAGTTCCTGAGCCGTGCCATCGATGCCCGCAGCATTGATCAGCTGCGGAAGGACCATGTGCGCCGGTTTCTGCTCACCTTCCAGAGAGAGGATCTTGAGAAGAAGTACTCCCGGAAGGTGGATCCCCGCTTCGGAGCCTACGTTGCCTGTGCCCTGTTGGTCTTCTGCTTCATCTGCTTCATCCAGCTTCTCATCTTCCCACACTCCACCCTGATGCTTGGGATCTATGCCAGCATCTTCCTGCTGCTGCTAATCACCGTGCTGATCTGTGCTGTGTACTCCTGTGGTTCTCTGTTCCCTAAGGCCCTGCAACGTCTGTCCCGCAGCATTGTCCGCTCACGGGCACATAGCACCGCAGTTGGCATCTTTTCCGTCCTGCTTGTGTTTACTTCTGCCATTGCCAACATGTTCACCTGTAACCACACCCCCATACGGAGCTGTGCAGCCCGGATGCTGAATTTAACACCTGCTGACATCACTGCCTGCCACCTGCAGCAGCTCAATTACTCTCTGGGCCTGGATGCTCCCCTGTGTGAGGGCACCATGCCCACCTGCAGCTTTCCTGAGTACTTCATCGGGAACATGCTGCTGAGTCTCTTGGCCAGCTCTGTCTTCCTGCACATCAGCAGCATCGGGAAGTTGGCCATGATCTTTGTCTTGGGGCTCATCTATTTGGTGCTGCTTCTGCTGGGTCCCCCAGCCACCATCTTTGACAACTATGACCTACTGCTTGGCGTCCATGGCTTGGCTTCTTCCAATGAGACCTTTGATGGGCTGGACTGTCCAGCTGCAGGGAGGGTGGCCCTCAAATATATGACCCCTGTGATTCTGCTGGTGTTTGCGCTGGCGCTGTATCTGCATGCTCAGCAGGTGGAGTCGACTGCCCGCCTAGACTTCCTCTGGAAACTACAGGCAACAGGGGAGAAGGAGGAGATGGAGGAGCTACAGGCATACAACCGGAGGCTGCTGCATAACATTCTGCCCAAGGACGTGGCGGCCCACTTCCTGGCCCGGGAGCGCCGCAATGATGAACTCTACTATCAGTCGTGTGAGTGTGTGGCTGTTATGTTTGCCTCCATTGCCAACTTCTCTGAGTTCTATGTGGAGCTGGAGGCAAACAATGAGGGTGTCGAGTGCCTGCGGCTGCTCAACGAGATCATCGCTGACTTTGATGAGATTATCAGCGAGGAGCGGTTCCGGCAGCTGGAAAAGATCAAGACGATTGGTAGCACCTACATGGCTGCCTCAGGGCTGAACGCCAGCACCTACGATCAGGTGGGCCGCTCCCACATCACTGCCCTGGCTGACTACGCCATGCGGCTCATGGAGCAGATGAAGCACATCAATGAGCACTCCTTCAACAATTTCCAGATGAAGATTGGGCTGAACATGGGCCCAGTCGTGGCAGGTGTCATCGGGGCTCGGAAGCCACAGTATGACATCTGGGGGAACACAGTGAATGTCTCTAGTCGTATGGACAGCACGGGGGTCCCCGACCGAATCCAGGTGACCACGGACCTGTACCAGGTTCTAGCTGCCAAGGGCTACCAGCTGGAGTGTCGAGGGGTGGTCAAGGTGAAGGGCAAGGGGGAGATGACCACCTACTTCCTCAATGGGGGCCCCAGCAGTTGA

SmBiT-linker-AC7

ATGGTCACCGGCTATCGGCTGTTTGAAGAGATTCTGGGGGGGTCAGGAGGAGGAGGCTCAGGGGGCTCATCATCAGGAGGACCAGCCAAGGGGCGCTACTTCCTCAACGAGGGCGAGGAGGGCCCTGACCAAGATGCGCTCTACGAGAAGTACCAGCTCACCAGCCAGCATGGGCCGCTGCTGCTCACGCTCCTGCTGGTGGCCGCCACTGCCTGCGTGGCCCTCATCATCATTGCCTTCAGCCAGGGGGACCCCTCCAGACACCAGGCCATTCTGGGCATGGCGTTCCTGGTGCTGGCGGTGTTTGCGGCCCTCTCTGTGCTGATGTACGTCGAGTGTCTCCTGCGGCGCTGGCTCAGGGCCTTGGCGCTGCTCACCTGGGCCTGCTTGGTGGCGCTGGGCTATGTGCTGGTGTTCGACGCATGGACAAAGGCGGCCTGTGCGTGGGAGCAGGTGCCCTTCTTCCTGTTCATTGTCTTCGTGGTGTACACACTACTGCCCTTCAGCATGCGGGGCGCTGTCGCCGTTGGGGCCGTCTCCACTGCCTCCCACCTCCTGGTGCTCGGTTCTTTGATGGGAGGCTTCACGACACCCAGTGTCCGGGTGGGGCTGCAGCTGCTGGCCAACGCAGTCATCTTCCTGTGTGGGAACCTGACAGGCGCCTTCCACAAGCACCAAATGCAGGATGCGTCCCGGGACCTCTTCACCTACACTGTGAAGTGCATCCAGATCCGCCGGAAGCTGCGCATCGAGAAGCGCCAGCAGGAGAACCTGCTGCTGTCAGTGCTTCCGGCCCACATCTCCATGGGCATGAAGCTGGCCATCATCGAACGGCTCAAGGAGCATGGTGACCGTCGCTGCATGCCTGACAACAACTTCCACAGCCTCTACGTCAAGAGGCACCAGAATGTCAGCATCCTCTATGCGGACATCGTGGGCTTCACGCAGCTGGCCAGCGACTGTTCTCCCAAGGAGCTGGTGGTGGTGCTGAATGAGCTCTTTGGCAAGTTCGACCAGATCGCCAAGGCCAACGAGTGCATGCGAATCAAGATCCTCGGCGACTGCTACTACTGTGTATCGGGCCTGCCCGTGTCGCTGCCTACCCACGCCCGGAACTGCGTGAAGATGGGGCTGGACATGTGCCAGGCCATCAAGCAGGTGCGGGAGGCCACGGGCGTGGACATCAACATGCGTGTGGGCATACACTCGGGGAATGTGCTGTGCGGGGTCATCGGGCTGCGCAAGTGGCAGTATGACGTGTGGTCCCACGACGTGTCCCTGGCCAACCGGATGGAGGCAGCCGGAGTACCCGGCCGGGTGCACATCACGGAGGCCACGCTAAAGCACCTGGACAAGGCGTACGAGGTGGAGGATGGGCACGGGCAGCAGCGGGACCCCTACCTCAAGGAGATGAACATCCGCACCTACCTGGTCATCGACCCCCGGAGCCAGCAGCCACCCCCGCCCAGCCAACACCTCCCCAGGCCCAAGGGGGACGCGGCCCTGAAGATGCGGGCGTCAGTGCGCATGACCCGGTACCTCGAGTCCTGGGGGGCGGCACGGCCCTTTGCACATCTCAACCACCGTGAGAGCGTGAGCAGTGGTGAGACCCACGTCCCCAACGGGCGGAGGCCTAAGAGCGTTCCCCAGCGCCACCGCCGGACCCCAGACAGAAGCATGTCCCCCAAGGGGCGGTCGGAGGATGACTCGTACGATGACGAGATGCTGTCAGCCATTGAGGGGCTCAGCTCCACGAGGCCCTGCTGCTCCAAGTCCGATGACTTCTACACCTTTGGGTCCATCTTCCTGGAGAAGGGCTTTGAGCGCGAGTACCGCCTGGCACCCATCCCCCGGGCCCGCCACGACTTTGCCTGCGCCAGCCTGATCTTCGTCTGCATCCTGCTCGTCCATGTCCTGCTCATGCCCAGGACGGCGGCACTGGGTGTGTCCTTCGGGCTGGTGGCCTGTGTACTGGGGCTGGTGCTGGGCCTGTGCTTTGCCACCAAGTTCTCGAGGTGCTGCCCAGCTCGGGGGACGCTCTGCACTATCTCTGAGAGGGTGGAGACACAGCCCCTGCTGAGGCTGACCCTGGCCGTCCTGACCATCGGCAGCCTGCTCACTGTGGCCATCATCAACCTGCCCCTGATGCCTTTCCAAGTTCCAGAGCTGCCTGTTGGCAATGAGACAGGCCTACTGGCCGCGAGCAGCAAGACAAGAGCCCTGTGTGAGCCCCTCCCGTACTACACCTGCAGCTGTGTCCTGGGCTTCATCGCCTGCTCGGTCTTCCTGAGGATGAGCCTGGAGCCAAAGGTTGTGCTGCTGACAGTGGCCCTGGTGGCCTACCTGGTGCTCTTCAACCTCTCCCCATGCTGGCAGTGGGACTGCTGCGGCCAAGGCCTGGGCAACCTCACCAAGCCCAACGGCACCACCAGTGGCACCCCTAGCTGTTCCTGGAAGGACCTGAAGACCATGACCAATTTCTACCTGGTCCTGTTCTACATCACCCTGCTTACACTCTCCAGACAGATTGACTATTACTGCCGCTTGGACTGCCTATGGAAGAAGAAGTTCAAGAAGGAGCACGAGGAGTTTGAGACCATGGAGAACGTGAACCGCCTTCTTCTGGAGAACGTCCTGCCAGCCCACGTGGCTGCCCACTTTATCGGTGACAAGTTAAACGAGGACTGGTACCATCAGTCCTATGACTGCGTCTGTGTCATGTTTGCCTCCGTGCCGGACTTCAAAGTGTTCTACACAGAGTGCGATGTCAACAAAGAAGGGCTGGAGTGCCTACGCCTGCTCAATGAGATCATTGCCGACTTCGACGAGCTCCTACTGAAGCCCAAGTTCAGCGGCGTGGAGAAGATCAAGACCATCGGCAGCACGTACATGGCAGCTGCAGGGCTCAGCGTCGCCTCAGGGCACGAGAACCAGGAGCTGGAGCGGCAGCATGCCCACATTGGTGTCATGGTGGAGTTCAGCATCGCCCTGATGAGTAAGCTGGACGGCATCAACAGGCACTCCTTCAACTCCTTCCGCCTCCGCGTCGGCATAAACCATGGGCCTGTGATTGCTGGAGTGATTGGGGCCCGAAAACCTCAGTATGACATCTGGGGAAACACTGTCAATGTGGCCAGCCGAATGGAAAGCACTGGAGAACTTGGGAAAATCCAGGTTACCGAGGAGACCTGCACCATCCTCCAGGGCCTCGGGTACTCTTGTGAATGCCGTGGCCTGATCAACGTCAAAGGCAAAGGCGAGCTGAGGACTTACTTTGTCTGTACGGACACTGCCAAGTTTCAGGGGCTGGGGCTGAACTGA

SmBiT-linker-AC8

ATGGTCACCGGCTATCGGCTGTTTGAAGAGATTCTGGGGGGGTCAGGAGGAGGAGGCTCAGGGGGCTCATCATCAGGAGGAGAGCTCTCCGATGTGCGCTGCCTTACAGGCAGCGAGGAACTCTACACCATCCACCCGACGCCCCCGGCCGGCGACGGCAGGAGCGCCTCCCGGCCGCAGCGGCTGCTGTGGCAGACGGCGGTGCGACACATCACGGAGCAGCGCTTCATTCACGGGCACCGGGGAGGCAGCGGCAGCGGGAGTGGAGGCTCGGGCAAAGCCTCGGACCCTGCGGGCGGCGGCCCCAACCACCACGCGCCGCAGCTGTCAGGCGACTCGGCGCTGCCCCTCTACTCGCTGGGCCCGGGAGAGCGAGCGCACAGCACCTGCGGCACCAAAGTCTTCCCGGAACGCAGCGGGAGCGGCAGTGCCAGCGGCAGCGGAGGCGGGGGCGACCTGGGCTTCCTGCACCTTGACTGTGCCCCTAGCAACTCGGATTTCTTTCTTAATGGGGGCTATAGCTACCGAGGGGTCATTTTCCCCACCCTGCGCAACTCCTTCAAATCTCGGGATTTGGAACGCCTCTACCAGCGCTATTTCTTGGGCCAAAGGCGCAAATCGGAAGTGGTGATGAACGTGCTGGACGTGCTGACCAAACTCACTCTCTTGGTCCTACACTTGAGCCTGGCCTCGGCCCCCATGGACCCGCTCAAGGGCATCCTGCTGGGCTTCTTCACCGGCATTGAGGTAGTGATCTGCGCCCTGGTGGTGGTCAGGAAGGACACCACCTCCCACACGTACCTGCAGTACAGCGGCGTGGTCACCTGGGTGGCCATGACCACCCAGATCCTGGCAGCAGGCCTCGGCTACGGGCTCCTGGGCGACGGCATAGGCTACGTGCTCTTCACGCTCTTCGCCACCTACAGTATGCTGCCGCTGCCGCTCACCTGGGCCATCCTGGCCGGCCTGGGCACCTCGCTGCTGCAGGTCATCCTCCAAGTGGTCATACCCCGGCTGGCGGTCATTTCCATCAACCAGGTTGTGGCCCAGGCAGTGCTATTCATGTGTATGAACACAGCTGGAATCTTCATCAGTTACCTGTCAGACCGGGCCCAGCGCCAAGCTTTCCTGGAGACTCGGAGGTGTGTGGAGGCCAGGCTGCGCCTGGAGACAGAGAACCAAAGACAGGAGCGGCTCGTGCTTTCTGTGCTCCCCCGGTTTGTTGTCCTGGAAATGATCAACGACATGACCAATGTGGAAGATGAGCACCTGCAGCACCAGTTCCATCGGATCTACATCCATCGCTATGAGAACGTCAGTATTCTTTTTGCAGATGTTAAAGGATTTACCAACCTCTCCACGACCTTGTCTGCTCAGGAGCTGGTCAGGATGCTCAACGAGCTCTTTGCCAGATTTGATCGACTGGCCCATGAGCATCACTGCCTTCGTATTAAAATCCTGGGGGACTGCTACTACTGCGTGTCTGGACTTCCTGAGCCCCGCCAGGACCATGCCCACTGCTGTGTTGAAATGGGTCTCAGCATGATCAAAACCATCAGGTATGTGCGGTCAAGGACAAAACACGATGTTGACATGAGGATTGGAATCCACTCCGGCTCGGTGCTGTGCGGTGTTTTGGGACTAAGGAAGTGGCAGTTTGATGTCTGGTCTTGGGATGTGGATATTGCAAACAAACTCGAATCTGGAGGAATCCCTGGGAGGATTCACATTTCCAAAGCCACGCTGGACTGTCTCAACGGTGACTATAACGTGGAAGAGGGCCATGGTAAAGAGAGGAATGAATTCCTGAGGAAGCATAATATCGAAACTTACTTAATTAAGCAGCCTGAGGACAGTCTGCTGTCCTTGCCTGAAGATATCGTCAAGGAGTCAGTGAGCTCCTCAGACCGGAGAAACAGTGGGGCCACATTCACTGAAGGATCCTGGAGCCCTGAACTGCCCTTTGATAATATCGTGGGGAAACAGAATACTCTGGCTGCCCTAACAAGAAATTCAATAAATCTGCTTCCAAACCATCTTGCACAAGCTTTGCATGTCCAGTCTGGGCCTGAGGAAATTAACAAGAGAATAGAACATACCATCGACTTGCGGAGTGGCGATAAATTGAGAAGAGAGCATATCAAGCCATTCTCACTGATGTTTAAAGACTCCAGCCTGGAGCACAAGTATTCTCAAATGAGGGATGAAGTGTTCAAGTCAAACTTGGTCTGTGCATTTATCGTTCTTCTATTTATCACGGCAATACAAAGTTTGCTTCCTTCTTCAAGAGTGATGCCAATGACCATCCAGTTCTCCATTCTGATTATGCTGCACTCGGCTCTGGTCCTCATCACCACAGCAGAGGATTATAAATGTTTGCCCCTCATCCTCCGGAAAACTTGCTGTTGGATTAATGAGACCTATTTGGCCCGGAACGTCATCATCTTTGCATCCATTTTGATTAATTTCCTGGGTGCCATCTTAAATATCCTGTGGTGTGATTTTGACAAGTCGATACCCTTGAAGAACCTGACTTTCAATTCCTCAGCTGTGTTTACAGATATCTGCTCCTACCCAGAGTACTTTGTCTTCACGGGGGTGTTGGCCATGGTGACCTGTGCAGTTTTCCTCCGGCTGAACTCCGTCCTGAAGCTGGCAGTGCTGCTGATCATGATTGCCATCTATGCCCTGCTCACTGAGACCGTCTACGCAGGCCTCTTTCTGCGTTATGACAACCTCAACCACAGTGGAGAAGATTTCCTGGGGACCAAGGAGGTATCACTGCTACTGATGGCCATGTTCCTCCTGGCTGTGTTCTACCATGGACAGCAGCTGGAGTACACAGCCCGCCTGGACTTCCTTTGGCGAGTACAGGCCAAAGAGGAGATCAATGAGATGAAGGAGCTGAGGGAACACAATGAGAACATGCTCCGGAATATCTTACCCAGCCATGTGGCCCGCCATTTCCTAGAGAAGGACCGAGACAATGAGGAGCTGTATTCTCAATCCTATGATGCTGTTGGGGTGATGTTTGCCTCCATCCCAGGATTTGCGGACTTTTACTCTCAGACTGAAATGAATAACCAGGGAGTGGAATGCCTGCGCTTGCTCAATGAGATCATTGCTGACTTCGATGAGTTGCTTGGTGAAGACCGATTTCAAGACATTGAAAAGATTAAGACCATTGGCAGCACCTACATGGCCGTGTCAGGCCTGTCACCTGAAAAACAGCAATGTGAAGACAAGTGGGGACATTTGTGTGCTCTGGCTGACTTCTCACTCGCCCTGACAGAAAGCATACAGGAGATCAACAAGCATTCATTCAACAATTTTGAACTCCGGATTGGCATCAGCCACGGCTCAGTGGTAGCTGGCGTTATCGGCGCTAAGAAACCACAGTATGACATTTGGGGCAAAACTGTGAACCTGGCAAGCCGAATGGACAGCACGGGGGTTAGTGGCCGGATCCAAGTCCCAGAGGAGACCTATCTCATCCTGAAGGACCAGGGCTTTGCCTTTGATTACCGAGGGGAGATCTATGTGAAGGGTATCAGTGAACAGGAAGGAAAAATCAAAACGTACTTTCTTCTGGGAAGAGTCCAACCCAACCCATTCATCTTGCCCCCAAGAAGACTGCCTGGGCAGTACTCCCTGGCCGCGGTTGTCCTGGGACTTGTCCAGTCCCTCAATAGGCAAAGGCAGAAGCAGCTACTCAATGAGAACAACAACACAGGAATCATCAAGGGTCATTACAACCGGCGGACTTTGTTGTCACCCAGCGGCACAGAGCCTGGAGCCCAGGCTGAAGGCACCGACAAATCTGATTTGCCATGA

SmBiT-linker-AC9

ATGGTCACCGGCTATCGGCTGTTTGAAGAGATTCTGGGGGGGTCAGGAGGAGGAGGCTCAGGGGGCTCATCATCAGGAGGAGCTTCCCCACCCCACCAGCAGCTGCTGCATCACCACAGCACCGAGGTGAGCTGCGACTCCAGCGGGGACAGCAACAGCGTGCGCGTCAAGATCAACCCCAAGCAGCTGTCCTCCAACAGCCACCCCAAGCACTGCAAATACAGCATCTCCTCTAGCTGCAGCAGCTCTGGGGACTCCGGGGGCGTCCCCCGGCGAGTGGGCGGCGGAGGCCGGCTGCGCAGGCAGAAGAAGCTGCCCCAGCTGTTCGAGAGGGCCTCCAGCCGCTGGTGGGACCCCAAGTTCGACTCGGTGAACCTGGAGGAGGCCTGCCTGGAGCGCTGCTTCCCGCAGACCCAGCGCCGGTTCCGGTATGCGCTCTTCTACATCGGCTTCGCCTGCCTTCTGTGGAGCATCTATTTTGCGGTCCACATGAGATCCAGACTGATCGTCATGGTCGCCCCCGCGCTGTGCTTCCTCCTGGTGTGTGTGGGCTTCTTTCTGTTTACCTTCACCAAGCTGTACGCCCGGCATTACGCGTGGACCTCGCTGGCTCTCACCCTGCTGGTGTTCGCCCTGACCCTGGCTGCGCAGTTCCAGGTCTTGACGCCTGTCTCAGGACGCGGCGACAGCTCCAACCTTACGGCCACAGCCCGGCCCACAGATACTTGCTTATCTCAAGTGGGGAGCTTCTCCATGTGCATCGAAGTGCTCTTTTTGCTCTATACCGTCATGCACTTACCTTTGTACCTGAGTTTGTGTCTGGGGGTGGCCTACTCTGTCCTTTTCGAGACCTTTGGCTACCATTTCCGGGATGAAGCCTGCTTCCCCTCGCCCGGAGCCGGGGCCCTGCACTGGGAGCTGCTGAGCAGGGGGCTGCTCCACGGCTGCATCCACGCCATCGGGGTCCACCTGTTCGTCATGTCCCAGGTGAGGTCCAGGAGCACCTTCCTCAAGGTGGGGCAATCCATTATGCACGGGAAGGACCTGGAAGTGGAAAAAGCCCTCAAAGAGAGGATGATTCATTCCGTGATGCCAAGAATCATAGCCGATGACTTAATGAAGCAGGGAGATGAGGAGAGTGAGAATTCTGTCAAGAGGCATGCCACCTCGAGCCCCAAGAACAGGAAGAAAAAGTCTTCCATCCAAAAAGCTCCTATAGCCTTCCGCCCTTTTAAGATGCAGCAGATCGAAGAAGTCAGTATTTTATTTGCAGATATCGTGGGCTTCACCAAGATGAGTGCCAACAAGTCTGCCCACGCCCTGGTGGGTCTCCTGAACGATCTGTTCGGTCGCTTCGACCGCCTGTGTGAGGAGACCAAGTGTGAGAAAATCAGCACCCTGGGAGACTGTTACTACTGCGTGGCGGGCTGTCCCGAGCCCCGGGCCGACCATGCCTACTGCTGCATCGAGATGGGCCTGGGCATGATCAAGGCCATCGAGCAGTTCTGCCAGGAGAAGAAGGAGATGGTGAACATGAGAGTCGGGGTGCACACGGGCACCGTCCTTTGCGGCATCCTGGGCATGAGGAGGTTTAAATTTGACGTGTGGTCCAACGATGTGAACCTGGCCAATCTCATGGAGCAGCTGGGAGTGGCCGGCAAAGTTCACATTTCTGAGGCCACCGCAAAATACTTAGATGACCGGTACGAAATGGAAGATGGGAAAGTTATTGAACGGCTGGGCCAGAGCGTGGTTGCTGACCAGTTGAAAGGTTTGAAGACATACCTGATATCGGGTCAGAGAGCCAAGGAGTCTCGCTGCAGCTGTGCAGAGGCCTTGCTTTCTGGCTTTGAGGTCATTGACGGCTCACAGGTGTCCTCAGGCCCTAGGGGACAGGGGACAGCGTCATCAGGGAATGTCAGTGACTTGGCGCAGACTGTCAAAACCTTTGATAACCTTAAGACCTGCCCTTCGTGCGGAATCACATTTGCTCCCAAATCTGAAGCCGGCGCCGAGGGAGGAGCACCTCAAAACGGCTGCCAAGACGAGCATAAAAACAGCACCAAGGCTTCTGGAGGACCTAATCCCAAAACTCAGAACGGGCTCCTCAGCCCTCCCCAAGAGGAGAAGCTCACCAACAGTCAGACTTCTCTGTGTGAGATCTTGCAGGAGAAGGGAAGGTGGGCAGGGGTGAGCCTGGACCAGTCGGCTCTCCTTCCGCTGAGGTTCAAGAACATCCGGGAGAAAACGGACGCCCACTTTGTGGACGTTATCAAAGAAGACAGCCTGATGAAAGATTACTTTTTTAAGCCGCCCATTAATCAGTTCAGCCTGAACTTCCTGGATCAGGAGCTGGAGCGATCCTACAGGACCAGCTATCAGGAAGAGGTCATAAAGAACTCCCCCGTGAAGACGTTTGCTAGTCCCACCTTCAGCTCCCTCCTGGATGTGTTTCTGTCGACCACAGTGTTTCTGACGCTGTCCACCACCTGCTTCCTGAAGTACGAGGCGGCCACCGTGCCTCCCCCGCCCGCCGCCCTGGCGGTCTTCAGTGCAGCCCTGCTGCTGGAGGTGCTGTCCCTCGCGGTGTCCATCAGGATGGTGTTCTTCCTGGAGGACGTCATGGCCTGCACCAAGCGCCTGCTGGAGTGGATCGCCGGCTGGCTACCACGTCACTGCATCGGGGCCATCCTGGTGTCGCTTCCCGCACTGGCCGTCTACTCCCATGTCACCTCCGAATATGAGACCAACATACACTTCCCAGTGTTCACAGGCTCGGCCGCGCTGATTGCCGTCGTGCACTACTGTAACTTCTGCCAGCTCAGCTCCTGGATGAGGTCCTCCCTCGCCACCGTCGTGGGGGCCGGGCCGCTGCTCCTGCTCTACGTCTCCCTGTGCCCAGACAGTTCTGTATTAACTTCGCCCCTTGACGCAGTACAGAATTTCAGTTCCGAGAGGAACCCGTGCAATAGTTCGGTGCCGCGTGACCTCCGGCGGCCCGCCAGCCTCATCGGCCAGGAGGTGGTTCTCGTCTTCTTTCTCCTGCTCTTGTTGGTCTGGTTCCTGAATCGCGAATTTGAAGTCAGCTACCGCCTCCACTACCACGGAGACGTGGAAGCGGATCTTCACCGCACCAAGATCCAGAGCATGCGGGACCAGGCAGACTGGCTGCTGAGGAACATCATCCCCTACCACGTGGCTGAGCAGCTGAAGGTGTCCCAGACCTACTCCAAGAACCATGACAGCGGAGGGGTGATCTTCGCCAGCATCGTCAACTTCAGCGAGTTCTACGAGGAGAACTACGAGGGCGGCAAGGAGTGCTACCGGGTCCTCAACGAGCTCATCGGGGACTTTGACGAGCTCCTAAGCAAGCCGGACTACAGCAGCATCGAGAAGATCAAGACCATCGGAGCCACGTACATGGCGGCGTCAGGGCTGAACACCGCGCAGGCCCAGGACGGCAGCCACCCGCAGGAGCACCTGCAGATCCTGTTCGAGTTCGCCAAGGAGATGATGCGCGTGGTGGACGACTTCAACAACAACATGCTGTGGTTCAACTTCAAGCTCCGCGTCGGCTTCAACCATGGGCCCCTCACGGCCGGGGTCATCGGCACCACCAAGCTGCTGTACGACATCTGGGGAGACACCGTCAACATCGCCAGCAGGATGGACACCACCGGCGTGGAGTGCCGCATCCAGGTGAGCGAAGAGAGCTACCGCGTCTTGAGCAAGATGGGCTATGACTTCGACTACAGAGGGACCGTGAATGTCAAGGGGAAAGGCCAGATGAAGACCTACCTGTACCCAAAGTGCACGGATCACAGGGTCATCCCACAGCACCAGCTGTCCATCTCCCCAGACATCCGCGTCCAGGTGGATGGCAGCATCGGACGGTCTCCCACAGACGAGATTGCCAACCTGGTGCCTTCTGTCCAGTATGTGGACAAGACATCTCTGGGTTCTGACAGCAGCACGCAGGCCAAGGATGCCCACCTGTCCCCCAAGAGACCGTGGAAGGAGCCCGTCAAAGCCGAAGAAAGGGGTCGATTTGGCAAAGCCATAGAGAAAGACGACTGTGACGAAACAGGAATAGAAGAAGCCAACGAACTCACCAAGCTCAACGTTTCAAAGAGTGTGTGA

SmBiT-linker-PLCβ1

ATGGTCACCGGCTATCGGCTGTTTGAAGAGATTCTGGGGGGGTCAGGAGGAGGAGGCTCAGGGGGCTCATCATCAGGAGGAGCCGGGGCTCAACCCGGAGTGCACGCCTTGCAACTCAAGCCCGTGTGCGTGTCCGACAGCCTCAAGAAGGGCACCAAATTCGTCAAGTGGGATGATGACTCAACTATTGTTACTCCAATTATTTTGAGGACTGACCCTCAGGGATTTTTCTTTTACTGGACAGATCAAAACAAGGAGACAGAGCTACTGGATCTCAGCCTTGTCAAAGATGCCAGATGTGGGAGACACGCCAAAGCTCCCAAGGACCCCAAATTACGTGAACTTTTGGATGTGGGGAACATCGGGCGCCTGGAGCAGCGCATGATCACAGTGGTGTATGGGCCTGACCTCGTGAACATCTCCCATTTGAATCTCGTGGCTTTCCAAGAAGAAGTGGCCAAGGAATGGACAAATGAGGTTTTCAGTTTGGCAACAAACCTGCTGGCCCAAAACATGTCCAGGGATGCATTTCTGGAAAAAGCCTATACTAAACTTAAGCTGCAAGTCACTCCAGAAGGGCGTATTCCTCTCAAAAACATATATCGCTTGTTTTCAGCAGATCGGAAGCGAGTTGAAACTGCTTTAGAGGCTTGTAGTCTTCCATCTTCAAGGAATGATTCAATACCTCAAGAAGATTTCACTCCAGAAGTGTACAGAGTTTTCCTCAACAACCTTTGCCCTCGACCTGAAATTGATAACATCTTTTCAGAATTTGGTGCAAAAAGCAAACCATATCTTACCGTTGATCAGATGATGGATTTTATCAACCTTAAGCAGCGAGATCCTCGGCTTAATGAAATACTTTATCCACCTCTAAAACAAGAGCAAGTCCAAGTATTGATTGAGAAGTATGAACCCAACAACAGCCTCGCCAGAAAAGGACAAATATCAGTGGATGGGTTCATGCGCTATCTGAGTGGAGAAGAAAACGGAGTCGTTTCACCTGAGAAACTGGATTTGAATGAAGACATGTCTCAGCCCCTTTCTCACTATTTCATTAATTCCTCGCACAACACCTACCTCACAGCTGGCCAACTGGCTGGAAACTCCTCTGTTGAGATGTATCGCCAAGTGCTCCTGTCTGGTTGTCGCTGTGTGGAGCTGGACTGCTGGAAGGGACGGACTGCAGAAGAGGAACCTGTCATCACCCATGGCTTCACCATGACAACTGAAATATCTTTCAAGGAAGTGATAGAAGCAATTGCGGAGTGTGCATTTAAGACTTCACCTTTTCCAATTCTCCTTTCGTTTGAGAACCATGTGGATTCCCCAAAGCAGCAAGCCAAGATGGCGGAGTACTGCCGACTGATCTTTGGGGATGCCCTTCTCATGGAGCCCCTGGAAAAATATCCACTGGAATCTGGAGTTCCTCTTCCAAGCCCTATGGATTTAATGTATAAAATTTTGGTGAAAAATAAGAAGAAATCACACAAGTCATCAGAAGGAAGCGGCAAAAAGAAGCTCTCAGAACAAGCCTCCAACACCTACAGTGACTCCTCCAGCATGTTCGAGCCCTCATCCCCAGGAGCCGGAGAAGCTGATACGGAAAGTGACGACGACGATGATGATGATGACTGTAAAAAATCTTCAATGGATGAGGGGACTGCTGGAAGTGAGGCTATGGCCACAGAAGAAATGTCTAATCTGGTGAACTATATTCAGCCAGTCAAGTTTGAGTCATTTGAAATTTCAAAAAAAAGAAATAAAAGTTTTGAAATGTCTTCCTTCGTGGAAACCAAAGGACTTGAACAACTCACCAAGTCTCCAGTGGAATTTGTAGAATATAACAAAATGCAGCTTAGCAGGATATATCCAAAAGGAACACGTGTGGATTCATCCAACTATATGCCTCAGCTCTTCTGGAATGCAGGTTGTCAGATGGTGGCACTTAATTTCCAGACAATGGACCTGGCTATGCAAATAAATATGGGGATGTATGAATACAACGGGAAGAGTGGCTACAGATTGAAGCCAGAGTTCATGAGGAGGCCTGACAAGCATTTTGATCCATTTACTGAAGGCATCGTAGATGGGATAGTGGCAAACACTTTGTCTGTTAAGATTATTTCAGGTCAGTTTCTTTCTGATAAGAAAGTTGGGACTTACGTGGAAGTAGATATGTTTGGTTTGCCTGTGGATACAAGGAGGAAGGCATTTAAGACCAAAACATCCCAAGGAAATGCTGTGAATCCTGTCTGGGAAGAAGAACCTATTGTGTTCAAAAAGGTGGTTCTTCCTACTCTGGCCTGTTTGAGAATAGCAGTTTATGAAGAAGGAGGTAAATTCATTGGCCACCGTATCTTGCCAGTGCAAGCCATTCGGCCAGGCTATCACTATATCTGTCTAAGGAATGAAAGGAACCAGCCTCTGACGCTGCCTGCTGTCTTTGTCTACATAGAAGTGAAAGACTATGTGCCAGACACATATGCAGATGTCATCGAAGCTTTATCAAACCCAATCCGATATGTGAACCTGATGGAACAGAGAGCTAAGCAATTGGCTGCTTTGACACTGGAAGATGAAGAAGAAGTAAAGAAAGAGGCTGATCCTGGAGAAACACCATCAGAGGCTCCAAGTGAAGCGAGAACGACTCCAGCAGAAAATGGGGTGAATCACACTACAACCCTGACACCCAAGCCACCCTCCCAGGCTCTCCACAGCCAGCCAGCTCCAGGTTCTGTAAAGGCACCTGCCAAAACAGAAGATCTTATTCAGAGTGTCTTAACAGAAGTGGAAGCACAGACCATCGAAGAACTAAAGCAACAGAAATCGTTTGTGAAACTTCAAAAGAAACACTACAAAGAAATGAAAGACCTGGTTAAGAGACACCACAAGAAAACCACTGACCTTATCAAAGAACACACTACCAAGTATAATGAAATTCAGAATGACTACTTGAGAAGGAGAGCCGCTTTGGAAAAGTCCGCCAAAAAGGACAGTAAGAAAAAATCGGAACCCAGCAGCCCTGATCATGGTTCATCAACGATTGAGCAAGACCTCGCTGCTCTGGATGCTGAAATGACCCAAAAGTTAATAGACTTGAAGGACAAACAACAGCAGCAGCTGCTTAATCTTCGGCAAGAACAGTATTATAGTGAAAAATACCAGAAGCGAGAACATATTAAACTGCTTATTCAAAAGTTGACGGATGTCGCAGAAGAGTGTCAGAACAATCAGTTAAAGAAGCTCAAAGAAATCTGTGAGAAAGAAAAGAAAGAATTAAAGAAGAAAATGGATAAAAAGAGGCAGGAGAAGATAACAGAAGCTAAATCCAAAGACAAAAGTCAGATGGAAGAGGAGAAGACAGAGATGATCCGGTCATATATCCAGGAAGTGGTGCAGTATATCAAGAGGCTAGAAGAAGCGCAAAGTAAACGGCAAGAAAAACTCGTAGAGAAACACAAGGAAATACGTCAGCAGATCCTGGATGAAAAGCCCAAGCTGCAGGTGGAGCTGGAGCAAGAATACCAAGACAAATTCAAAAGACTGCCCCTCGAGATTTTGGAATTCGTGCAGGAAGCCATGAAAGGAAAGATCAGTGAAGACAGCAATCACGGTTCTGCCCCTCTCTCCCTGTCCTCAGACCCTGGAAAAGTGAACCACAAGACTCCCTCCAGTGAGGAGCTGGGAGGAGACATCCCAGGAAAAGAATTTGATACTCCTCTGTGA

SmBiT-linker-PLCβ2

ATGGTCACCGGCTATCGGCTGTTTGAAGAGATTCTGGGGGGGTCAGGAGGAGGAGGCTCAGGGGGCTCATCATCAGGAGGATCTCTGCTCAACCCTGTCCTGCTGCCCCCCAAGGTGAAGGCCTATCTGAGCCAAGGGGAGCGCTTCATCAAATGGGATGATGAAACTACAGTTGCCTCTCCAGTTATCCTCCGTGTGGATCCTAAGGGCTACTACTTATACTGGACGTATCAAAGTAAGGAGATGGAGTTTCTGGATATCACCAGCATCCGGGATACTCGCTTTGGGAAGTTTGCCAAGATGCCCAAGAGCCAGAAGCTCCGGGACGTCTTCAACATGGACTTTCCTGATAACAGTTTCCTGCTGAAGACACTCACGGTGGTGTCCGGCCCGGACATGGTGGACCTCACCTTCCACAACTTCGTCTCCTACAAGGAGAACGTGGGCAAGGCCTGGGCTGAGGACGTACTGGCCCTAGTCAAACATCCGCTGACGGCCAACGCCTCCCGCAGCACCTTCCTGGACAAGATCCTTGTGAAGCTCAAGATGCAGCTCAACTCTGAAGGGAAGATTCCGGTGAAGAACTTTTTCCAGATGTTTCCTGCTGACCGCAAGCGGGTGGAAGCTGCTCTCAGTGCCTGCCACCTCCCCAAAGGCAAAAATGACGCCATCAATCCTGAGGACTTCCCAGAACCTGTCTACAAGAGTTTCCTCATGAGCCTCTGTCCTCGGCCAGAAATAGATGAGATCTTCACTTCTTACCATGCTAAGGCCAAACCCTACATGACGAAGGAGCACCTGACCAAATTCATCAACCAGAAACAGCGGGACTCCCGGCTTAACTCCCTGCTGTTCCCGCCAGCACGGCCTGACCAGGTGCAGGGCCTCATCGACAAGTATGAGCCCAGTGGCATCAATGCACAGAGGGGCCAGCTGTCACCTGAAGGCATGGTCTGGTTTCTCTGTGGGCCAGAGAACAGCGTGCTGGCCCAGGACAAGCTGCTGCTCCACCACGACATGACGCAGCCACTCAATCATTACTTCATCAACTCGTCCCACAACACCTACCTGACAGCCGGCCAGTTCTCAGGCCTCTCCTCGGCTGAGATGTACCGCCAGGTGCTGCTCTCTGGCTGCCGTTGCGTGGAGCTAGACTGCTGGAAGGGGAAACCCCCTGACGAGGAGCCCATTATCACCCATGGCTTCACCATGACCACAGACATCTTCTTCAAAGAAGCAATTGAGGCTATTGCAGAAAGCGCCTTTAAGACCTCCCCCTATCCCATCATCCTGTCGTTTGAGAACCATGTGGACTCACCCCGCCAGCAGGCTAAGATGGCTGAGTATTGCCGGACGATCTTTGGGGATATGCTGCTCACAGAGCCCCTGGAAAAGTTCCCACTAAAACCAGGTGTCCCCCTGCCCAGCCCTGAGGATCTCAGGGGCAAGATCCTCATCAAGAACAAGAAGAACCAGTTTTCTGGCCCCACCTCCTCCAGTAAGGATACTGGTGGGGAGGCTGAGGGCAGCAGCCCACCCAGTGCCCCTGCAGTGTGGGCTGGCGAGGAAGGGACTGAGCTGGAGGAGGAGGAGGTGGAAGAGGAAGAGGAGGAGGAGTCAGGAAACCTGGATGAAGAAGAGATTAAGAAGATGCAGTCGGATGAGGGCACAGCGGGCCTGGAAGTGACGGCTTATGAGGAGATGTCCAGCCTAGTCAATTACATCCAGCCCACCAAGTTCGTCTCCTTTGAGTTCTCTGCCCAAAAGAACCGAAGTTATGTCATCTCGTCCTTCACAGAGCTCAAGGCATATGACCTGCTCTCCAAGGCCTCGGTGCAGTTTGTGGACTACAACAAGCGCCAGATGAGCCGCATTTACCCCAAGGGAACCCGCATGGACTCCTCCAACTACATGCCCCAGATGTTCTGGAATGCTGGATGCCAGATGGTTGCCCTCAACTTCCAGACGATGGACTTGCCCATGCAGCAGAACATGGCAGTATTTGAGTTCAACGGGCAGAGCGGCTACCTCCTCAAGCATGAGTTCATGCGCCGGCCGGACAAGCAGTTCAACCCCTTCTCAGTGGACCGCATCGACGTGGTGGTGGCCACCACCCTTTCCATTACGGTGATCTCTGGGCAGTTCCTGTCAGAACGCAGCGTGCGCACCTATGTAGAAGTGGAGCTGTTTGGCCTTCCTGGGGACCCCAAGAGGCGCTATCGAACTAAGCTGTCACCCAGTACTAACTCCATCAATCCTGTCTGGAAGGAGGAGCCCTTTGTCTTTGAGAAGATCTTGATGCCTGAGCTGGCCTCCCTCAGAGTGGCTGTGATGGAGGAAGGCAACAAGTTTCTTGGACACCGCATCATCCCCATCAATGCCCTAAATTCTGGGTACCACCACCTGTGCCTGCACAGTGAGAGCAACATGCCCCTCACCATGCCTGCGCTCTTCATCTTCCTGGAGATGAAGGACTACATACCTGGTGCTTGGGCAGATCTCACTGTGGCCCTCGCCAACCCCATTAAGTTCTTCAGTGCCCATGACACGAAGTCTGTGAAGCTCAAGGAGGCCATGGGAGGTCTGCCTGAGAAGCCCTTCCCACTGGCGAGTCCAGTTGCCAGCCAGGTCAATGGGGCGTTGGCCCCAACGAGCAATGGGTCACCAGCAGCCAGGGCCGGGGCCAGGGAAGAGGCTATGAAAGAAGCTGCGGAGCCGCGGACCGCCAGCCTGGAGGAGCTCCGGGAGCTAAAGGGCGTGGTGAAGCTGCAGCGGCGGCACGAGAAGGAGCTGCGAGAGTTGGAGCGGCGCGGAGCGCGGCGCTGGGAGGAGCTGCTGCAGCGGGGCGCGGCGCAGCTGGCGGAGCTCGGGCCACCGGGCGTGGGGGGCGTCGGGGCCTGCAAGCTCGGTCCCGGCAAGGGCTCTCGCAAGAAGAGGAGCCTGCCCCGCGAGGAGAGCGCCGGAGCCGCGCCGGGCGAGGGCCCTGAGGGCGTGGACGGGCGCGTGCGGGAGCTGAAAGACAGGCTGGAGCTGGAGCTGCTGCGGCAGGGCGAGGAGCAGTACGAGTGCGTTCTGAAGCGCAAGGAGCAGCACGTGGCCGAGCAAATCTCCAAAATGATGGAGCTGGCCAGAGAGAAACAGGCGGCAGAGCTGAAGGCCCTGAAGGAGACGTCGGAGAACGACACCAAAGAGATGAAGAAAAAGCTGGAGACAAAGAGACTGGAGCGGATCCAGGGCATGACCAAAGTCACCACAGACAAGATGGCCCAGGAGAGGTTGAAGAGAGAGATTAACAACTCCCACATCCAGGAAGTAGTGCAGGTGATCAAGCAGATGACGGAGAACTTGGAGAGGCACCAGGAGAAGCTGGAGGAGAAGCAGGCGGCTTGCCTGGAACAGATACGGGAGATGGAAAAGCAGTTCCAGAAGGAGGCGCTGGCAGAGTACGAGGCCAGGATGAAGGGTCTGGAGGCAGAGGTGAAGGAGTCGGTGAGGGCCTGCCTCAGGACCTGCTTTCCCTCCGAGGCCAAGGACAAGCCTGAGAGGGCCTGCGAGTGCCCCCCAGAGCTGTGTGAGCAGGACCCACTCATAGCAAAGGCAGATGCCCAGGAGAGCCGCCTCTGA

SmBiT-linker-PLCβ3

ATGGTCACCGGCTATCGGCTGTTTGAAGAGATTCTGGGGGGGTCAGGAGGAGGAGGCTCAGGGGGCTCATCATCAGGAGGAGCGGGCGCCCAGCCCGGCGTCCACGCGCTGCAGTTGGAGCCGCCCACCGTGGTGGAGACCCTGCGGCGCGGGAGTAAGTTCATCAAATGGGACGAGGAGACCTCCAGTCGGAACCTGGTGACCCTGCGTGTGGACCCCAATGGCTTCTTCTTGTACTGGACGGGCCCCAACATGGAGGTGGACACACTGGACATCAGTTCCATCAGGGACACACGGACAGGCCGGTACGCCCGCCTGCCCAAGGACCCCAAGATCCGGGAAGTTCTGGGCTTTGGGGGTCCCGATGCCCGGCTGGAGGAGAAGCTGATGACGGTGGTGTCTGGGCCAGACCCAGTGAACACAGTGTTCTTGAACTTCATGGCCGTGCAGGATGACACAGCCAAGGTCTGGTCTGAGGAGCTATTCAAGCTGGCTATGAACATCCTGGCTCAGAACGCCTCCCGGAACACCTTCCTGCGCAAAGCATACACGAAGCTGAAGCTGCAGGTGAACCAGGATGGTCGGATCCCCGTCAAGAACATCCTGAAGATGTTCTCAGCAGACAAGAAGCGGGTGGAGACTGCGCTGGAATCCTGTGGCCTCAAATTCAACCGGAGTGAGTCCATCCGGCCTGATGAGTTTTCCTTGGAAATCTTTGAGCGGTTCCTGAACAAGCTGTGTCTGCGGCCGGACATTGACAAGATCCTGCTGGAGATAGGCGCCAAGGGCAAGCCATACCTGACGCTGGAGCAGCTCATGGACTTCATCAACCAGAAGCAACGCGACCCGAGACTCAACGAAGTGCTGTACCCGCCCCTGCGGCCCTCCCAGGCCCGGCTGCTCATCGAAAAGTATGAGCCCAACCAGCAGTTTCTGGAGCGAGACCAGATGTCCATGGAGGGCTTTAGCCGCTACCTGGGAGGCGAGGAGAATGGCATCCTGCCCCTGGAAGCCCTGGATCTGAGCACGGACATGACCCAGCCACTGAGTGCCTACTTCATCAACTCCTCGCATAACACCTATCTCACTGCGGGGCAGCTGGCTGGGACCTCGTCGGTGGAGATGTACCGCCAGGCACTACTATGGGGCTGCCGCTGCGTGGAGCTGGACGTGTGGAAGGGACGGCCGCCTGAGGAGGAACCCTTCATTACCCACGGCTTCACCATGACCACAGAGGTGCCTCTGCGCGACGTGCTGGAGGCCATTGCCGAGACTGCCTTCAAGACCTCGCCCTACCCCGTCATCCTCTCCTTCGAGAACCATGTGGACTCGGCAAAGCAACAGGCAAAGATGGCTGAGTACTGCCGCTCCATCTTTGGAGACGCGCTACTCATCGAGCCTCTGGACAAGTACCCGCTGGCCCCAGGCGTTCCCCTGCCCAGCCCCCAGGACCTGATGGGCCGTATCCTGGTGAAGAACAAGAAGCGGCACCGACCCAGCGCAGGTGGCCCAGACAGCGCCGGGCGCAAGCGGCCCCTGGAGCAGAGCAATTCTGCCCTGAGCGAGAGCTCCGCGGCCACCGAGCCCTCCTCCCCGCAGCTGGGGTCTCCCAGCTCTGACAGCTGCCCAGGCCTGAGCAATGGGGAGGAGGTAGGGCTTGAGAAGCCCAGCCTGGAGCCTCAGAAGTCTCTGGGTGACGAGGGCCTGAACCGAGGCCCCTATGTTCTTGGACCTGCTGACCGTGAGGATGAGGAGGAAGATGAGGAAGAGGAGGAACAGACAGACCCCAAAAAGCCAACTACAGATGAGGGCACAGCCAGCAGCGAGGTGAATGCCACTGAGGAGATGTCCACGCTTGTCAACTACATCGAACCTGTCAAGTTCAAGTCCTTTGAGGCTGCTCGAAAGAGGAACAAATGCTTCGAGATGTCGTCCTTTGTGGAGACCAAGGCCATGGAGCAACTGACCAAGAGCCCCATGGAGTTTGTGGAATACAACAAGCAGCAGCTCAGCCGCATCTACCCCAAGGGCACCCGCGTGGACTCCTCCAACTACATGCCCCAGCTCTTCTGGAACGTAGGGTGCCAGCTTGTTGCGCTCAACTTCCAGACCCTCGATGTGGCGATGCAGCTCAACGCGGGCGTTTTTGAGTACAACGGGCGCAGCGGGTACCTGCTCAAGCCGGAGTTCATGCGGCGGCCGGACAAGTCCTTCGACCCCTTCACTGAGGTCATCGTGGATGGCATCGTGGCCAATGCCTTGCGGGTCAAGGTGATCTCAGGGCAGTTCCTGTCCGACAGGAAGGTGGGCATCTACGTGGAGGTGGACATGTTTGGCCTCCCTGTTGATACGCGGCGCAAGTACCGCACCCGGACCTCTCAGGGGAACTCGTTCAACCCCGTGTGGGACGAAGAGCCCTTCGACTTCCCCAAGGTGGTGCTGCCCACGCTGGCTTCACTTCGCATTGCAGCCTTTGAGGAGGGGGGTAAATTCGTAGGGCACCGGATCCTGCCTGTCTCTGCCATCCGCTCCGGATACCACTACGTCTGCCTGCGGAACGAGGCCAACCAACCGCTGTGCCTGCCGGCCCTGCTCATCTACACCGAAGCCTCGGACTACATTCCTGACGACCACCAGGACTATGCGGAGGCCCTGATCAACCCCATTAAGCACGTCAGCCTGATGGACCAGAGGGCCCGGCAGCTGGCCGCCCTCATTGGGGAGAGTGAGGCTCAGGCTGGCCAAGAGACGTGCCAGGACACCCAGTCTCAGCAGCTGGGGTCTCAGCCGTCCTCAAACCCCACCCCCAGCCCACTGGATGCCTCCCCCCGCCGGCCCCCTGGCCCCACCACCTCCCCTGCCAGCACCTCCCTCAGCAGCCCAGGGCAGCGTGATGATCTCATCGCCAGCATCCTCTCAGAGGTGGCCCCCACCCCGCTGGATGAGCTCCGAGGTCACAAGGCTCTGGTCAAGCTCCGGAGCCGGCAAGAGCGAGACCTGCGGGAGCTGCGCAAGAAGCATCAGCGGAAGGCAGTCACCCTCACCCGCCGCCTGCTGGATGGCCTGGCTCAGGCACAGGCTGAGGGCAGGTGCCGGCTGCGGCCAGGTGCCCTAGGTGGGGCCGCTGATGTGGAGGACACGAAGGAGGGGGAGGACGAGGCAAAGCGGTATCAGGAGTTCCAGAACAGACAGGTGCAGAGCCTGCTGGAGCTGCGGGAGGCCCAGGTGGACGCAGAGGCCCAGCGGAGGCTGGAACACCTGAGACAGGCTCTGCAGCGGCTCAGGGAGGTCGTCCTTGATGCAAACACAACTCAGTTCAAGAGGCTGAAAGAGATGAACGAGAGGGAGAAGAAGGAGCTGCAGAAGATCCTGGACAGAAAGCGCCATAACAGCATCTCGGAGGCCAAGATGAGGGACAAGCATAAGAAGGAGGCGGAACTGACGGAGATTAACCGTCGGCACATCACTGAGTCAGTCAACTCCATCCGTCGGCTGGAGGAGGCCCAGAAGCAGCGGCATGACCGTCTTGTGGCTGGGCAGCAGCAGGTCCTGCAACAGCTGGCAGAAGAGGAGCCCAAGCTGCTGGCCCAGCTGGCCCAGGAGTGTCAGGAGCAGCGGGCGAGGCTCCCCCAGGAGATCCGCCGGAGCCTGCTGGGCGAGATGCCGGAGGGGCTGGGGGACGGGCCTCTGGTGGCCTGTGCCAGCAACGGTCACGCACCCGGGAGCAGCGGGCACCTGTCGGGCGCTGACTCGGAGAGCCAGGAGGAGAACACGCAGCTCTGA

SmBiT-linker-PLCβ4

ATGGTCACCGGCTATCGGCTGTTTGAAGAGATTCTGGGGGGGTCAGGAGGAGGAGGCTCAGGGGGCTCATCATCAGGAGGAGCCAAACCTTATGAATTTAACTGGCAGAAGGAAGTTCCCTCCTTTTTGCAAGAAGGAGCAGTTTTTGACAGATACGAGGAGGAATCCTTTGTGTTTGAACCCAACTGCCTCTTCAAAGTGGATGAGTTTGGCTTCTTTCTGACATGGAGAAGTGAAGGCAAGGAAGGACAGGTGCTAGAATGCTCCCTCATCAACAGTATTCGGTCGGGAGCCATACCAAAGGATCCCAAAATCTTGGCTGCTCTTGAAGCTGTTGGAAAATCAGAAAATGATCTGGAAGGGCGGATAGTTTGTGTCTGCAGTGGCACAGATCTAGTGAACATTAGTTTTACCTACATGGTGGCTGAAAATCCAGAAGTAACTAAGCAATGGGTAGAAGGCCTGAGATCAATCATACACAACTTCAGGGCCAACAACGTCAGTCCAATGACATGCCTCAAGAAACACTGGATGAAATTGGCATTTATGACCAACACAAATGGTAAAATTCCAGTTAGGAGTATTACTAGAACATTTGCATCGGGAAAAACAGAAAAGGTGATCTTTCAAGCACTCAAGGAGTTAGGTCTTCCCAGTGGAAAGAATGATGAAATTGAGCCCACAGCATTTTCTTATGAAAAGTTCTATGAACTGACACAAAAGATTTGTCCTCGGACAGATATAGAAGATCTTTTCAAAAAAATCAATGGAGACAAAACTGATTATTTAACGGTAGACCAATTAGTGAGCTTTCTAAATGAACATCAACGAGATCCTCGATTGAATGAAATTTTATTTCCATTTTATGATGCCAAAAGGGCAATGCAGATCATTGAGATGTATGAACCTGATGAAGATTTGAAGAAAAAAGGCCTTATATCAAGTGATGGGTTTTGCAGATATCTGATGTCAGATGAAAACGCCCCAGTCTTCCTAGATCGTTTAGAACTTTACCAAGAAATGGACCATCCTCTGGCTCACTACTTCATCAGTTCTTCCCATAACACTTATCTCACTGGCAGACAGTTCGGCGGGAAGTCTTCGGTAGAAATGTACAGACAGGTTCTCCTGGCTGGTTGCAGATGTGTTGAACTTGACTGCTGGGATGGAAAAGGTGAAGACCAAGAACCAATAATAACTCATGGAAAAGCAATGTGTACAGATATCCTTTTTAAGGATGTAATTCAAGCCATCAAGGAAACTGCATTTGTCACATCAGAATATCCTGTAATTCTCTCCTTTGAAAATCACTGCAGCAAATATCAACAGTACAAGATGTCCAAATATTGCGAAGATCTATTTGGGGATCTCCTGTTGAAACAAGCACTTGAATCACATCCACTTGAACCAGGCAGGGCTTTGCCATCCCCCAATGACCTCAAAAGAAAAATACTCATAAAAAACAAGCGGCTGAAACCTGAAGTTGAAAAAAAACAGCTGGAAGCTTTGAGAAGCATGATGGAAGCTGGAGAATCTGCCTCCCCAGCAAACATCTTAGAGGACGATAATGAAGAGGAGATCGAAAGTGCTGACCAAGAGGAGGAAGCTCACCCCGAATTCAAATTTGGAAATGAACTTTCTGCTGATGACTTGGGTCACAAGGAAGCTGTTGCAAATAGCGTCAAGAAGGGCCTGGTCACTGTAGAAGATGAGCAGGCGTGGATGGCATCTTATAAATATGTAGGTGCTACCACTAATATCCATCCATATTTGTCCACAATGATCAACTACGCCCAGCCTGTAAAGTTTCAAGGTTTCCATGTGGCAGAAGAACGCAATATTCATTATAACATGTCTTCTTTTAATGAATCAGTCGGTCTTGGCTACTTGAAGACACATGCAATTGAATTTGTCAATTATAACAAACGGCAAATGAGTCGCATTTACCCCAAGGGAGGCCGAGTCGATTCCAGTAATTACATGCCTCAGATTTTCTGGAACGCTGGCTGCCAGATGGTTTCACTGAACTATCAAACCCCAGATTTAGCGATGCAATTGAATCAGGGAAAATTTGAGTATAATGGATCGTGCGGGTACCTTCTCAAACCAGATTTCATGAGGCGGCCTGATCGAACATTTGACCCCTTCTCTGAAACTCCTGTTGATGGTGTTATTGCAGCCACTTGCTCAGTGCAGGTTATATCAGGTCAATTCTTATCAGATAAGAAAATTGGCACCTACGTAGAGGTGGATATGTATGGGTTGCCCACTGACACCATACGTAAGGAATTCCGAACTCGCATGGTTATGAATAATGGACTCAATCCAGTTTACAATGAAGAGTCATTTGTATTTCGGAAGGTGATCCTGCCGGACCTGGCTGTCTTGAGAATAGCTGTGTATGATGATAACAACAAGCTGATTGGCCAGAGGATCCTCCCGCTTGATGGCCTCCAAGCCGGATATCGACACATTTCCCTTCGAAATGAGGGAAATAAACCATTATCACTACCAACAATTTTCTGCAATATTGTTCTTAAAACATATGTGCCTGATGGATTTGGAGATATCGTGGATGCTTTATCAGATCCAAAGAAATTTCTCTCAATTACAGAAAAGAGAGCAGACCAAATGAGAGCTATGGGCATTGAAACTAGTGACATAGCCGACGTGCCCAGTGACACTTCCAAAAATGACAAGAAAGGAAAGGCCAACACCGCCAAAGCAAATGTGACCCCTCAGAGTAGCTCTGAGCTCAGACCAACCACCACGGCTGCCCTGGCCTCTGGTGTGGAAGCCAAGAAAGGTATTGAACTTATCCCTCAAGTAAGGATAGAAGACTTAAAGCAGATGAAGGCTTACTTGAAGCATTTAAAGAAACAGCAGAAGGAGCTAAATTCTTTAAAGAAGAAACATGCAAAGGAACACAGTACCATGCAGAAGTTACACTGCACGCAAGTTGACAAAATTGTGGCACAGTATGACAAAGAGAAGTCGACTCATGAGAAAATCCTAGAGAAGGCAATGAAGAAGAAGGGGGGAAGTAATTGTCTCGAAATGAAAAAAGAAACAGAAATCAAAATTCAGACGCTGACATCAGATCACAAATCTAAGGTCAAAGAGATTGTAGCACAGCACACAAAGGAATGGTCAGAAATGATCAATACCCACAGTGCTGAGGAGCAAGAAATCCGAGACCTGCACCTCAGCCAGCAGTGTGAGCTGCTGAAAAAGCTACTCATCAATGCCCACGAGCAGCAAACCCAGCAGCTGAAACTGTCCCATGACAGGGAAAGCAAGGAAATGCGAGCACACCAGGCTAAGATTTCTATGGAAAATAGCAAAGCCATCAGCCAAGATAAATCTATCAAGAATAAAGCAGAACGGGAAAGGCGAGTCAGGGAGTTAAACAGCAGCAACACTAAAAAGTTTCTGGAAGAAAGAAAGAGACTTGCCATGAAGCAGTCCAAAGAAATGGATCAGTTGAAAAAAGTCCAGCTTGAACATCTAGAATTCCTAGAGAAACAGAATGAGCAGCTTTTGAAATCCTGTCATGCAGTGTCCCAAACGCAAGGCGAAGGAGATGCAGCAGATGGTGAAATTGGAAGCCGAGATGGACCGCAGACCAGCAACAGTAGTATGAAACTCCAAAATGCAAACTGA

LgBiT-linker-RhoA

ATGGTGTTTACTCTGGAGGACTTCGTCGGAGACTGGGAACAGACTGCTGCTTACAATCTGGATCAGGTGCTGGAACAGGGGGGGGTCAGCTCCCTGCTCCAGAACCTGGCCGTGTCTGTGACACCTATCCAGCGGATCGTGAGAAGCGGCGAGAATGCCCTGAAGATCGACATCCACGTGATCATCCCATACGAGGGCCTGTCCGCCGATCAGATGGCCCAGATCGAGGAGGTGTTCAAGGTGGTGTACCCAGTGGACGATCACCACTTCAAAGTGATCCTGCCCTATGGCACCCTGGTCATCGACGGAGTGACCCCAAACATGCTGAATTACTTCGGCAGGCCTTATGAGGGCATCGCCGTGTTTGATGGCAAGAAGATCACCGTGACAGGCACCCTGTGGAACGGCAATAAGATCATCGACGAGCGGCTGATCACCCCCGATGGCTCTATGCTGTTCAGAGTGACCATCAATAGCGGGGGGAGCGGCGGGGGGGGGAGCGGGGGAAGCAGTAGTGGCGGTACCGCAGCAATCCGGAAGAAGCTGGTCATCGTGGGCGACGGAGCATGCGGCAAGACCTGTCTGCTGATCGTGTTCAGCAAGGATCAGTTTCCCGAGGTGTACGTGCCTACAGTGTTCGAGAACTATGTGGCCGACATCGAGGTGGATGGCAAGCAGGTGGAGCTGGCCCTGTGGGACACCGCCGGCCAGGAGGACTACGATAGGCTGCGCCCTCTGTCCTATCCAGACACAGATGTGATCCTGATGTGCTTCAGCATCGACTCCCCTGATTCTCTGGAGAACATCCCAGAGAAGTGGACCCCCGAGGTGAAGCACTTTTGTCCCAATGTGCCTATCATCCTGGTGGGCAACAAGAAGGACCTGAGGAATGATGAGCACACACGGAGAGAGCTGGCCAAGATGAAGCAGGAGCCAGTGAAGCCAGAGGAGGGAAGGGACATGGCAAATAGAATCGGCGCCTTCGGCTACATGGAGTGCTCTGCCAAGACCAAGGATGGCGTGCGCGAGGTGTTTGAGATGGCAACAAGAGCCGCCCTGCAAGCAAGGAGAGGAAAGAAGAAAAGTGGATGTCTGGTGCTGTGA

SmBiT-linker-PKN1-GBD

ATGGTCACTGGATATAGGCTGTTCGAGGAGATTCTGGGGGGCTCAGGAGGAGGAGGCTCAGGAGGCTCATCATCAGGGGGGGAAAGCGAGCCCAGGTCTTGGAGCCTGCTGGAGCAGCTGGGCCTGGCAGGAGCAGACCTGGCAGCACCTGGCGTGCAGCAGCAGCTGGAGCTGGAGAGGGAGCGCCTGAGGAGAGAGATCCGGAAGGAGCTGAAGCTGAAGGAGGGAGCAGAGAACCTGAGGAGGGCAACCACAGACCTGGGCCGGAGCCTGGGACCAGTGGAGCTGCTGCTGAGAGGCAGCTCCCGGAGACTGGATCTGCTGCACCAGCAGCTGCAAGAACTGCACGCACACGTCGTCCTGCCAGACCCCGCCGCAACCTGA

SmBiT-linker-p115-RhoGEF

ATGGTCACCGGCTATCGGCTGTTTGAAGAGATTCTGGGGGGGTCAGGAGGAGGAGGCTCAGGGGGCTCATCATCAGGAGGAGAAGACTTCGCCCGAGGGGCGGCCTCCCCAGGCCCCTCCCGGCCTGGCCTGGTTCCCGTCAGCATCATCGGGGCTGAGGATGAGGATTTTGAGAACGAGCTGGAGACAAACTCAGAAGAGCAAAACAGCCAGTTCCAGAGCCTGGAGCAGGTGAAGCGGCGCCCAGCCCACCTCATGGCCCTCCTGCAGCACGTGGCCCTGCAGTTTGAGCCAGGACCCCTGCTTTGCTGTCTGCATGCCGACATGCTGGGCTCACTGGGCCCCAAGGAGGCCAAGAAGGCCTTCCTGGACTTCTACCACAGCTTCCTGGAGAAGACAGCGGTTCTCCGGGTGCCGGTCCCTCCCAACGTCGCCTTTGAACTTGACCGCACTAGGGCTGACCTCATCTCCGAGGATGTCCAGCGGCGGTTCGTGCAGGAGGTGGTGCAAAGCCAGCAGGTAGCCGTGGGCCGGCAGCTGGAGGACTTCCGTTCCAAGCGGCTCATGGGCATGACGCCCTGGGAGCAGGAGCTGGCCCAGCTGGAGGCTTGGGTTGGGCGGGACCGAGCCAGCTACGAGGCCCGGGAGCGGCACGTGGCGGAGCGGCTGCTCATGCACCTGGAGGAGATGCAACATACCATCTCTACCGACGAAGAAAAGAGTGCTGCCGTGGTCAACGCCATTGGCCTGTACATGCGCCACCTTGGGGTGCGGACCAAGAGTGGAGACAAGAAGTCGGGGAGGAACTTCTTCCGGAAAAAGGTGATGGGGAACCGGCGGTCGGACGAGCCTGCCAAGACCAAGAAGGGGCTGAGCAGCATCCTGGATGCCGCCCGCTGGAACCGGGGAGAGCCCCAGGTTCCAGATTTTCGACACCTCAAAGCAGAGGTTGATGCCGAGAAGCCAGGTGCTACAGACCGGAAGGGAGGCGTGGGGATGCCCTCTCGGGACCGGAATATCGGGGCTCCTGGGCAGGACACCCCTGGAGTCTCTCTGCACCCTCTGTCCCTGGACAGCCCAGACCGGGAACCAGGTGCTGACGCCCCCCTGGAGCTGGGGGACTCATCCCCGCAGGGCCCAATGAGCCTGGAGTCCTTGGCGCCCCCAGAGAGTACCGACGAGGGGGCCGAAACCGAGAGCCCCGAGCCTGGAGATGAGGGGGAGCCGGGGCGGTCGGGACTGGAGCTTGAACCAGAAGAGCCTCCCGGCTGGCGGGAACTCGTCCCCCCAGACACCCTGCACAGCCTGCCCAAGAGCCAGGTGAAGCGGCAGGAGGTCATCAGCGAGCTGCTGGTGACAGAGGCGGCCCACGTGCGCATGCTGCGGGTGCTGCACGACCTCTTCTTCCAGCCCATGGCAGAATGCCTGTTCTTCCCCTTGGAGGAGCTGCAGAACATCTTCCCCAGCCTGGACGAGCTCATCGAGGTGCATTCCCTGTTCCTCGATCGCCTGATGAAGCGGAGGCAGGAGAGTGGCTACCTCATCGAGGAGATCGGAGACGTGCTGCTGGCCCGGTTTGATGGTGCTGAGGGCTCCTGGTTCCAGAAAATCTCCTCCCGCTTCTGCAGCCGCCAGTCATTTGCCTTAGAGCAGCTCAAAGCCAAGCAACGCAAGGACCCTCGGTTCTGTGCCTTCGTGCAGGAAGCTGAGAGCCGCCCGCGGTGCCGCCGCCTGCAGCTGAAGGACATGATCCCCACGGAGATGCAGCGGCTGACCAAGTACCCCCTGCTCCTGCAGAGCATCGGGCAGAACACAGAAGAGCCCACAGAACGGGAGAAAGTGGAGCTGGCAGCCGAGTGCTGCCGGGAAATTCTACACCACGTCAACCAAGCCGTGCGTGACATGGAGGACCTGCTGAGGCTCAAGGACTATCAGCGGCGCCTGGACTTGTCCCACCTTCGGCAGAGCAGCGACCCTATGCTGAGCGAGTTCAAGAACCTGGACATCACCAAGAAGAAATTGGTCCACGAGGGCCCACTGACGTGGCGGGTGACTAAGGACAAGGCAGTGGAGGTGCATGTGCTGCTGCTGGACGACCTGCTGCTGCTGCTCCAGCGCCAGGACGAGCGGCTGCTGCTCAAGTCCCATAGCCGGACACTGACGCCCACGCCCGATGGCAAGACCATGCTGCGGCCCGTGCTGCGGCTCACCTCCGCCATGACCCGCGAGGTGGCCACCGATCACAAAGCCTTCTACGTCCTTTTTACCTGGGACCAGGAGGCCCAGATATACGAGCTGGTGGCACAGACTGTGTCGGAGCGGAAAAACTGGTGTGCTCTCATCACTGAGACTGCCGGATCCCTGAAAGTCCCTGCCCCTGCCTCTCGCCCTAAGCCCCGGCCCAGCCCGAGCAGCACCCGAGAACCCCTCCTCAGCAGCTCTGAGAACGGCAATGGTGGCCGAGAGACGTCTCCAGCTGATGCCCGGACCGAGAGAATCCTCAGTGACCTCCTGCCCTTCTGCAGACCAGGCCCCGAGGGCCAGCTCGCTGCCACGGCCCTTCGGAAAGTGCTGTCCCTGAAGCAGCTTCTGTTTCCGGCGGAGGAAGACAATGGGGCGGGGCCTCCTCGAGATGGGGATGGGGTCCCAGGGGGCGGCCCCCTGAGCCCAGCACGGACCCAGGAAATCCAGGAGAACCTGCTCAGCTTGGAGGAGACCATGAAGCAGCTGGAGGAGTTGGAGGAGGAATTTTGCCGCCTGAGACCCCTCCTGTCTCAGCTTGGGGGGAACTCTGTCCCCCAGCCTGGCTGCACTTGA

SmBiT-linker-PDZ-RhoGEF

ATGGTCACCGGCTATCGGCTGTTTGAAGAGATTCTGGGGGGGTCAGGAGGAGGAGGCTCAGGGGGCTCATCATCAGGAGGAAGTGTAAGGTTACCCCAGAGTATAGACAGGTTAAGTAGCCTGTCTTCTCTGGGAGATTCTGCACCAGAGCGCAAGTCCCCTTCCCACCATCGCCAGCCTTCGGATGCCTCTGAGACAACAGGTCTCGTTCAACGCTGTGTCATTATCCAAAAGGACCAGCATGGCTTCGGCTTCACAGTCAGTGGGGATCGCATTGTTCTGGTGCAGTCTGTGCGGCCTGGAGGTGCAGCCATGAAGGCCGGTGTGAAAGAGGGCGACCGGATCATCAAAGTCAACGGCACCATGGTGACCAATAGCTCACACCTGGAAGTGGTAAAGCTGATCAAATCTGGCGCCTATGTCGCACTCACCCTCCTGGGCTCTTCACCTTCATCCATGGGCATCTCTGGGCTCCAGCAGGACCCATCCCCAGCAGGAGCTCCCCGAATCACGTCAGTGATCCCCTCACCACCACCTCCTCCACCTCTACCACCTCCACAACGCATCACAGGACCCAAACCTCTGCAGGATCCCGAAGTTCAAAAACATGCCACCCAGATCCTCAGGAATATGCTGAGGCAGGAAGAAAAAGAATTACAGGACATACTTCCACTATATGGTGACACCAGCCAGAGACCATCAGAAGGCCGGCTCTCTCTGGATTCCCAGGAGGGGGACAGTGGCTTGGACTCTGGGACAGAACGCTTTCCTTCCCTCAGTGAGTCATTGATGAATCGGAACTCGGTACTGTCAGACCCTGGGCTAGACAGTCCTCGAACCTCCCCTGTGATCATGGCCAGGGTGGCCCAGCACCACAGGCGGCAGGGCTCGGATGCAGCAGTCCCCTCAACCGGTGACCAGGGTGTAGATCAAAGCCCAAAGCCTTTAATTATTGGCCCAGAGGAAGACTATGACCCGGGTTATTTCAACAACGAGAGCGACATCATATTCCAGGATCTGGAGAAACTGAAGTCTCGGCCAGCTCACCTGGGGGTTTTTCTACGTTACATCTTCTCTCAGGCGGACCCCAGTCCACTGCTTTTTTACCTGTGTGCAGAAGTTTATCAGCAGGCAAGCCCCAAGGATTCCCGAAGCTTGGGGAAAGACATCTGGAATATTTTCCTGGAGAAAAATGCGCCTCTGAGAGTGAAGATCCCTGAGATGCTACAGGCTGAAATTGACTCGCGCCTGCGGAACAGCGAAGATGCCCGTGGTGTTCTCTGTGAAGCTCAAGAGGCAGCCATGCCTGAGATCCAAGAGCAGATCCACGACTACAGAACGAAGCGCACACTGGGGCTGGGCAGCCTGTATGGTGAAAATGACCTGCTGGACCTGGATGGGGACCCTCTCCGAGAGCGCCAAGTGGCTGAGAAGCAGCTGGCTGCCCTTGGAGATATTTTGTCCAAGTATGAGGAAGACAGGAGCGCCCCCATGGACTTCGCCCTCAATACCTACATGAGCCATGCTGGGATCCGTCTTCGAGAGGCACGACCTTCCAACACAGCTGAAAAGGCCCAGTCTGCTCCTGACAAGGACAAGTGGCTACCGTTCTTCCCTAAGACCAAGAAGAGCAGCAATTCCAAGAAAGAAAAGGATGCCTTGGAGGACAAGAAGCGAAACCCTATCCTCAAATACATTGGGAAGCCCAAAAGCTCTTCTCAAAGCACATTTCATATTCCCTTGTCCCCTGTGGAAGTCAAACCAGGCAATGTGAGGAACATCATTCAGCACTTTGAGAACAACCAGCAGTATGATGCCCCAGAACCTGGGACACAACGACTCTCGACCGGAAGCTTTCCTGAGGACCTGCTGGAGAGTGACAGTTCACGCTCAGAGATTCGCCTGGGCCGCTCTGAAAGCCTCAAGGGCCGGGAAGAGATGAAACGGTCTCGAAAGGCAGAGAACGTGCCCCGCTCTCGCAGTGATGTTGACATGGATGCTGCTGCGGAGGCTACTCGCCTGCACCAGTCAGCCTCGTCCTCTACCTCCAGCCTCTCCACCAGGTCTCTTGAGAACCCAACCCCTCCATTCACTCCCAAAATGGGCCGCAGGAGCATTGAGTCCCCCAGTTTGGGGTTCTGCACAGATACCCTCCTTCCCCACCTCCTAGAGGATGATCTGGGCCAGCTGTCTGACCTGGAGCCAGAGCCAGATGCCCAAAATTGGCAGCATACAGTGGGCAAGGATGTGGTGGCTGGGCTAACCCAGCGGGAGATTGACCGGCAAGAGGTCATCAATGAGCTGTTTGTGACTGAAGCTTCCCACCTGCGCACACTCCGGGTCCTGGACCTGATCTTCTACCAGCGAATGAAGAAGGAGAACCTGATGCCCCGGGAGGAGCTGGCCCGGCTCTTCCCGAACCTGCCTGAACTCATAGAGATTCACAATTCCTGGTGTGAAGCCATGAAGAAGCTCCGGGAGGAAGGCCCCATCATCAAAGAGATCAGTGACCTCATGCTGGCCCGGTTTGATGGCCCTGCCCGAGAGGAACTCCAGCAAGTGGCTGCACAGTTCTGTTCCTATCAGTCAATAGCCCTAGAGCTAATCAAGACCAAGCAACGCAAGGAGAGTCGATTCCAGCTCTTCATGCAGGAGGCTGAGAGCCACCCTCAGTGTCGGCGGCTGCAGCTGAGAGACCTCATCATCTCTGAGATGCAGCGGCTCACCAAGTACCCGCTGCTGCTGGAGAGCATCATCAAGCACACAGAGGGTGGCACCTCTGAGCATGAGAAGCTGTGCCGGGCCCGGGACCAGTGCCGGGAGATTCTCAAGTATGTGAATGAAGCGGTAAAACAAACAGAGAACCGCCACCGTTTAGAGGGCTACCAGAAACGCCTGGATGCCACCGCCCTGGAGAGGGCCAGCAACCCCCTGGCAGCAGAGTTCAAGAGCCTGGATCTTACAACCAGAAAAATGATCCATGAGGGACCCCTGACCTGGAGGATCAGCAAGGATAAGACCTTGGACCTCCACGTGCTGCTGCTGGAGGACCTCCTAGTGCTGCTACAGAAACAGGATGAGAAGCTATTGCTGAAGTGCCACAGCAAGACTGCTGTGGGCTCCTCAGACAGCAAGCAGACCTTCAGCCCCGTGCTCAAGCTCAATGCTGTGCTCATCCGCTCTGTGGCCACAGATAAACGGGCCTTCTTCATCATCTGCACCTCCAAGCTGGGCCCACCCCAGATCTATGAGCTGGTTGCATTGACGTCATCAGACAAGAACACATGGATGGAGCTCTTAGAAGAGGCCGTGCGGAATGCCACCAGGCACCCCGGAGCTGCCCCAATGCCCGTCCATCCTCCACCCCCAGGTCCCCGGGAGCCAGCCCAGCAGGGCCCCACACCCAGCAGGGTAGAACTGGATGACTCAGACGTGTTCCATGGTGAACCTGAACCTGAGGAGCTGCCTGGAGGCACTGGGTCCCAGCAGAGGGTCCAAGGGAAGCACCAGGTCCTGCTAGAGGACCCTGAGCAGGAGGGCAGTGCAGAGGAAGAGGAACTGGGTGTCCTGCCTTGCCCTTCCACATCCCTGGATGGAGAGAACAGGGGCATCAGGACAAGGAACCCCATCCACTTGGCCTTCCCAGGCCCTCTGTTCATGGAAGGGCTCGCTGACTCCGCTCTGGAAGATGTGGAGAACCTGCGACATCTGATCCTGTGGAGCCTGCTGCCAGGTCACACCATGGAAACTCAGGCTGCCCAGGAGCCCGAGGACGACCTGACACCCACACCTTCTGTCATCAGCGTCACCTCTCACCCCTGGGACCCAGGCTCCCCAGGGCAAGCACCCCCTGGGGGTGAAGGGGACAACACCCAGCTTGCAGGGCTGGAGGGGGAACGGCCAGAGCAGGAAGACATGGGTCTCTGTTCTCTGGAACACCTACCCCCAAGGACCAGAAATTCTGGGATATGGGAGTCTCCAGAACTGGACAGGAATCTGGCTGAAGATGCTTCAAGCACAGAGGCAGCAGGAGGTTACAAAGTTGTGAGAAAAGCTGAGGTGGCAGGCAGCAAGGTTGTCCCTGCACTACCAGAGAGTGGCCAGTCAGAGCCTGGGCCACCTGAAGTGGAAGGCGGAACAAAGGCTACGGGGAACTGCTTTTATGTCAGCATGCCATCAGGACCCCCGGACTCAAGCACCGACCACTCAGAGGCACCCATGAGCCCCCCTCAGCCTGACAGCCTCCCTGCAGGGCAGACAGAGCCTCAGCCTCAGCTGCAGGGAGGCAACGATGATCCAAGACGCCCCAGCCGCTCTCCTCCAAGCCTGGCCCTCAGGGACGTGGGCATGATCTTCCATACCATTGAGCAGCTCACTCTCAAGCTCAACAGGCTCAAGGATATGGAGCTGGCCCACAGAGAGCTGCTCAAGTCCCTTGGGGGAGAGTCATCTGGTGGCACCACGCCTGTGGGCAGTTTCCACACAGAAGCAGCTAGATGGACAGATGGCTCCCTCTCACCTCCCGCTAAGGAGCCCCTAGCTTCTGACTCCAGGAACAGCCATGAACTGGGGCCCTGCCCTGAGGATGGCTCTGACGCCCCCCTGGAAGACAGCACAGCAGACGCAGCCGCGTCACCAGGACCATGA

SmBiT-linker-LARG

ATGGTCACCGGCTATCGGCTGTTTGAAGAGATTCTGGGGGGGTCAGGAGGAGGAGGCTCAGGGGGCTCATCATCAGGAGGAAGTGGCACACAGTCTACTATCACCGACAGGTTTCCCCTCAAAAAACCTATAAGGCATGGAAGTATTTTGAACCGAGAGTCACCAACAGATAAGAAGCAGAAAGTTGAGCGCATTGCATCACATGATTTTGACCCCACAGATAGCTCCTCCAAGAAGACAAAGTCTAGTTCAGAGGAGAGTAGATCCGAGATATATGGTCTTGTTCAGCGTTGCGTAATCATCCAGAAAGATGACAATGGATTTGGGCTGACGGTCAGTGGAGACAATCCAGTCTTCGTACAGTCTGTCAAAGAAGATGGAGCAGCCATGCGGGCTGGAGTACAGACAGGTGATCGAATCATCAAGGTGAATGGAACTCTGGTGACTCATTCAAATCATCTGGAGGTGGTGAAGCTAATCAAATCTGGTTCCTATGTAGCTCTCACTGTTCAGGGACGCCCACCTGGGTCGCCCCAGATTCCACTTGCCGACTCTGAAGTAGAGCCGTCAGTCATTGGACATATGTCTCCCATCATGACATCTCCTCATTCACCTGGAGCATCTGGGAATATGGAGAGAATCACTAGTCCTGTGCTCATGGGGGAGGAAAACAATGTGGTTCATAACCAGAAAGTAGAAATTCTGAGAAAAATGTTACAGAAAGAACAGGAACGGCTACAGTTATTGCAGGAAGATTACAACCGAACACCTGCCCAAAGATTGCTAAAAGAGATCCAAGAGGCCAAGAAACACATTCCTCAGCTGCAAGAGCAGTTATCCAAAGCCACAGGCTCTGCTCAGGATGGAGCTGTAGTTACACCCTCCAGACCTTTAGGGGACACCCTAACAGTCAGTGAGGCAGAAACAGATCCTGGAGATGTACTGGGCAGGACTGACTGTAGCAGTGGAGATGCTTCTCGGCCCAGTAGTGACAATGCAGATAGTCCCAAGAGTGGCCCAAAAGAGAGAATTTATCTAGAGGAAAACCCAGAGAAAAGTGAAACAATTCAGGACACTGACACTCAATCACTTGTCGGAAGTCCCTCAACCCGTATAGCACCTCATATTATTGGAGCAGAAGATGATGATTTTGGTACTGAACATGAACAGATCAATGGACAGTGCAGCTGTTTCCAGAGCATTGAATTACTAAAATCTCGCCCGGCTCATTTGGCTGTTTTCTTACACCATGTAGTTTCACAATTTGACCCTGCGACTTTGCTCTGTTATCTCTATTCAGACCTGTATAAACATACCAATTCCAAAGAAACTCGTCGCATCTTCCTTGAGTTTCATCAGTTCTTTCTAGATCGATCAGCACACCTGAAAGTTTCTGTTCCTGATGAAATGTCTGCAGATCTAGAAAAGAGAAGACCTGAGCTCATTCCTGAGGATCTGCATCGCCACTATATCCAAACTATGCAAGAAAGAGTCCATCCAGAAGTTCAAAGGCACTTAGAAGATTTTCGGCAGAAACGTAGTATGGGACTGACCTTGGCTGAAAGCGAGCTGACTAAACTTGATGCAGAGCGAGACAAGGACCGATTGACTTTGGAGAAGGAGCGGACATGTGCAGAACAGATTGTTGCCAAAATTGAAGAAGTATTGATGACTGCTCAGGCTGTAGAGGAAGATAAGAGCTCCACCATGCAGTATGTTATTCTCATGTATATGAAGCATTTGGGAGTAAAAGTGAAAGAGCCTCGAAATTTGGAGCACAAACGGGGTCGGATTGGATTTCTTCCCAAAATCAAGCAAAGTATGAAGAAAGATAAAGAAGGGGAAGAAAAAGGGAAGCGAAGAGGATTCCCCAGCATCCTGGGACCCCCACGGAGACCAAGCCGTCATGACAACAGTGCAATTGGCAGAGCCATGGAACTACAGAAGGCGCGCCACCCTAAGCACTTATCCACACCCTCATCTGTGAGTCCTGAACCTCAGGACTCTGCCAAGTTGCGCCAGAGTGGGTTAGCAAATGAAGGAACAGACGCTGGATACCTGCCTGCCAATTCCATGTCTTCTGTAGCTTCAGGGGCCTCTTTTTCCCAGGAAGGAGGGAAAGAGAATGATACAGGATCAAAGCAAGTTGGAGAAACATCAGCACCTGGAGACACCTTAGATGGCACACCTCGTACTCTCAATACTGTCTTTGATTTCCCACCACCTCCATTAGACCAAGTGCAGGAGGAGGAATGTGAAGTAGAAAGGGTGACTGAACATGGGACACCAAAGCCCTTTCGAAAGTTTGACAGTGTAGCTTTTGGAGAAAGTCAAAGTGAGGATGAACAATTTGAAAATGACTTAGAGACAGATCCACCCAACTGGCAGCAGCTTGTTAGTCGAGAAGTGTTACTGGGACTAAAACCTTGTGAAATCAAAAGACAGGAAGTGATTAATGAATTGTTCTACACTGAAAGAGCTCATGTTCGAACACTGAAGGTTCTTGATCAAGTGTTCTATCAGCGAGTATCCAGAGAAGGAATTCTGTCACCCTCAGAGCTACGGAAAATTTTTTCAAACTTGGAAGATATTCTTCAACTTCATATTGGATTGAATGAACAAATGAAGGCTGTTCGAAAGAGAAATGAGACCTCTGTTATCGATCAGATTGGGGAAGATTTGCTGACATGGTTCAGCGGACCAGGAGAGGAGAAATTGAAACATGCTGCTGCTACCTTTTGCAGTAACCAACCTTTCGCCCTGGAAATGATCAAATCTCGTCAGAAAAAGGATTCTCGATTTCAGACTTTTGTGCAAGATGCTGAAAGTAATCCACTGTGTCGTCGTCTTCAACTGAAGGATATTATTCCCACTCAAATGCAAAGGCTTACTAAGTACCCACTTCTGTTGGATAATATTGCCAAATACACAGAATGGCCAACAGAAAGGGAGAAGGTGAAGAAAGCTGCAGATCACTGTCGTCAGATCTTAAATTATGTAAATCAGGCTGTCAAGGAGGCAGAAAACAAGCAGCGCCTAGAAGATTATCAGCGTCGCCTTGATACCTCCAGCCTGAAGTTGTCAGAGTACCCAAATGTTGAAGAGCTCAGGAATTTGGATTTAACAAAAAGGAAGATGATTCATGAAGGGCCATTGGTTTGGAAGGTGAATAGAGATAAAACTATTGATTTATACACGTTGCTGCTGGAAGACATTCTTGTATTGTTACAAAAGCAGGATGATAGACTGGTTTTAAGGTGTCATAGTAAGATTCTGGCATCTACAGCTGATAGCAAACACACGTTTAGCCCTGTCATTAAGTTGAGTACAGTGTTGGTTCGACAAGTGGCAACAGATAACAAAGCTTTATTCGTCATTTCCATGTCAGACAATGGCGCTCAGATTTATGAACTGGTGGCACAGACAGTTTCTGAAAAGACTGTCTGGCAGGACCTAATCTGTCGGATGGCTGCATCAGTGAAGGAGCAATCCACAAAGCCAATTCCATTACCACAGTCAACACCTGGCGAAGGAGATAATGATGAAGAAGATCCTTCAAAATTAAAAGAGGAGCAGCATGGCATTTCAGTCACTGGTTTGCAGAGTCCAGACAGAGATTTGGGATTAGAATCTACCTTAATATCGTCAAAACCTCAGTCTCATTCACTGAGTACCTCTGGGAAATCAGAGGTACGTGATCTGTTTGTGGCTGAGAGACAGTTTGCAAAGGAACAACATACAGATGGGACACTAAAGGAAGTTGGAGAAGATTATCAAATCGCAATCCCAGATTCACACCTGCCTGTCTCAGAAGAACGGTGGGCATTGGATGCACTAAGAAATTTGGGTTTGTTGAAGCAGTTGCTGGTGCAACAGCTAGGTTTGACTGAGAAGAGCGTTCAGGAAGACTGGCAACATTTCCCAAGATACAGAACAGCCTCTCAGGGGCCGCAGACAGACAGTGTCATCCAGAACTCTGAAAATATTAAGGCCTATCATTCTGGTGAAGGACATATGCCCTTTAGAACTGGAACTGGTGACATTGCAACTTGTTACAGTCCACGGACTTCAACTGAATCTTTTGCTCCACGGGATTCAGTGGGACTGGCACCCCAGGATAGCCAGGCAAGTAACATTTTAGTAATGGACCACATGATTATGACCCCAGAGATGCCTACCATGGAGCCAGAAGGGGGTCTTGATGACAGTGGAGAGCACTTTTTTGATGCCCGTGAAGCACATAGTGATGAGAATCCATCAGAAGGTGATGGAGCAGTTAACAAGGAAGAGAAGGATGTTAATTTACGCATCTCAGGAAACTATTTGATCCTTGATGGCTATGACCCAGTGCAGGAGAGTTCCACAGATGAGGAGGTTGCTTCCTCACTTACCCTGCAGCCCATGACAGGCATCCCTGCTGTGGAATCCACCCACCAGCAGCAACATTCTCCTCAGAATACTCACTCCGATGGGGCAATTTCACCATTCACCCCCGAATTTCTGGTCCAGCAGCGCTGGGGAGCTATGGAGTATTCCTGTTTTGAGATCCAGAGTCCCTCCTCTTGTGCAGATTCACAGAGCCAGATCATGGAGTACATTCATAAGATAGAGGCTGACCTTGAACACTTAAAGAAGGTGGAGGAAAGTTACACCATTCTTTGCCAAAGGCTGGCTGGATCAGCCCTCACAGACAAGCACTCAGATAAAAGTTGA

SmBiT-linker-p63-RhoGEF

ATGGTCACCGGCTATCGGCTGTTTGAAGAGATTCTGGGGGGGTCAGGAGGAGGAGGCTCAGGGGGCTCATCATCAGGAGGACGGGGGGGGCACAAAGGGGGTCGCTGTGCCTGTCCCCGTGTGATCCGAAAAGTGCTGGCAAAATGCGGCTGCTGCTTCGCCCGGGGGGGACGTGAATCCTATTCCATTGCGGGCAGTGAGGGGAGTATATCGGCTTCTGCTGCCTCCGGTCTGGCTGCCCCCTCTGGCCCCAGCTCTGGCCTCAGCTCTGGCCCCTGTTCCCCAGGCCCCCCAGGGCCCGTCAGTGGCCTGAGGAGATGGTTGGATCATTCCAAACATTGTCTCAGTGTGGAAACTGAGGCAGACAGTGGTCAGGCAGGACCATATGAGAACTGGATGTTGGAGCCAGCTCTAGCCACAGGAGAGGAGCTGCCGGAACTGACCTTGCTGACCACACTGTTGGAGGGCCCTGGAGATAAGACGCAGCCACCTGAAGAGGAGACTTTGTCCCAAGCCCCTGAGAGTGAGGAGGAACAGAAGAAGAAGGCTCTGGAAAGGAGTATGTATGTCCTGAGTGAACTGGTAGAAACAGAGAAAATGTACGTGGACGACTTGGGGCAGATTGTGGAGGGTTATATGGCCACCATGGCTGCTCAGGGGGTCCCCGAGAGTCTTCGAGGCCGTGACAGGATTGTGTTTGGGAATATCCAGCAAATCTATGAGTGGCACCGAGACTATTTCTTGCAAGAGCTACAACGGTGTCTGAAAGATCCTGATTGGCTGGCTCAGCTATTCATCAAACACGAGCGCCGGCTGCATATGTATGTGGTGTACTGTCAGAATAAGCCCAAGTCAGAGCATGTGGTGTCAGAGTTTGGGGACAGCTACTTTGAGGAGCTCCGGCAGCAGCTGGGGCACCGCCTGCAGCTGAACGACCTCCTCATCAAACCTGTGCAGCGGATCATGAAATACCAGCTGCTGCTCAAGGATTTTCTCAAGTATTACAATAGAGCTGGGATGGATACTGCAGACCTAGAGCAAGCTGTGGAGGTCATGTGCTTTGTGCCCAAGCGCTGCAACGATATGATGACGCTGGGGAGATTGCGGGGATTTGAGGGCAAACTGACTGCTCAGGGGAAGCTCTTGGGCCAGGACACTTTCTGGGTCACCGAGCCTGAGGCTGGAGGGCTGCTGTCTTCCCGAGGTCGAGAGAGGCGCGTCTTCCTCTTTGAGCAAATCATCATCTTCAGTGAAGCCCTGGGAGGAGGAGTGAGAGGTGGAACACAGCCTGGATATGTATACAAGAACAGCATTAAGGTGAGCTGCCTGGGACTGGAGGGGAACCTCCAAGGTGACCCTTGCCGCTTTGCACTGACCTCCAGAGGGCCAGAGGGTGGGATCCAGCGCTATGTCCTGCAGGCTGCAGACCCTGCTATCAGTCAGGCCTGGATCAAGCATGTGGCTCAGATCTTGGAGAGCCAACGGGACTTCCTCAACGCATTGCAGTCACCCATTGAGTACCAGAGACGGGAGAGCCAGACCAACAGCCTGGGGCGGCCAAGAGGGCCTGGAGTGGGGAGCCCTGGAAGAATTCAGCTTGGAGATCAGGCCCAGGGCAGCACACACACACCCATCAATGGCTCTCTCCCCTCTCTGCTGCTGTCACCCAAAGGGGAGGTGGCCAGAGCCCTCTTGCCACTGGATAAACAGGCCCTTGGTGACATCCCCCAGGCTCCCCATGACTCTCCTCCAGTCTCTCCAACTCCAAAAACCCCTCCCTGCCAAGCCAGACTTGCCAAGCTGGATGAAGATGAGCTGTGA

LgBiT-linker-Ib4

ATGGTGTTTACTCTGGAGGACTTCGTCGGAGACTGGGAACAGACTGCTGCTTACAATCTGGATCAGGTGCTGGAACAGGGGGGGGTCAGCTCCCTGCTCCAGAACCTGGCCGTGTCTGTGACACCTATCCAGCGGATCGTGAGAAGCGGCGAGAATGCCCTGAAGATCGACATCCACGTGATCATCCCATACGAGGGCCTGTCCGCCGATCAGATGGCCCAGATCGAGGAGGTGTTCAAGGTGGTGTACCCAGTGGACGATCACCACTTCAAAGTGATCCTGCCCTATGGCACCCTGGTCATCGACGGAGTGACCCCAAACATGCTGAATTACTTCGGCAGGCCTTATGAGGGCATCGCCGTGTTTGATGGCAAGAAGATCACCGTGACAGGCACCCTGTGGAACGGCAATAAGATCATCGACGAGCGGCTGATCACCCCCGATGGCTCTATGCTGTTCAGAGTGACCATCAATAGCGGGGGGAGCGGCGGGGGGGGGAGCGGGGGAAGCAGTAGTGGCGGTGGCAGTGATATTCAGATGACCCAGAGTCCTTCTAGCCTGTCCGCTTCCGTCGGGGACCGCGTGACTATTACCTGTCGGGCTTCACAGAGCGTGAGCAGCGCCGTGGCCTGGTATCAGCAGAAGCCCGGCAAGGCCCCTAAGCTGCTGATCTACAGCGCCTCTAGCCTGTATAGCGGCGTGCCATCCCGGTTCTCCGGCAGCCGGAGCGGCACCGACTTTACCCTGACAATCTCCTCTCTCCAGCCCGAGGATTTCGCCACATACTATTGCCAGCAGTACTATTCCCGGCTGATCACCTTTGGCCAGGGCACAAAGGTGGAGATCAAGGGCACCACAGCCGCCTCTGGCTCTGAGTTCAGCGGCGGCAGCTCCTCTGGAGCAGAGGTGCAGCTCGTGGAAAGCGGCGGCGGCCTGGTGCAGCCTGGCGGCTCTCTGAGACTGAGCTGTGCAGCATCCGGCTTTAACGTGTACAGCTCCTATATCCACTGGGTGAGACAGGCACCAGGCAAGGGCCTGGAGTGGGTGGCCTCCATCTCTAGCTACTATGGCTACACCTACTATGCCGACTCTGTGAAGGGCAGGTTCACAATCTCTGCCGATACCAGCAAGAACACAGCCTACCTCCAGATGAATAGCCTGAGAGCAGAGGACACCGCCGTGTACTATTGCGCCAGGGAGCAGAACTGGCCCTACCGCTGGTATGCCCTGGACTACTGGGGACAGGGGACACTGGTGACCGTGAGTTCAGCA

LgBiT-linker-Ib30

ATGGTGTTTACTCTGGAGGACTTCGTCGGAGACTGGGAACAGACTGCTGCTTACAATCTGGATCAGGTGCTGGAACAGGGGGGGGTCAGCTCCCTGCTCCAGAACCTGGCCGTGTCTGTGACACCTATCCAGCGGATCGTGAGAAGCGGCGAGAATGCCCTGAAGATCGACATCCACGTGATCATCCCATACGAGGGCCTGTCCGCCGATCAGATGGCCCAGATCGAGGAGGTGTTCAAGGTGGTGTACCCAGTGGACGATCACCACTTCAAAGTGATCCTGCCCTATGGCACCCTGGTCATCGACGGAGTGACCCCAAACATGCTGAATTACTTCGGCAGGCCTTATGAGGGCATCGCCGTGTTTGATGGCAAGAAGATCACCGTGACAGGCACCCTGTGGAACGGCAATAAGATCATCGACGAGCGGCTGATCACCCCCGATGGCTCTATGCTGTTCAGAGTGACCATCAATAGCGGGGGGAGCGGCGGGGGGGGGAGCGGGGGAAGCAGTAGTGGCGGTGGATCAGATATTCAGATGACTCAGAGCCCTTCCTCACTGTCCGCCTCCGTCGGGGACCGCGTCACAATCACTTGCAGAGCCTCCCAGAGCGTGAGCAGCGCCGTGGCCTGGTATCAGCAGAAGCCCGGCAAGGCCCCTAAGCTGCTGATCTACTCCGCCTCTAGCCTGTATTCCGGCGTGCCCTCTAGGTTCTCCGGCTCTCGCAGCGGCACCGACTTTACCCTGACAATCTCCTCTCTCCAGCCAGAGGATTTCGCCACATACTATTGCCAGCAGTACAAGTATGTGCCCGTGACCTTTGGCCAGGGCACAAAGGTGGAGATCAAGGGCACCACAGCCGCCTCTGGCAGCTCCGGCGGCTCTAGCTCCGGAGCAGAGGTGCAGCTCGTGGAAAGCGGCGGCGGCCTGGTGCAGCCTGGCGGCAGCCTGCGGCTGTCCTGTGCCGCCTCTGGCTTCAACGTGTACTCTAGCTCCATCCACTGGGTGAGACAGGCACCAGGCAAGGGCCTGGAGTGGGTGGCCTCCATCTCTAGCTACTATGGCTACACCTACTATGCCGACTCTGTGAAGGGCAGGTTCACAATCAGCGCCGATACCTCCAAGAACACAGCCTACCTCCAGATGAATAGCCTGAGAGCCGAGGACACCGCCGTGTACTATTGTGCACGGAGCCGGCAGTTTTGGTATAGCGGCCTGGATTACTGGGGACAGGGAACTCTGGTGACCGTGAGCAGCGCATGA

LgBiT-linker-β-arrestin1

ATGGTGTTTACTCTGGAGGACTTCGTCGGAGACTGGGAACAGACTGCTGCTTACAATCTGGATCAGGTGCTGGAACAGGGGGGGGTCAGCTCCCTGCTCCAGAACCTGGCCGTGTCTGTGACACCTATCCAGCGGATCGTGAGAAGCGGCGAGAATGCCCTGAAGATCGACATCCACGTGATCATCCCATACGAGGGCCTGTCCGCCGATCAGATGGCCCAGATCGAGGAGGTGTTCAAGGTGGTGTACCCAGTGGACGATCACCACTTCAAAGTGATCCTGCCCTATGGCACCCTGGTCATCGACGGAGTGACCCCAAACATGCTGAATTACTTCGGCAGGCCTTATGAGGGCATCGCCGTGTTTGATGGCAAGAAGATCACCGTGACAGGCACCCTGTGGAACGGCAATAAGATCATCGACGAGCGGCTGATCACCCCCGATGGCTCTATGCTGTTCAGAGTGACCATCAATAGCGGGGGGAGCGGCGGGGGGGGGAGCGGGGGAAGCAGTAGTGGCGGTACCGGCGACAAAGGGACCCGAGTGTTCAAGAAGGCCAGTCCAAATGGAAAGCTCACCGTCTACCTGGGAAAGCGGGACTTTGTGGACCACATCGACCTCGTGGACCCTGTGGATGGTGTGGTCCTGGTGGATCCTGAGTATCTCAAAGAGCGGAGAGTCTATGTGACGCTGACCTGCGCCTTCCGCTATGGCCGGGAGGACCTGGATGTCCTGGGCCTGACCTTTCGCAAGGACCTGTTTGTGGCCAACGTACAGTCGTTCCCACCGGCCCCCGAGGACAAGAAGCCCCTGACGCGGCTGCAGGAACGCCTCATCAAGAAGCTGGGCGAGCACGCTTACCCTTTCACCTTTGAGATCCCTCCAAACCTTCCATGTTCTGTGACACTGCAGCCGGGGCCCGAAGACACGGGGAAGGCTTGCGGTGTGGACTATGAAGTCAAAGCCTTCTGCGCGGAGAATTTGGAGGAGAAGATCCACAAGCGGAATTCTGTGCGTCTGGTCATCCGGAAGGTTCAGTATGCCCCAGAGAGGCCTGGCCCCCAGCCCACAGCCGAGACCACCAGGCAGTTCCTCATGTCGGACAAGCCCTTGCACCTAGAAGCCTCTCTGGATAAGGAGATCTATTACCATGGAGAACCCATCAGCGTCAACGTCCACGTCACCAACAACACCAACAAGACGGTGAAGAAGATCAAGATCTCAGTGCGCCAGTATGCAGACATCTGCCTTTTCAACACAGCTCAGTACAAGTGCCCTGTTGCCATGGAAGAGGCTGATGACACTGTGGCACCCAGCTCGACGTTCTGCAAGGTCTACACACTGACCCCCTTCCTAGCCAATAACCGAGAGAAGCGGGGCCTCGCCTTGGACGGGAAGCTCAAGCACGAAGACACGAACTTGGCCTCTAGCACCCTGTTGAGGGAAGGTGCCAACCGTGAGATCCTGGGGATCATTGTTTCCTACAAAGTGAAAGTGAAGCTGGTGGTGTCTCGGGGCGGCCTGTTGGGAGATCTTGCATCCAGCGACGTGGCCGTGGAACTGCCCTTCACCCTAATGCACCCCAAGCCCAAAGAGGAACCCCCGCATCGGGAAGTTCCAGAGAACGAGACGCCAGTAGATACCAATCTCATAGAACTTGACACAAATGATGACGACATTGTATTTGAGGACTTTGCTCGCCAGAGACTGAAAGGCATGAAGGATGACAAGGAGGAAGAGGAGGATGGTACCGGCTCTCCACAGCTCAACAACAGATGA

LgBiT-linker-β-arrestin1 (ΔC382)

ATGGTGTTTACTCTGGAGGACTTCGTCGGAGACTGGGAACAGACTGCTGCTTACAATCTGGATCAGGTGCTGGAACAGGGGGGGGTCAGCTCCCTGCTCCAGAACCTGGCCGTGTCTGTGACACCTATCCAGCGGATCGTGAGAAGCGGCGAGAATGCCCTGAAGATCGACATCCACGTGATCATCCCATACGAGGGCCTGTCCGCCGATCAGATGGCCCAGATCGAGGAGGTGTTCAAGGTGGTGTACCCAGTGGACGATCACCACTTCAAAGTGATCCTGCCCTATGGCACCCTGGTCATCGACGGAGTGACCCCAAACATGCTGAATTACTTCGGCAGGCCTTATGAGGGCATCGCCGTGTTTGATGGCAAGAAGATCACCGTGACAGGCACCCTGTGGAACGGCAATAAGATCATCGACGAGCGGCTGATCACCCCCGATGGCTCTATGCTGTTCAGAGTGACCATCAATAGCGGGGGGAGCGGCGGGGGGGGGAGCGGGGGAAGCAGTAGTGGCGGTACCGGCGACAAAGGGACCCGAGTGTTCAAGAAGGCCAGTCCAAATGGAAAGCTCACCGTCTACCTGGGAAAGCGGGACTTTGTGGACCACATCGACCTCGTGGACCCTGTGGATGGTGTGGTCCTGGTGGATCCTGAGTATCTCAAAGAGCGGAGAGTCTATGTGACGCTGACCTGCGCCTTCCGCTATGGCCGGGAGGACCTGGATGTCCTGGGCCTGACCTTTCGCAAGGACCTGTTTGTGGCCAACGTACAGTCGTTCCCACCGGCCCCCGAGGACAAGAAGCCCCTGACGCGGCTGCAGGAACGCCTCATCAAGAAGCTGGGCGAGCACGCTTACCCTTTCACCTTTGAGATCCCTCCAAACCTTCCATGTTCTGTGACACTGCAGCCGGGGCCCGAAGACACGGGGAAGGCTTGCGGTGTGGACTATGAAGTCAAAGCCTTCTGCGCGGAGAATTTGGAGGAGAAGATCCACAAGCGGAATTCTGTGCGTCTGGTCATCCGGAAGGTTCAGTATGCCCCAGAGAGGCCTGGCCCCCAGCCCACAGCCGAGACCACCAGGCAGTTCCTCATGTCGGACAAGCCCTTGCACCTAGAAGCCTCTCTGGATAAGGAGATCTATTACCATGGAGAACCCATCAGCGTCAACGTCCACGTCACCAACAACACCAACAAGACGGTGAAGAAGATCAAGATCTCAGTGCGCCAGTATGCAGACATCTGCCTTTTCAACACAGCTCAGTACAAGTGCCCTGTTGCCATGGAAGAGGCTGATGACACTGTGGCACCCAGCTCGACGTTCTGCAAGGTCTACACACTGACCCCCTTCCTAGCCAATAACCGAGAGAAGCGGGGCCTCGCCTTGGACGGGAAGCTCAAGCACGAAGACACGAACTTGGCCTCTAGCACCCTGTTGAGGGAAGGTGCCAACCGTGAGATCCTGGGGATCATTGTTTCCTACAAAGTGAAAGTGAAGCTGGTGGTGTCTCGGGGCGGCCTGTTGGGAGATCTTGCATCCAGCGACGTGGCCGTGGAACTGCCCTTCACCCTAATGCACCCCAAGCCCAAAGAGGAACCCCCGCATCGGGAAGTTCCAGAGAACGAGACGCCAGTAGATACCAATCTCATAGAACTTGACACAAATTGA

LgBiT-linker-β-arrestin2

ATGGTGTTTACTCTGGAGGACTTCGTCGGAGACTGGGAACAGACTGCTGCTTACAATCTGGATCAGGTGCTGGAACAGGGGGGGGTCAGCTCCCTGCTCCAGAACCTGGCCGTGTCTGTGACACCTATCCAGCGGATCGTGAGAAGCGGCGAGAATGCCCTGAAGATCGACATCCACGTGATCATCCCATACGAGGGCCTGTCCGCCGATCAGATGGCCCAGATCGAGGAGGTGTTCAAGGTGGTGTACCCAGTGGACGATCACCACTTCAAAGTGATCCTGCCCTATGGCACCCTGGTCATCGACGGAGTGACCCCAAACATGCTGAATTACTTCGGCAGGCCTTATGAGGGCATCGCCGTGTTTGATGGCAAGAAGATCACCGTGACAGGCACCCTGTGGAACGGCAATAAGATCATCGACGAGCGGCTGATCACCCCCGATGGCTCTATGCTGTTCAGAGTGACCATCAATAGCGGGGGGAGCGGCGGGGGGGGGAGCGGGGGAAGCAGTAGTGGCGGTACCGGGGAGAAACCCGGGACCAGGGTCTTCAAGAAGTCGAGCCCTAACTGCAAGCTCACCGTGTACTTGGGCAAGCGGGACTTCGTAGATCACCTGGACAAAGTGGACCCTGTAGATGGCGTGGTGCTTGTGGACCCTGACTACCTGAAGGACCGCAAAGTGTTTGTGACCCTCACCTGCGCCTTCCGCTATGGCCGTGAAGACCTGGATGTGCTGGGCTTGTCCTTCCGCAAAGACCTGTTCATCGCCACCTACCAGGCCTTCCCCCCGGTGCCCAACCCACCCCGGCCCCCCACCCGCCTGCAGGACCGGCTGCTGAGGAAGCTGGGCCAGCATGCCCACCCCTTCTTCTTCACCATACCCCAGAATCTTCCATGCTCCGTCACACTGCAGCCAGGCCCAGAGGATACAGGAAAGGCCTGCGGCGTAGACTTTGAGATTCGAGCCTTCTGTGCTAAATCACTAGAAGAGAAAAGCCACAAAAGGAACTCTGTGCGGCTGGTGATCCGAAAGGTGCAGTTCGCCCCGGAGAAACCCGGCCCCCAGCCTTCAGCCGAAACCACACGCCACTTCCTCATGTCTGACCGGTCCCTGCACCTCGAGGCTTCCCTGGACAAGGAGCTGTACTACCATGGGGAGCCCCTCAATGTAAATGTCCACGTCACCAACAACTCCACCAAGACCGTCAAGAAGATCAAAGTCTCTGTGAGACAGTACGCCGACATCTGCCTCTTCAGCACCGCCCAGTACAAGTGTCCTGTGGCTCAACTCGAACAAGATGACCAGGTATCTCCCAGCTCCACATTCTGTAAGGTGTACACCATAACCCCACTGCTCAGCGACAACCGGGAGAAGCGGGGTCTCGCCCTGGATGGGAAACTCAAGCACGAGGACACCAACCTGGCTTCCAGCACCATCGTGAAGGAGGGTGCCAACAAGGAGGTGCTGGGAATCCTGGTGTCCTACAGGGTCAAGGTGAAGCTGGTGGTGTCTCGAGGCGGGGATGTCTCTGTGGAGCTGCCTTTTGTTCTTATGCACCCCAAGCCCCACGACCACATCCCCCTCCCCAGACCCCAGTCAGCCGCTCCGGAGACAGATGTCCCTGTGGACACCAACCTCATTGAATTTGATACCAACTATGCCACAGATGATGACATTGTGTTTGAGGACTTTGCCCGGCTTCGGCTGAAGGGGATGAAGGATGACGACTATGATGATCAACTCTGCTGA

LgBiT-linker-β-arrestin2 (ΔC382)

ATGGTGTTTACTCTGGAGGACTTCGTCGGAGACTGGGAACAGACTGCTGCTTACAATCTGGATCAGGTGCTGGAACAGGGGGGGGTCAGCTCCCTGCTCCAGAACCTGGCCGTGTCTGTGACACCTATCCAGCGGATCGTGAGAAGCGGCGAGAATGCCCTGAAGATCGACATCCACGTGATCATCCCATACGAGGGCCTGTCCGCCGATCAGATGGCCCAGATCGAGGAGGTGTTCAAGGTGGTGTACCCAGTGGACGATCACCACTTCAAAGTGATCCTGCCCTATGGCACCCTGGTCATCGACGGAGTGACCCCAAACATGCTGAATTACTTCGGCAGGCCTTATGAGGGCATCGCCGTGTTTGATGGCAAGAAGATCACCGTGACAGGCACCCTGTGGAACGGCAATAAGATCATCGACGAGCGGCTGATCACCCCCGATGGCTCTATGCTGTTCAGAGTGACCATCAATAGCGGGGGGAGCGGCGGGGGGGGGAGCGGGGGAAGCAGTAGTGGCGGTACCGGGGAGAAACCCGGGACCAGGGTCTTCAAGAAGTCGAGCCCTAACTGCAAGCTCACCGTGTACTTGGGCAAGCGGGACTTCGTAGATCACCTGGACAAAGTGGACCCTGTAGATGGCGTGGTGCTTGTGGACCCTGACTACCTGAAGGACCGCAAAGTGTTTGTGACCCTCACCTGCGCCTTCCGCTATGGCCGTGAAGACCTGGATGTGCTGGGCTTGTCCTTCCGCAAAGACCTGTTCATCGCCACCTACCAGGCCTTCCCCCCGGTGCCCAACCCACCCCGGCCCCCCACCCGCCTGCAGGACCGGCTGCTGAGGAAGCTGGGCCAGCATGCCCACCCCTTCTTCTTCACCATACCCCAGAATCTTCCATGCTCCGTCACACTGCAGCCAGGCCCAGAGGATACAGGAAAGGCCTGCGGCGTAGACTTTGAGATTCGAGCCTTCTGTGCTAAATCACTAGAAGAGAAAAGCCACAAAAGGAACTCTGTGCGGCTGGTGATCCGAAAGGTGCAGTTCGCCCCGGAGAAACCCGGCCCCCAGCCTTCAGCCGAAACCACACGCCACTTCCTCATGTCTGACCGGTCCCTGCACCTCGAGGCTTCCCTGGACAAGGAGCTGTACTACCATGGGGAGCCCCTCAATGTAAATGTCCACGTCACCAACAACTCCACCAAGACCGTCAAGAAGATCAAAGTCTCTGTGAGACAGTACGCCGACATCTGCCTCTTCAGCACCGCCCAGTACAAGTGTCCTGTGGCTCAACTCGAACAAGATGACCAGGTATCTCCCAGCTCCACATTCTGTAAGGTGTACACCATAACCCCACTGCTCAGCGACAACCGGGAGAAGCGGGGTCTCGCCCTGGATGGGAAACTCAAGCACGAGGACACCAACCTGGCTTCCAGCACCATCGTGAAGGAGGGTGCCAACAAGGAGGTGCTGGGAATCCTGGTGTCCTACAGGGTCAAGGTGAAGCTGGTGGTGTCTCGAGGCGGGGATGTCTCTGTGGAGCTGCCTTTTGTTCTTATGCACCCCAAGCCCCACGACCACATCCCCCTCCCCAGACCCCAGTCAGCCGCTCCGGAGACAGATGTCCCTGTGGACACCAACCTCATTGAATTTGATACCAACTATGCCACATGA

SmBiT-linker-β-arrestin1

ATGGTCACCGGCTATCGGCTGTTTGAAGAGATTCTGGGGGGGTCAGGAGGAGGAGGCTCAGGGGGCTCATCATCAGGAGGAACCGGCGACAAAGGGACCCGAGTGTTCAAGAAGGCCAGTCCAAATGGAAAGCTCACCGTCTACCTGGGAAAGCGGGACTTTGTGGACCACATCGACCTCGTGGACCCTGTGGATGGTGTGGTCCTGGTGGATCCTGAGTATCTCAAAGAGCGGAGAGTCTATGTGACGCTGACCTGCGCCTTCCGCTATGGCCGGGAGGACCTGGATGTCCTGGGCCTGACCTTTCGCAAGGACCTGTTTGTGGCCAACGTACAGTCGTTCCCACCGGCCCCCGAGGACAAGAAGCCCCTGACGCGGCTGCAGGAACGCCTCATCAAGAAGCTGGGCGAGCACGCTTACCCTTTCACCTTTGAGATCCCTCCAAACCTTCCATGTTCTGTGACACTGCAGCCGGGGCCCGAAGACACGGGGAAGGCTTGCGGTGTGGACTATGAAGTCAAAGCCTTCTGCGCGGAGAATTTGGAGGAGAAGATCCACAAGCGGAATTCTGTGCGTCTGGTCATCCGGAAGGTTCAGTATGCCCCAGAGAGGCCTGGCCCCCAGCCCACAGCCGAGACCACCAGGCAGTTCCTCATGTCGGACAAGCCCTTGCACCTAGAAGCCTCTCTGGATAAGGAGATCTATTACCATGGAGAACCCATCAGCGTCAACGTCCACGTCACCAACAACACCAACAAGACGGTGAAGAAGATCAAGATCTCAGTGCGCCAGTATGCAGACATCTGCCTTTTCAACACAGCTCAGTACAAGTGCCCTGTTGCCATGGAAGAGGCTGATGACACTGTGGCACCCAGCTCGACGTTCTGCAAGGTCTACACACTGACCCCCTTCCTAGCCAATAACCGAGAGAAGCGGGGCCTCGCCTTGGACGGGAAGCTCAAGCACGAAGACACGAACTTGGCCTCTAGCACCCTGTTGAGGGAAGGTGCCAACCGTGAGATCCTGGGGATCATTGTTTCCTACAAAGTGAAAGTGAAGCTGGTGGTGTCTCGGGGCGGCCTGTTGGGAGATCTTGCATCCAGCGACGTGGCCGTGGAACTGCCCTTCACCCTAATGCACCCCAAGCCCAAAGAGGAACCCCCGCATCGGGAAGTTCCAGAGAACGAGACGCCAGTAGATACCAATCTCATAGAACTTGACACAAATGATGACGACATTGTATTTGAGGACTTTGCTCGCCAGAGACTGAAAGGCATGAAGGATGACAAGGAGGAAGAGGAGGATGGTACCGGCTCTCCACAGCTCAACAACAGATGA

SmBiT-linker-β-arrestin1 (ΔC382)

ATGGTCACCGGCTATCGGCTGTTTGAAGAGATTCTGGGGGGGTCAGGAGGAGGAGGCTCAGGGGGCTCATCATCAGGAGGAACCGGCGACAAAGGGACCCGAGTGTTCAAGAAGGCCAGTCCAAATGGAAAGCTCACCGTCTACCTGGGAAAGCGGGACTTTGTGGACCACATCGACCTCGTGGACCCTGTGGATGGTGTGGTCCTGGTGGATCCTGAGTATCTCAAAGAGCGGAGAGTCTATGTGACGCTGACCTGCGCCTTCCGCTATGGCCGGGAGGACCTGGATGTCCTGGGCCTGACCTTTCGCAAGGACCTGTTTGTGGCCAACGTACAGTCGTTCCCACCGGCCCCCGAGGACAAGAAGCCCCTGACGCGGCTGCAGGAACGCCTCATCAAGAAGCTGGGCGAGCACGCTTACCCTTTCACCTTTGAGATCCCTCCAAACCTTCCATGTTCTGTGACACTGCAGCCGGGGCCCGAAGACACGGGGAAGGCTTGCGGTGTGGACTATGAAGTCAAAGCCTTCTGCGCGGAGAATTTGGAGGAGAAGATCCACAAGCGGAATTCTGTGCGTCTGGTCATCCGGAAGGTTCAGTATGCCCCAGAGAGGCCTGGCCCCCAGCCCACAGCCGAGACCACCAGGCAGTTCCTCATGTCGGACAAGCCCTTGCACCTAGAAGCCTCTCTGGATAAGGAGATCTATTACCATGGAGAACCCATCAGCGTCAACGTCCACGTCACCAACAACACCAACAAGACGGTGAAGAAGATCAAGATCTCAGTGCGCCAGTATGCAGACATCTGCCTTTTCAACACAGCTCAGTACAAGTGCCCTGTTGCCATGGAAGAGGCTGATGACACTGTGGCACCCAGCTCGACGTTCTGCAAGGTCTACACACTGACCCCCTTCCTAGCCAATAACCGAGAGAAGCGGGGCCTCGCCTTGGACGGGAAGCTCAAGCACGAAGACACGAACTTGGCCTCTAGCACCCTGTTGAGGGAAGGTGCCAACCGTGAGATCCTGGGGATCATTGTTTCCTACAAAGTGAAAGTGAAGCTGGTGGTGTCTCGGGGCGGCCTGTTGGGAGATCTTGCATCCAGCGACGTGGCCGTGGAACTGCCCTTCACCCTAATGCACCCCAAGCCCAAAGAGGAACCCCCGCATCGGGAAGTTCCAGAGAACGAGACGCCAGTAGATACCAATCTCATAGAACTTGACACAAATTGA

SmBiT-linker-β-arrestin2

ATGGTCACCGGCTATCGGCTGTTTGAAGAGATTCTGGGGGGGTCAGGAGGAGGAGGCTCAGGGGGCTCATCATCAGGAGGAACCGGGGAGAAACCCGGGACCAGGGTCTTCAAGAAGTCGAGCCCTAACTGCAAGCTCACCGTGTACTTGGGCAAGCGGGACTTCGTAGATCACCTGGACAAAGTGGACCCTGTAGATGGCGTGGTGCTTGTGGACCCTGACTACCTGAAGGACCGCAAAGTGTTTGTGACCCTCACCTGCGCCTTCCGCTATGGCCGTGAAGACCTGGATGTGCTGGGCTTGTCCTTCCGCAAAGACCTGTTCATCGCCACCTACCAGGCCTTCCCCCCGGTGCCCAACCCACCCCGGCCCCCCACCCGCCTGCAGGACCGGCTGCTGAGGAAGCTGGGCCAGCATGCCCACCCCTTCTTCTTCACCATACCCCAGAATCTTCCATGCTCCGTCACACTGCAGCCAGGCCCAGAGGATACAGGAAAGGCCTGCGGCGTAGACTTTGAGATTCGAGCCTTCTGTGCTAAATCACTAGAAGAGAAAAGCCACAAAAGGAACTCTGTGCGGCTGGTGATCCGAAAGGTGCAGTTCGCCCCGGAGAAACCCGGCCCCCAGCCTTCAGCCGAAACCACACGCCACTTCCTCATGTCTGACCGGTCCCTGCACCTCGAGGCTTCCCTGGACAAGGAGCTGTACTACCATGGGGAGCCCCTCAATGTAAATGTCCACGTCACCAACAACTCCACCAAGACCGTCAAGAAGATCAAAGTCTCTGTGAGACAGTACGCCGACATCTGCCTCTTCAGCACCGCCCAGTACAAGTGTCCTGTGGCTCAACTCGAACAAGATGACCAGGTATCTCCCAGCTCCACATTCTGTAAGGTGTACACCATAACCCCACTGCTCAGCGACAACCGGGAGAAGCGGGGTCTCGCCCTGGATGGGAAACTCAAGCACGAGGACACCAACCTGGCTTCCAGCACCATCGTGAAGGAGGGTGCCAACAAGGAGGTGCTGGGAATCCTGGTGTCCTACAGGGTCAAGGTGAAGCTGGTGGTGTCTCGAGGCGGGGATGTCTCTGTGGAGCTGCCTTTTGTTCTTATGCACCCCAAGCCCCACGACCACATCCCCCTCCCCAGACCCCAGTCAGCCGCTCCGGAGACAGATGTCCCTGTGGACACCAACCTCATTGAATTTGATACCAACTATGCCACAGATGATGACATTGTGTTTGAGGACTTTGCCCGGCTTCGGCTGAAGGGGATGAAGGATGACGACTATGATGATCAACTCTGCTGA

SmBiT-linker-β-arrestin2 (ΔC382)

ATGGTCACCGGCTATCGGCTGTTTGAAGAGATTCTGGGGGGGTCAGGAGGAGGAGGCTCAGGGGGCTCATCATCAGGAGGAACCGGGGAGAAACCCGGGACCAGGGTCTTCAAGAAGTCGAGCCCTAACTGCAAGCTCACCGTGTACTTGGGCAAGCGGGACTTCGTAGATCACCTGGACAAAGTGGACCCTGTAGATGGCGTGGTGCTTGTGGACCCTGACTACCTGAAGGACCGCAAAGTGTTTGTGACCCTCACCTGCGCCTTCCGCTATGGCCGTGAAGACCTGGATGTGCTGGGCTTGTCCTTCCGCAAAGACCTGTTCATCGCCACCTACCAGGCCTTCCCCCCGGTGCCCAACCCACCCCGGCCCCCCACCCGCCTGCAGGACCGGCTGCTGAGGAAGCTGGGCCAGCATGCCCACCCCTTCTTCTTCACCATACCCCAGAATCTTCCATGCTCCGTCACACTGCAGCCAGGCCCAGAGGATACAGGAAAGGCCTGCGGCGTAGACTTTGAGATTCGAGCCTTCTGTGCTAAATCACTAGAAGAGAAAAGCCACAAAAGGAACTCTGTGCGGCTGGTGATCCGAAAGGTGCAGTTCGCCCCGGAGAAACCCGGCCCCCAGCCTTCAGCCGAAACCACACGCCACTTCCTCATGTCTGACCGGTCCCTGCACCTCGAGGCTTCCCTGGACAAGGAGCTGTACTACCATGGGGAGCCCCTCAATGTAAATGTCCACGTCACCAACAACTCCACCAAGACCGTCAAGAAGATCAAAGTCTCTGTGAGACAGTACGCCGACATCTGCCTCTTCAGCACCGCCCAGTACAAGTGTCCTGTGGCTCAACTCGAACAAGATGACCAGGTATCTCCCAGCTCCACATTCTGTAAGGTGTACACCATAACCCCACTGCTCAGCGACAACCGGGAGAAGCGGGGTCTCGCCCTGGATGGGAAACTCAAGCACGAGGACACCAACCTGGCTTCCAGCACCATCGTGAAGGAGGGTGCCAACAAGGAGGTGCTGGGAATCCTGGTGTCCTACAGGGTCAAGGTGAAGCTGGTGGTGTCTCGAGGCGGGGATGTCTCTGTGGAGCTGCCTTTTGTTCTTATGCACCCCAAGCCCCACGACCACATCCCCCTCCCCAGACCCCAGTCAGCCGCTCCGGAGACAGATGTCCCTGTGGACACCAACCTCATTGAATTTGATACCAACTATGCCACATGA

LgBiT-linker-CAAX

ATGGTGTTTACTCTGGAGGACTTCGTCGGAGACTGGGAACAGACTGCTGCTTACAATCTGGATCAGGTGCTGGAACAGGGGGGGGTCAGCTCCCTGCTCCAGAACCTGGCCGTGTCTGTGACACCTATCCAGCGGATCGTGAGAAGCGGCGAGAATGCCCTGAAGATCGACATCCACGTGATCATCCCATACGAGGGCCTGTCCGCCGATCAGATGGCCCAGATCGAGGAGGTGTTCAAGGTGGTGTACCCAGTGGACGATCACCACTTCAAAGTGATCCTGCCCTATGGCACCCTGGTCATCGACGGAGTGACCCCAAACATGCTGAATTACTTCGGCAGGCCTTATGAGGGCATCGCCGTGTTTGATGGCAAGAAGATCACCGTGACAGGCACCCTGTGGAACGGCAATAAGATCATCGACGAGCGGCTGATCACCCCCGATGGCTCTATGCTGTTCAGAGTGACCATCAATAGCGGGGGGTCCGGGGGGGGCGGATCAGGCGGCAGCTCCTCTGGAGGAGGCAAGAAGAAGAAGAAGAAGAGCAAGACCAAGTGCGTGATCATGTGA

Lyn-linker-LgBiT

ATGGGATGTATAAAATCAAAAGGGAAAGACAGCGGGGGATCTGGGGGGGGGGGCTCTGGAGGCTCTTCATCAGGGGGAGTGTTTACTCTGGAGGACTTCGTCGGAGACTGGGAACAGACTGCTGCTTACAATCTGGATCAGGTGCTGGAACAGGGGGGGGTCAGCTCCCTGCTCCAGAACCTGGCCGTGTCTGTGACACCTATCCAGCGGATCGTGAGAAGCGGCGAGAATGCCCTGAAGATCGACATCCACGTGATCATCCCATACGAGGGCCTGTCCGCCGATCAGATGGCCCAGATCGAGGAGGTGTTCAAGGTGGTGTACCCAGTGGACGATCACCACTTCAAAGTGATCCTGCCCTATGGCACCCTGGTCATCGACGGAGTGACCCCAAACATGCTGAATTACTTCGGCAGGCCTTATGAGGGCATCGCCGTGTTTGATGGCAAGAAGATCACCGTGACAGGCACCCTGTGGAACGGCAATAAGATCATCGACGAGCGGCTGATCACCCCCGATGGCTCTATGCTGTTCAGAGTGACCATCAATAGCTGA

GAP43-linker-LgBiT

ATGTGCTGTCTGAGAAGAACCAAACAGGTTGAAAAGAATGATGAGGACCAAAAGATCATGGTGAGCAAGGGGGGATCTGGGGGGGGGGGCTCTGGAGGCTCTTCATCAGGGGGAGTGTTTACTCTGGAGGACTTCGTCGGAGACTGGGAACAGACTGCTGCTTACAATCTGGATCAGGTGCTGGAACAGGGGGGGGTCAGCTCCCTGCTCCAGAACCTGGCCGTGTCTGTGACACCTATCCAGCGGATCGTGAGAAGCGGCGAGAATGCCCTGAAGATCGACATCCACGTGATCATCCCATACGAGGGCCTGTCCGCCGATCAGATGGCCCAGATCGAGGAGGTGTTCAAGGTGGTGTACCCAGTGGACGATCACCACTTCAAAGTGATCCTGCCCTATGGCACCCTGGTCATCGACGGAGTGACCCCAAACATGCTGAATTACTTCGGCAGGCCTTATGAGGGCATCGCCGTGTTTGATGGCAAGAAGATCACCGTGACAGGCACCCTGTGGAACGGCAATAAGATCATCGACGAGCGGCTGATCACCCCCGATGGCTCTATGCTGTTCAGAGTGACCATCAATAGCTGA

ssHA-FLAG-linker-CD86-linker-LgBiT

ATGAAGACTATTATCGCTCTGAGTTACATTTTCTGCCTGGTGTTCGCTGATTACAAGGACGACGACGACAAAGGGGGCTCTGGGGGGGGAGGCTCCGGAGGCAGCTCCTCTGGAGGAGGTGATCCCCAGTGCACTATGGGACTGAGTAACATTCTCTTTGTGATGGCCTTCCTGCTCTCTGGTGCTGCTCCTCTGAAGATTCAAGCTTATTTCAATGAGACTGCAGACCTGCCATGCCAATTTGCAAACTCTCAAAACCAAAGCCTGAGTGAGCTAGTAGTATTTTGGCAGGACCAGGAAAACTTGGTTCTGAATGAGGTATACTTAGGCAAAGAGAAATTTGACAGTGTTCATTCCAAGTATATGGGCCGCACAAGTTTTGATTCGGACAGTTGGACCCTGAGACTTCACAATCTTCAGATCAAGGACAAGGGCTTGTATCAATGTATCATCCATCACAAAAAGCCCACAGGAATGATTCGCATCCACCAGATGAATTCTGAACTGTCAGTGCTTGCTAACTTCAGTCAACCTGAAATAGTACCAATTTCTAATATAACAGAAAATGTGTACATAAATTTGACCTGCTCATCTATACACGGTTACCCAGAACCTAAGAAGATGAGTGTTTTGCTAAGAACCAAGAATTCAACTATCGAGTATGATGGTATTATGCAGAAATCTCAAGATAATGTCACAGAACTGTACGACGTTTCCATCAGCTTGTCTGTTTCATTCCCTGATGTTACGAGCAATATGACCATCTTCTGTATTCTGGAAACTGACAAGACGCGGCTTTTATCTTCACCTTTCTCTATAGAGCTTGAGGACCCTCAGCCTCCCCCAGACCACATTCCTTGGATTACAGCTGTACTTCCAACAGTTATTATATGTGTGATGGTTTTCTGTCTAATTCTATGGAAATGGAAGAAGAAGAAGCGGCCTCGCGGGGGATCTGGGGGGGGGGGCTCTGGAGGCTCTTCATCAGGGGGAGTGTTTACTCTGGAGGACTTCGTCGGAGACTGGGAACAGACTGCTGCTTACAATCTGGATCAGGTGCTGGAACAGGGGGGGGTCAGCTCCCTGCTCCAGAACCTGGCCGTGTCTGTGACACCTATCCAGCGGATCGTGAGAAGCGGCGAGAATGCCCTGAAGATCGACATCCACGTGATCATCCCATACGAGGGCCTGTCCGCCGATCAGATGGCCCAGATCGAGGAGGTGTTCAAGGTGGTGTACCCAGTGGACGATCACCACTTCAAAGTGATCCTGCCCTATGGCACCCTGGTCATCGACGGAGTGACCCCAAACATGCTGAATTACTTCGGCAGGCCTTATGAGGGCATCGCCGTGTTTGATGGCAAGAAGATCACCGTGACAGGCACCCTGTGGAACGGCAATAAGATCATCGACGAGCGGCTGATCACCCCCGATGGCTCTATGCTGTTCAGAGTGACCATCAATAGCTGA

Endofin-linker-LgBiT

ATGCAGAAGCAGCCAACATGGGTCCCCGATAGCGAAGCCCCTAACTGTATGAATTGCCAGGTCAAGTTTACTTTCACTAAGCGAAGACACCACTGCCGGGCATGTGGCAAGGTGTTCTGCGGCGTGTGCTGTAACCGGAAGTGTAAGCTCCAGTACCTGGAGAAGGAGGCCCGGGTGTGCGTGGTGTGCTATGAGACCATCAGCAAGGGCGGCAGCGGCGGCGGCGGGAGCGGCGGAAGTAGTAGCGGAGGCGTGTTTACTCTGGAGGACTTCGTCGGAGACTGGGAACAGACTGCTGCTTACAATCTGGATCAGGTGCTGGAACAGGGGGGGGTCAGCTCCCTGCTCCAGAACCTGGCCGTGTCTGTGACACCTATCCAGCGGATCGTGAGAAGCGGCGAGAATGCCCTGAAGATCGACATCCACGTGATCATCCCATACGAGGGCCTGTCCGCCGATCAGATGGCCCAGATCGAGGAGGTGTTCAAGGTGGTGTACCCAGTGGACGATCACCACTTCAAAGTGATCCTGCCCTATGGCACCCTGGTCATCGACGGAGTGACCCCAAACATGCTGAATTACTTCGGCAGGCCTTATGAGGGCATCGCCGTGTTTGATGGCAAGAAGATCACCGTGACAGGCACCCTGTGGAACGGCAATAAGATCATCGACGAGCGGCTGATCACCCCCGATGGCTCTATGCTGTTCAGAGTGACCATCAATAGCTGA

LgBiT-linker-Rab4

ATGGTGTTTACTCTGGAGGACTTCGTCGGAGACTGGGAACAGACTGCTGCTTACAATCTGGATCAGGTGCTGGAACAGGGGGGGGTCAGCTCCCTGCTCCAGAACCTGGCCGTGTCTGTGACACCTATCCAGCGGATCGTGAGAAGCGGCGAGAATGCCCTGAAGATCGACATCCACGTGATCATCCCATACGAGGGCCTGTCCGCCGATCAGATGGCCCAGATCGAGGAGGTGTTCAAGGTGGTGTACCCAGTGGACGATCACCACTTCAAAGTGATCCTGCCCTATGGCACCCTGGTCATCGACGGAGTGACCCCAAACATGCTGAATTACTTCGGCAGGCCTTATGAGGGCATCGCCGTGTTTGATGGCAAGAAGATCACCGTGACAGGCACCCTGTGGAACGGCAATAAGATCATCGACGAGCGGCTGATCACCCCCGATGGCTCTATGCTGTTCAGAGTGACCATCAATAGCGGGGGGAGCGGCGGGGGGGGGAGCGGGGGAAGCAGTAGTGGCGGAATGTCGCAGACGGCCATGTCCGAAACCTACGATTTTTTGTTTAAGTTCTTGGTTATTGGAAATGCAGGAACTGGCAAATCTTGCTTACTTCATCAGTTTATTGAAAAAAAATTCAAAGATGACTCAAATCATACAATAGGAGTGGAATTTGGTTCAAAGATAATAAATGTTGGTGGTAAATATGTAAAGTTACAAATATGGGATACAGCAGGACAAGAACGATTCAGGTCCGTGACGAGAAGTTATTACCGAGGCGCGGCCGGGGCTCTCCTCGTCTATGATATCACCAGCCGAGAAACCTACAATGCGCTTACTAATTGGTTAACAGATGCCCGAATGCTGGCGAGCCAGAACATTGTGATCATCCTTTGTGGAAACAAGAAGGACCTGGATGCAGATCGTGAAGTTACCTTCTTAGAAGCCTCCAGATTTGCTCAAGAAAATGAGCTGATGTTTTTGGAAACAAGTGCGCTCACAGGGGAGAATGTAGAAGAGGCTTTTGTACAGTGTGCAAGAAAAATACTTAACAAAATCGAATCAGGTGAGCTGGACCCAGAAAGAATGGGCTCAGGTATTCAGTACGGAGATGCTGCCTTGAGACAGCTGAGGTCACCGCGGCGCGCACAGGCCCCGAACGCTCAGGAGTGTGGTTGTTGA

LgBiT-linker-Rab5

ATGGTGTTTACTCTGGAGGACTTCGTCGGAGACTGGGAACAGACTGCTGCTTACAATCTGGATCAGGTGCTGGAACAGGGGGGGGTCAGCTCCCTGCTCCAGAACCTGGCCGTGTCTGTGACACCTATCCAGCGGATCGTGAGAAGCGGCGAGAATGCCCTGAAGATCGACATCCACGTGATCATCCCATACGAGGGCCTGTCCGCCGATCAGATGGCCCAGATCGAGGAGGTGTTCAAGGTGGTGTACCCAGTGGACGATCACCACTTCAAAGTGATCCTGCCCTATGGCACCCTGGTCATCGACGGAGTGACCCCAAACATGCTGAATTACTTCGGCAGGCCTTATGAGGGCATCGCCGTGTTTGATGGCAAGAAGATCACCGTGACAGGCACCCTGTGGAACGGCAATAAGATCATCGACGAGCGGCTGATCACCCCCGATGGCTCTATGCTGTTCAGAGTGACCATCAATAGCGGGGGGAGCGGCGGGGGGGGGAGCGGGGGAAGCAGTAGTGGCGGAGCTAGTCGAGGCGCAACAAGACCCAACGGGCCAAATACGGGAAATAAAATATGCCAGTTCAAACTAGTACTTCTGGGAGAGTCCGCTGTTGGCAAATCAAGCCTAGTGCTTCGTTTTGTGAAAGGCCAATTTCATGAATTTCAAGAGAGTACCATTGGGGCTGCTTTTCTAACCCAAACTGTATGTCTTGATGACACTACAGTAAAGTTTGAAATATGGGATACAGCTGGTCAAGAACGATACCATAGCCTAGCACCAATGTACTACAGAGGAGCACAAGCAGCCATAGTTGTATATGATATCACAAATGAGGAGTCCTTTGCAAGAGCAAAAAATTGGGTTAAAGAACTTCAGAGGCAAGCAAGTCCTAACATTGTAATAGCTTTATCGGGAAACAAGGCCGACCTAGCAAATAAAAGAGCAGTAGATTTCCAGGAAGCACAGTCCTATGCAGATGACAATAGTTTATTATTCATGGAGACATCCGCTAAAACATCAATGAATGTAAATGAAATATTCATGGCAATAGCTAAAAAATTGCCAAAGAATGAACCACAAAATCCAGGAGCAAATTCTGCCAGAGGAAGAGGAGTAGACCTTACCGAACCCACACAACCAACCAGGAATCAGTGTTGTAGTAACTAA

LgBiT-linker-Rab7

ATGGTGTTTACTCTGGAGGACTTCGTCGGAGACTGGGAACAGACTGCTGCTTACAATCTGGATCAGGTGCTGGAACAGGGGGGGGTCAGCTCCCTGCTCCAGAACCTGGCCGTGTCTGTGACACCTATCCAGCGGATCGTGAGAAGCGGCGAGAATGCCCTGAAGATCGACATCCACGTGATCATCCCATACGAGGGCCTGTCCGCCGATCAGATGGCCCAGATCGAGGAGGTGTTCAAGGTGGTGTACCCAGTGGACGATCACCACTTCAAAGTGATCCTGCCCTATGGCACCCTGGTCATCGACGGAGTGACCCCAAACATGCTGAATTACTTCGGCAGGCCTTATGAGGGCATCGCCGTGTTTGATGGCAAGAAGATCACCGTGACAGGCACCCTGTGGAACGGCAATAAGATCATCGACGAGCGGCTGATCACCCCCGATGGCTCTATGCTGTTCAGAGTGACCATCAATAGCGGGGGGAGCGGCGGGGGGGGGAGCGGGGGAAGCAGTAGTGGCGGAATGACCTCTAGGAAGAAAGTGTTGCTGAAGGTTATCATCCTGGGAGATTCTGGAGTCGGGAAGACATCACTCATGAACCAGTATGTGAATAAGAAATTCAGCAATCAGTACAAAGCCACAATAGGAGCTGACTTTCTGACCAAGGAGGTGATGGTGGATGACAGGCTAGTCACAATGCAGATATGGGACACAGCAGGACAGGAACGGTTCCAGTCTCTCGGTGTGGCCTTCTACAGAGGTGCAGACTGCTGCGTTCTGGTATTTGATGTGACTGCCCCCAACACATTCAAAACCCTAGATAGCTGGAGAGATGAGTTTCTCATCCAGGCCAGTCCCCGAGATCCTGAAAACTTCCCATTTGTTGTGTTGGGAAACAAGATTGACCTCGAAAACAGACAAGTGGCCACAAAGCGGGCACAGGCCTGGTGCTACAGCAAAAACAACATTCCCTACTTTGAGACCAGTGCCAAGGAGGCCATCAACGTGGAGCAGGCGTTCCAGACGATTGCACGGAATGCACTTAAGCAGGAAACGGAGGTGGAGCTGTACAACGAATTTCCTGAACCTATCAAACTGGACAAGAATGACCGGGCCAAGGCCTCGGCAGAAAGCTGCAGTTGCTGA

LgBiT-linker-Rab11

ATGGTGTTTACTCTGGAGGACTTCGTCGGAGACTGGGAACAGACTGCTGCTTACAATCTGGATCAGGTGCTGGAACAGGGGGGGGTCAGCTCCCTGCTCCAGAACCTGGCCGTGTCTGTGACACCTATCCAGCGGATCGTGAGAAGCGGCGAGAATGCCCTGAAGATCGACATCCACGTGATCATCCCATACGAGGGCCTGTCCGCCGATCAGATGGCCCAGATCGAGGAGGTGTTCAAGGTGGTGTACCCAGTGGACGATCACCACTTCAAAGTGATCCTGCCCTATGGCACCCTGGTCATCGACGGAGTGACCCCAAACATGCTGAATTACTTCGGCAGGCCTTATGAGGGCATCGCCGTGTTTGATGGCAAGAAGATCACCGTGACAGGCACCCTGTGGAACGGCAATAAGATCATCGACGAGCGGCTGATCACCCCCGATGGCTCTATGCTGTTCAGAGTGACCATCAATAGCGGGGGGAGCGGCGGGGGGGGGAGCGGGGGAAGCAGTAGTGGCGGAggtacccgcgacgacgagtacgactacctctttaaagttgtccttattggagattctggtgttggaaagagtaatctcctgtctcgatttactcgaaatgagtttaatctggaaagcaagagcaccattggagtagagtttgcaacaagaagcatccaggttgatggaaaaacaataaaggcacagatatgggacacagcagggcaagagcgatatcgagctataacatcagcatattatcgtggagctgtaggtgccttattggtttatgacattgctaaacatctcacatatgaaaatgtagagcgatggctgaaagaactgagagatcatgctgatagtaacattgttatcatgcttgtgggcaataagagtgatctacgtcatctcagggcagttcctacagatgaagcaagagcttttgcagaaaagaatggtttgtcattcattgaaacttcggccctagactctacaaatgtagaagctgcttttcagacaattttaacagagatttaccgcattgtttctcagaagcaaatgtcagacagacgcgaaaatgacatgtctccaagcaacaatgtggttcctattcatgttccaccaaccactgaaaacaagccaaaggtgcagtgctgtcagaacatctga

GRK2-linker-LgBiT

ATGGCTGACCTGGAGGCTGTCCTGGCCGATGTCTCCTATCTGATGGCTATGGAAAAGAGTAAGGCAACACCTGCTGCAAGGGCAAGCAAGAAGATCCTGCTGCCTGAGCCAAGCATCAGGTCCGTGATGCAGAAGTACCTGGAGGACCGCGGCGAGGTGACCTTCGAGAAGATCTTTTCCCAGAAGCTGGGCTATCTGCTGTTCCGGGATTTTTGCCTGAACCACCTGGAGGAGGCCAGACCCCTGGTGGAGTTTTACGAGGAGATCAAGAAGTATGAGAAGCTGGAGACAGAGGAGGAGAGGGTGGCACGGAGCAGAGAGATCTTTGACTCCTACATCATGAAGGAGCTGCTGGCCTGTTCTCACCCTTTCTCTAAGAGCGCCACAGAGCACGTGCAGGGACACCTGGGCAAGAAGCAGGTGCCACCTGATCTGTTCCAGCCATATATCGAGGAGATCTGCCAGAACCTGAGGGGCGACGTGTTCCAGAAGTTTATCGAGTCTGATAAGTTCACCCGCTTTTGTCAGTGGAAGAACGTGGAGCTGAATATCCACCTGACAATGAATGACTTTAGCGTGCACAGGATCATCGGAAGGGGAGGATTCGGAGAGGTGTACGGCTGCAGGAAGGCCGACACCGGCAAGATGTATGCCATGAAGTGTCTGGATAAGAAGCGGATCAAGATGAAGCAGGGCGAGACACTGGCCCTGAACGAGAGAATCATGCTGTCTCTGGTGAGCACAGGCGACTGCCCCTTCATCGTGTGCATGTCTTACGCCTTTCACACACCAGATAAGCTGAGCTTCATCCTGGACCTGATGAATGGCGGCGATCTGCACTACCACCTGTCTCAGCACGGCGTGTTCAGCGAGGCCGATATGCGGTTTTATGCCGCCGAGATCATCCTGGGCCTGGAGCACATGCACAACAGGTTTGTGGTGTATCGCGACCTGAAGCCTGCCAATATCCTGCTGGATGAGCACGGCCACGTGAGAATCAGCGACCTGGGCCTGGCCTGCGACTTCAGCAAGAAGAAGCCACACGCATCTGTGGGAACCCACGGCTACATGGCACCTGAGGTGCTGCAGAAGGGAGTGGCCTATGACAGCTCCGCCGATTGGTTCTCCCTGGGCTGTATGCTGTTTAAGCTGCTGCGGGGCCACTCTCCCTTTAGACAGCACAAGACAAAGGACAAGCACGAGATCGATAGGATGACCCTGACAATGGCCGTGGAGCTGCCCGACTCCTTCTCTCCTGAGCTGCGGAGCCTGCTGGAGGGCCTGCTGCAGAGAGATGTGAACCGGAGACTGGGATGCCTGGGAAGGGGAGCACAGGAGGTGAAGGAGTCCCCCTTCTTTCGCTCTCTGGACTGGCAGATGGTGTTCCTGCAGAAGTACCCACCACCTCTGATCCCACCAAGGGGAGAGGTGAACGCAGCAGACGCCTTCGATATCGGCTCCTTTGACGAGGAGGATACCAAGGGCATCAAGCTGCTGGACAGCGATCAGGAGCTGTATAGGAATTTTCCTCTGACAATCTCCGAGCGGTGGCAGCAGGAGGTGGCAGAGACAGTGTTCGACACAATCAACGCCGAGACAGATCGGCTGGAGGCCAGAAAGAAGGCCAAGAATAAGCAGCTGGGACACGAGGAGGACTACGCACTGGGCAAGGATTGTATCATGCACGGCTATATGAGCAAGATGGGCAACCCATTTCTGACACAGTGGCAGAGGCGCTACTTCTATCTGTTTCCAAATAGGCTGGAGTGGAGGGGAGAGGGAGAGGCACCACAGAGCCTGCTGACCATGGAGGAGATCCAGTCCGTGGAGGAGACACAGATCAAGGAGAGGAAGTGCCTGCTGCTGAAGATCCGCGGCGGCAAGCAGTTCATCCTGCAGTGTGACTCCGATCCCGAGCTGGTGCAGTGGAAGAAGGAGCTGCGGGACGCCTACAGAGAGGCACAGCAGCTGGTGCAGAGGGTGCCAAAGATGAAGAATAAGCCCAGAAGCCCTGTCGTGGAACTGAGCAAAGTGCCTCTGGTGCAGCGGGGAAGCGCAAACGGCCTGGGGGGATCTGGGGGGGGGGGCTCTGGAGGCTCTTCATCAGGGGGAGTGTTTACTCTGGAGGACTTCGTCGGAGACTGGGAACAGACTGCTGCTTACAATCTGGATCAGGTGCTGGAACAGGGGGGGGTCAGCTCCCTGCTCCAGAACCTGGCCGTGTCTGTGACACCTATCCAGCGGATCGTGAGAAGCGGCGAGAATGCCCTGAAGATCGACATCCACGTGATCATCCCATACGAGGGCCTGTCCGCCGATCAGATGGCCCAGATCGAGGAGGTGTTCAAGGTGGTGTACCCAGTGGACGATCACCACTTCAAAGTGATCCTGCCCTATGGCACCCTGGTCATCGACGGAGTGACCCCAAACATGCTGAATTACTTCGGCAGGCCTTATGAGGGCATCGCCGTGTTTGATGGCAAGAAGATCACCGTGACAGGCACCCTGTGGAACGGCAATAAGATCATCGACGAGCGGCTGATCACCCCCGATGGCTCTATGCTGTTCAGAGTGACCATCAATAGCTTGA

GRK2 (K220M)-linker-LgBiT

ATGGCTGACCTGGAGGCTGTCCTGGCCGATGTCTCCTATCTGATGGCTATGGAAAAGAGTAAGGCAACACCTGCTGCAAGGGCAAGCAAGAAGATCCTGCTGCCTGAGCCAAGCATCAGGTCCGTGATGCAGAAGTACCTGGAGGACCGCGGCGAGGTGACCTTCGAGAAGATCTTTTCCCAGAAGCTGGGCTATCTGCTGTTCCGGGATTTTTGCCTGAACCACCTGGAGGAGGCCAGACCCCTGGTGGAGTTTTACGAGGAGATCAAGAAGTATGAGAAGCTGGAGACAGAGGAGGAGAGGGTGGCACGGAGCAGAGAGATCTTTGACTCCTACATCATGAAGGAGCTGCTGGCCTGTTCTCACCCTTTCTCTAAGAGCGCCACAGAGCACGTGCAGGGACACCTGGGCAAGAAGCAGGTGCCACCTGATCTGTTCCAGCCATATATCGAGGAGATCTGCCAGAACCTGAGGGGCGACGTGTTCCAGAAGTTTATCGAGTCTGATAAGTTCACCCGCTTTTGTCAGTGGAAGAACGTGGAGCTGAATATCCACCTGACAATGAATGACTTTAGCGTGCACAGGATCATCGGAAGGGGAGGATTCGGAGAGGTGTACGGCTGCAGGAAGGCCGACACCGGCAAGATGTATGCCATGATGTGTCTGGATAAGAAGCGGATCAAGATGAAGCAGGGCGAGACACTGGCCCTGAACGAGAGAATCATGCTGTCTCTGGTGAGCACAGGCGACTGCCCCTTCATCGTGTGCATGTCTTACGCCTTTCACACACCAGATAAGCTGAGCTTCATCCTGGACCTGATGAATGGCGGCGATCTGCACTACCACCTGTCTCAGCACGGCGTGTTCAGCGAGGCCGATATGCGGTTTTATGCCGCCGAGATCATCCTGGGCCTGGAGCACATGCACAACAGGTTTGTGGTGTATCGCGACCTGAAGCCTGCCAATATCCTGCTGGATGAGCACGGCCACGTGAGAATCAGCGACCTGGGCCTGGCCTGCGACTTCAGCAAGAAGAAGCCACACGCATCTGTGGGAACCCACGGCTACATGGCACCTGAGGTGCTGCAGAAGGGAGTGGCCTATGACAGCTCCGCCGATTGGTTCTCCCTGGGCTGTATGCTGTTTAAGCTGCTGCGGGGCCACTCTCCCTTTAGACAGCACAAGACAAAGGACAAGCACGAGATCGATAGGATGACCCTGACAATGGCCGTGGAGCTGCCCGACTCCTTCTCTCCTGAGCTGCGGAGCCTGCTGGAGGGCCTGCTGCAGAGAGATGTGAACCGGAGACTGGGATGCCTGGGAAGGGGAGCACAGGAGGTGAAGGAGTCCCCCTTCTTTCGCTCTCTGGACTGGCAGATGGTGTTCCTGCAGAAGTACCCACCACCTCTGATCCCACCAAGGGGAGAGGTGAACGCAGCAGACGCCTTCGATATCGGCTCCTTTGACGAGGAGGATACCAAGGGCATCAAGCTGCTGGACAGCGATCAGGAGCTGTATAGGAATTTTCCTCTGACAATCTCCGAGCGGTGGCAGCAGGAGGTGGCAGAGACAGTGTTCGACACAATCAACGCCGAGACAGATCGGCTGGAGGCCAGAAAGAAGGCCAAGAATAAGCAGCTGGGACACGAGGAGGACTACGCACTGGGCAAGGATTGTATCATGCACGGCTATATGAGCAAGATGGGCAACCCATTTCTGACACAGTGGCAGAGGCGCTACTTCTATCTGTTTCCAAATAGGCTGGAGTGGAGGGGAGAGGGAGAGGCACCACAGAGCCTGCTGACCATGGAGGAGATCCAGTCCGTGGAGGAGACACAGATCAAGGAGAGGAAGTGCCTGCTGCTGAAGATCCGCGGCGGCAAGCAGTTCATCCTGCAGTGTGACTCCGATCCCGAGCTGGTGCAGTGGAAGAAGGAGCTGCGGGACGCCTACAGAGAGGCACAGCAGCTGGTGCAGAGGGTGCCAAAGATGAAGAATAAGCCCAGAAGCCCTGTCGTGGAACTGAGCAAAGTGCCTCTGGTGCAGCGGGGAAGCGCAAACGGCCTGGGGGGATCTGGGGGGGGGGGCTCTGGAGGCTCTTCATCAGGGGGAGTGTTTACTCTGGAGGACTTCGTCGGAGACTGGGAACAGACTGCTGCTTACAATCTGGATCAGGTGCTGGAACAGGGGGGGGTCAGCTCCCTGCTCCAGAACCTGGCCGTGTCTGTGACACCTATCCAGCGGATCGTGAGAAGCGGCGAGAATGCCCTGAAGATCGACATCCACGTGATCATCCCATACGAGGGCCTGTCCGCCGATCAGATGGCCCAGATCGAGGAGGTGTTCAAGGTGGTGTACCCAGTGGACGATCACCACTTCAAAGTGATCCTGCCCTATGGCACCCTGGTCATCGACGGAGTGACCCCAAACATGCTGAATTACTTCGGCAGGCCTTATGAGGGCATCGCCGTGTTTGATGGCAAGAAGATCACCGTGACAGGCACCCTGTGGAACGGCAATAAGATCATCGACGAGCGGCTGATCACCCCCGATGGCTCTATGCTGTTCAGAGTGACCATCAATAGCTTGA

GRK3-linker-LgBiT

ATGGCTGACCTGGAAGCAGTCCTGGCTGACGTGTCCTATCTGATGGCTATGGAAAAGAGCAAGGCAACTCCCGCCGCAAGGGCAAGCAAGAGGATCGTGCTGCCAGAGCCCTCCATCCGCTCTGTGATGCAGAAGTACCTGGCCGAGAGGAACGAGATCACCTTTGACAAGATTTTCAACCAGAAGATCGGCTTCCTGCTGTTTAAGGATTTCTGCCTGAACGAGATCAATGAGGCCGTGCCCCAGGTGAAGTTCTACGAGGAGATCAAGGAGTATGAGAAGCTGGACAACGAGGAGGATAGGCTGTGCCGGAGCCGGCAAATCTACGACGCCTATATCATGAAGGAGCTGCTGTCTTGTAGCCACCCATTCAGCAAGCAGGCCGTGGAGCACGTGCAGAGCCACCTGTCCAAGAAGCAGGTGACCTCCACACTGTTTCAGCCCTACATCGAGGAGATTTGCGAGAGCCTGAGGGGCGACATCTTTCAGAAGTTCATGGAGTCCGATAAGTTTACCCGCTTCTGTCAGTGGAAGAACGTGGAGCTGAATATCCACCTGACAATGAATGAGTTCTCTGTGCACCGGATCATCGGCAGAGGAGGATTTGGAGAGGTGTACGGCTGCCGGAAGGCAGACACCGGCAAGATGTATGCCATGAAGTGTCTGGATAAGAAGCGGATCAAGATGAAGCAGGGCGAGACACTGGCCCTGAACGAGAGAATCATGCTGTCTCTGGTGAGCACAGGCGACTGCCCCTTCATCGTGTGCATGACCTACGCCTTCCACACACCCGATAAGCTGTGCTTTATCCTGGACCTGATGAATGGCGGCGATCTGCACTACCACCTGAGCCAGCACGGCGTGTTTTCCGAGAAGGAGATGAGGTTCTATGCCACCGAGATCATCCTGGGCCTGGAGCACATGCACAACCGGTTCGTGGTGTATCGGGACCTGAAGCCTGCCAATATCCTGCTGGATGAGCACGGACACGCAAGGATCAGCGACCTGGGCCTGGCCTGCGACTTCAGCAAGAAGAAGCCTCACGCAAGCGTGGGCACCCACGGATACATGGCACCAGAGGTGCTCCAGAAGGGCACAGCCTATGACAGCTCCGCCGATTGGTTTTCCCTGGGCTGTATGCTGTTCAAGCTGCTGCGGGGCCACTCTCCATTCAGACAGCACAAGACCAAGGACAAGCACGAGATCGATAGAATGACCCTGACAGTGAACGTGGAGCTGCCTGACACATTTTCCCCAGAGCTGAAGTCTCTGCTGGAGGGCCTGCTCCAGAGGGACGTGAGCAAGCGCCTGGGATGTCACGGAGGAGGCTCCCAGGAGGTGAAGGAGCACTCTTTCTTTAAGGGCGTGGACTGGCAGCACGTGTACCTCCAGAAGTATCCACCTCCACTGATCCCACCTCGGGGCGAGGTGAACGCCGCCGACGCCTTTGATATCGGCTCTTTCGACGAGGAGGATACCAAGGGCATCAAGCTGCTGGACTGCGATCAGGAGCTGTACAAGAATTTCCCCCTGGTCATCAGCGAGAGATGGCAGCAGGAGGTGACCGAGACAGTGTATGAGGCCGTGAACGCCGACACAGATAAGATCGAGGCCAGGAAGCGCGCCAAGAATAAGCAGCTCGGACACGAGGAGGACTACGCACTGGGCAAGGATTGTATCATGCACGGCTATATGCTGAAGCTGGGCAATCCCTTTCTGACCCAGTGGCAGCGGAGATACTTTTATCTGTTCCCTAACAGGCTGGAGTGGAGGGGAGAGGGAGAGTCTCGGCAGAATCTGCTGACCATGGAGCAGATCCTGAGCGTGGAGGAGACACAGATCAAGGACAAGAAGTGCATCCTGTTCAGAATCAAGGGCGGCAAGCAGTTCGTGCTCCAGTGTGAGAGCGATCCTGAGTTCGTGCAGTGGAAGAAGGAGCTGAACGAGACATTCAAGGAGGCACAGCGGCTGCTGAGGAGAGCCCCCAAGTTCCTGAATAAGCCTAGGAGCGGGACTGTGGAACTGCCCAAACCCTCACTGTGCCATAGAAACTCAAACGGACTGGGGGGATCTGGGGGGGGGGGCTCTGGAGGCTCTTCATCAGGGGGAGTGTTTACTCTGGAGGACTTCGTCGGAGACTGGGAACAGACTGCTGCTTACAATCTGGATCAGGTGCTGGAACAGGGGGGGGTCAGCTCCCTGCTCCAGAACCTGGCCGTGTCTGTGACACCTATCCAGCGGATCGTGAGAAGCGGCGAGAATGCCCTGAAGATCGACATCCACGTGATCATCCCATACGAGGGCCTGTCCGCCGATCAGATGGCCCAGATCGAGGAGGTGTTCAAGGTGGTGTACCCAGTGGACGATCACCACTTCAAAGTGATCCTGCCCTATGGCACCCTGGTCATCGACGGAGTGACCCCAAACATGCTGAATTACTTCGGCAGGCCTTATGAGGGCATCGCCGTGTTTGATGGCAAGAAGATCACCGTGACAGGCACCCTGTGGAACGGCAATAAGATCATCGACGAGCGGCTGATCACCCCCGATGGCTCTATGCTGTTCAGAGTGACCATCAATAGCTTGA

GRK3 (K220M)-linker-LgBiT

ATGGCTGACCTGGAAGCAGTCCTGGCTGACGTGTCCTATCTGATGGCTATGGAAAAGAGCAAGGCAACTCCCGCCGCAAGGGCAAGCAAGAGGATCGTGCTGCCAGAGCCCTCCATCCGCTCTGTGATGCAGAAGTACCTGGCCGAGAGGAACGAGATCACCTTTGACAAGATTTTCAACCAGAAGATCGGCTTCCTGCTGTTTAAGGATTTCTGCCTGAACGAGATCAATGAGGCCGTGCCCCAGGTGAAGTTCTACGAGGAGATCAAGGAGTATGAGAAGCTGGACAACGAGGAGGATAGGCTGTGCCGGAGCCGGCAAATCTACGACGCCTATATCATGAAGGAGCTGCTGTCTTGTAGCCACCCATTCAGCAAGCAGGCCGTGGAGCACGTGCAGAGCCACCTGTCCAAGAAGCAGGTGACCTCCACACTGTTTCAGCCCTACATCGAGGAGATTTGCGAGAGCCTGAGGGGCGACATCTTTCAGAAGTTCATGGAGTCCGATAAGTTTACCCGCTTCTGTCAGTGGAAGAACGTGGAGCTGAATATCCACCTGACAATGAATGAGTTCTCTGTGCACCGGATCATCGGCAGAGGAGGATTTGGAGAGGTGTACGGCTGCCGGAAGGCAGACACCGGCAAGATGTATGCCATGATGTGTCTGGATAAGAAGCGGATCAAGATGAAGCAGGGCGAGACACTGGCCCTGAACGAGAGAATCATGCTGTCTCTGGTGAGCACAGGCGACTGCCCCTTCATCGTGTGCATGACCTACGCCTTCCACACACCCGATAAGCTGTGCTTTATCCTGGACCTGATGAATGGCGGCGATCTGCACTACCACCTGAGCCAGCACGGCGTGTTTTCCGAGAAGGAGATGAGGTTCTATGCCACCGAGATCATCCTGGGCCTGGAGCACATGCACAACCGGTTCGTGGTGTATCGGGACCTGAAGCCTGCCAATATCCTGCTGGATGAGCACGGACACGCAAGGATCAGCGACCTGGGCCTGGCCTGCGACTTCAGCAAGAAGAAGCCTCACGCAAGCGTGGGCACCCACGGATACATGGCACCAGAGGTGCTCCAGAAGGGCACAGCCTATGACAGCTCCGCCGATTGGTTTTCCCTGGGCTGTATGCTGTTCAAGCTGCTGCGGGGCCACTCTCCATTCAGACAGCACAAGACCAAGGACAAGCACGAGATCGATAGAATGACCCTGACAGTGAACGTGGAGCTGCCTGACACATTTTCCCCAGAGCTGAAGTCTCTGCTGGAGGGCCTGCTCCAGAGGGACGTGAGCAAGCGCCTGGGATGTCACGGAGGAGGCTCCCAGGAGGTGAAGGAGCACTCTTTCTTTAAGGGCGTGGACTGGCAGCACGTGTACCTCCAGAAGTATCCACCTCCACTGATCCCACCTCGGGGCGAGGTGAACGCCGCCGACGCCTTTGATATCGGCTCTTTCGACGAGGAGGATACCAAGGGCATCAAGCTGCTGGACTGCGATCAGGAGCTGTACAAGAATTTCCCCCTGGTCATCAGCGAGAGATGGCAGCAGGAGGTGACCGAGACAGTGTATGAGGCCGTGAACGCCGACACAGATAAGATCGAGGCCAGGAAGCGCGCCAAGAATAAGCAGCTCGGACACGAGGAGGACTACGCACTGGGCAAGGATTGTATCATGCACGGCTATATGCTGAAGCTGGGCAATCCCTTTCTGACCCAGTGGCAGCGGAGATACTTTTATCTGTTCCCTAACAGGCTGGAGTGGAGGGGAGAGGGAGAGTCTCGGCAGAATCTGCTGACCATGGAGCAGATCCTGAGCGTGGAGGAGACACAGATCAAGGACAAGAAGTGCATCCTGTTCAGAATCAAGGGCGGCAAGCAGTTCGTGCTCCAGTGTGAGAGCGATCCTGAGTTCGTGCAGTGGAAGAAGGAGCTGAACGAGACATTCAAGGAGGCACAGCGGCTGCTGAGGAGAGCCCCCAAGTTCCTGAATAAGCCTAGGAGCGGGACTGTGGAACTGCCCAAACCCTCACTGTGCCATAGAAACTCAAACGGACTGGGGGGATCTGGGGGGGGGGGCTCTGGAGGCTCTTCATCAGGGGGAGTGTTTACTCTGGAGGACTTCGTCGGAGACTGGGAACAGACTGCTGCTTACAATCTGGATCAGGTGCTGGAACAGGGGGGGGTCAGCTCCCTGCTCCAGAACCTGGCCGTGTCTGTGACACCTATCCAGCGGATCGTGAGAAGCGGCGAGAATGCCCTGAAGATCGACATCCACGTGATCATCCCATACGAGGGCCTGTCCGCCGATCAGATGGCCCAGATCGAGGAGGTGTTCAAGGTGGTGTACCCAGTGGACGATCACCACTTCAAAGTGATCCTGCCCTATGGCACCCTGGTCATCGACGGAGTGACCCCAAACATGCTGAATTACTTCGGCAGGCCTTATGAGGGCATCGCCGTGTTTGATGGCAAGAAGATCACCGTGACAGGCACCCTGTGGAACGGCAATAAGATCATCGACGAGCGGCTGATCACCCCCGATGGCTCTATGCTGTTCAGAGTGACCATCAATAGCTTGA

GRK5-linker-LgBiT

ATGGAGCTGGAAAACATCGTGGCCAACACGGTCTTGCTGAAAGCCAGGGAAGGGGGCGGAGGAAAGCGCAAAGGGAAAAGCAAGAAGTGGAAAGAAATCCTGAAGTTCCCTCACATTAGCCAGTGTGAAGACCTCCGAAGGACCATAGACAGAGATTACTGCAGTTTATGTGACAAGCAGCCAATCGGGAGGCTGCTTTTCCGGCAGTTTTGTGAAACCAGGCCTGGGCTGGAGTGTTACATTCAGTTCCTGGACTCCGTGGCAGAATATGAAGTTACTCCAGATGAAAAACTGGGAGAGAAAGGGAAGGAAATTATGACCAAGTACCTCACCCCAAAGTCCCCTGTTTTCATAGCCCAAGTTGGCCAAGACCTGGTCTCCCAGACGGAGGAGAAGCTCCTACAGAAGCCGTGCAAAGAACTCTTTTCTGCCTGTGCACAGTCTGTCCACGAGTACCTGAGGGGAGAACCATTCCACGAATATCTGGACAGCATGTTTTTTGACCGCTTTCTCCAGTGGAAGTGGTTGGAAAGGCAACCGGTGACCAAAAACACTTTCAGGCAGTATCGAGTGCTAGGAAAAGGGGGCTTCGGGGAGGTCTGTGCCTGCCAGGTTCGGGCCACGGGTAAAATGTATGCCTGCAAGCGCTTGGAGAAGAAGAGGATCAAAAAGAGGAAAGGGGAGTCCATGGCCCTCAATGAGAAGCAGATCCTCGAGAAGGTCAACAGTCAGTTTGTGGTCAACCTGGCCTATGCCTACGAGACCAAGGATGCACTGTGCTTGGTCCTGACCATCATGAATGGGGGTGACCTGAAGTTCCACATCTACAACATGGGCAACCCTGGCTTCGAGGAGGAGCGGGCCTTGTTTTATGCGGCAGAGATCCTCTGCGGCTTAGAAGACCTCCACCGTGAGAACACCGTCTACCGAGATCTGAAACCTGAAAACATCCTGTTAGATGATTATGGCCACATTAGGATCTCAGACCTGGGCTTGGCTGTGAAGATCCCCGAGGGAGACCTGATCCGCGGCCGGGTGGGCACTGTTGGCTACATGGCTCCAGAGGTCCTGAACAACCAGAGGTACGGCCTGAGCCCCGACTACTGGGGCCTTGGCTGCCTCATCTATGAGATGATCGAGGGCCAGTCGCCGTTCCGCGGCCGCAAGGAGAAGGTGAAGCGGGAGGAGGTGGACCGCCGGGTCCTGGAGACGGAGGAGGTGTACTCCCACAAGTTCTCCGAGGAGGCCAAGTCCATCTGCAAGATGCTGCTCACGAAAGATGCGAAGCAGAGGCTGGGCTGCCAGGAGGAGGGGGCTGCAGAGGTCAAGAGACACCCCTTCTTCAGGAACATGAACTTCAAGCGCTTAGAAGCCGGGATGTTGGACCCTCCCTTCGTTCCAGACCCCCGCGCTGTGTACTGTAAGGACGTGCTGGACATCGAGCAGTTCTCCACTGTGAAGGGCGTCAATCTGGACCACACAGACGACGACTTCTACTCCAAGTTCTCCACGGGCTCTGTGTCCATCCCATGGCAAAACGAGATGATAGAAACAGAATGCTTTAAGGAGCTGAACGTGTTTGGACCTAATGGTACCCTCCCGCCAGATCTGAACAGAAACCACCCTCCGGAACCGCCCAAGAAAGGGCTGCTCCAGAGACTCTTCAAGCGGCAGCATCAGAACAATTCCAAGAGTTCGCCCAGCTCCAAGACCAGTTTTAACCACCACATAAACTCAAACCATGTCAGCTCGAACTCCACCGGAAGCAGCGGGGGATCTGGGGGGGGGGGCTCTGGAGGCTCTTCATCAGGGGGAGTGTTTACTCTGGAGGACTTCGTCGGAGACTGGGAACAGACTGCTGCTTACAATCTGGATCAGGTGCTGGAACAGGGGGGGGTCAGCTCCCTGCTCCAGAACCTGGCCGTGTCTGTGACACCTATCCAGCGGATCGTGAGAAGCGGCGAGAATGCCCTGAAGATCGACATCCACGTGATCATCCCATACGAGGGCCTGTCCGCCGATCAGATGGCCCAGATCGAGGAGGTGTTCAAGGTGGTGTACCCAGTGGACGATCACCACTTCAAAGTGATCCTGCCCTATGGCACCCTGGTCATCGACGGAGTGACCCCAAACATGCTGAATTACTTCGGCAGGCCTTATGAGGGCATCGCCGTGTTTGATGGCAAGAAGATCACCGTGACAGGCACCCTGTGGAACGGCAATAAGATCATCGACGAGCGGCTGATCACCCCCGATGGCTCTATGCTGTTCAGAGTGACCATCAATAGCTTGA

GRK5 (K215M)-linker-LgBiT

ATGGAGCTGGAAAACATCGTGGCCAACACGGTCTTGCTGAAAGCCAGGGAAGGGGGCGGAGGAAAGCGCAAAGGGAAAAGCAAGAAGTGGAAAGAAATCCTGAAGTTCCCTCACATTAGCCAGTGTGAAGACCTCCGAAGGACCATAGACAGAGATTACTGCAGTTTATGTGACAAGCAGCCAATCGGGAGGCTGCTTTTCCGGCAGTTTTGTGAAACCAGGCCTGGGCTGGAGTGTTACATTCAGTTCCTGGACTCCGTGGCAGAATATGAAGTTACTCCAGATGAAAAACTGGGAGAGAAAGGGAAGGAAATTATGACCAAGTACCTCACCCCAAAGTCCCCTGTTTTCATAGCCCAAGTTGGCCAAGACCTGGTCTCCCAGACGGAGGAGAAGCTCCTACAGAAGCCGTGCAAAGAACTCTTTTCTGCCTGTGCACAGTCTGTCCACGAGTACCTGAGGGGAGAACCATTCCACGAATATCTGGACAGCATGTTTTTTGACCGCTTTCTCCAGTGGAAGTGGTTGGAAAGGCAACCGGTGACCAAAAACACTTTCAGGCAGTATCGAGTGCTAGGAAAAGGGGGCTTCGGGGAGGTCTGTGCCTGCCAGGTTCGGGCCACGGGTAAAATGTATGCCTGCATGCGCTTGGAGAAGAAGAGGATCAAAAAGAGGAAAGGGGAGTCCATGGCCCTCAATGAGAAGCAGATCCTCGAGAAGGTCAACAGTCAGTTTGTGGTCAACCTGGCCTATGCCTACGAGACCAAGGATGCACTGTGCTTGGTCCTGACCATCATGAATGGGGGTGACCTGAAGTTCCACATCTACAACATGGGCAACCCTGGCTTCGAGGAGGAGCGGGCCTTGTTTTATGCGGCAGAGATCCTCTGCGGCTTAGAAGACCTCCACCGTGAGAACACCGTCTACCGAGATCTGAAACCTGAAAACATCCTGTTAGATGATTATGGCCACATTAGGATCTCAGACCTGGGCTTGGCTGTGAAGATCCCCGAGGGAGACCTGATCCGCGGCCGGGTGGGCACTGTTGGCTACATGGCTCCAGAGGTCCTGAACAACCAGAGGTACGGCCTGAGCCCCGACTACTGGGGCCTTGGCTGCCTCATCTATGAGATGATCGAGGGCCAGTCGCCGTTCCGCGGCCGCAAGGAGAAGGTGAAGCGGGAGGAGGTGGACCGCCGGGTCCTGGAGACGGAGGAGGTGTACTCCCACAAGTTCTCCGAGGAGGCCAAGTCCATCTGCAAGATGCTGCTCACGAAAGATGCGAAGCAGAGGCTGGGCTGCCAGGAGGAGGGGGCTGCAGAGGTCAAGAGACACCCCTTCTTCAGGAACATGAACTTCAAGCGCTTAGAAGCCGGGATGTTGGACCCTCCCTTCGTTCCAGACCCCCGCGCTGTGTACTGTAAGGACGTGCTGGACATCGAGCAGTTCTCCACTGTGAAGGGCGTCAATCTGGACCACACAGACGACGACTTCTACTCCAAGTTCTCCACGGGCTCTGTGTCCATCCCATGGCAAAACGAGATGATAGAAACAGAATGCTTTAAGGAGCTGAACGTGTTTGGACCTAATGGTACCCTCCCGCCAGATCTGAACAGAAACCACCCTCCGGAACCGCCCAAGAAAGGGCTGCTCCAGAGACTCTTCAAGCGGCAGCATCAGAACAATTCCAAGAGTTCGCCCAGCTCCAAGACCAGTTTTAACCACCACATAAACTCAAACCATGTCAGCTCGAACTCCACCGGAAGCAGCGGGGGATCTGGGGGGGGGGGCTCTGGAGGCTCTTCATCAGGGGGAGTGTTTACTCTGGAGGACTTCGTCGGAGACTGGGAACAGACTGCTGCTTACAATCTGGATCAGGTGCTGGAACAGGGGGGGGTCAGCTCCCTGCTCCAGAACCTGGCCGTGTCTGTGACACCTATCCAGCGGATCGTGAGAAGCGGCGAGAATGCCCTGAAGATCGACATCCACGTGATCATCCCATACGAGGGCCTGTCCGCCGATCAGATGGCCCAGATCGAGGAGGTGTTCAAGGTGGTGTACCCAGTGGACGATCACCACTTCAAAGTGATCCTGCCCTATGGCACCCTGGTCATCGACGGAGTGACCCCAAACATGCTGAATTACTTCGGCAGGCCTTATGAGGGCATCGCCGTGTTTGATGGCAAGAAGATCACCGTGACAGGCACCCTGTGGAACGGCAATAAGATCATCGACGAGCGGCTGATCACCCCCGATGGCTCTATGCTGTTCAGAGTGACCATCAATAGCTTGA

GRK6-linker-LgBiT

ATGGAGCTCGAGAACATCGTAGCGAACACGGTGCTACTCAAGGCCCGGGAAGGTGGCGGTGGAAATCGCAAAGGCAAAAGCAAGAAATGGCGGCAGATGCTCCAGTTCCCTCACATCAGCCAGTGCGAAGAGCTGCGGCTCAGCCTCGAGCGTGACTATCACAGCCTGTGCGAGCGGCAGCCCATTGGGCGCCTGCTGTTCCGAGAGTTCTGTGCCACGAGGCCGGAGCTGAGCCGCTGCGTCGCCTTCCTGGATGGGGTGGCCGAGTATGAAGTGACCCCGGATGACAAGCGGAAGGCATGTGGGCGGCAGCTAACGCAGAATTTTCTGAGCCACACGGGTCCTGACCTCATCCCTGAGGTCCCCCGGCAGCTGGTGACGAACTGCACCCAGCGGCTGGAGCAGGGTCCCTGCAAAGACCTTTTCCAGGAACTCACCCGGCTGACCCACGAGTACCTGAGCGTGGCCCCTTTTGCCGACTACCTCGACAGCATCTACTTCAACCGTTTCCTGCAGTGGAAGTGGCTGGAAAGGCAGCCAGTGACCAAAAACACCTTCAGGCAATACCGAGTCCTGGGCAAAGGTGGCTTTGGGGAGGTGTGCGCCTGCCAGGTGCGGGCCACAGGTAAGATGTATGCCTGCAAGAAGCTAGAGAAAAAGCGGATCAAGAAGCGGAAAGGGGAGGCCATGGCGCTGAACGAGAAGCAGATCCTGGAGAAAGTGAACAGTAGGTTTGTAGTGAGCTTGGCCTACGCCTATGAGACCAAGGACGCGCTGTGCCTGGTGCTGACACTGATGAACGGGGGCGACCTCAAGTTCCACATCTACCACATGGGCCAGGCTGGCTTCCCCGAAGCGCGGGCCGTCTTCTACGCCGCCGAGATCTGCTGTGGCCTGGAGGACCTGCACCGGGAGCGCATCGTGTACAGGGACCTGAAGCCCGAGAACATCTTGCTGGATGACCACGGCCACATCCGCATCTCTGACCTGGGACTAGCTGTGCATGTGCCCGAGGGCCAGACCATCAAAGGGCGTGTGGGCACCGTGGGTTACATGGCTCCGGAGGTGGTGAAGAATGAACGGTACACGTTCAGCCCTGACTGGTGGGCGCTCGGCTGCCTCCTGTACGAGATGATCGCAGGCCAGTCGCCCTTCCAGCAGAGGAAGAAGAAGATCAAGCGGGAGGAGGTGGAGCGGCTGGTGAAGGAGGTCCCCGAGGAGTATTCCGAGCGCTTTTCCCCGCAGGCCCGCTCACTTTGCTCACAGCTCCTCTGCAAGGACCCTGCCGAACGCCTGGGGTGTCGTGGGGGCAGTGCCCGCGAGGTGAAGGAGCACCCCCTCTTTAAGAAGCTGAACTTCAAGCGGCTGGGAGCTGGCATGCTGGAGCCGCCGTTCAAGCCTGACCCCCAGGCCATTTACTGCAAGGATGTTCTGGACATTGAACAGTTCTCTACGGTCAAGGGCGTGGAGCTGGAGCCTACCGACCAGGACTTCTACCAGAAGTTTGCCACAGGCAGTGTGCCCATCCCCTGGCAGAACGAGATGGTGGAGACCGAGTGCTTCCAAGAGCTGAATGTCTTTGGGCTGGATGGCTCAGTTCCCCCAGACCTGGACTGGAAGGGCCAGCCACCTGCACCTCCTAAAAAGGGACTGCTGCAGAGACTCTTCAGTCGCCAAAGGATTGCTGTGGAAACTGCAGCGACAGCGAGGAAGAGCTCCCCACCCGCCTCTAGCCCCCAGCCCGAGGCCCCCACCAGCAGTTGGCGGGGGGGATCTGGGGGGGGGGGCTCTGGAGGCTCTTCATCAGGGGGAGTGTTTACTCTGGAGGACTTCGTCGGAGACTGGGAACAGACTGCTGCTTACAATCTGGATCAGGTGCTGGAACAGGGGGGGGTCAGCTCCCTGCTCCAGAACCTGGCCGTGTCTGTGACACCTATCCAGCGGATCGTGAGAAGCGGCGAGAATGCCCTGAAGATCGACATCCACGTGATCATCCCATACGAGGGCCTGTCCGCCGATCAGATGGCCCAGATCGAGGAGGTGTTCAAGGTGGTGTACCCAGTGGACGATCACCACTTCAAAGTGATCCTGCCCTATGGCACCCTGGTCATCGACGGAGTGACCCCAAACATGCTGAATTACTTCGGCAGGCCTTATGAGGGCATCGCCGTGTTTGATGGCAAGAAGATCACCGTGACAGGCACCCTGTGGAACGGCAATAAGATCATCGACGAGCGGCTGATCACCCCCGATGGCTCTATGCTGTTCAGAGTGACCATCAATAGCTTGA

GRK6 (K215M)-linker-LgBiT

ATGGAGCTCGAGAACATCGTAGCGAACACGGTGCTACTCAAGGCCCGGGAAGGTGGCGGTGGAAATCGCAAAGGCAAAAGCAAGAAATGGCGGCAGATGCTCCAGTTCCCTCACATCAGCCAGTGCGAAGAGCTGCGGCTCAGCCTCGAGCGTGACTATCACAGCCTGTGCGAGCGGCAGCCCATTGGGCGCCTGCTGTTCCGAGAGTTCTGTGCCACGAGGCCGGAGCTGAGCCGCTGCGTCGCCTTCCTGGATGGGGTGGCCGAGTATGAAGTGACCCCGGATGACAAGCGGAAGGCATGTGGGCGGCAGCTAACGCAGAATTTTCTGAGCCACACGGGTCCTGACCTCATCCCTGAGGTCCCCCGGCAGCTGGTGACGAACTGCACCCAGCGGCTGGAGCAGGGTCCCTGCAAAGACCTTTTCCAGGAACTCACCCGGCTGACCCACGAGTACCTGAGCGTGGCCCCTTTTGCCGACTACCTCGACAGCATCTACTTCAACCGTTTCCTGCAGTGGAAGTGGCTGGAAAGGCAGCCAGTGACCAAAAACACCTTCAGGCAATACCGAGTCCTGGGCAAAGGTGGCTTTGGGGAGGTGTGCGCCTGCCAGGTGCGGGCCACAGGTAAGATGTATGCCTGCATGAAGCTAGAGAAAAAGCGGATCAAGAAGCGGAAAGGGGAGGCCATGGCGCTGAACGAGAAGCAGATCCTGGAGAAAGTGAACAGTAGGTTTGTAGTGAGCTTGGCCTACGCCTATGAGACCAAGGACGCGCTGTGCCTGGTGCTGACACTGATGAACGGGGGCGACCTCAAGTTCCACATCTACCACATGGGCCAGGCTGGCTTCCCCGAAGCGCGGGCCGTCTTCTACGCCGCCGAGATCTGCTGTGGCCTGGAGGACCTGCACCGGGAGCGCATCGTGTACAGGGACCTGAAGCCCGAGAACATCTTGCTGGATGACCACGGCCACATCCGCATCTCTGACCTGGGACTAGCTGTGCATGTGCCCGAGGGCCAGACCATCAAAGGGCGTGTGGGCACCGTGGGTTACATGGCTCCGGAGGTGGTGAAGAATGAACGGTACACGTTCAGCCCTGACTGGTGGGCGCTCGGCTGCCTCCTGTACGAGATGATCGCAGGCCAGTCGCCCTTCCAGCAGAGGAAGAAGAAGATCAAGCGGGAGGAGGTGGAGCGGCTGGTGAAGGAGGTCCCCGAGGAGTATTCCGAGCGCTTTTCCCCGCAGGCCCGCTCACTTTGCTCACAGCTCCTCTGCAAGGACCCTGCCGAACGCCTGGGGTGTCGTGGGGGCAGTGCCCGCGAGGTGAAGGAGCACCCCCTCTTTAAGAAGCTGAACTTCAAGCGGCTGGGAGCTGGCATGCTGGAGCCGCCGTTCAAGCCTGACCCCCAGGCCATTTACTGCAAGGATGTTCTGGACATTGAACAGTTCTCTACGGTCAAGGGCGTGGAGCTGGAGCCTACCGACCAGGACTTCTACCAGAAGTTTGCCACAGGCAGTGTGCCCATCCCCTGGCAGAACGAGATGGTGGAGACCGAGTGCTTCCAAGAGCTGAATGTCTTTGGGCTGGATGGCTCAGTTCCCCCAGACCTGGACTGGAAGGGCCAGCCACCTGCACCTCCTAAAAAGGGACTGCTGCAGAGACTCTTCAGTCGCCAAAGGATTGCTGTGGAAACTGCAGCGACAGCGAGGAAGAGCTCCCCACCCGCCTCTAGCCCCCAGCCCGAGGCCCCCACCAGCAGTTGGCGGGGGGGATCTGGGGGGGGGGGCTCTGGAGGCTCTTCATCAGGGGGAGTGTTTACTCTGGAGGACTTCGTCGGAGACTGGGAACAGACTGCTGCTTACAATCTGGATCAGGTGCTGGAACAGGGGGGGGTCAGCTCCCTGCTCCAGAACCTGGCCGTGTCTGTGACACCTATCCAGCGGATCGTGAGAAGCGGCGAGAATGCCCTGAAGATCGACATCCACGTGATCATCCCATACGAGGGCCTGTCCGCCGATCAGATGGCCCAGATCGAGGAGGTGTTCAAGGTGGTGTACCCAGTGGACGATCACCACTTCAAAGTGATCCTGCCCTATGGCACCCTGGTCATCGACGGAGTGACCCCAAACATGCTGAATTACTTCGGCAGGCCTTATGAGGGCATCGCCGTGTTTGATGGCAAGAAGATCACCGTGACAGGCACCCTGTGGAACGGCAATAAGATCATCGACGAGCGGCTGATCACCCCCGATGGCTCTATGCTGTTCAGAGTGACCATCAATAGCTTGA

SRC-linker-LgBiT

atgggtagcaacaagagcaagcccaaggatgccagccagcggcgccgcagcctggagcccgccgagaacgtgcacggcgctggcgggggcgctttccccgcctcgcagacccccagcaagccagcctcggccgacggccaccgcggccccagcgcggccttcgcccccgcggccgccgagcccaagctgttcggaggcttcaactcctcggacaccgtcacctccccgcagagggcgggcccgctggccggtggagtgaccacctttgtggccctctatgactatgagtctaggacggagacagacctgtccttcaagaaaggcgagcggctccagattgtcaacaacacagagggagactggtggctggcccactcgctcagcacaggacagacaggctacatccccagcaactacgtggcgccctccgactccatccaggctgaggagtggtattttggcaagatcaccagacgggagtcagagcggttactgctcaatgcagagaacccgagagggaccttcctcgtgcgagaaagtgagaccacgaaaggtgcctactgcctctcagtgtctgacttcgacaacgccaagggcctcaacgtgaagcactacaagatccgcaagctggacagcggcggcttctacatcacctcccgcacccagttcaacagcctgcagcagctggtggcctactactccaaacacgccgatggcctgtgccaccgcctcaccaccgtgtgccccacgtccaagccgcagactcagggcctggccaaggatgcctgggagatccctcgggagtcgctgcggctggaggtcaagctgggccagggctgctttggcgaggtgtggatggggacctggaacggtaccaccagggtggccatcaaaaccctgaagcctggcacgatgtctccagaggccttcctgcaggaggcccaggtcatgaagaagctgaggcatgagaagctggtgcagttgtatgctgtggtttcagaggagcccatttacatcgtcacggagtacatgagcaaggggagtttgctggactttctcaagggggagacaggcaagtacctgcggctgcctcagctggtggacatggctgctcagatcgcctcaggcatggcgtacgtggagcggatgaactacgtccaccgggaccttcgtgcagccaacatcctggtgggagagaacctggtgtgcaaagtggccgactttgggctggctcggctcattgaagacaatgagtacacggcgcggcaaggtgccaaattccccatcaagtggacggctccagaagctgccctctatggccgcttcaccatcaagtcggacgtgtggtccttcgggatcctgctgactgagctcaccacaaagggacgggtgccctaccctgggatggtgaaccgcgaggtgctggaccaggtggagcggggctaccggatgccctgcccgccggagtgtcccgagtccctgcacgacctcatgtgccagtgctggcggaaggagcctgaggagcggcccaccttcgagtacctgcaggccttcctggaggactacttcacgtccaccgagccccagtaccagcccggggagaacctcgggGGATCTGGGGGGGGGGGCTCTGGAGGCTCTTCATCAGGGGGAGTGTTTACTCTGGAGGACTTCGTCGGAGACTGGGAACAGACTGCTGCTTACAATCTGGATCAGGTGCTGGAACAGGGGGGGGTCAGCTCCCTGCTCCAGAACCTGGCCGTGTCTGTGACACCTATCCAGCGGATCGTGAGAAGCGGCGAGAATGCCCTGAAGATCGACATCCACGTGATCATCCCATACGAGGGCCTGTCCGCCGATCAGATGGCCCAGATCGAGGAGGTGTTCAAGGTGGTGTACCCAGTGGACGATCACCACTTCAAAGTGATCCTGCCCTATGGCACCCTGGTCATCGACGGAGTGACCCCAAACATGCTGAATTACTTCGGCAGGCCTTATGAGGGCATCGCCGTGTTTGATGGCAAGAAGATCACCGTGACAGGCACCCTGTGGAACGGCAATAAGATCATCGACGAGCGGCTGATCACCCCCGATGGCTCTATGCTGTTCAGAGTGACCATCAATAGCtga
